# Exploring the Influence of Chemical Exposures in Breast Cancer Disparities: High-Throughput Transcriptomic Analysis in Normal Breast Cells from Diverse Donors

**DOI:** 10.64898/2026.02.23.707203

**Authors:** Neil Zhao, Peiyao Zhao, Anagha Tapaswi, Katelyn Polemi, Tasha Thong, Jonathan Sexton, Simone Charles, Max S. Wicha, Laurie Svoboda, Xiang Zhou, Justin A. Colacino

**Author notes:** These authors contributed equally (Neil Zhao, Peiyao Zhao).

## Abstract

Racial disparities in the incidence of, and mortality from, aggressive breast cancers are a pressing public health issue. Many factors have been investigated in these inequities; however, the role of toxicant exposures is not well characterized. We and others have identified substantial inequities in chemical biomarker concentrations by race. The goal of this study was to test the hypothesis that exposure to these chemicals is linked to biological changes relevant to aggressive breast cancers, such as dysregulation of the Hallmarks of Cancer. We used high throughput transcriptomic profiling of normal primary human breast epithelial cells from diverse donors (*n=6*) to test effects of 8 chemicals (cadmium, lead, arsenic, copper, PFNA, BPA, BPS, p,p’-DDE) with documented exposure disparities by race/ethnicity across 3 concentrations (100nM, 1µM, 10µM). Across chemicals, we identified that pathways related to cell cycle regulation and protein secretion were commonly affected. Through bioinformatic estimation of cell type proportions, we found that metals like lead and cadmium induced cell-type shifts, consistent with the dysregulated cellular plasticity cancer hallmark. Lead and arsenic response genes were enriched for genes associated with poor breast cancer survival in the Cancer Genome Atlas. Integrating concentration-response modeling and chemical biomonitoring data, BPA, p,p’-DDE, copper, and lead elicited expression changes at concentrations relevant to the US population. Finally, we identified substantial interindividual heterogeneity in response to organic compounds, but less so in metals. These findings highlight the value of high-throughput transcriptomics as a New Approach Methodology (NAM) in quantifying how common exposures may impact aggressive breast cancer associated biological processes.

## Introduction

Breast cancer disparities by race are a pressing issue in public health. In the United States, women of African ancestry face a 40% higher likelihood of dying from breast cancer than other racial and ethnic groups.^1,2^ Of note, triple-negative breast cancer (TNBC), a heterogeneous and highly aggressive subtype of cancer, has the worst overall survival rates and is 2-3 times more likely to be diagnosed in African American than European American women.^3,4^ These aggressive breast cancers are characterized by the acquisition of common “Hallmark” traits, including replicative immortality, activation of invasion & metastatic pathways, evasion of growth suppressors, and phenotypic plasticity.^5^ However, the etiological drivers of these biological capabilities underlying racial disparities are multifactorial and complex. Although epidemiological studies have examined links to ancestry, genetics, diet, exercise, reproductive factors, and social determinants of health, the impacts of chemical exposures are less understood. Growing evidence suggests environmental chemicals encountered at human-relevant doses promote breast cancer-associated molecular and functional bioactivities.^6^ Recent paradigm shifts also highlight the parallels between cancer and stem cells through the existence of hybrid cell-state transitions—the ability of cells to express a mixed gradient of epithelial-to-mesenchymal and luminal-to-basal characteristics to adapt and respond to environmental signals and stressors.^7–10^ Within invasive breast cancers, studies have documented luminal-to-basal-like hybrid signatures and “stem-like” epigenetic configurations that display dysregulated developmental trajectories, lineage infidelities, self-renewal capacity, and increased metastatic potential.^11,12^

Furthermore, population-level biomonitoring efforts, such as the US National Health and Nutrition Examination Survey (NHANES), have shown that African American and low-income women are disparately exposed to a wide range of chemicals, including toxicants such as heavy metals, pesticides, PFAS compounds, and endocrine-disrupting chemicals.^13,14^ For instance, previous work demonstrated the ability of bisphenols to elicit non-monotonic dose-response of breast cancer stem-cell signatures through both estrogenic and non-canonical signaling pathways.^15–18^ Recent experimental data also implicates a wide range of chemicals, including cadmium, arsenic, lead, copper, the DDT metabolite dichlorodiphenyldichloroethylene (p,p’-DDE), and perfluorononanoic acid (PFNA), in their ability to promote carcinogenesis within mammary epithelial cells through distinct mechanisms across rodents, non-tumorigenic MCF10A *in vitro* cultures, and bioactivity data from the EPA’s ToxCast Program.^6,14,19^ Although animal and cell line models are commonly utilized to assess the risks of environmental toxicants, they are inadequate in representing the genetic diversity, physiology, and inter-individual differences found in human primary tissues. Given the disproportionate exposures of African American women to environmental toxicants, increasing number of data-poor chemicals in circulation, and the need to address breast cancer health disparities, the goal of this project is to better understand how chemical exposures impact personalized breast cancer risk and elucidate mechanisms of action. To achieve this, we expanded testing of prioritized chemicals using high-throughput transcriptomic characterization on conditionally reprogrammed normal mammary cells from genetically diverse individuals. Chemical concentrations, spanning 100 nM to 10 μM, were selected based on prior benchmark concentration modeling from our group that integrated *in-vitro* transcriptomic responses with biomarker concentrations from population-level exposure data (NHANES), establishing biologically active concentration ranges that correspond to human-relevant internal doses.^6,14^ Our overarching hypothesis is that exposure disparity chemicals promote aggressive breast cancer-associated biological effects linked to poor clinical outcomes at human-relevant doses through distinct and overlapping mechanisms.

## Materials and Methods

### Sample Selection

Normal human mammary samples were obtained from the Susan G. Komen normal tissue bank and established and expanded into conditional reprogramming (CR) cultures according to the protocol of Liu et al. (*2017*) and Thong et al. (*2020*).^20,21^ All samples were nulliparous and self-identified as African American (AA) or European American (EA). 6 sample cell lines (*n*=3 AA, *n*=3 EA) matched by age, BMI, and days since last menstrual period were selected (*see Supplementary Table S-1*).

### Compound Preparation

The 8 chemicals were purchased from Sigma-Aldrich (Saint Louis, MO, USA) and Cayman Chemical (Ann Arbor, MI, USA) and processed according to the protocol of Sala-Hamrick et al. (2024) (*see Supplementary Table S-2*).^6^ In brief, the chemicals were weighed and dissolved in either Dimethyl sulfoxide (DMSO) (BPA, BPS, p,p’-DDE, PFNA) or Sterile Water (sodium arsenite, cadmium chloride, copper chloride, lead acetate) at a concentration of 5mg/mL and stored at −20^°^C for long term storage. An intermediate stock concentration of 5mM was prepared for each of the chemicals, which was further diluted into final concentrations for dosing the CR cells: 0.1 µM, 1 µM, 10 µM. These final concentrations were prepared fresh in CR culture F-media for each experiment. For vehicle controls and chemicals dissolved in DMSO, the final DMSO concentration was 0.5%. For chemicals dissolved in water, an equivalent amount of water was included in the vehicle control.

### Cell Culture and Plating

Conditionally reprogrammed cells from each sample (KCR7889, KCR7953, KCR7518, KCR8195, KCR8580, KCR8519) were thawed from cryopreservation, cultured individually in CR culture, and expanded at passage 2 for all rounds of cell culture according to the protocol of Schroeder et al (*2024*), where this same exposure paradigm was performed and impacts on cell number and cellular state were assessed via high content imaging.^22^ Once confluence was reached at ∼70%, cells were differentially trypsinized (Gibco 0.05% Trypsin/EDTA), counted, assessed for viability using acridine orange/propidium iodide staining (LUNA FL Dual Fluorescence Cell Counter), and plated in black, clear flat bottom 384 well plates (Corning, Corning, NY, USA) using a 2.5-125uL multi-channel pipette according to the protocol of Sala-Hamrick et al. and Shroeder et al. Cells were then exposed to test chemicals for 48 hours with 3 biological triplicates per dose (0.1, 1, 10µM). For vehicle controls (*n_DMSO_ = 6*, *n_water_ = 6*), cells from each individual were exposed, yielding 6 biological replicates for 48 hours to control across all doses of water- or DMSO-dissolved chemicals (*see Supplementary Table S-3*). Experiments were run in 2 batches to include experimental replicates for all individual cell line-by-chemical-by-dose combinations.

### RNA-Sequencing Overview

Samples were processed using an adapted version of the plexWell (SeqWell, Beverly, MA, USA) plate-based sequencing method according to the protocol of Sala-Hamrick et al. (*2024*).^6^ Briefly, sequencing reactions are prepared in custom 96 well plates, with each well containing a unique oligo barcode. Cells are lysed and cDNA prepared *in situ* directly from the cellular lysate. A tagmentation reaction adds the well specific oligo barcode to the cDNA, which provides the sample from each well with a unique combination of i5 and i7 indices. Next, cDNA from the multiple wells is pooled for RNA-seq library preparation via the plexWell 384 Rapid Single Cell RNA Library Prep Kit. Library QC was performed on the Agilent Bioanalyzer (High Sensitivity DNA 5000 kit) prior to paired-end sequencing on an Illumina NovaSeq 6000 at the University of Michigan Advanced Genomics Core.

### RNA-Sequencing Data Processing

Sequencing FASTQ reads were first demultiplexed back to individual samples based on the i5 and i7 indices. Sequencing reads were transferred to the University of Michigan Great Lakes high performance computing cluster for further analysis. We assessed sequencing read quality via *fastQC and multiQC*. Because the conditional reprogramming process cultures human cells on a feeder layer of irradiated mouse embryonic fibroblasts, we aligned the reads to a splice junction aware build of a combined human-mouse genome (hg19-mm10) using STAR. Aligned reads were assigned to genes using *featureCounts*, excluding reads which multimapped and multi-overlapped. Counts which aligned to the mouse genome were discarded for downstream analyses, leaving only counts aligning to the human genome.

### QC Differential Gene Expression Testing

Read count matrices from featureCounts were imported into R/Bioconductor (version 4.4.3) package edgeR (version 4.2.2), and samples with fewer than 5,000,000 total mapped reads (library size) were excluded from downstream analysis. Samples were further filtered for excess experimental replicates to reflect the standard group size and were selected based on the highest library size. Trimmed mean of M values (TMM) normalization factors was calculated to adjust for sample composition biases in library size. Genes with low expression were excluded from further analysis using the edgeR *filterByExpr*() function with default settings. The TMM size factors and filtered count matrices were imported into the R/Bioconductor package limma (version 3.60.6). Multidimensional scaling (MDS) was performed on the normalized log-transformed counts per million (cpm) data using the top 500 most variable genes to identify the top principal coordinates and any confounding batch effects. After undergoing QC and filtering, a total of 20,389 genes passed the selection criteria and were used for downstream bulk differential expression analyses.

The *voom()* function was applied to estimate the mean-variance relationship of the data and to assign appropriate weights to each observation. Replicates were treated as random effects, and *duplicateCorrelation()* was used to account for this random effect. Linear models were fitted to the data, and empirical Bayes moderation of the t-statistics was applied using the *eBayes()* function, which shrinks the gene- and sample-specific error variances towards a pooled global estimate for more robust inference. To adjust for potential biases introduced by the small sample size, *n=*50 permutation replicates were performed for each combination. Adjusted p-values were calculated as the proportion of permuted statistics greater than or equal to the observed statistic. Multiple testing correction was then applied across all genes using the Benjamini–Hochberg (BH) procedure, and genes with an adjusted p-value less than 0.1 were considered significantly differentially expressed between the treatment and control groups.

### Functional Enrichment and Gene Ontology Analysis

For each chemical, differentially expressed (DE) genes that were consistently upregulated or downregulated across all three doses were selected for gene set enrichment analysis (GSEA) (*see supplementary Table S-7*). The *gost()* function from the gprofiler2 package (version 0.2.3) was used to perform enrichment analysis across multiple gene set collections with the parameter organism = “gp__7ie1_G7XX_XJw” for the MSigDB Hallmark (H) collection.

### Benchmark Dose Analyses

Best fit benchmark dose-response models and gene- and pathway-level benchmark doses were identified using BMDExpress (version 3.20.141), a free software package developed by the National Institute for Environmental Health Sciences (NIEHS) and available for download (https://github.com/auerbachs/BMDExpress-3/releases). Best practices for dose-response modeling for each chemical were conducted according to BMDExpress online documentation (https://github.com/auerbachs/BMDExpress-3/wiki#basic-workflow). Normalized log2-transformed cpm reads from the RNA-seq data were imported into BMDExpress and pre-filtered using One-Way ANOVA for significance at a Benjamini-Hochberg FDR adjusted p-value ≤ 0.05 to identify genes showing significantly increasing or decreasing concentration responses. To determine concentration-response relationships, the filtered data were modeled in BMDExpress EPA Benchmark Dose Software (BMDS) with hill, linear, power, exponential (3, 5), and polynomial (2°, 3°, 4°) models, and the best fit model was chosen at a confidence level of 0.95. Default model settings were selected for EPA BMDS. The benchmark response (BMR) was set to 1 standard deviation (1.021, 5%) with non-constant variance relative to control response and “Profile Likelihood” as the BMDU/L Estimation Method (upper/lower bound of the BMD estimate). Best fit models were chosen for each dose-response relationship per gene by computing the estimated BMD, BMDU, and BMDL and using nested chi square followed by lowest Akaike information criterion (AIC) to select for the best model. Hill models were flagged if its ‘k’ parameter was 1/3 less than the lowest positive dose and flagged hill models were replaced with the next best model that met both the minimum AIC value and a goodness-of-fit p-value > 0.05. Benchmark concentrations with values above the highest assayed dose (10 µM) were excluded from further analysis.

### Cell Type Composition Prediction using Normal Breast Single Cell Atlases

To determine if the chemical treatments had an effect on the cellular composition of the samples, bioinformatic deconvolution of the bulk RNA-seq data was performed based on single cell-atlases of the normal human mammary gland from Pal et al. (*2021*) (*n=10*).^23^ Data from Pal et al. (*2021*) was pre-processed according to the protocol from Sala-Hamrick et al. (*2024*)^6^ and includes cell cluster annotations labeled as “Luminal Progenitor” and “Myoepithelial” based on marker gene expression signatures. This annotated single-cell gene expression dataset was then used to estimate cell type proportions in the bulk RNA-seq data using the Multi-subject Single Cell (MuSiC) deconvolution method.^24^ Statistical significance for multiple comparisons was assessed using Dunnett’s post-hoc test (α ≤ 0.05), with the vehicle control group (0 µM) serving as the reference. Dose-dependent changes in predicted cell-type proportions were then modelled for each chemical individually with multivariate generalized linear mixed-effects (GLME) models, treating dose as a fixed effect, the matched control (DMSO or water) as the reference level, and including donor-specific random intercepts, coefficients, and slopes to account for inter-individual variability.

### Cell Viability Assessment and Cytotoxicity Analysis

Cell viability data were obtained from our parallel cytotoxicity study, conducted by Schroeder et al. (*2024*), using identical experimental plates, culture conditions, and chemical exposure protocols as the transcriptomic analysis.^22^ Briefly, CR cultures from the same six donors (KCR7518, KCR7889, KCR7953, KCR8195, KCR8519, KCR8580) were exposed to the same eight chemicals at matching doses (0.1, 1, and 10 µM) in HTTr 48-well plates. Following 48-hour exposure, cells were fixed and stained with DAPI to quantify nuclei counts using automated high-content imaging. Cell viability for each treatment condition was calculated as the percentage of nuclei counts relative to vehicle controls, with water serving as the control for inorganic chemicals (cadmium chloride, copper chloride, lead acetate, sodium arsenite) and DMSO for organic chemicals (BPA, BPS, DDE, PFNA). Control values were averaged across technical replicates for each donor to establish baseline cell counts, and treatment replicates were normalized to these donor-specific control means.

To assess whether overt cytotoxicity contributed to observed transcriptomic changes, we performed correlation analyses between relative viability and the number of differentially expressed genes (DEGs) identified for each treatment condition. Technical replicates for viability measurements were averaged to match the structure of RNA-sequencing data. Pearson correlation coefficients were calculated for the overall dataset (*n = 144* treatment conditions) as well as stratified by dose (0.1, 1, and 10 µM). Chemical- and donor-specific correlations were computed using individual technical replicate measurements to assess whether viability-DEG relationships varied systematically across chemical classes or donor backgrounds. Statistical significance was determined using two-tailed t-tests.

To quantify the relative contribution of viability versus chemical identity and dose to transcriptomic variance, we performed analysis of covariance (ANCOVA) with DEG count as the dependent variable and relative viability, chemical, and dose as independent variables. Variance partitioning was conducted by calculating the proportion of total sum of squares explained by each factor. We also compared mean viability between high-DEG conditions (DEG count > median) and low-DEG conditions (DEG count ≤ median) using unpaired two-sample t-tests. All statistical analyses were performed in R (version 4.4.3) while using the tidyverse (version 2.0.0) suite of packages.

### Contextualizing Benchmark Doses to Human Population Exposure Levels

To understand whether cancer-associated gene and pathway effects due to toxicant exposures occur at population relevant levels, BMD estimates of assayed chemicals were compared to NHANES human biomarker concentrations measured in the blood, urine, or serum of US women according to the methodology from Nguyen et al. (*2021*) and Sala-Hamrick et al. (*2024*).^6,13^ This curated NHANES dataset contains chemical biomarker measurements of the 8 assessed chemicals in up to 57786 women recruited from 1999-2018.^25^ Briefly, human biomarker concentrations were converted to molarity units and then compared to the transcriptional BMD concentrations identified through benchmark dose analyses using boxplots. Metadata between chemicals assayed *in vitro* and their corresponding exposure biomarkers in NHANES are shown in *Supplementary Table S-4*.

### Comparison of Transcriptomic Differences between Komen Dataset and TCGA BRCA Cohorts

To compare our dataset with publicly available molecular data from breast cancer patients, we examined studies conducted by The Cancer Genome Atlas (TCGA), with a specific focus on Breast Invasive Carcinoma (BRCA) RNA-seq data and its relationship with patient outcomes. Cox regression analysis results, assessing the association between survival outcomes and mRNA expression, were obtained from OncoLnc.^26^ Genes with a Benjamini-Hochberg adjusted p-value smaller than or equal to 0.2 were selected for further investigation. Fisher’s exact test was conducted to assess the significant association of the detected DEGs between the Komen dataset and the TCGA-BRCA dataset.

### Data Availability

RNA-seq data have been submitted to the gene expression omnibus (GSE308852). Original expression data, ANOVA filtered results, and subsequent BMDExpress dose-response modeling analyses are included in a .bm2 file available in supplementary information.

## Results

### Transcriptomes Cluster by Individual, Highlighting Inter-Individual Variability in Global Expression Profiles

To investigate whether exposure disparity chemicals induce distinct transcriptomic responses in normal mammary cells from diverse individuals, we tested three human-relevant doses of eight chemicals using HTTr plate-based bulk RNA-sequencing on six CR cultures (**Table S-1**). After sequencing and QC, we retained 20,389 expressed genes in each experimental condition for downstream analyses. Multidimensional scaling (MDS) analysis of the 500 most variable genes was conducted to assess how inter-individual differences and chemical exposures shape the global transcriptional landscape. Principal coordinate analysis of log-normalized CPM gene counts revealed that donor identity exerted a dominant influence on transcriptomic variation, with samples clustering primarily by individual rather than treatment or dose (**Figure 1A**). KCR7953 was the most transcriptionally distinct donor, separating along the first principal coordinate (PC1), while the remaining donors formed two sub-groups: KCR7518, KCR8519, and KCR8580 clustering together, and KCR7889 grouping closely with KCR8195. Treatment- and dose-related effects were detectable along PC2 but were relatively modest compared to inter-individual variability, with considerable overlap among different exposure conditions.

**Figure 1:**
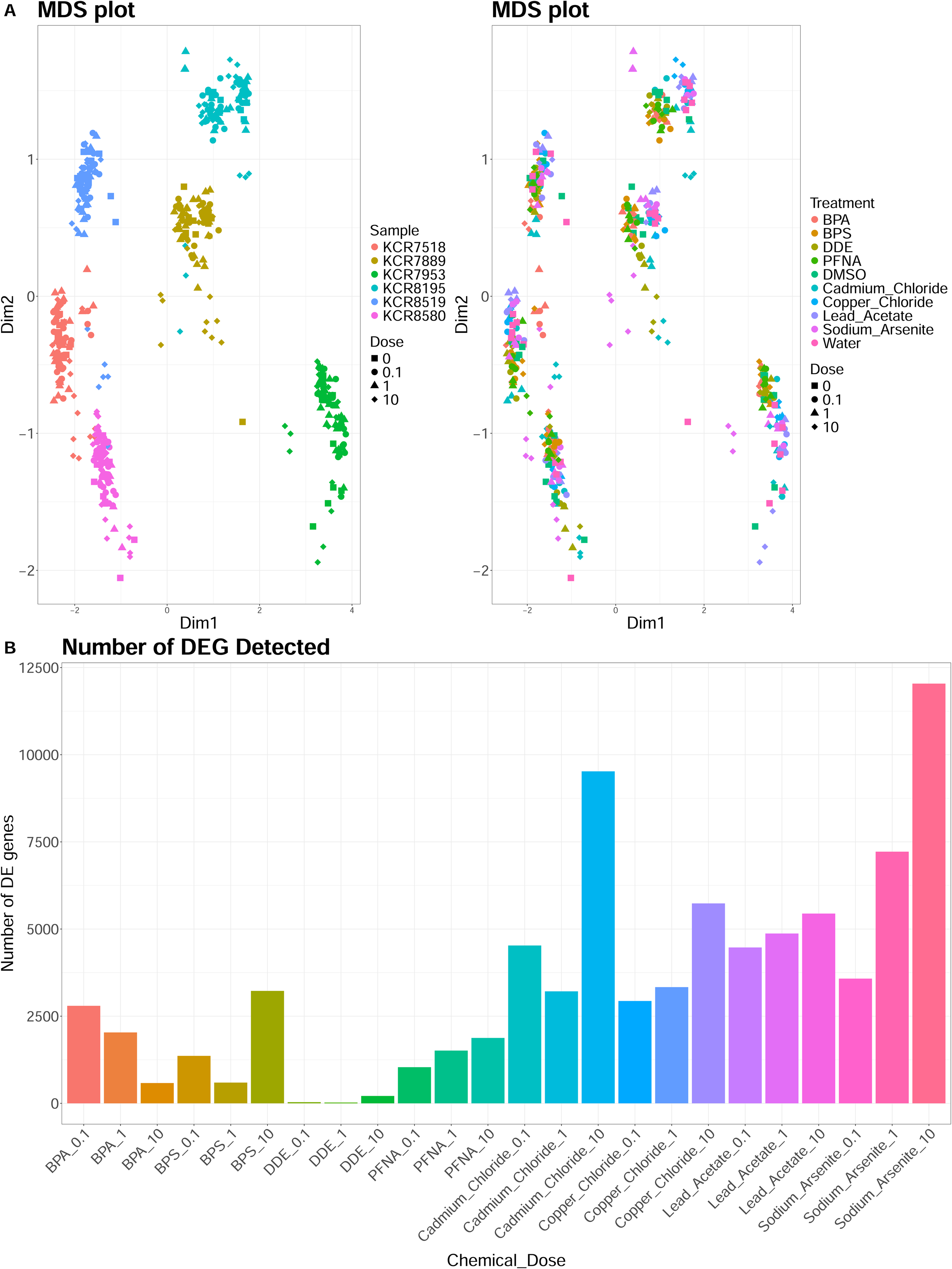
Multidimensional Scaling and Overall Differential Expression. **A.)** Multidimensional scaling plot (MDS) of depicting global transcriptomic variation among conditionally reprogrammed mammary cells (CR cultures). Left: Experimental samples colored by individual donor. Right: Experiment samples colored by treatment group. Each point represents the normalized expression profile of a single sample, with shape indicating dose (0, 0.1, 1, or 10 µM). The first two dimensions (Dim1 and Dim2) capture the greatest variance in the top 500 most variable genes. **B.)** Bar chart summarizing the total number of significantly differentially expressed genes (DEGs) by chemical–dose combination compared to vehicle control, as determined using a permutation-based approach.

To address potential confounding effects of overt cytotoxicity on transcriptomic profiles, we integrated cell counts measurements from a parallel matched design conducted by Schroeder et al. (*2024*) on the same experimental plates and culture conditions.^22^ Relative cell counts were calculated for each treatment condition by normalizing cell counts to respective vehicle controls and compared against the magnitude of transcriptomic response, quantified as the total number of differentially expressed genes (DEGs). Across all 144 treatment conditions spanning six donors, eight chemicals, and three doses, we observed a weak negative correlation between relative cell counts and DEG count (*r = -0.244*, *p = 0.0032*), with 97% of conditions maintaining ≥50% cell counts and 82% maintaining ≥80% relative to control (**Figure S-1A**). Dose-stratified analysis revealed this correlation strengthened only at the highest dose (10 µM: *r = -0.380*, *p = 0.0076*), while lower doses showed minimal relationship (0.1 µM: *r = 0.051, p = 0.73*; 1 µM: *r = -0.135, p = 0.36*) (**Figure S-1C-E**). Importantly, 97.2% of all treatment conditions (*n = 140/144*) maintained ≥50% viability relative to controls, well above the threshold where cytotoxicity-driven transcriptional changes become non-specific.^27^

Chemical- and donor-specific correlation analysis identified 13 significant viability-DEG correlations (*p < 0.05*) among 48 chemical × donor combinations (27.1%) (**Figure S-1B**). Notably, all significant correlations were negative and concentrated in specific chemical-donor pairs, particularly sodium arsenite (significant in all 6 donors). Three chemical-donor combinations could not be evaluated due to absence of DEGs across all doses (copper chloride in donors KCR7518 and KCR8519; BPS in KCR8519) (**Figure S-1B**; **Table S-5**). ANCOVA confirmed that chemical identity (17.7%) explained ∼3-fold more transcriptomic variance than viability (5.9%), with dose contributing 4.1%. Although high-DEG conditions showed slightly lower mean viability than low-DEG conditions (94.6% vs. 99.8%), this difference was not statistically significant (*p = 0.17*), and both groups maintained viability well above cytotoxicity thresholds.

Together, these findings demonstrate that observed transcriptomic variation reflect donor-and chemical-specific biological responses rather than non-specific cytotoxic artifacts. While inter-individual heterogeneity exerted the strongest influence on global transcriptomic variation, chemical exposures still introduced distinct variation, prompting further investigation into the dose-dependent nature of these effects.

### Differential Gene Expression Analysis Reveals Chemical- and Dose-Specific Effects

To further characterize and quantify treatment-specific transcriptomic alterations, we applied quality-weighted *limma-Voom* differential gene expression analysis to identify DEGs across chemical treatment doses relative to vehicle control for each donor and aggregated across all donors. The number, magnitude, and directionality of DEGs varied substantially by sample donor, chemical, and dose (**Figure 1B**; **Figure S-2**; **Figure S-3**).

Heavy metals induced the most robust transcriptional responses across all concentrations. At the donor level, the total number of DEGs across all sample-by-chemical dose combinations (**Table S-5**) varied widely from 0 (KCR8519 at 0.1 µM) to 15,201 (KCR7953 at 10 µM, 60.1% upregulated) following arsenic exposure; 0 (KCR7518 and KCR8519 at 0.1 µM) to 7,896 (KCR7953 at 10 µM, 58.4% downregulated) following cadmium exposure; 0 (KCR7889 at 0.1 µM) to 12,251 (KCR8195 at 10 µM, 59.9% downregulated) following copper exposure; and 0 (KCR7518, KCR7889 at 0.1 µM) to 10,063 (KCR8195 at 0.1 µM, 50.3% upregulated) following lead exposure. In contrast, organic compounds such as bisphenols (BPA, BPS) and PFNA induced fewer DEGs and consistently showed upregulation-dominant profiles across doses (**Table S-5**). The total number of DEGs ranged from 0 (KCR8519 at 0.1 µM) to 2,338 (KCR8580 at 10 µM, 71.5% upregulated) following BPA exposure; 0 (KCR8580 at 1 µM and KCR7518 at 10 µM, and KCR8519 at all doses) to 2,146 (KCR8580 at 10 µM, 71.7% upregulated) following BPS exposure; 0 (KCR7889 and KCR8519 at 0.1 µM, KCR7889 at 1 µM, and KCR8580 at 10 µM) to 5,593 (KCR7889 at 10 µM, 82.9% upregulated) following p,p’- DDE exposure; and 0 (KCR8519 at 0.1 µM) to 1,993 (KCR8580 at 0.1 µM, 68.2% upregulated) following PFNA exposure.

Among the exposure disparity chemicals, arsenic, lead, and BPS consistently elicited significant and substantial DEGs (adj. p-value ≤ 0.05, |log₂FC| ≥ 2) across all three doses in all six individuals (**Figure S-3**). Notably, the directionality of differential gene expression shifted in a dose-dependent manner, with lower doses predominantly showing downregulation while higher doses favored upregulation across donors. Arsenic induced the highest number of downregulated genes in KCR7953 at 0.1 µM (*n = 7,434*, 65.0% downregulated), while shifting towards a maximal upregulation at 10 µM (*n = 9,135*, 60.1% upregulated) (**Table S-5**). Similarly, BPS exposure led to the highest number of downregulated DEGs in KCR8580 at 0.1 µM (*n=563*, 27.4% downregulated), whereas the same donor exhibited maximal upregulated transcriptional effects at 10 µM (*n=1,538*, 71.7% upregulated). Interestingly, lead induced the highest number of downregulated genes in KCR8195 at 0.1 µM (*n=5,008*, 49.8% downregulated), with relatively balanced up- and downregulation across doses.

To examine the overall directional bias of transcriptional responses, we generated a heatmap of consistently upregulated and downregulated genes across all doses (**Figure 2A**; **Table S-6**). All chemicals displayed prominent fractions of genes consistently upregulated across all three doses, where lead acetate (98.82%) and copper chloride (95.97%) demonstrated the highest proportion of consistently upregulated genes. To a lesser extent, BPA (83.41%), BPS (88.85%), PFNA (89.22%) and sodium arsenite (89.97%) exhibited moderate proportions of consistently upregulated genes. In contrast, cadmium chloride (78.42%) and p,p′-DDE (75%) showed relatively smaller fractions, though still representing high proportions of consistently upregulated genes. Together, these observations illustrate substantial inter-individual variability in transcriptional response and differences in sensitivity across chemicals.

**Figure 2:**
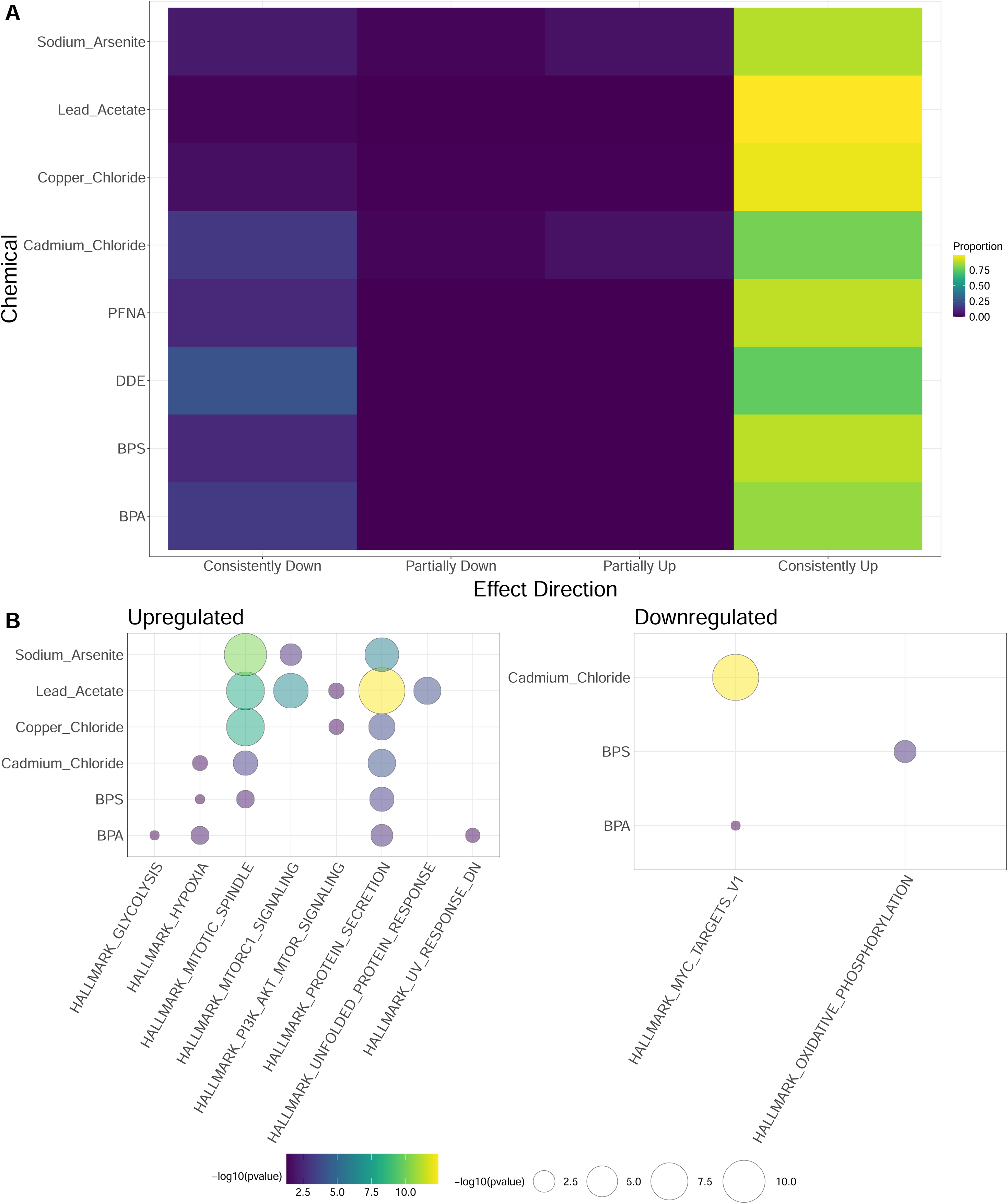
Directionality and Pathway-Level Effects of Exposure Disparity Chemicals. **(A)** Heatmap showing the fraction of DEGs that were (*left-to-right*) consistently downregulated, partially down, partially up, or consistently upregulated across all three doses of each chemical compared to vehicle control. Rows correspond to each chemical, and columns represent the effect direction category. The color scale ranges from purple (proportion = 0.00) to yellow (proportion ≥ 0.75). **(B)** Bubble plot showing the top MSigDB Hallmark (H) gene sets significantly enriched by genes consistently upregulated across all doses per chemical. Bubble size represents the relative magnitude of enrichment (e.g., proportion of upregulated DEGs), and bubble color indicates the overall significance or strength of the response ( -log10 p-value). Pathways on the x-axis include hallmark processes implicated in breast cancer biology. Rows represent chemicals tested.

### Benchmark Dose Modeling Reveals Chemical-Specific and Inter-Individual Variability in Transcriptional Sensitivity

To further evaluate the transcriptional sensitivity and dose-dependent gene expression changes induced by exposure disparity chemicals, best-fit benchmark dose (BMD) modeling was conducted on DEGs identified across individual donor samples. The median BMD for each gene was determined per chemical-treatment combination using best-fit dose-response models (*see supplementary .bm2 file*). The accumulation plots depict the cumulative number of DEGs as a function of increasing median BMD values, revealing distinct chemical-specific and inter-individual variability in transcriptional sensitivity (**Figure 3**).

**Figure 3.**
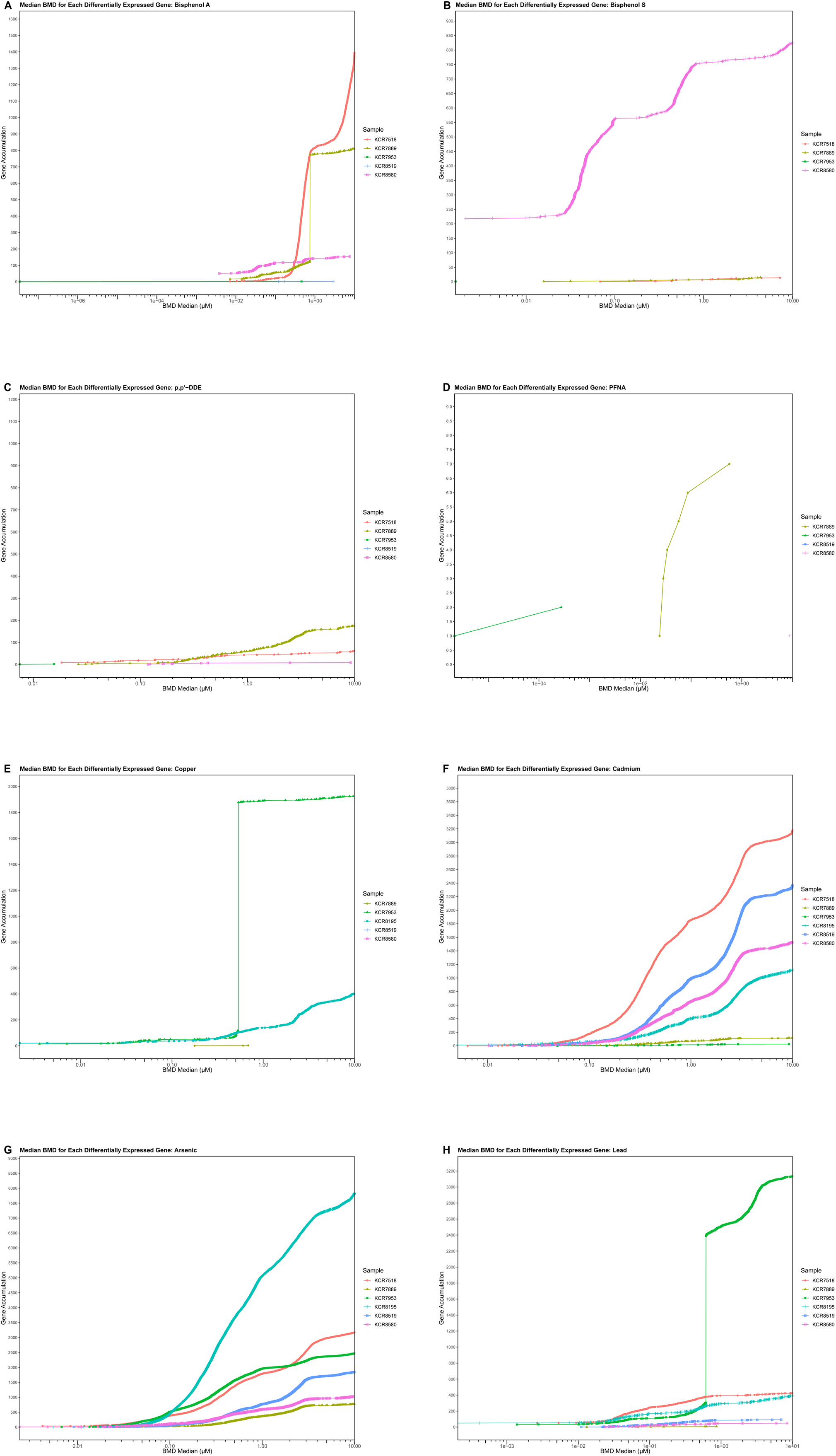
(A-H): Individual Benchmark Dose Response Modeling Graphs Using DEGs for Each Exposure Disparity Chemical. In the accumulation plot, each plotted point represents the number of DEGs that have a median BMD less than or equal to the corresponding value on the horizontal axis. Each colored line represents an individual donor sample. DEGs and/or samples are only shown for statistically significant dose responses filtered using a one-way ANOVA at an FDR adjusted p-value cutoff of ≤0.05.

Organic chemicals such as BPA (**Figure 3A**), BPS (**Figure 3B**), and p,p’-DDE (**Figure 3C**) displayed more variable and individual-specific responses. BPA and BPS induced sharp gene accumulation and maximal effects in KCR7518 and KCR8580, respectively, whereas other individuals exhibited weaker or minimal response. p,p’-DDE showed the most pronounced transcriptional effects in KCR7889 but had limited impact on other samples. In contrast, PFNA (**Figure 3D**) displayed minimal enrichment, with few dose-responsive genes across all individuals.

Heavy metals such as sodium arsenite, cadmium chloride, and lead acetate induced the most potent dose-responsive gene accumulation across all individuals, with sodium arsenite (**Figure 3G**) and cadmium chloride (**Figure 3F**) exhibiting the steepest increases in gene accumulation at ∼0.1 µM. Notably, KCR8195 and KCR7518 respectively demonstrated greater sensitivity and maximal response to sodium arsenite and cadmium chloride, accumulating a high number of DEGs at lower BMD values. Lead acetate (**Figure 3H**) displayed moderate effects, with KCR7518, KCR7953, and KCR8195 exhibiting greater sensitivity and maximal response, while other individuals showed more gradual gene accumulation and limited maximal response. In contrast, copper chloride (**Figure 3E**) induced a more restricted transcriptional response, with only a subset of individuals exhibiting significant dose-dependent changes (KCR7889, KCR7953). Collectively, these findings highlight the consistent potency of heavy metals in triggering widespread dose-responsive transcriptional effects, while organic compounds elicited more heterogeneous and individual-dependent effects.

### Gene Set Enrichment Analysis Identifies Breast Cancer Related Biological Processes and Breast Cancer Mortality Associated-Pathways Affected by Exposure Disparity Chemicals

To determine whether exposure disparity chemicals perturb biological pathways relevant to cancer and breast cancer mortality, we performed differential gene expression and gene set enrichment analysis (GSEA) (**Figure 2B**). Hallmark-level GSEA, performed on the subset of DEGs that were consistently upregulated or downregulated across all doses for each chemical, is summarized in *Supplementary Table S-7*. A core set of DEGs exhibited consistent enrichment (≥ 3 conditions, FDR-adj. *p ≤ 0.05*, |logFC|≥2) in cancer-related gene signatures. Despite spanning different protein-encoding gene families, differential expression converges on adaptive processes—exporting toxicants, reshaping metabolism, and activating pro-survival signaling.

Among the dose-specific DEGs, oxidative stress and detoxification pathways showed the most consistent activation across chemical-dose conditions. *HMOX1* (heme oxygenase 1), a critical cytoprotective enzyme upregulated in chemoresistant breast cancers,^28^ was induced across 5 conditions (cadmium 10, lead 1-10, arsenic 1-10 µM) (**Figure S-2**). Similarly, aldehyde dehydrogenases *ALDH3A1* (6 conditions: cadmium 1-10, lead 1-10, arsenic 1-10 µM) and *ALDH2* (6 conditions: BPS 10, PFNA 10, copper 1-10, lead 1-10, arsenic 10 µM), and aldo-keto reductases *AKR1C1/C2* (5 conditions: cadmium 10, lead 1-10, arsenic 1-10 µM) and *AKR1C3* (3 conditions: cadmium 10, lead 10, arsenic 10 µM), which have been implicated in chemotherapy resistance and hormone-dependent malignancies, demonstrated coordinated upregulation.^29,30^ Oxidative stress response genes, including glutathione biosynthesis enzymes *GCLM* and *GCLC* (4 conditions: cadmium 10, lead 1-10, arsenic 1-10 µM) and thioredoxin system genes *NQO1* (5 conditions: cadmium 10, lead 1-10, arsenic 1-10 µM) and *TXNRD1* (4 conditions: cadmium 10, lead 10, arsenic 1-10 µM), exhibited coordinated upregulation (**Figure S-2**). Metallothioneins, a family of metal-binding chaperone proteins linked to cadmium accumulation in breast tissue and oxidative stress, were among the most consistently induced genes across 4 conditions (cadmium 0.1-10, arsenic 10 µM).^31^ Their co-enrichment reflects a coordinated, cancer-relevant stress-adaptation signature rather than isolated pathway hits.

Additionally, pro-inflammatory signaling genes demonstrated consistent upregulation, with IL6 upregulated across 8 dose conditions (BPA 10, BPS 10, PFNA 10, cadmium 10, copper 1-10, lead 1-10, arsenic 1-10 µM) and TNF across 4 conditions (cadmium 10, copper 1-10, arsenic 10 µM) (**Figure S-2**). Both cytokines are associated with breast cancer progression and promoting therapeutic resistance.^32,33^ Growth factor gene (*FGF19*), a marker associated with basal-like breast cancers,^34^ demonstrated coordinated expression changes across similar overlapping exposures (4 conditions: BPA 10, cadmium 1-10, arsenic 10 µM). N-cadherin (CDH2), an epithelial-to-mesenchymal transition marker associated with invasiveness^7^, was upregulated across 4 conditions (BPS 10, cadmium 1-10, copper 1-10, arsenic 10 µM). Conversely, SOX10, a transcription factor that regulates stem/progenitor cell states in mammary epithelium,^35^ showed predominantly downregulation across 6 conditions (PFNA 10, cadmium 1-10, copper 1-10, arsenic 1-10 µM), indicating chemical-induced alterations in mammary epithelial cell-state regulation.

Across the 594 Hallmark gene set intersections (FDR < 0.1) for directionally consistent DEGs across all three doses, 96.97% of the total intersections were gained through upregulated genes (**Table S-7**). Pathway-level enrichment analysis further revealed that mitotic spindle assembly and protein secretion recurred as the most pervasive stress-response signatures across all chemicals at every dose (except p,p’-DDE and PFNA). Heavy metals contributed the largest share: lead acetate (*n_MITOTIC_SPINDLE_ = 61; n_PROTEIN_SECRETION_=44*), sodium arsenite (*n_MITOTIC_SPINDLE_ = 58; n_PROTEIN_SECRETION_ = 31*), and copper chloride (*n_MITOTIC_SPINDLE_ = 47; n_PROTEIN_SECRETION_ = 23*), exhibiting the most extensive and consistent DEG enrichment, as reflected by large bubble sizes and strong coloration (**Figure 2B; Table S-7**). PI3K/AKT/mTOR signaling and activation of the mTORC1 complex, key oncogenic pathways involved in cell growth and survival, were predominantly enriched in heavy metal exposures, with notable similarities in DEG intersection and upregulated directionality between lead acetate (*n_PI3K_AKT_MTOR_SIGNALING_= 27; n_MTORC1_SIGNALING_ = 58*), sodium arsenite (*n_MTORC1_SIGNALING_ = 41*), and copper chloride (*n_PI3K_AKT_MTOR_SIGNALING_= 20*). To a lesser extent, cadmium chloride also produced upregulated DEG enrichments (*n_MITOTIC_SPINDLE_ = 28; n_PROTEIN_SECRETION_ = 19*), but exhibited the single largest proportion of downregulated Hallmark terms driven by a subgroup of genes regulated by MYC (*n_MYC_TARGETS_V1_* _=_ 12). These heavy metals also demonstrated broader transcriptomic perturbations, significantly activating 3-5 hallmark pathways each and preferentially producing upregulated enrichments (93.33% across all heavy metal pathways).

In contrast, organic compounds (BPA, BPS, PFNA, and p,p’-DDE) exhibited fewer transcriptional overlaps, with each chemical associated with ≤11 DEG intersections and variable pathway-level enrichment (**Figure 2B; Table S-7**). Specifically, significant pathway enrichment was detected for BPA (*n_total_ =* 5 pathways) and BPS (*n_total_ =* 4 pathways), whereas p,p’-DDE and PFNA showed no directionally consistent DEG enrichment across all dose exposures (**Figure 2B**). Both BPA and BPS elicited distinct yet overlapping transcriptional responses characterized by direction-specific activation of multiple hallmark pathways (**Table S-**^7^). BPA and BPS predominantly induced upregulated enrichments (BPA: n_total_ = 4/5 pathways; BPS: n_total_ = 3/4 pathways). Genes involved in protein secretion pathways (BPA: *n_PROTEIN_SECRETION_ =7*; BPS: *n_PROTEIN_SECRETION_ =9*) and canonical upregulation in response to low oxygen levels (BPA: *n_HYPOXIA_= 9*; BPS: *n_HYPOXIA_ = 10*) were preferentially upregulated across all bisphenol dose exposures. BPA uniquely activated stress-related metabolic reprogramming toward glycolytic metabolism, such as upregulated enrichments of genes encoding proteins involved in glycolysis and gluconeogenesis (*n_GLYCOLYSIS_ = 8*), and genes canonically downregulated in response to ultraviolet radiation (*n_UV_RESPONSE_DN_ =7*). Whereas BPS triggered upregulated enrichment of genes important for mitotic spindle assembly (*n_MITOTIC_SPINDLE_ =11*). Downregulated pathways were mutually exclusive and limited to enrichment of genes regulated by MYC (BPA: *n_MYC_TARGETS_V1_* _=_ 2) and genes encoding proteins involved in oxidative phosphorylation (BPS: *n*_OXIDATIVE_PHOSPHORYLATION_ = 4). Collectively, these direction-aware findings reveal a shared molecular signature in which bisphenols may impose a dual oncogenic stress—amplifying proliferative and survival circuits while dampening protective DNA-repair module checkpoints—and signify a metabolic shift away from oxidative metabolism.

### Mammary Cells Exhibit Distinct Changes in Cellular Plasticity in Response to Exposure Disparity Chemicals

Next, we investigated whether transcriptional alterations induced by exposure disparity chemicals are linked to shifts in mammary cell-type composition. To assess the cellular composition of mammary epithelial cultures, we performed bioinformatic deconvolution of bulk RNA-seq data using a normal human breast single-cell reference atlas. Cell-type proportions were estimated for myoepithelial and luminal progenitor populations across individual donor samples and compared across three doses (0.1, 1, and 10 µM) for each treatment condition relative to vehicle-matched controls (0 µM). Baseline cell-type proportions in control samples exhibited substantial inter-individual variability in the relative abundance of myoepithelial and luminal progenitor cells (**Figure 4A**). Donors showed a higher proportion of myoepithelial cells and KCR7953 and KCR8195 were predominantly composed of myoepithelial cells compared to others.

**Figure 4:**
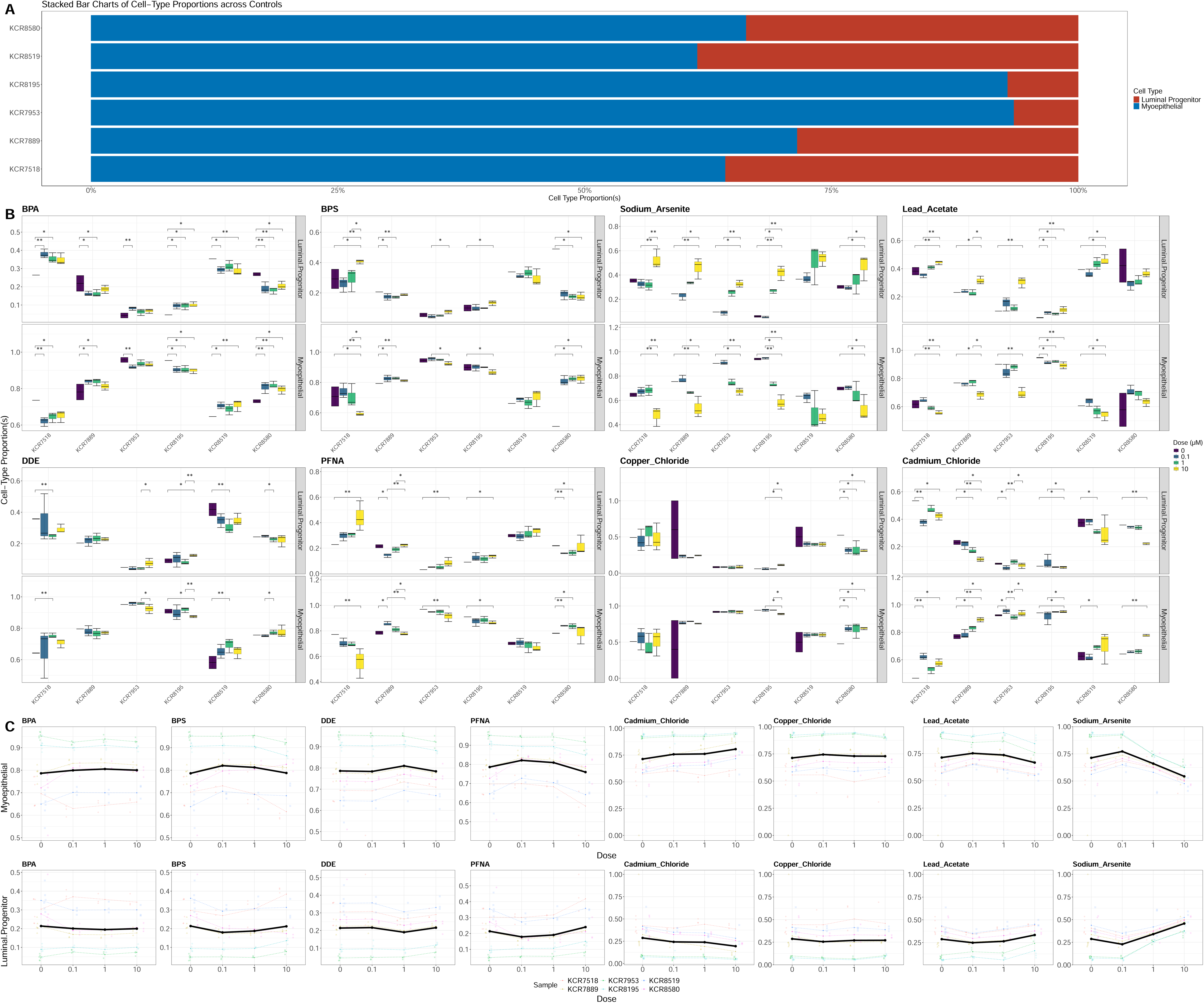
Estimated Baseline and Dose-Responsive Shifts in Cell-Type Proportions. **(A)** Stacked bar charts showing baseline (vehicle control) cell-type proportions—myoepithelial (blue) versus luminal progenitor (red)—deconvoluted from bulk RNA-seq data using a normal human breast single-cell reference atlas. Each bar represents an individual donor sample. **(B)** Boxplots of estimated myoepithelial (*bottom grid*) and luminal progenitor (*top grid*) proportions across three doses (0.1, 1, 10 µM) of each chemical and vehicle control. Colors denote dose, and each facet corresponds to one chemical. Significant pairwise comparisons are indicated by an asterisk (Dunnett’s post-hoc test; *p< 0.05, **p< 0.01). **(C)** Multivariate generalized linear mixed-effects (GLME) modeling of dose-responsive shifts in myoepithelial (*top row*) and luminal progenitor (*bottom row*) proportions. Each line represents a donor (colored by sample), and solid black lines depict the overall GLME trend.

Exposure disparity chemicals induce dose-dependent shifts in predicted cell-type composition, though the magnitude and directionality varies by both chemical and donor (**Figure 4B**). Heavy metals such as cadmium chloride and lead acetate induced pronounced shifts, with cadmium chloride driving an enrichment of myoepithelial cells and lead acetate increasing luminal progenitor fractions at higher doses. In contrast, sodium arsenite exhibited biphasic effects, with an increase in myoepithelial cells at 0.1 µM followed by a reversal decrease at higher doses. Organic chemicals such as BPA and BPS exhibited heterogeneous responses across individuals, with non-monotonic trends in cell-type proportion alterations. No clear association was observed between baseline myoepithelial or luminal progenitor proportions and the direction or significance of dose-dependent shifts. To formally quantify these trends across individuals, we applied multivariate generalized linear mixed-effects (GLME) modeling, treating donor identity as a random effect while modeling cell-type proportions as a function of chemical dose (**Figure 4B, Figure S-4**). When examining trends across all samples, the only significant changes in the cell-type proportions response were observed when comparing 10 µM to control group for sodium arsenite (*p = 0.001*) and when compared to dose level 10 to control group for cadmium chloride (*p = 0.026*) (**Figure 4B, Figure S-4**). Together, these data suggest that subsets of exposure disparity chemicals induce expression changes that are phenotypically anchored to cell-type composition and are highly variable by individual, chemical, dose, and cell-type.

### NHANES Chemical Biomarker and TCGA-BRCA Cohort Comparisons Indicate Cancer-Associated Transcriptional Signatures Occur at Human-Relevant Concentrations

To assess whether exposure-induced gene expression changes occur at human-relevant exposure levels, we compared transcriptional BMD estimates from the normal primary mammary cultures with biomarker concentrations from the National Health and Nutrition Examination Survey (NHANES). NHANES biomarker data were converted to molarity units to facilitate direct comparison with BMD estimates derived from bulk RNA-seq best-fit dose-response modeling.^6^

The comparison revealed substantial overlap in the distributions between NHANES biomarker concentrations and transcriptional BMDs for BPA, p,p’-DDE, copper, and lead, suggesting that these chemicals may elicit transcriptomic changes at exposure levels commonly observed in the general US population (**Figure 5**). Notably, in at least four donor samples, chemical biomarker concentrations in the US population were comparable to, or higher than, the BMDs associated with biological activity for BPA, p,p’-DDE, and lead. In contrast, the interquartile ranges of NHANES biomarker concentrations for BPS, PFNA, cadmium, and arsenic were generally lower than their respective transcriptional BMDs, suggesting that transcriptomic perturbations for these chemicals occur at concentrations higher than those typically observed in human population biomonitoring.

**Figure 5:**
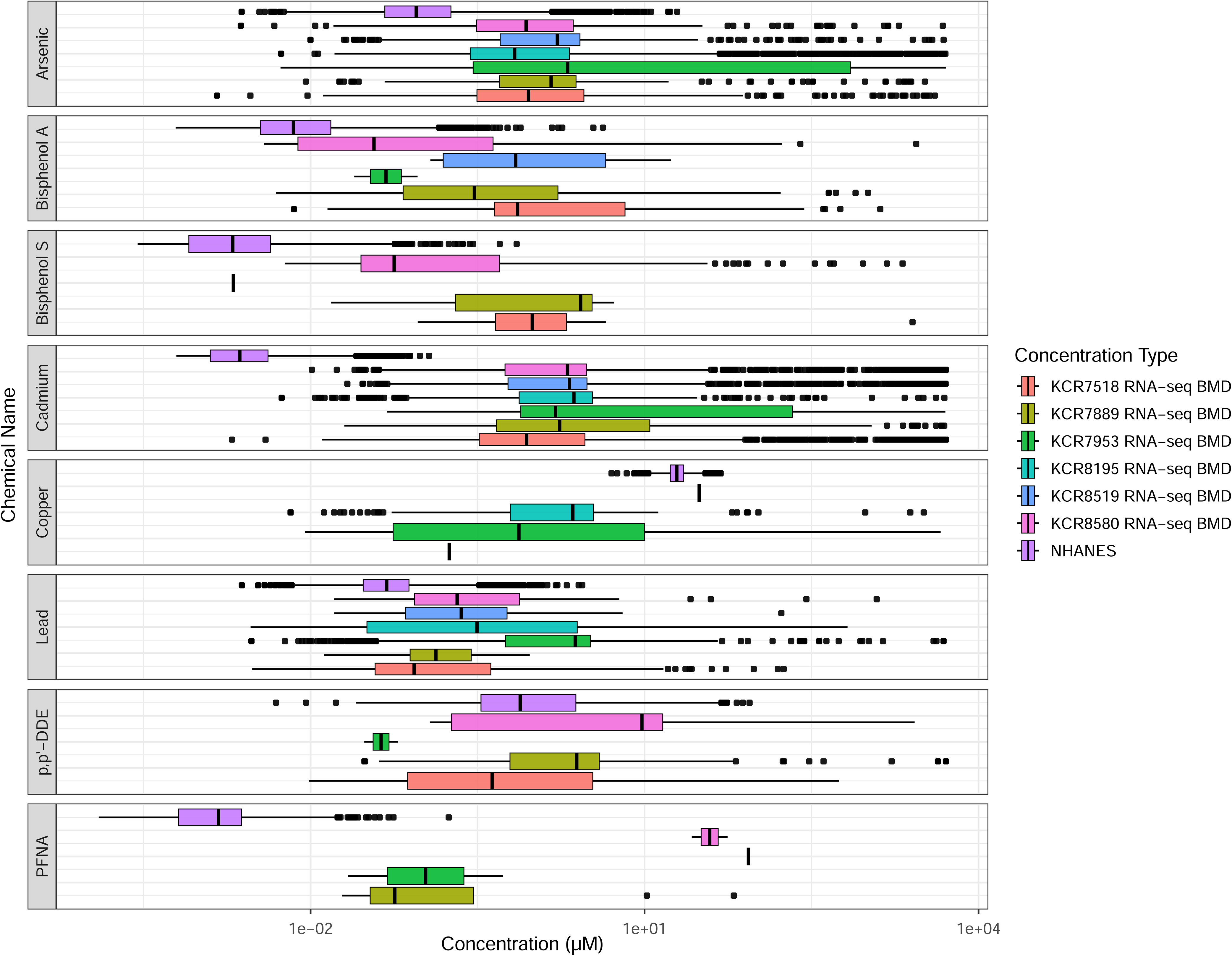
Comparison of DEG Benchmark Dose Modeling with NHANES Exposure Biomarker Levels. NHANES chemical biomarker concentrations and median transcriptional BMDs for DEGs were converted to molarity units and plotted for each donor-derived primary mammary epithelial cell line (color-coded) across the assessed chemicals (faceted grid panels). Boxplots represent the distribution of BMD values across individual samples, while NHANES biomarker concentrations illustrate the range of US human population exposures in women (*n=57786*) from 1999-2018.

Finally, to contextualize the relevance of exposure-induced transcriptional changes, we assessed whether DEGs from the normal primary mammary cultures overlapped with genes significantly associated with survival outcomes in The Cancer Genome Atlas Breast Invasive Carcinoma Collection (TCGA-BRCA) (**Table 1**). Genes from TCGA-BRCA were selected based on an FDR threshold of 0.2. To determine whether chemical exposure-induced DEGs were enriched for these cancer-associated genes, we performed Fisher’s exact tests for each chemical*dose combination (0.1 µM, 1 µM, and 10 µM). An adjusted significant p-value (≤0.05) indicates a non-random association between the two gene sets, suggesting that exposure to the tested chemical alters transcriptional programs relevant to breast cancer survival.

**Table 1.** Enrichment Testing of the DEGs Detected in the Komen Dataset with Chemical Exposure vs. TCGA BRCA Survival Genes Across Concentrations.

| Dose | BPA | BPS | DDE | PFNA | Cadmium<br>Chloride | Copper<br>Chloride | Lead_<br>Acetate | Sodium<br>Arsenite |
| --- | --- | --- | --- | --- | --- | --- | --- | --- |
| 0.1 | 0.584 | 0.388 | 0.110 | 0.806 | 0.897 | 0.591 | 0.011<br>(*n*=97) | 0.033<br>(*n*=77) |
| 1 | 0.653 | 0.630 | 0.318 | 0.537 | 0.301 | 0.716 | 0.377 | 0.008<br>(*n*=148) |
| 10 | 1 | 0.883 | 0.418 | 0.780 | 0.031<br>(*n*=184) | 0.031<br>(*n*=117) | 0.088 | 0.352 |

*Note:* p-values from Fisher's exact test assessing number of DEGs detected association between the Komen dataset and TCGA BRCA dataset are shown. Results are stratified by chemical and dose level (0.1 μM, 1 μM, and 10 μM). The number of genes detected in the Komen dataset (*n*) are indicated for significant doses ( $p \leq 0.05$ ).

The results indicate that exposure to specific chemicals, particularly heavy metals, induced gene expression changes that significantly overlapped with survival-associated genes in TCGA-BRCA (**Table 1**).^26^ Lead acetate and sodium arsenite exhibited the strongest associations, with significant enrichment at multiple dose levels. At 0.1 µM, lead acetate showed a significant overlap (*p = 0.011, n = 97*) as well as sodium arsenite (*p=0.033, n=77*), while at 1 µM, sodium arsenite exhibited a more pronounced association (*p = 0.008, n = 148*). At the highest dose (10 µM), cadmium chloride (*p = 0.031, n = 184*) and copper chloride (*p = 0.031, n = 117*) maintained significant enrichment compared to TCGA-BRCA gene sets. In contrast, chemical exposures such as BPA, and BPS, pp’ DDE and PFNA did not exhibit significant enrichment, suggesting that their induced transcriptional changes may not be directly associated with cancer mortality-associated gene expression patterns.

These findings suggest that exposure disparity chemicals, particularly heavy metals, induce gene expression alterations that are significantly associated with genes linked to breast cancer survival. This overlap underscores the potential for these chemicals to disrupt key transcriptional programs relevant to tumor progression and disease outcomes.

## Discussion

To the best of our knowledge, this is the first study that leverages a high-throughput integrated analysis of multimodal bulk sequencing data to investigate environmental contributions to breast cancer risk across 8 chemicals with racial exposure disparities, 3 human-relevant doses, and 6 normal human primary cell lines from diverse genetic backgrounds. This experimental design exemplifies a New Approach Methodology (NAM) towards modern toxicological approaches, such as high-throughput transcriptomics, which enable chemical hazard assessment in a human-relevant context without reliance on traditional animal models. Our high-throughput nested design enables comprehensive genome-wide analysis with computational rigor at human-relevant doses in primary human cells and exemplifies a cost-effective approach to mechanistic toxicity evaluation. Such NAM-based approaches align closely with evolving regulatory paradigms, including the FDA Modernization Act 2.0, underscoring their relevance and translatability to both scientific and public health communities.

From these transcriptomic analyses, we identified several key findings: 1.) the magnitude, significance, direction, and dose-responsiveness of transcriptomic alterations highly vary by individual, chemical, dose, and estimated cell-type proportions; 2.) exposure disparity chemicals elicit transcriptional responses that are phenotypically anchored with changes in cell-type distributions; 3.) exposure disparity chemicals commonly enriched cancer- and stemness-related biological processes and pleiotropic signaling pathways regulated by distinct and convergent signal transduction mechanisms; 4.) response to inorganic heavy metal exposures shared greater overlaps in the magnitude of transcriptomic alterations across individuals as opposed to organic compounds; 5.) and median transcriptional BMD values in at least 4 out of 6 diverse primary cell lines overlapped with the range of NHANES biomarker levels for BPA, p,p’-DDE, copper, and lead; 6.) heavy metals had gene expression alterations significantly enriched for those observed in tumors where TCGA patients experienced breast cancer related mortality. These findings support the growing body of evidence towards a personalized understanding of breast cancer risk and precancerous conditions, emphasize human-relevant exposures with etiological contributions to breast cancer, and offer new insights and validations into the mechanisms underlying mutagenic and non-mutational drivers of chemical carcinogenesis.

Examination of individual genes and biological pathways perturbed by exposure disparity chemicals revealed consistent alterations of key regulatory and functional processes, particularly those governing proliferation, protein secretion, and metabolic effects. First, a core proliferative program is affected in 5 of the 8 exposures, reflected by significant enrichment in the “mitotic spindle” hallmark. Second, heavy metals converged on a shared mitogenic-oncogenic signature marked by coordinated activation of growth-factor and survival pathways. Such conserved adaptive responses may inadvertently promote aggressive cancer traits: by enhancing survival, proliferation, and stress resistance, they create conditions in which damaged cells can persist. For example, high metallothionein expression has been linked to invasiveness and poorer outcomes in breast cancer, illustrating that the very mechanisms evolved to cope with toxic heavy metals can also support the survival and progression of malignant cells under stress.^36^ Third, genes associated with the “protein secretion” hallmark were commonly upregulated in 6 of the 8 exposures. A recent study investigating biological pathways altered in invasive breast cancers likewise identified the “protein secretion” hallmark as the most strongly enriched pathway, suggesting that positive enrichment of this gene set represents a key process in the development of invasive capacity and that it can be modulated by common environmental exposures.^37^ Fourth, the organic compounds assayed elicit comparatively sparse, chemical-specific enrichments that are confined largely to metabolic-centered effects. Taken together, these patterns provide insight into potential mechanisms and signatures underlying toxicant-induced transcriptional changes.

Notably, the directionality of transcriptional responses varied substantially by chemical and dose. While 96.97% of consistently altered hallmark genes across all doses were upregulated (reflecting activation of stress-response and survival pathways) (**Table S-7**), dose-stratified analysis revealed more complex patterns. At the highest dose tested (10 µM), sodium arsenite induced the most pronounced cytotoxic effects across all chemicals, with some individuals (particularly KCR7953 and KCR8195) showing enhanced sensitivity to heavy metals (**Table S-5**). At 10 µM arsenite, the near-universal transcriptional downregulation observed in some samples (e.g., KCR7953: 65% downregulation at 0.1 µM transitioning to 60% upregulation at 10 µM) may reflect overt cytotoxicity and general cellular shutdown rather than chemical-specific transcriptional programs, similar to translation inhibition effects. However, it is critical to note that 97% of all treatment conditions maintained ≥50% viability, with only 3% of conditions (primarily 10 µM arsenite in sensitive donors) approaching cytotoxic thresholds where non-specific stress responses dominate (**Figure S-1**).^27^ Furthermore, the pathways consistently enriched across chemicals—mitotic spindle assembly, PI3K/AKT/mTOR signaling, oxidative stress response—are mechanistically distinct from general xenobiotic stress responses. Notably, HALLMARK_XENOBIOTIC_METABOLISM was not significantly enriched by directionally consistent genes across all doses for any chemical (**Figure 2B**; **Table S-7**), indicating preferential activation of cancer-relevant adaptive pathways over generic detoxification responses. This pattern demonstrates that the observed transcriptional changes reflect pathway-specific biological responses aligned with key characteristics of carcinogens, including oxidative stress induction, altered cell proliferation, and survival signaling, rather than non-specific cellular injury.^38^

A growing body of literature has characterized cell populations with luminal/basal hybrid phenotypes in both normal and cancerous human and murine mammary tissue.^6,10,20^ Transcriptome-based meta-analyses recently discovered that these hybrid cells exhibit developmentally immature and embryonic stem cell-like molecular signatures, which are implicated in the development of aggressive basal-like cancers.^20^ While we were unable to characterize transcriptomic differences by cell-type, the bulk data alignment with single-cell RNA-seq atlases of the normal human mammary gland allowed us to estimate and test for dose-responsive transcriptional alterations in cellular heterogeneity. This high-throughput nested experimental design format enabled us to achieve cost-effective high genome coverage and computational rigor in our analysis. Using cell-type deconvolution and benchmark dose analyses, we found evidence of largely non-monotonic changes in cell-type proportions across the vast majority of treatments, and most notably, luminal-to-basal transitions in exposures to cadmium. Surprisingly, arsenic exposures exhibited slight increases in the proportion of estimated myoepithelial cells at 0.1µM, followed by steep decreases at higher doses, while the opposite trend was observed for luminal progenitor proportions. These findings corroborate previous studies that documented basal-like phenotype transitions following treatments with chronic low-level cadmium (2.5µM) and arsenic (0.1µM), and provide new insights into complex multi-partner interactions with PI3K/Akt/mTOR signaling that may underlie the biphasic transcriptional effects of arsenic.^31,39–42^

The observed transcriptomic overlap between chemically exposed mammary epithelial cultures and genes associated with poor survival in TCGA-BRCA tumors provide additional insight into the potential relevance of exposure-induced gene expression changes. Significant overlap in gene expression suggests that certain chemicals induce transcriptional shifts that resemble gene expression signatures related to breast cancer metastasis and poor survival.

Notably, heavy metal chemicals such as sodium arsenite, cadmium chloride, copper chloride, and lead acetate exhibited the strongest mortality signature overlap (**Table 1**). While these findings do not establish causation, they highlight the potential for exposure to these chemicals to disrupt gene expression patterns relevant to breast cancer progression. Given that the genes tested were selected based on their association with survival outcomes in breast cancer patients (FDR < 0.2), significant enrichment indicates that exposure may perturb key pathways implicated in tumorigenesis. However, a lack of significant overlap does not necessarily imply an absence of effect, as dose-dependent or non-linear transcriptomic responses may not be captured in this specific statistical framework. These findings support a growing body of epidemiological evidence linking exposure to some of these common chemicals and breast cancer mortality.

For example, a meta-analysis of 36 studies encompassing 5,747 women demonstrated that breast cancer patients exhibited substantially elevated serum copper levels and copper/zinc ratios compared to both healthy controls and those with benign breast disease.^43^ These cross-sectional findings are complemented by prospective survival data from the Sweden Cancerome Analysis Network-Breast Initiative, where individuals with serum copper/zinc ratios in the highest quartile experienced a 1.58-fold increased risk of overall mortality compared to those in the lowest quartile, even after adjusting for age, menopausal status, tumor characteristics, and treatment factors.^44^ Notably, the predictive capacity of the copper/zinc ratio approached that of tumor size and exceeded lymph node status in time-dependent analyses. At the molecular level, transcriptomic profiling has revealed that breast cancer patients with elevated expression of copper-related genes demonstrate diminished survival, altered immune cell infiltration profiles, and differential therapeutic vulnerabilities.^45^ Mechanistic studies have demonstrated that copper depletion impairs mitochondrial oxidative phosphorylation and suppresses metastasis in triple-negative breast cancer models,^46^ supporting ongoing clinical trials targeting copper metabolism in treatment-resistant breast cancers.^47^ In an analysis of 209 patients from the Cancer Hospital of Shantou Medical Hospital, blood cadmium concentrations of greater than 3 µg/L were associated with a 2.25-fold increased odds of distant metastasis compared to individuals with blood cadmium less than 3 µg/L.^48^ Data examining impacts of arsenic and lead and their impacts of breast cancer mortality remain equivocal^49–52^ and limited, and represent an area for future epidemiologic investigation, particularly in the context of racial disparities.

While our study highlights cancer-associated biological pathways which are dysregulated in primary human breast cells at human relevant doses, further analyses, such as pathway-level enrichment and integrative network modeling, are needed to determine whether exposure-induced changes activate oncogenic programs or reflect broader stress and adaptation responses. The overlap between population-relevant exposures and transcriptional perturbations underscores the importance of linking *in vitro* models with epidemiological data to assess the real-world impact of environmental chemicals on breast cancer susceptibility.

Additionally, while our present study provides a high-throughput screening of cancer initiation and promotion processes, earlier biochemical changes induced by environmental chemical stressors frequently coincide with physiological mechanisms of cellular repair and adaptation, which may not be unique to toxicity. Many of these signal transduction mechanisms also differ markedly in the spatiotemporal course of induction of their cellular effects and share complex interactions within and across pathways. Our experimental design employed an acute exposure paradigm (48 hours at 0.1–10 µM), which, while enabling high-throughput analyses across multiple chemicals and individuals, does not fully capture the chronic, low-dose exposure scenarios more representative of real-world human exposures. Acute exposure models at higher concentrations are valuable for identifying initial transcriptional responses and establishing concentration-response relationships, but they may not reflect the cumulative, adaptive, or compensatory cellular responses that emerge over prolonged exposures. Chronic exposure models using microfluidic organ-on-chip systems could provide more physiologically relevant insights by enabling longer perturbation times (weeks to months) at lower, environmentally relevant concentrations while maintaining cellular architecture and dynamic exposure conditions.^53,54^ Such systems would be particularly valuable for distinguishing transient stress responses from persistent epigenetic and phenotypic alterations that may drive cancer initiation. While our dose range (100 nM–10 µM) was established through benchmark concentration modeling linking in vitro transcriptomic responses to NHANES biomarker data^6,14^, the concentrations tested span three orders of magnitude and may not uniformly reflect physiological tissue concentrations following environmental exposures. In-vitro-to-in-vivo extrapolation (IVIVE) considerations are critical when interpreting these findings in a human health context, including differences between administered dose and internal tissue concentrations, metabolic activation/detoxification, protein binding, and cellular uptake kinetics.^55,56^ In vitro concentrations (particularly at 10 µM) may exceed steady-state tissue concentrations, though peak tissue levels following acute exposures or localized accumulation in lipophilic tissues may approach these ranges, particularly for persistent chemicals like p,p’-DDE (half-life ∼6-10 years) and metals that bioaccumulate in tissues.^31,57^ Future work should consider effective integrations of organotypic models, single-cell multi-omic technologies, and transient-dynamic data modalities to identify and capture specific contributions to mutational and non-mutational signatures of cancer risk.

Moreover, although we observed dose-dependent inter-individual differences in response to chemicals, our data did not identify clear patterns in molecular signatures by race, likely due to limited study power (**Table S-1**). The substantial inter-individual variability in baseline cell-type proportions (**Figure 4A**) and chemical responsiveness likely reflects biological heterogeneity among donors rather than technical artifacts from tissue collection or processing. All samples were collected, processed, and expanded using standardized conditional reprogramming protocols, and inter-individual differences persisted across multiple experimental batches.^21^ Factors contributing to donor-specific responses may include genetic polymorphisms in xenobiotic metabolism enzymes (e.g., CYP450s, glutathione-S-transferases), epigenetic differences established during development or prior environmental exposures, and inherent variation in mammary epithelial cell-type composition and differentiation states.^10,23,58^ For instance, KCR8195 demonstrated heightened sensitivity to multiple heavy metals (arsenic, cadmium, copper), while KCR8519 showed minimal transcriptional responses to several chemicals, patterns consistent across independent experimental replicates. As mentioned by Thong (*2022*), despite a moderately sized donor pool, the successful CR establishment and expansion of primary human samples was further complicated by self-defined race versus genetic ancestry.^15^ Future cohort studies and data sharing protocols could be designed to expand racially diverse normal tissue biobanks and gain more insights into environmental carcinogenic processes.

Overall, this study expands HTTr testing of prioritized chemicals with exposure disparities and examines their effects on aggressive breast cancer-associated biology in normal primary mammary cells from diverse individuals at human-relevant doses. Future research will involve additional characterizations across single-cell multi-omics, functional- and marker-based validation assays, and molecular epidemiology cohorts to better understand the health impacts of chemical exposure disparities, guide the development of molecular biomarkers, and inform targeted interventions and personalized risk assessments for highly exposed populations.

## Acknowledgements.

This work was supported by grants from the National Institutes of Health R01 ES028802, P30 ES017885, UH3 CA267907, P30 CA046592, and T32GM141746.

## Appendix

### Supplementary Figures

**Figure S-1: Relationship Between Cell Viability and Differential Gene Expression Across Chemical Exposures.**

**(A)** Scatterplot of relative cell viability (% of vehicle control) versus number of differentially expressed genes (DEGs) across all treatment conditions (*n = 144*; 6 donors × 8 chemicals × 3 doses). Each point represents technical replicates averaged per condition. Background shading indicates viability ranges: green (≥80%), yellow (50-80%), red (<50%). Dashed line shows linear regression fit. Pearson correlation: *r = -0.244, p = 0.0032*. **(B)** Heatmap of correlation coefficients between viability and DEG count for each chemical × donor combination. Correlations calculated using individual technical replicates per combination. Diagonal cross (×) indicates non-significant correlations (*p ≥ 0.05*). Numbers indicate correlation coefficient with significance level (*\*, p < 0.05; **, p < 0.01; ***, p < 0.001*). Gray cells (*NA*) indicate no variance for correlation calculation. **(C-E)** Scatterplots of viability versus DEG count stratified by dose: 0.1 µM **(C)**, 1 µM **(D)**, and 10 µM **(E)**. Each point represents technical replicates averaged per condition. Points colored by chemical and shaped by donor. Dashed line shows linear regression fit with Pearson correlation coefficient and p-value reported for each dose.

**Figure S-2: Differentially Expressed Genes for Treatment Doses Aggregated across Samples.**

Differential gene expression between each aggregated chemical dose and control was calculated using *limma-Voom* and empirical Bayes quality and precision weighted generalized linear modeling. Vertical dotted-black lines mark a log_2_ fold-change cutoff of >|2|. Horizontal dotted-black lines mark a false discovery rate (FDR) adjusted p-value of ≤0.05.

**Figure S-3: Differentially Expressed Genes for 6 Samples Treated with Treatment Doses.**

Differential gene expression between each sample-specific chemical dose and control was calculated using limma-Voom and empirical Bayes quality and precision weighted generalized linear modeling. Vertical dotted-black lines mark a log2 fold-change cutoff of >|2|. Horizontal dotted-black lines mark a FDR adjusted p-value of ≤0.05.

**Figure S-4: GLME Model Diagnostics**

Residuals versus fitted values for luminal progenitor (*left*) and myoepithelial (*right*) cell-type proportion models. Points represent individual observations colored by treatment. Horizontal dashed line at zero. Blue curve shows LOESS smoothing of residuals. Cell-type proportions sum to 1.0.

### Supplementary Tables

**Table S-1.** Susan G. Komen Normal Human Primary Breast Cell Line Demographic Data (Adapted from Thong et al. 2022).^15^.

| Cell Line | Self-identified Race/Ethnicity | Age | BMI (kg/m2) | Days since last menstrual period |
| --- | --- | --- | --- | --- |
| KCR8195 | AA | 27 | 27.37 | 10 |
| KCR8519 | AA | 25 | 21.3 | 9 |
| KCR8580 | AA | 25 | 36.32 | 7 |
| KCR7518 | EA | 27 | 30.6 | 19 |
| KCR7889 | EA | 24 | 20.2 | 17 |
| KCR7953 | EA | 24 | 32.3 | 11 |

*Note:* Conditionally reprogrammed, nulliparous normal mammary tissues were acquired from the Susan G. Komen tissue bank according to the protocol of Thong et al. (2022). Samples either self-identified as African American (AA) or European American (EA). 6 sample cell lines ( $n=3$ AA, $n=3$ EA) were matched (Mean (s.d.)) by age (AA= 25.6 (1.2), EA= 25.0 (1.7)), BMI (AA =27.7 (7.6), EA = 27.1 (6.6)), and days since last menstrual period (AA= 8.6 (1.5), EA= 15.3 (4.2)).<sup>15</sup>

**Table S-2.** Exposure Disparity Chemical Information (Adapted from Sala-Hamrick et al. 2024).^6^.

| Chemical | CAS Number | Vendor; Catalog Number | Solvent |
| --- | --- | --- | --- |
| Cadmium Chloride | 10108-64-2 | Sigma Aldrich; 202908 | Water |
| Copper (II) Chloride | 7447-39-4 | Sigma Aldrich; 222011 | Water |
| Lead (II) Acetate Trihydrate | 6080-56-4 | Sigma Aldrich; 316512 | Water |
| Sodium (meta) Arsenite | 7784-46-5 | Sigma Aldrich; S7400 | Water |
| Bisphenol A | 9980-05-7 | Chem Service; S-F7295S | Dimethyl sulfoxide (DMSO) |
| Bisphenol S | 80-09-1 | Sigma Aldrich; 103039 | Dimethyl sulfoxide (DMSO) |
| p,p'-DDE | 72-55-9 | Chem Service; N-10875 | Dimethyl sulfoxide (DMSO) |
| PFNA | 375-95-1 | Sigma Aldrich; 394459 | Dimethyl sulfoxide (DMSO) |

*Note:* 8 chemicals prioritized based on NHANES chemical biomarker concentration data and their associated CAS number, vendor, catalog number, and solvent in which each was dissolved for cell treatments and vehicle controls.<sup>6,13</sup>

**Table S-3.** Example Plate and *plexWell* cDNA Library Preparation Layout for each Individual Cell Line.

|  | 1 | 2 | 3 | 4 | 5 | 6 | 7 | 8 | 9 | 10 | 11 |
| --- | --- | --- | --- | --- | --- | --- | --- | --- | --- | --- | --- |
| A | Control 1-Water | Cadmium 100nM 3 | Copper 100nM 2 | Arsenic 100nM 1 | Arsenic 10µM 3 | Lead 10µM 2 | DDE 100nM 1 | DDE 10µM 3 | BPS 10µM 2 | BPA 10µM 1 | PFNA 1µM 3 |
| B | Control 2-Water | Cadmium 1µM 1 | Copper 100nM 3 | Arsenic 100nM 2 | Lead 100nM 1 | Lead 10µM 3 | DDE 100nM 2 | BPS 100nM 1 | BPS 10µM 3 | BPA 10µM 2 | PFNA 10µM 1 |
| c | Control 3-Water | Cadmium 1µM 2 | Copper 1µM 1 | Arsenic 100nM 3 | Lead 100nM 2 | Control 7-DMSO | DDE 100nM 3 | BPS 100nM 2 | BPA 100nM 1 | BPA 10µM 3 | PFNA 10µM 2 |
| D | Control 4-Water | Cadmium 1µM 3 | Copper 1µM 2 | Arsenic 1µM 1 | Lead 100nM 3 | Control 8-DMSO | DDE 1µM 1 | BPS 100nM 3 | BPA 100nM 2 | PFNA 100nM 1 | PFNA 10µM 3 |
| E | Control 5-Water | Cadmium 10µM 1 | Copper 1µM 3 | Arsenic 1µM 2 | Lead 1µM 1 | Control 9-DMSO | DDE 1µM 2 | BPS 1µM 1 | BPA 100nM 3 | PFNA 100nM 2 |  |
| F | Control 6-Water | Cadmium 10µM 2 | Copper 10µM 1 | Arsenic 1µM 3 | Lead 1µM 2 | Control 10-DMSO | DDE 1µM 3 | BPS 1µM 2 | BPA 1µM 1 | PFNA 100nM 3 |  |
| G | Cadmium 100nM 1 | Cadmium 10µM 3 | Copper 10µM 2 | Arsenic 10µM 1 | Lead 1µM 3 | Control 11-DMSO | DDE 10µM 1 | BPS 1µM 3 | BPA 1µM 2 | PFNA 1µM 1 |  |
| H | Cadmium 100nM 2 | Copper 100nM 1 | Copper 10µM 3 | Arsenic 10µM 2 | Lead 10µM 1 | Control 12-DMSO | DDE 10µM 2 | BPS 10µM 1 | BPA 1µM 3 | PFNA 1µM 2 |  |

**Table S-4.**
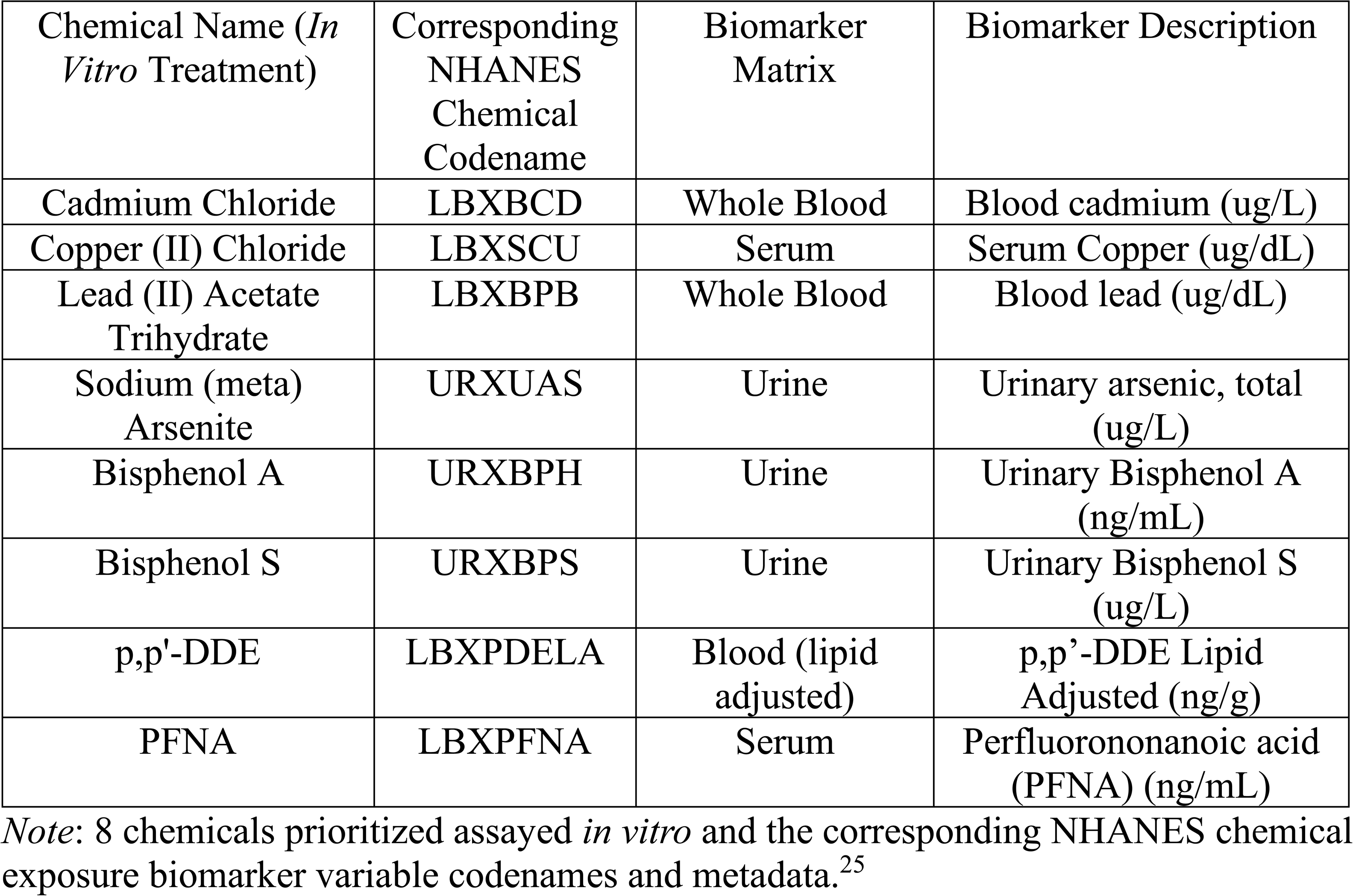
NHANES Biomarker Variable and Metadata Information (Adapted from Sala-Hamrick et al. 2022). ^6^

| Chemical Name (*In Vitro* Treatment) | Corresponding NHANES Chemical Codename | Biomarker Matrix | Biomarker Description |
| --- | --- | --- | --- |
| Cadmium Chloride | LBXBCD | Whole Blood | Blood cadmium (ug/L) |
| Copper (II) Chloride | LBXSCU | Serum | Serum Copper (ug/dL) |
| Lead (II) Acetate Trihydrate | LBXBPB | Whole Blood | Blood lead (ug/dL) |
| Sodium (meta) Arsenite | URXUAS | Urine | Urinary arsenic, total (ug/L) |
| Bisphenol A | URXBPH | Urine | Urinary Bisphenol A (ng/mL) |
| Bisphenol S | URXBPS | Urine | Urinary Bisphenol S (ug/L) |
| p,p'-DDE | LBXPDELA | Blood (lipid adjusted) | p,p'-DDE Lipid Adjusted (ng/g) |
| PFNA | LBXPFNA | Serum | Perfluorononanoic acid (PFNA) (ng/mL) |

*Note:* 8 chemicals prioritized assayed *in vitro* and the corresponding NHANES chemical exposure biomarker variable codenames and metadata.<sup>25</sup>

**Table S-5.**
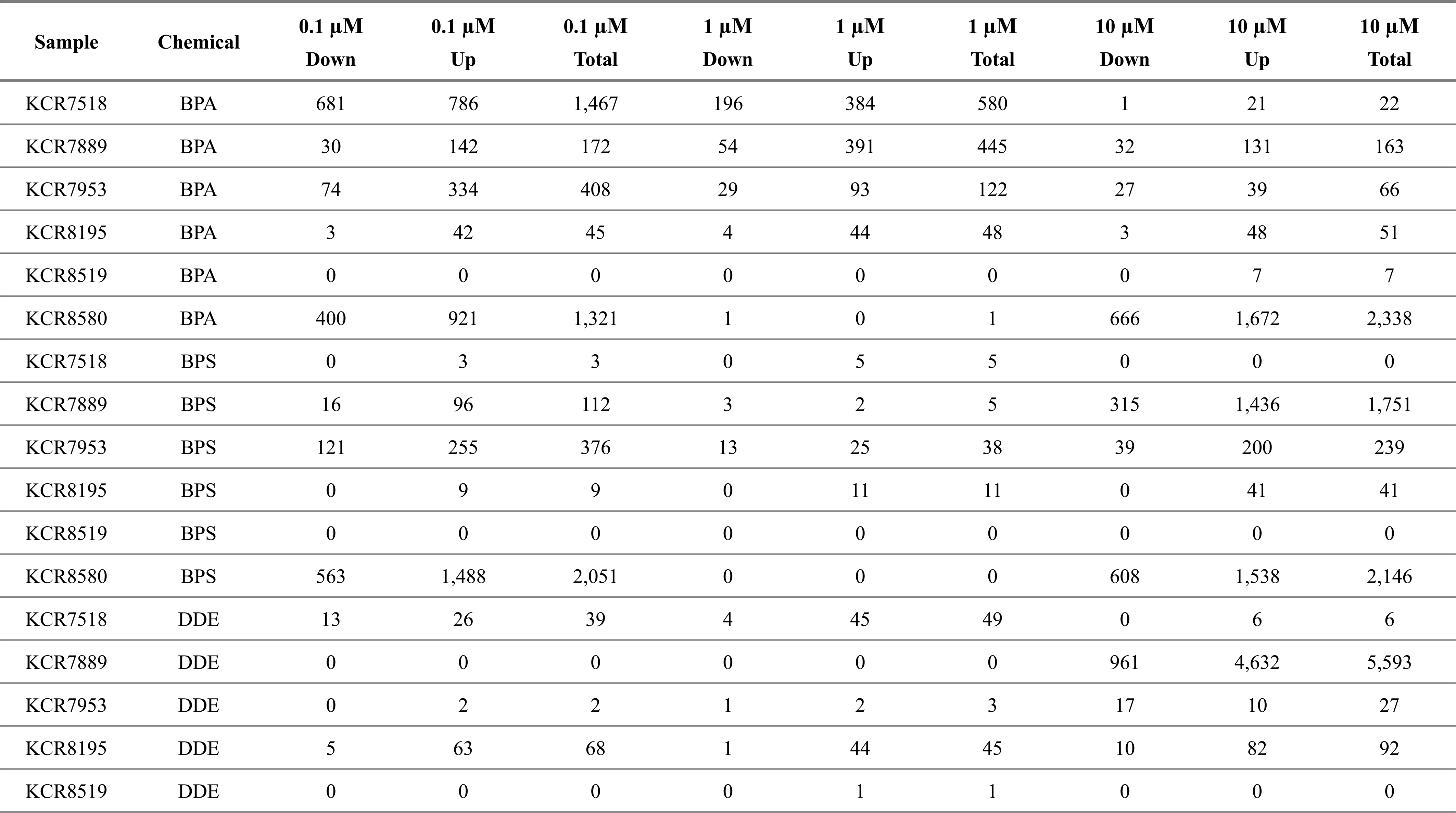
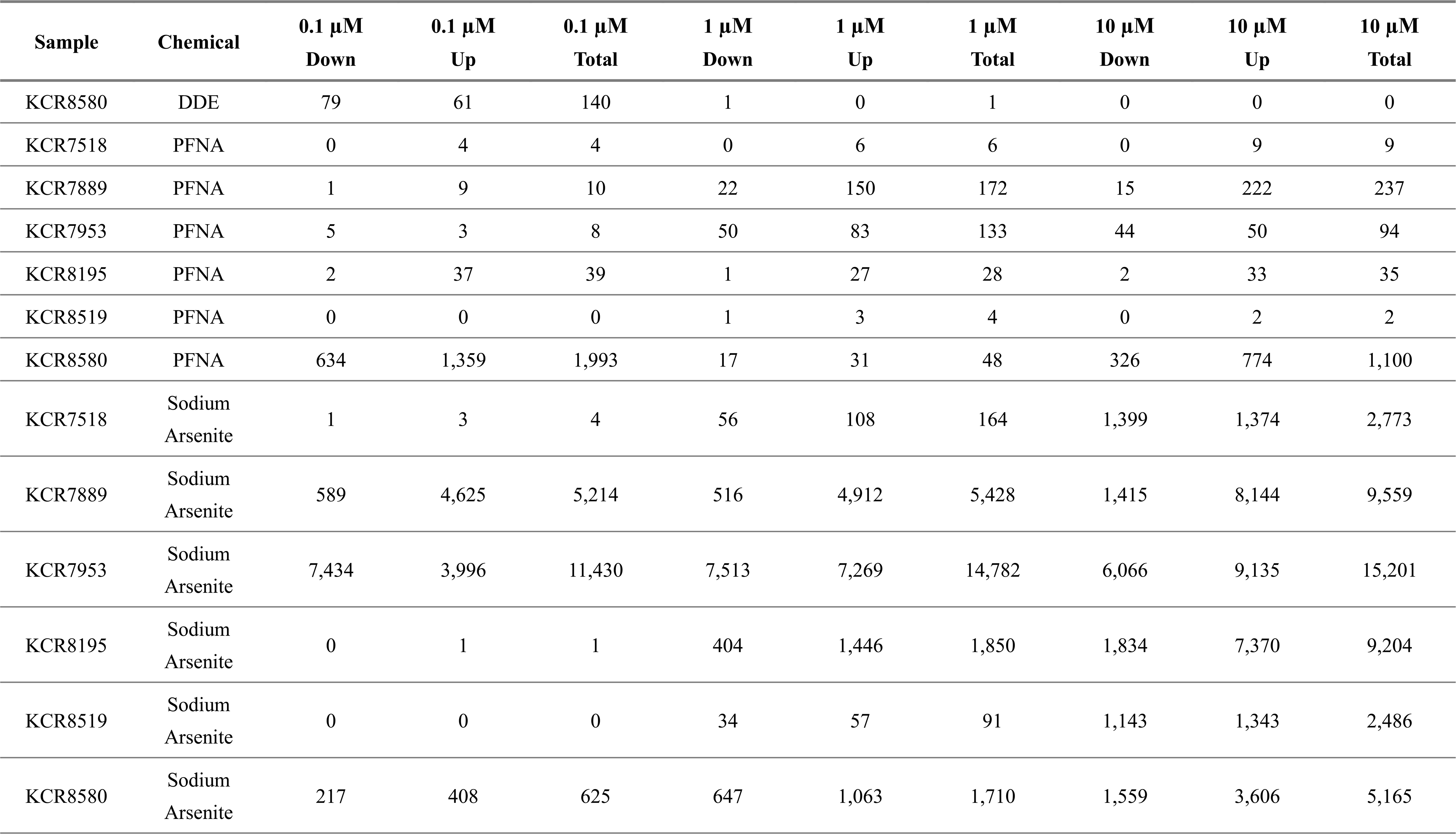
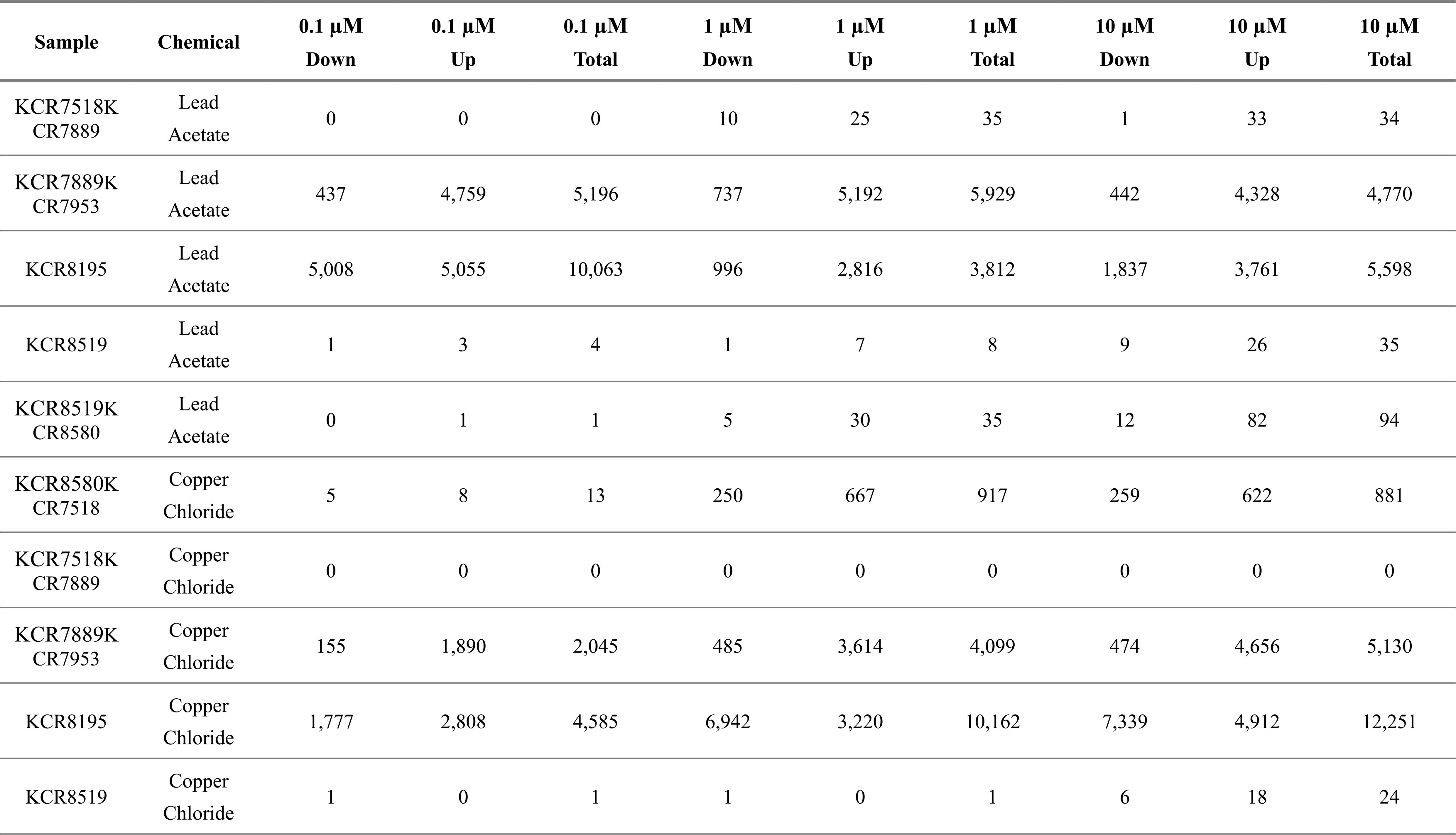
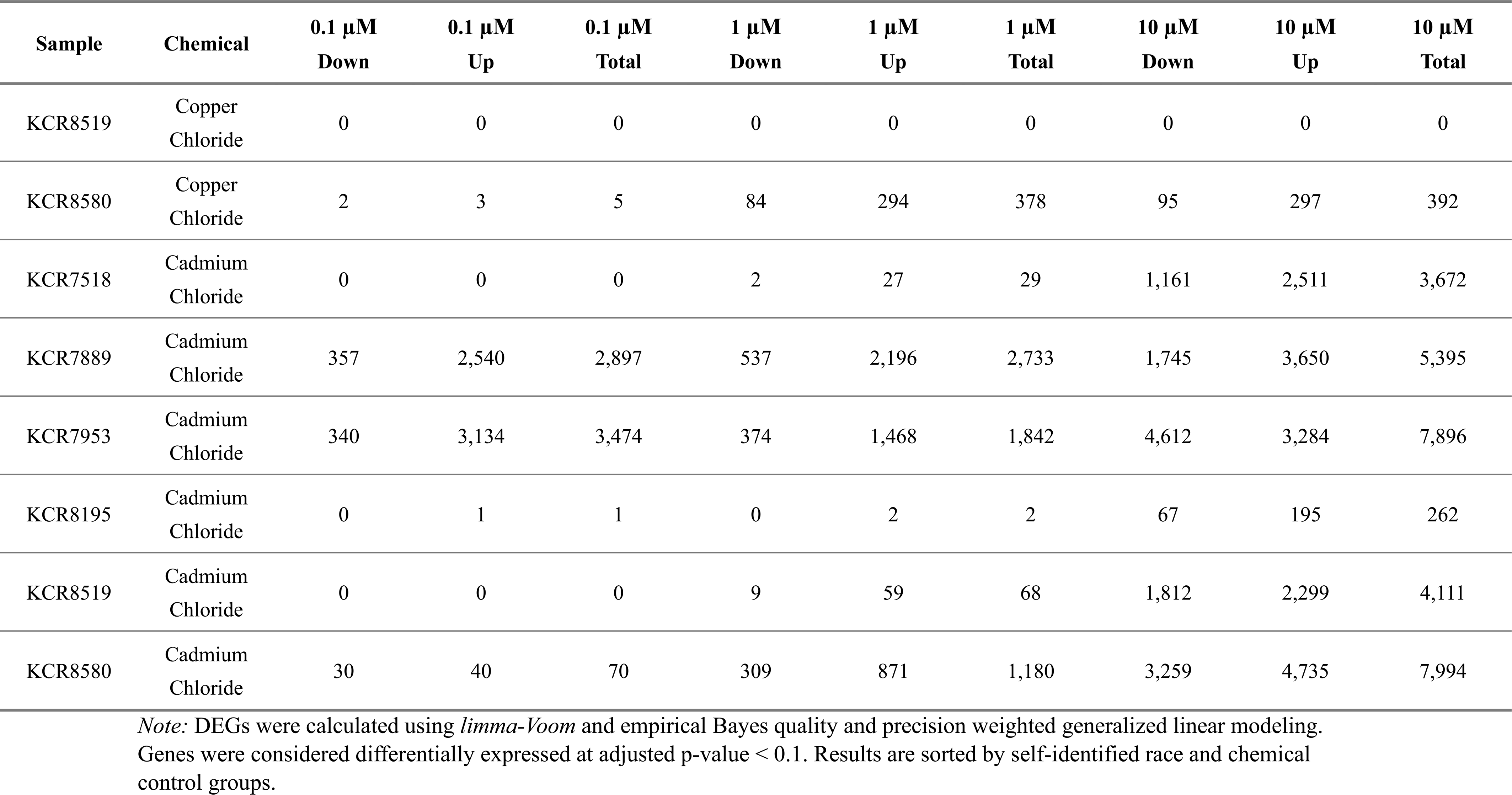
Differentially Expressed Genes (DEGs) Between Each Chemical Dose Relative to Control for Each Individual.

**Table S-6.** Proportion of DEGs That Are Consistently Up- or Down-Regulated Across All Three Doses of Each Exposure-Disparity Chemical.

| **Direction** | **Freq** | **Proportion** | **Chem** |
| --- | --- | --- | --- |
| Consistently Down | 35 | 0.1659 | BPA |
| Consistently Up | 176 | 0.8341 | BPA |
| Consistently Down | 34 | 0.1115 | BPS |
| Consistently Up | 271 | 0.8885 | BPS |
| Consistently Down | 1 | 0.25 | DDE |
| Consistently Up | 3 | 0.75 | DDE |
| Consistently Down | 36 | 0.1078 | PFNA |
| Consistently Up | 298 | 0.8922 | PFNA |
| Partially Down | 13 | 0.0117 | Cadmium Chloride |
| Consistently Down | 178 | 0.1601 | Cadmium Chloride |
| Partially Up | 49 | 0.0441 | Cadmium Chloride |
| Consistently Up | 872 | 0.7842 | Cadmium Chloride |
| Partially Down | 2 | 0.0014 | Copper Chloride |
| Consistently Down | 53 | 0.0374 | Copper Chloride |
| Partially Up | 2 | 0.0014 | Copper Chloride |
| Consistently Up | 1359 | 0.9597 | Copper Chloride |
| Consistently Down | 25 | 0.0114 | Lead Acetate |
| Partially Up | 1 | 0.0005 | Lead Acetate |
| Consistently Up | 2175 | 0.9882 | Lead Acetate |
| Partially Down | 28 | 0.0131 | Sodium Arsenite |
| Consistently Down | 141 | 0.0658 | Sodium Arsenite |
| Partially Up | 89 | 0.0415 | Sodium Arsenite |
| Consistently Up | 1886 | 0.8797 | Sodium Arsenite |

**Table S-7. Hallmark Gene-Set Enrichment (g:Profiler) for DEGs Consistently Up- or Down-Regulated Across All Doses of Each Chemical.**

