## Supplementary figures and images for "Exploring the Influence of Chemical Exposures in Breast Cancer Disparities: High-Throughput Transcriptomic Analysis in Normal Breast Cells from Diverse Donors"

### Figure S-1

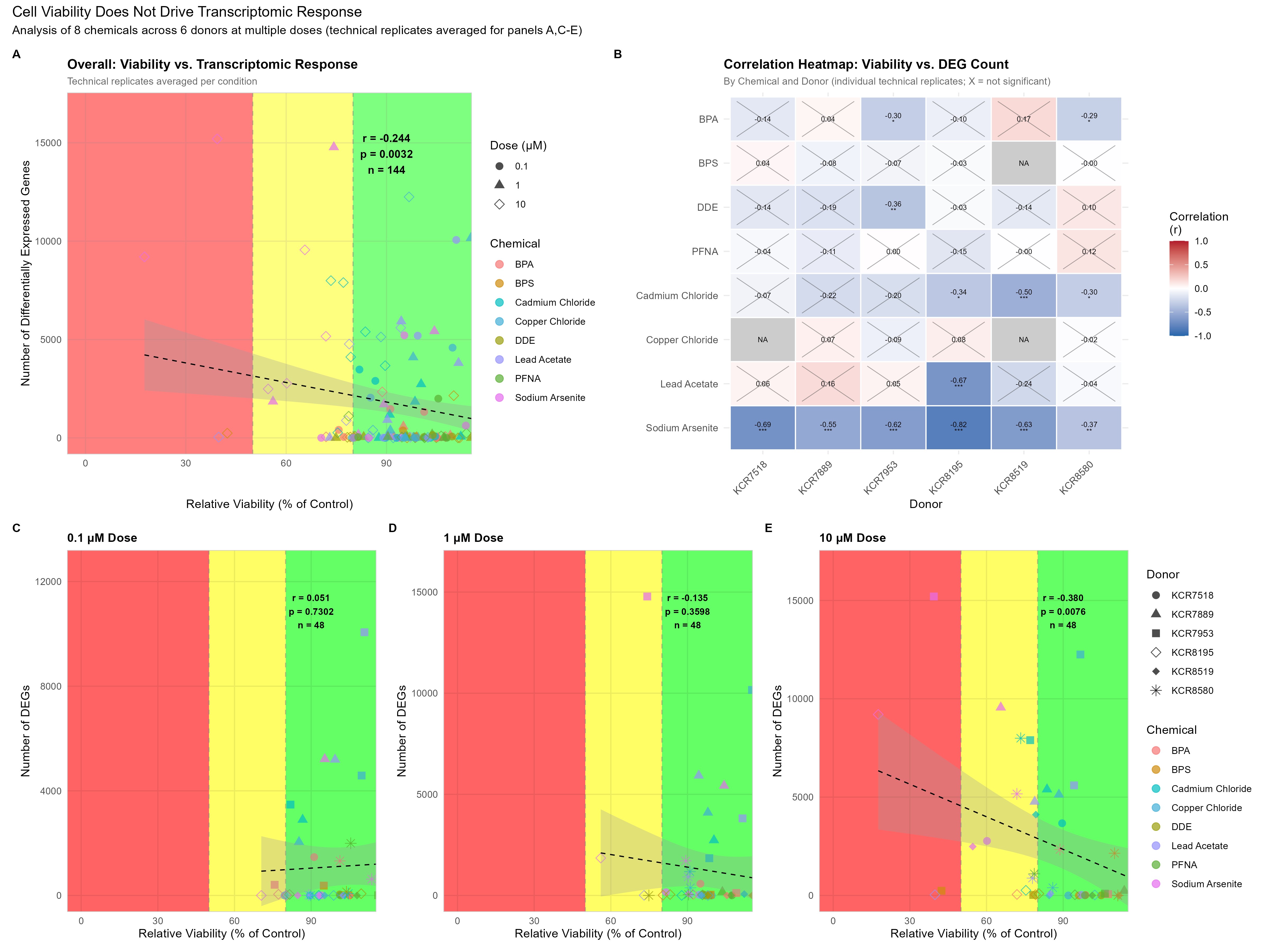

### Figure S-4

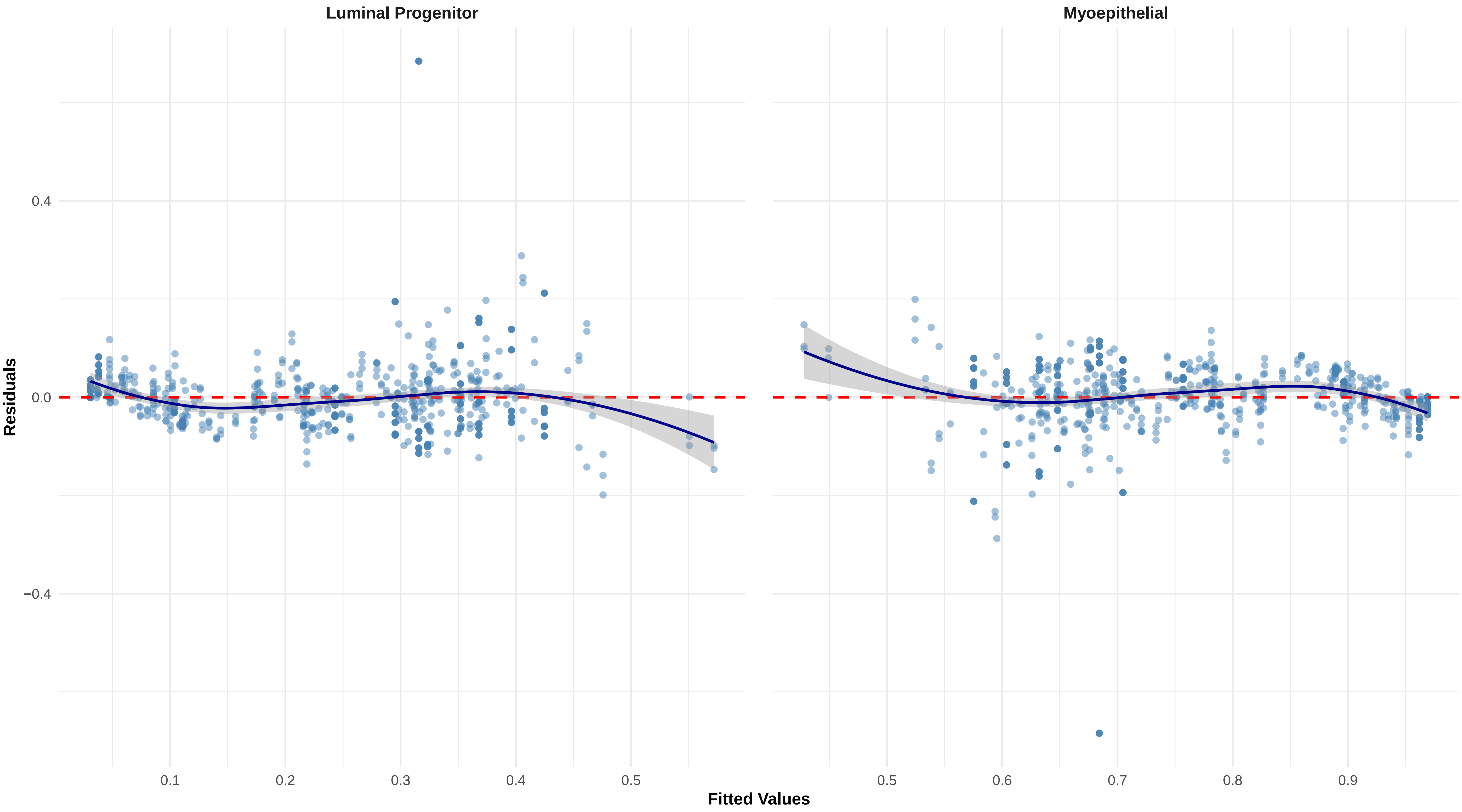
