## Supplementary material for "Exploring the Influence of Chemical Exposures in Breast Cancer Disparities: High-Throughput Transcriptomic Analysis in Normal Breast Cells from Diverse Donors": Figure S-2

### Differential Expression

BPA_0.1 − DMSO_0

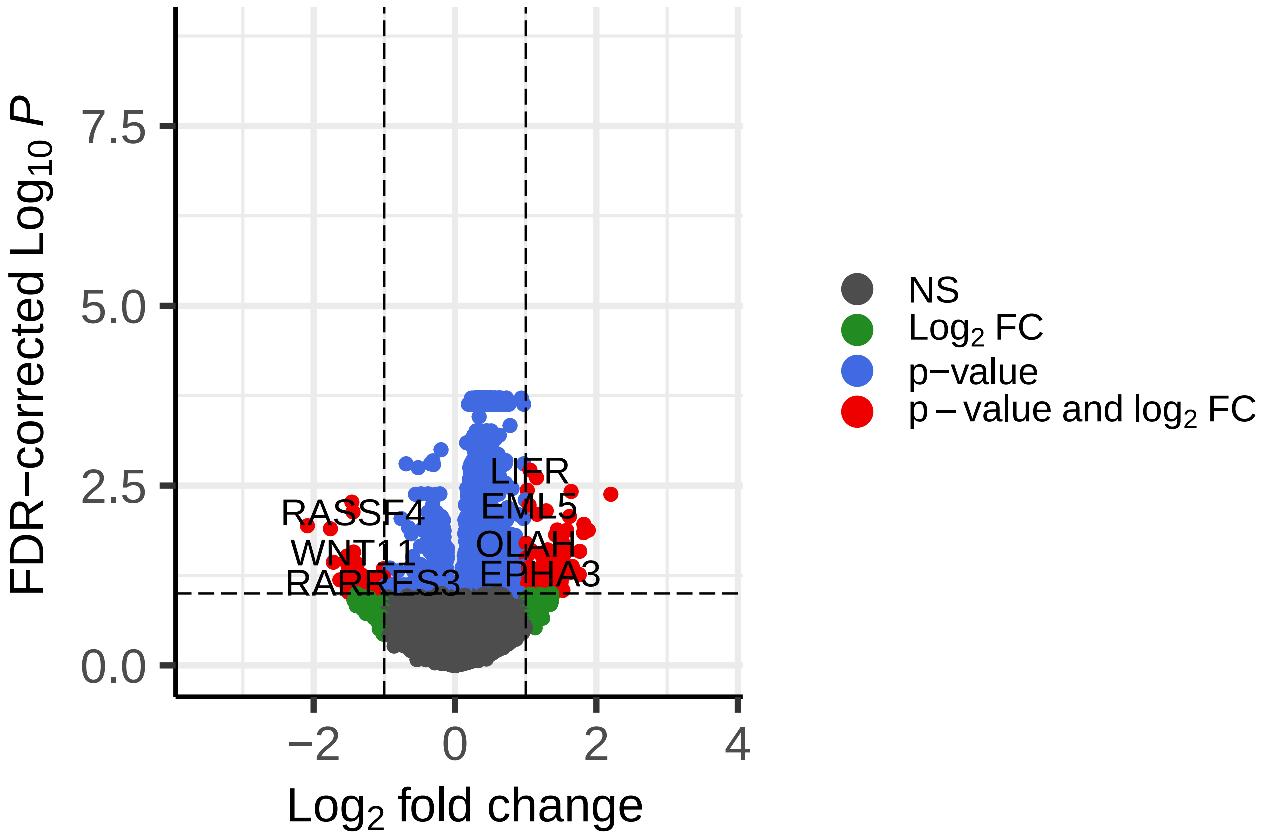


FDR−corrected permutation p−values

### Differential Expression

BPA_1 − DMSO_0

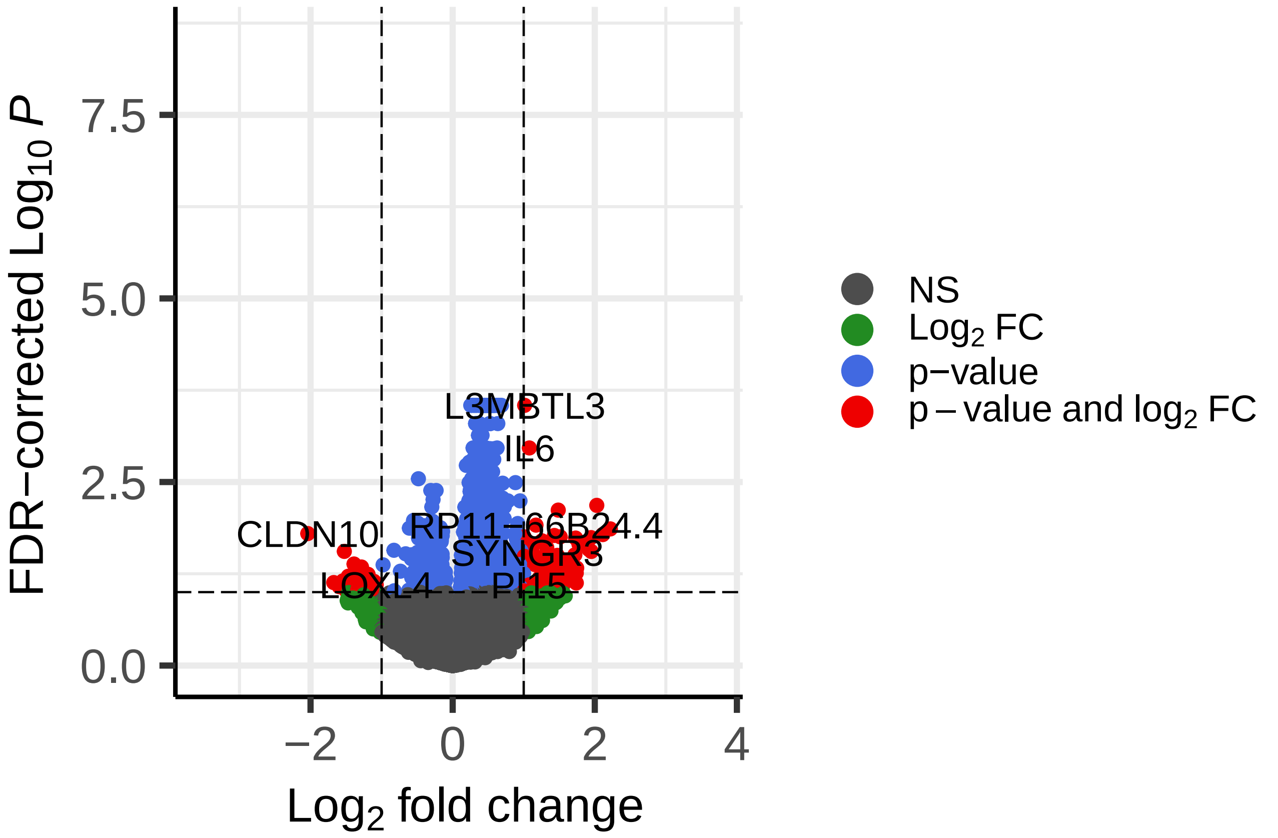


FDR−corrected permutation p−values

### Differential Expression

BPA_10 − DMSO_0

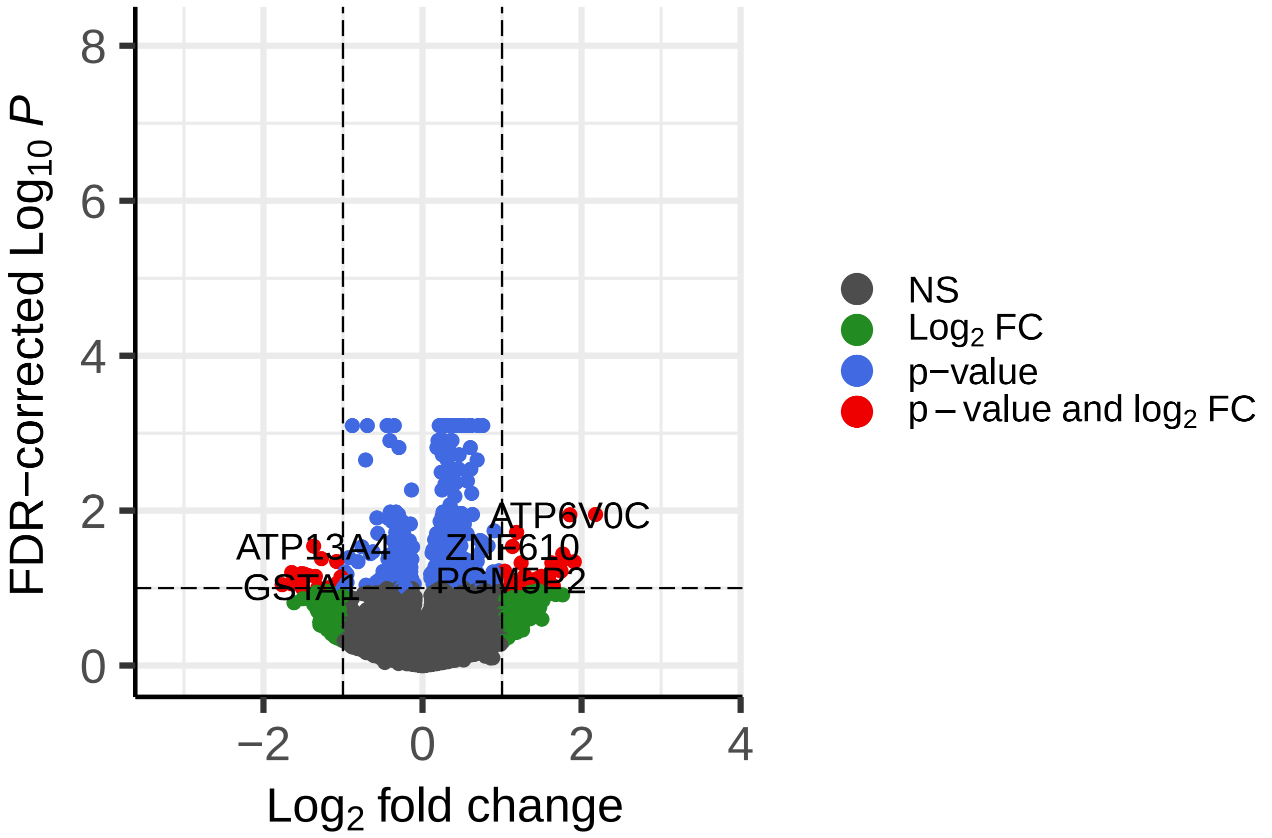


FDR−corrected permutation p−values

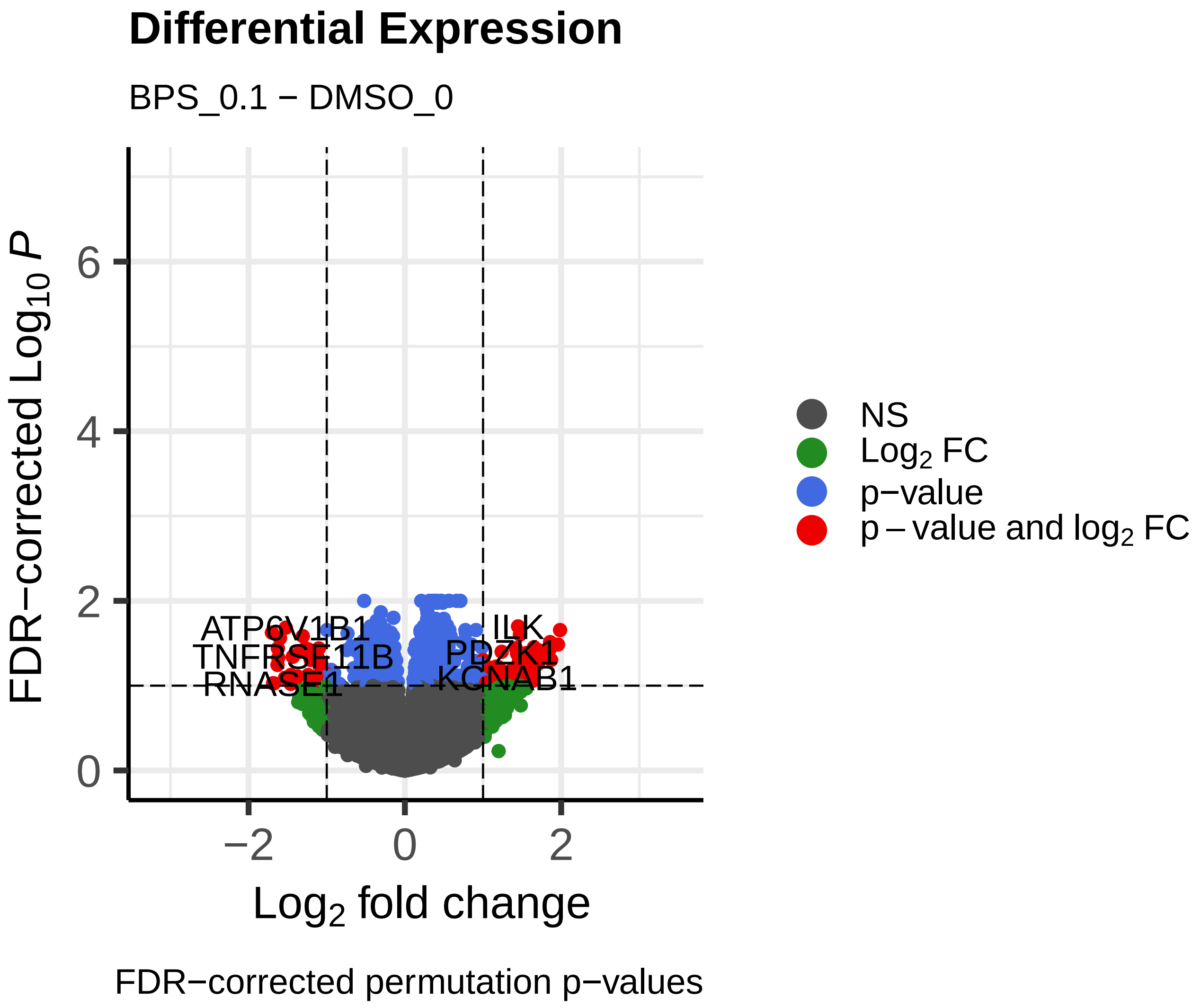
Differential Expression
BPS_0.1 − DMSO_0
FDR−corrected Log10 P
Log2 fold change
ATP6V1B1
TNFRSF11B
RNASE1
ILK
PDZK1
KCNAB1
NS
Log2 FC
p−value
p − value and log2 FC
FDR−corrected permutation p−values

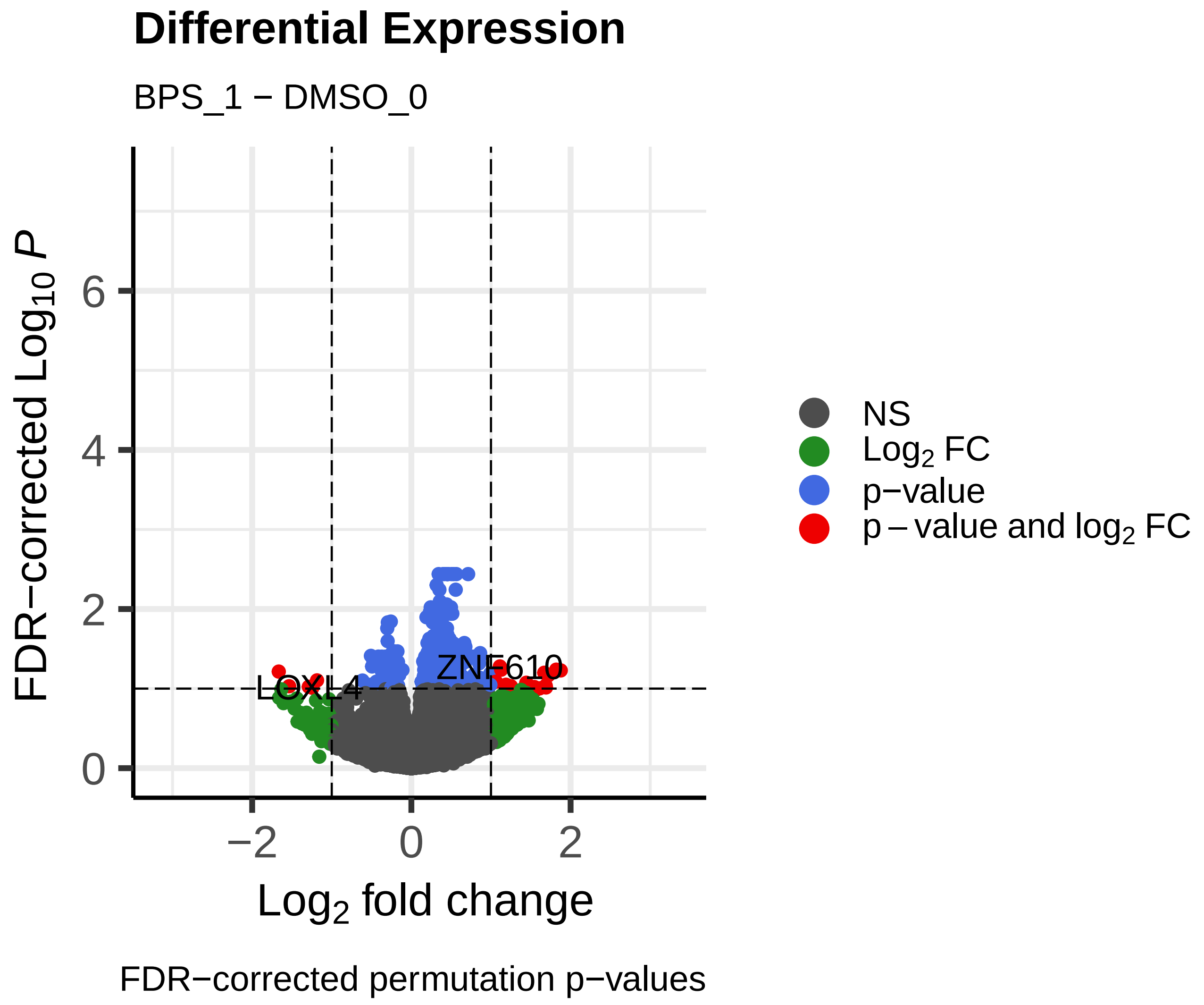
Differential Expression
BPS_1 − DMSO_0
FDR−corrected Log10 P
Log2 fold change
LOXL4
ZNF610
NS
Log2 FC
p−value
p − value and log2 FC
FDR−corrected permutation p−values

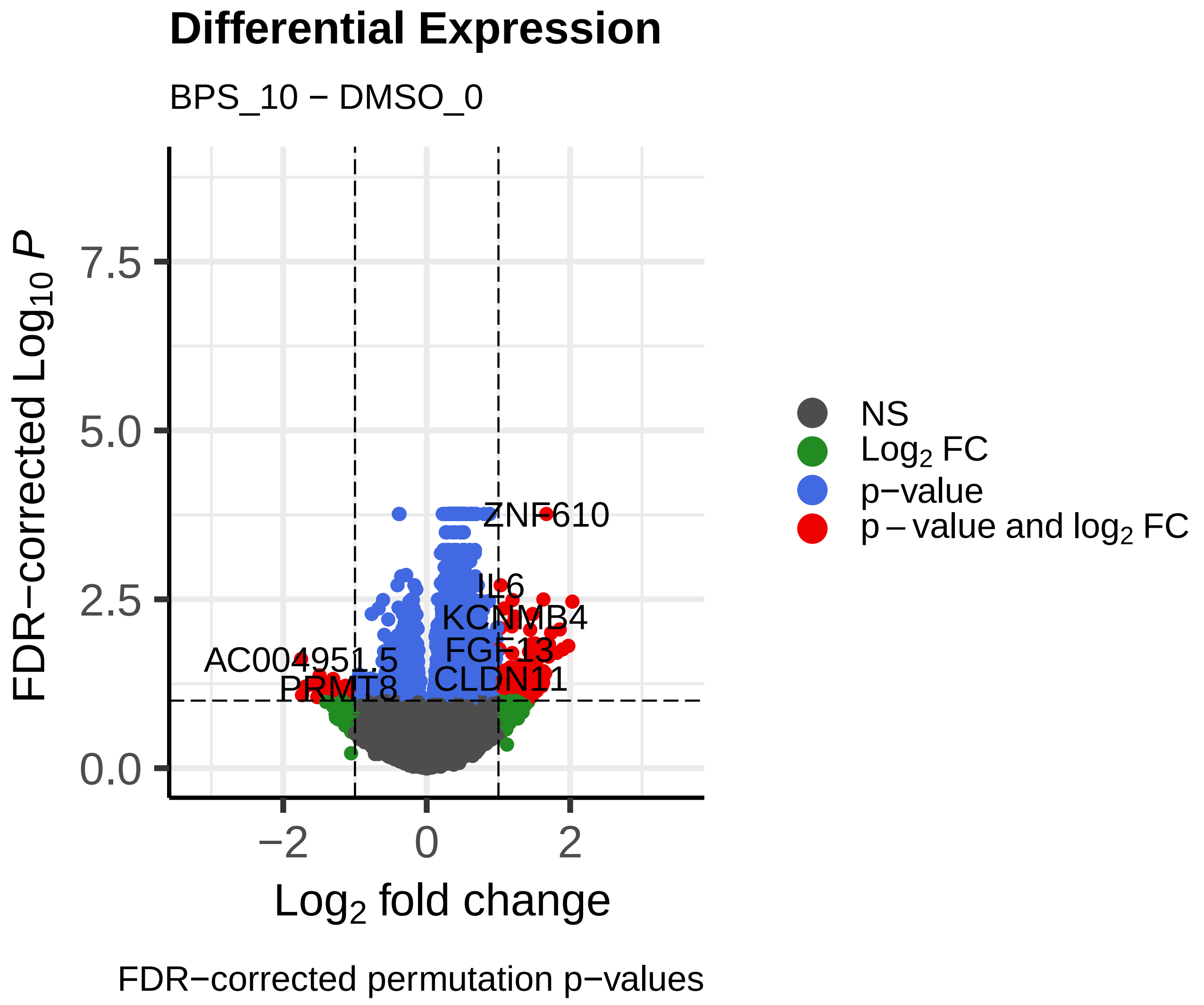
Differential Expression
BPS_10 − DMSO_0
FDR−corrected Log10 P
Log2 fold change
ZNF610
IL6
KCNMB4
AC004951.5
FGF13
PRMT8
CLDN11
NS
Log2 FC
p−value
p − value and log2 FC
FDR−corrected permutation p−values

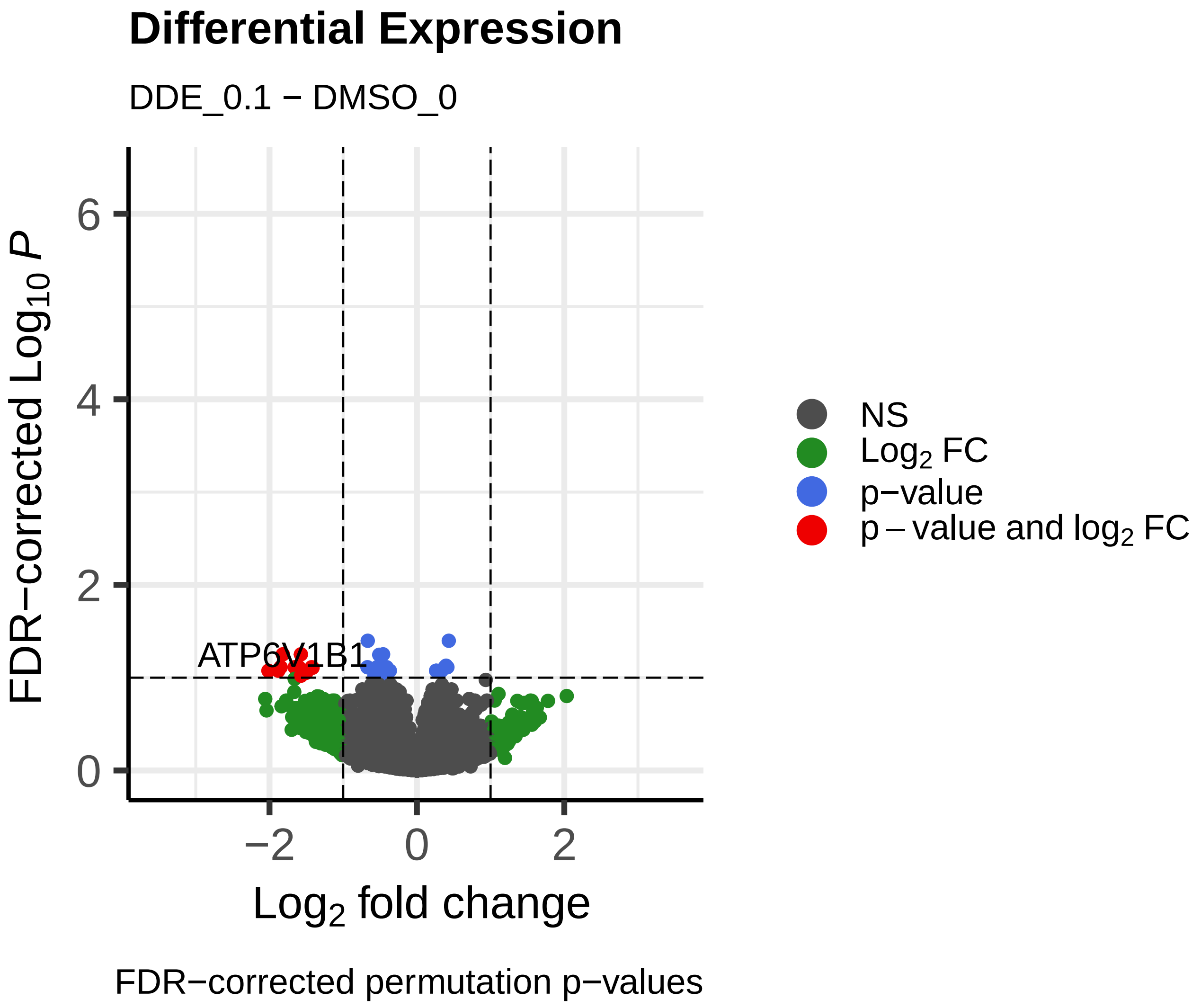
Differential Expression
DDE_0.1 − DMSO_0
FDR−corrected Log10 P
Log2 fold change
ATP6V1B1
NS
Log2 FC
p−value
p − value and log2 FC
FDR−corrected permutation p−values

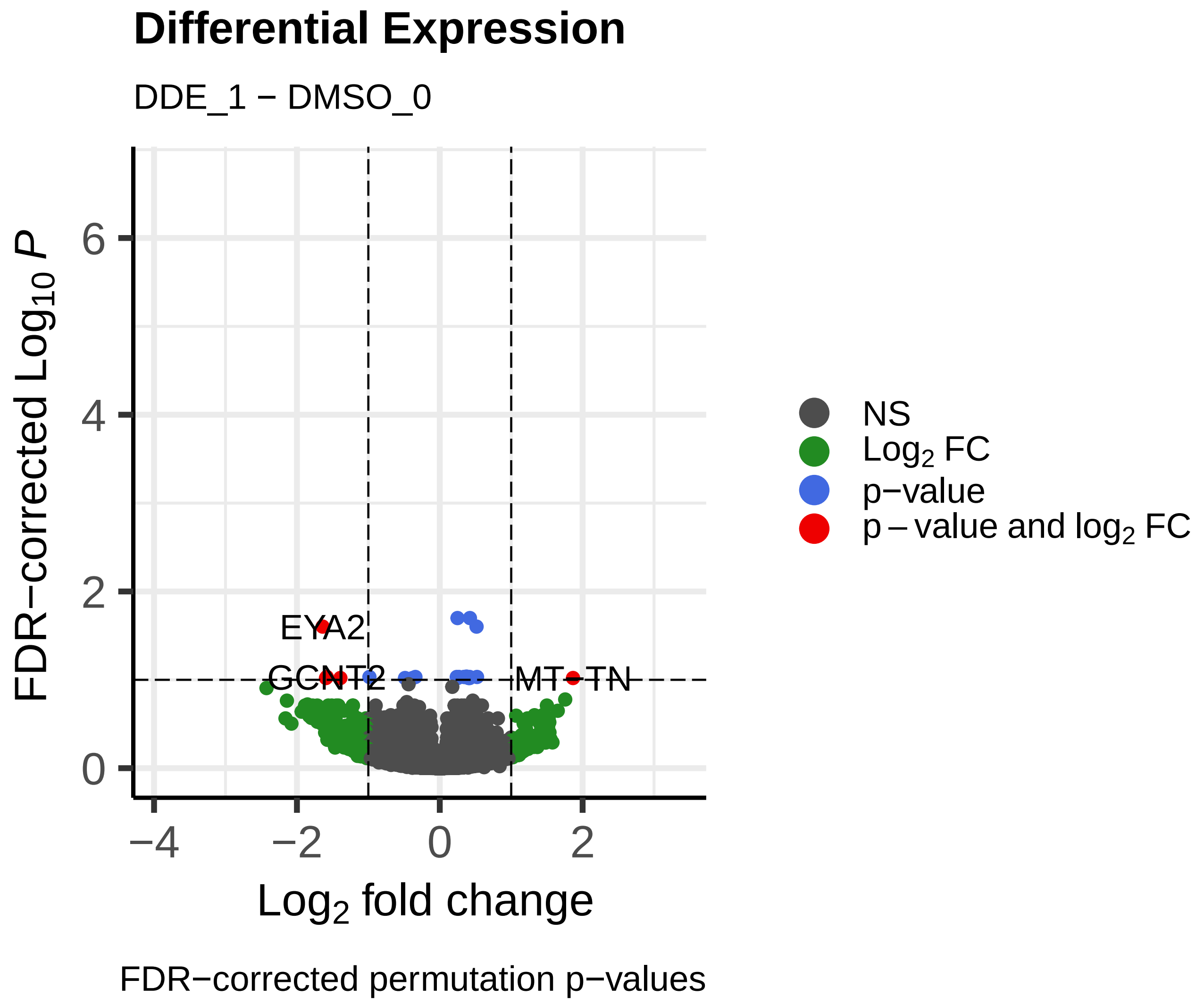
Differential Expression
DDE_1 − DMSO_0
FDR−corrected Log10 P
Log2 fold change
EYA2
GCNT2
MT−TN
NS
Log2 FC
p−value
p − value and log2 FC
FDR−corrected permutation p−values

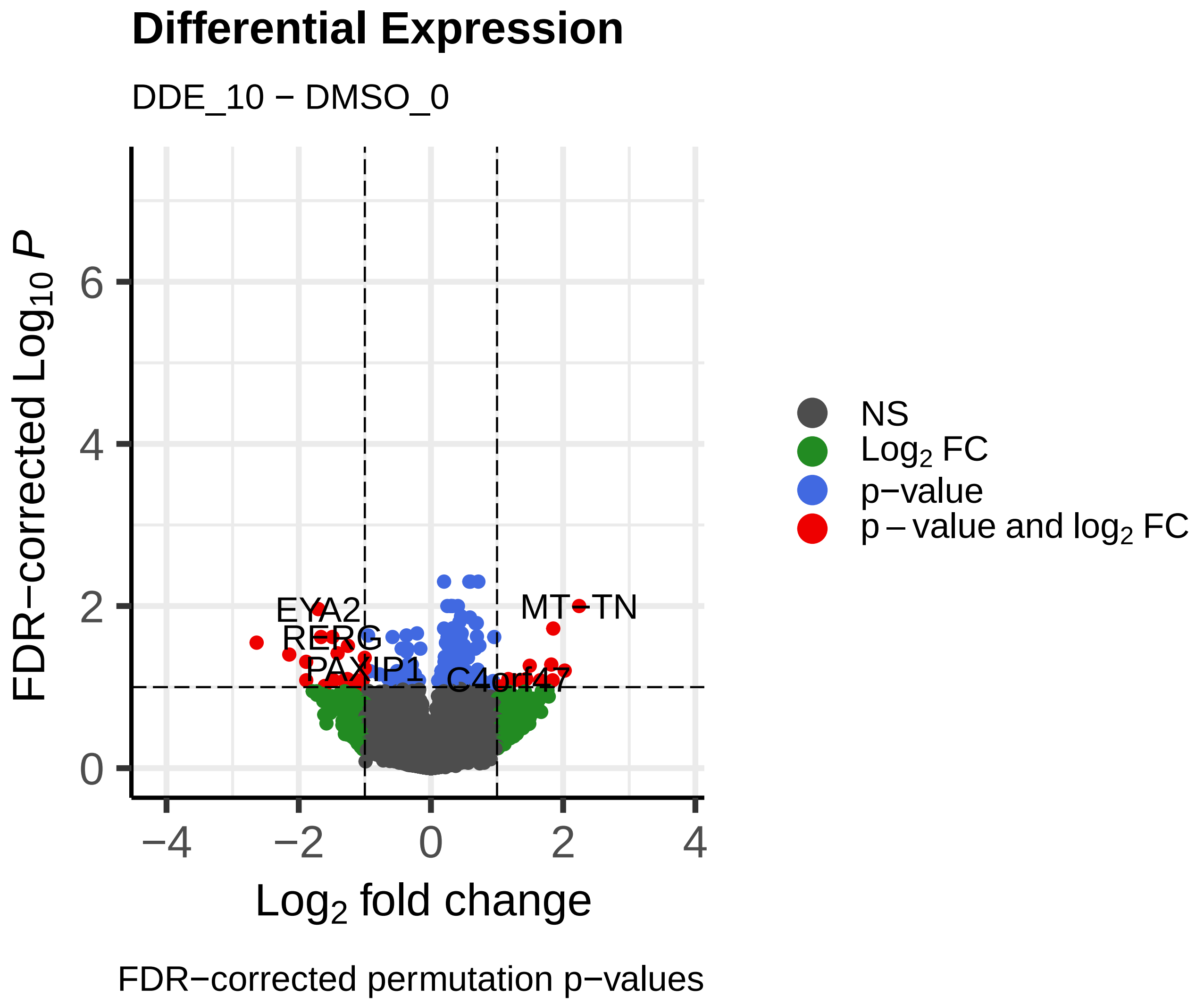
Differential Expression
DDE_10 − DMSO_0
FDR−corrected Log10 P
Log2 fold change
EYA2
RERG
PAXIP1
MT−TN
NS
Log2 FC
p−value
p − value and log2 FC
FDR−corrected permutation p−values

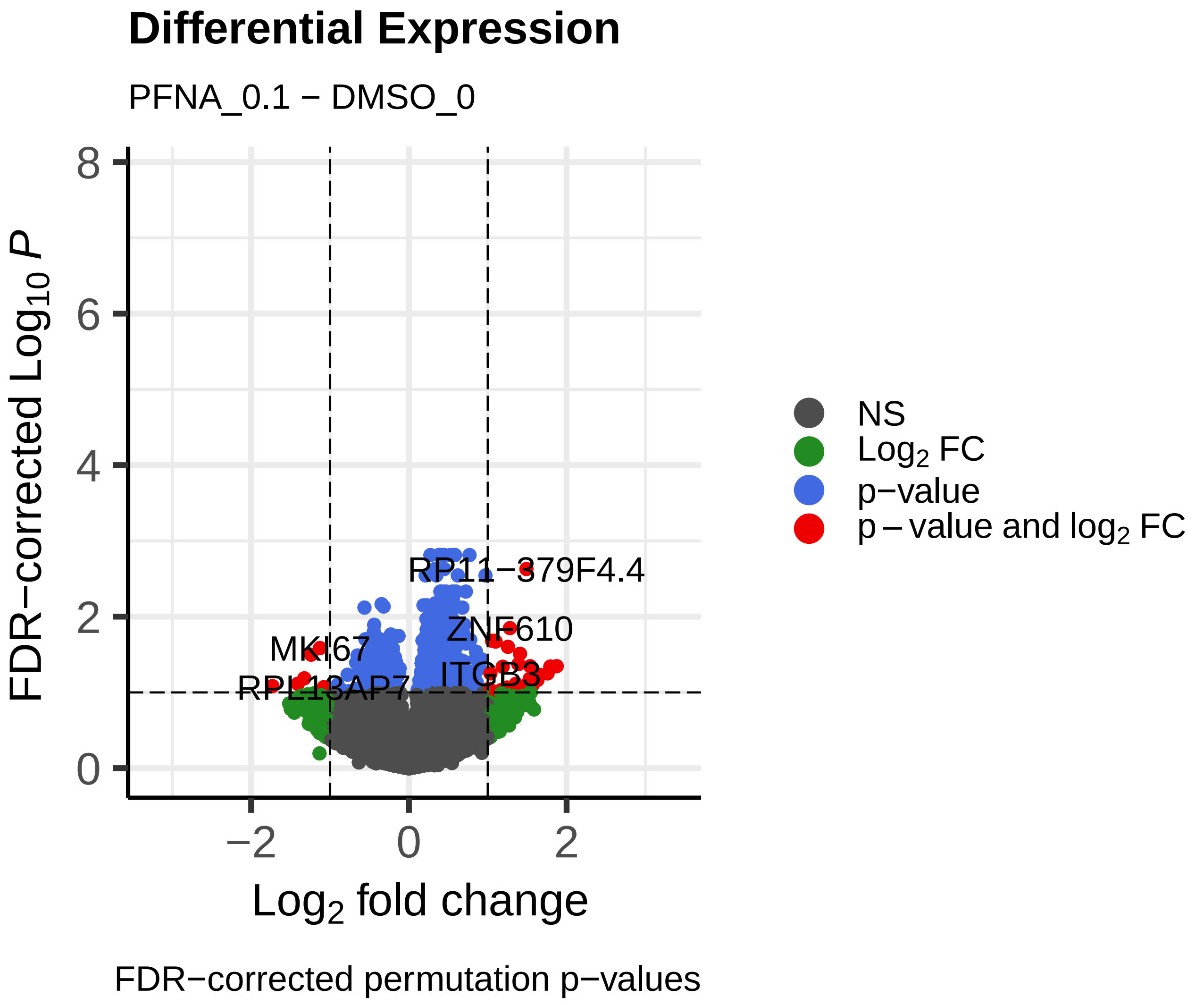
Differential Expression
PFNA_0.1 − DMSO_0
FDR−corrected Log10 P
Log2 fold change
NS
Log2 FC
p−value
p − value and log2 FC
RP11−379F4.4
ZNF610
MKI67
ITGB3
RPL13AP7
FDR−corrected permutation p−values

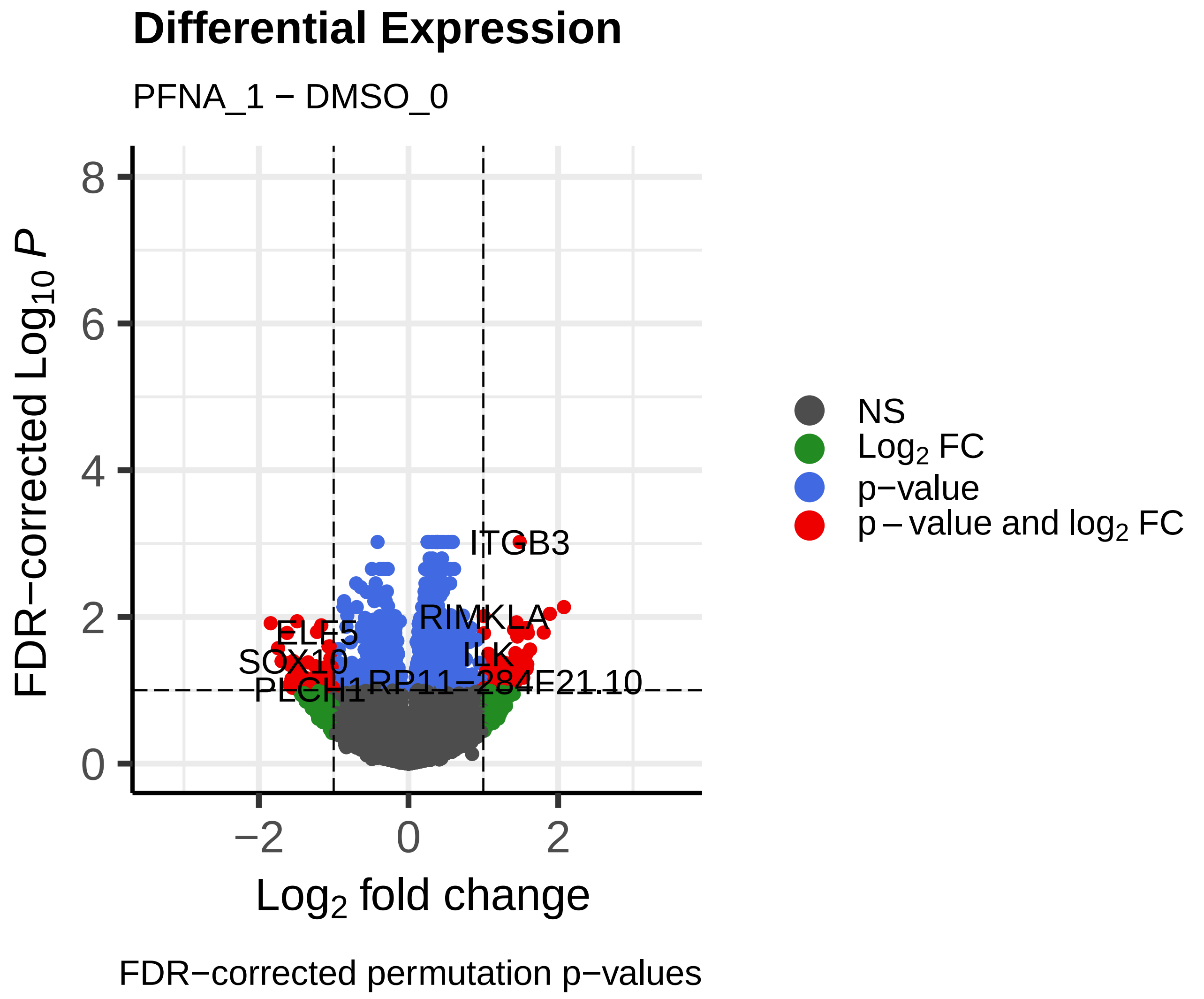
Differential Expression
PFNA_1 − DMSO_0
FDR−corrected Log10 P
Log2 fold change
NS
Log2 FC
p−value
p − value and log2 FC
ITGB3
RIMKLA
ELF5
SOX10
PLCH1
RP11−284F21.10
FDR−corrected permutation p−values

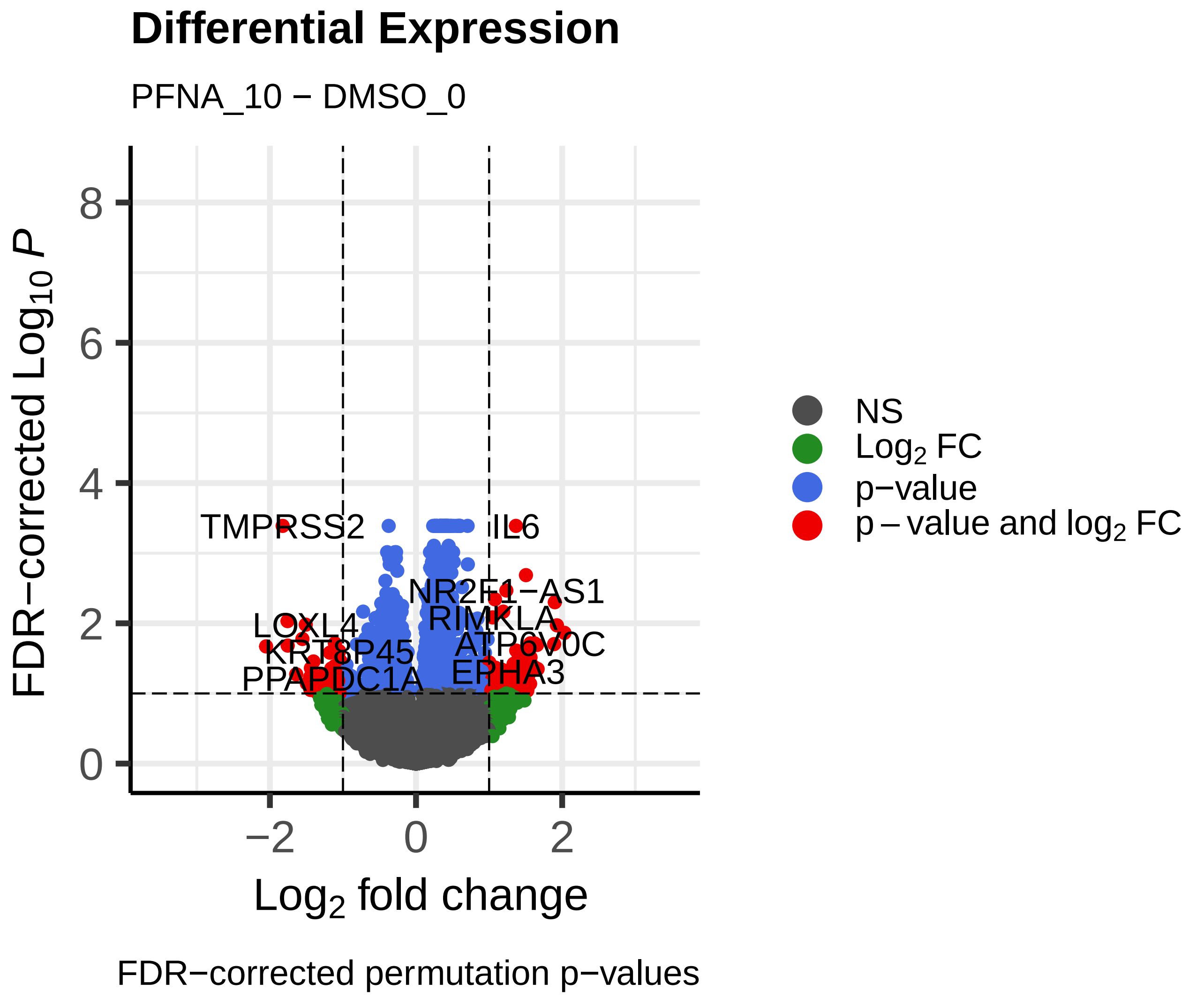
Differential Expression
PFNA_10 − DMSO_0
FDR−corrected Log10 P
Log2 fold change
NS
Log2 FC
p−value
p − value and log2 FC
TMPRSS2
IL6
NR2F1−AS1
LOXL4
RIMKLA
ATP6V0C
PPAPDC1A
EPHA3
FDR−corrected permutation p−values

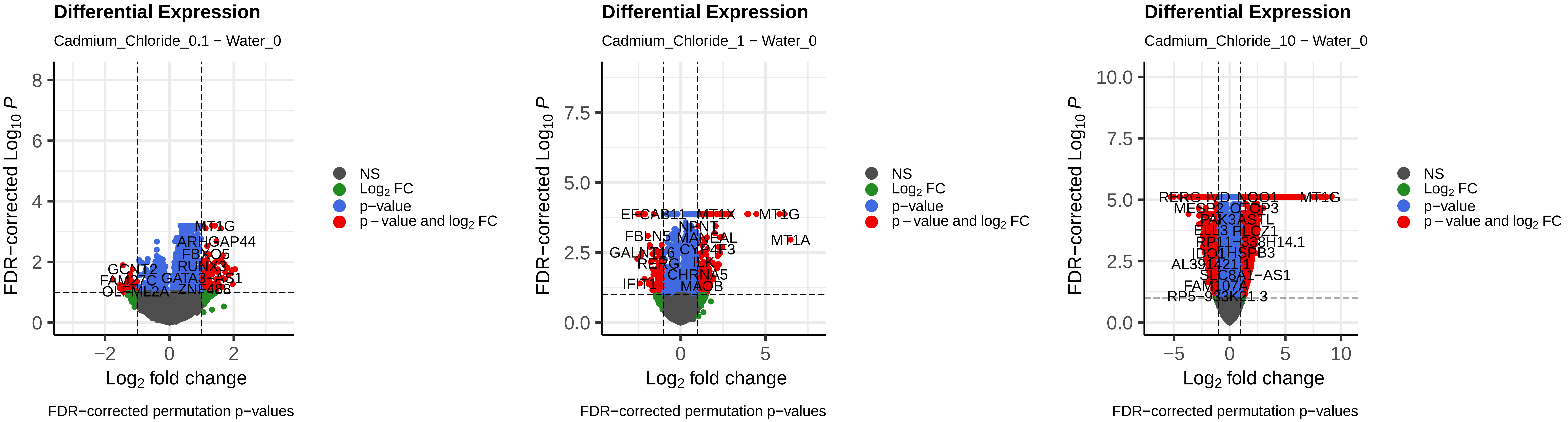
Differential Expression
Cadmium_Chloride_0.1 − Water_0
FDR−corrected Log10 P
Log2 fold change
MT1G
ARHGAP44
FBXO5
RUNX3
GATA3−AS1
FDR−corrected permutation p−values
Differential Expression
Cadmium_Chloride_1 − Water_0
FDR−corrected Log10 P
Log2 fold change
EFCAB11
MT1X
MT1G
MT1A
NPNT
MANEAL
CYP4F3
RERG
CHRNA5
MAOB
FDR−corrected permutation p−values
Differential Expression
Cadmium_Chloride_10 − Water_0
FDR−corrected Log10 P
Log2 fold change
MT1G
RP11−338H14.1
SLC8A1−AS1
FDR−corrected permutation p−values
NS
Log2 FC
p−value
p − value and log2 FC

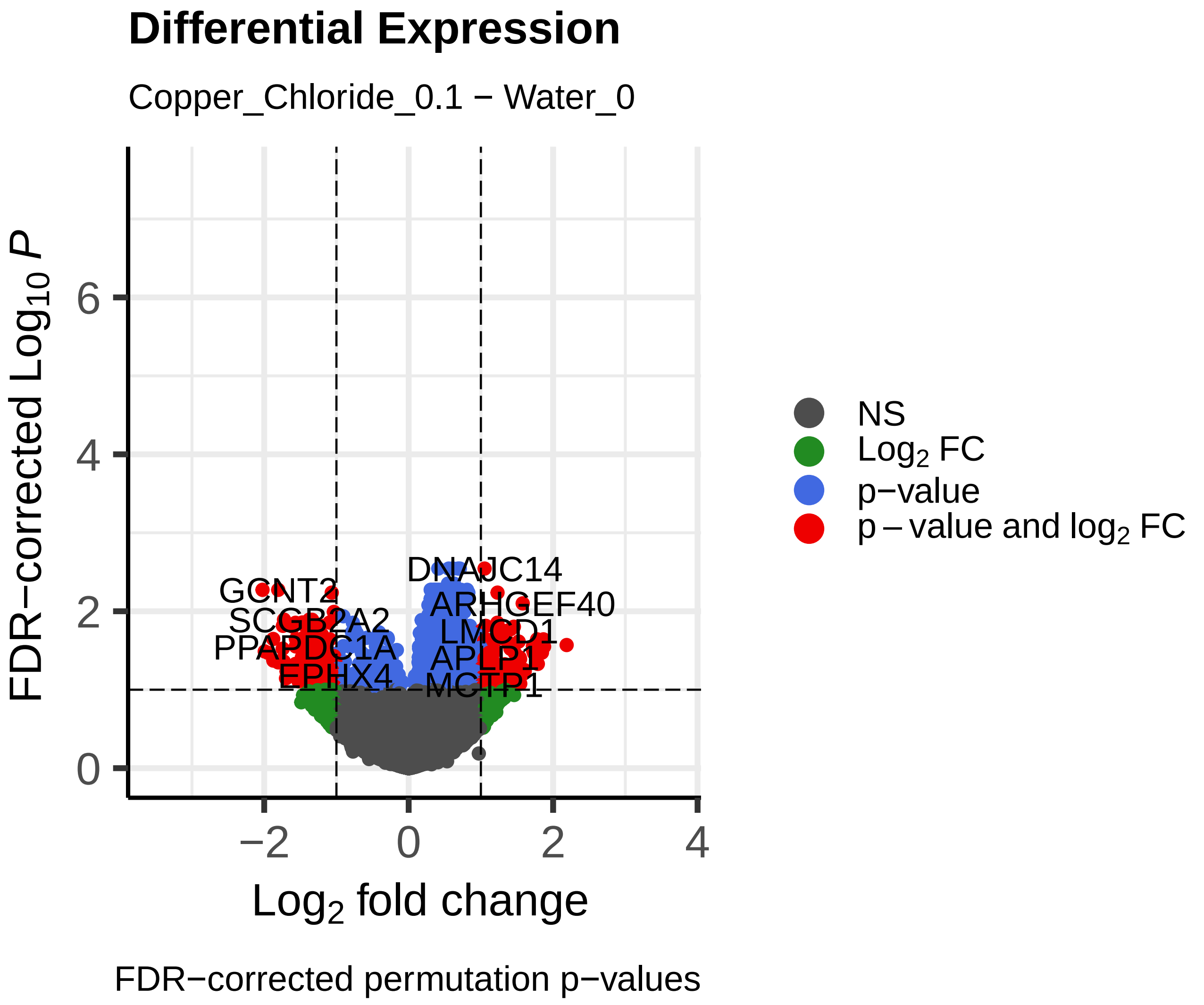
Differential Expression
Copper_Chloride_0.1 − Water_0
FDR−corrected Log10 P
Log2 fold change
NS
Log2 FC
p−value
p − value and log2 FC
FDR−corrected permutation p−values

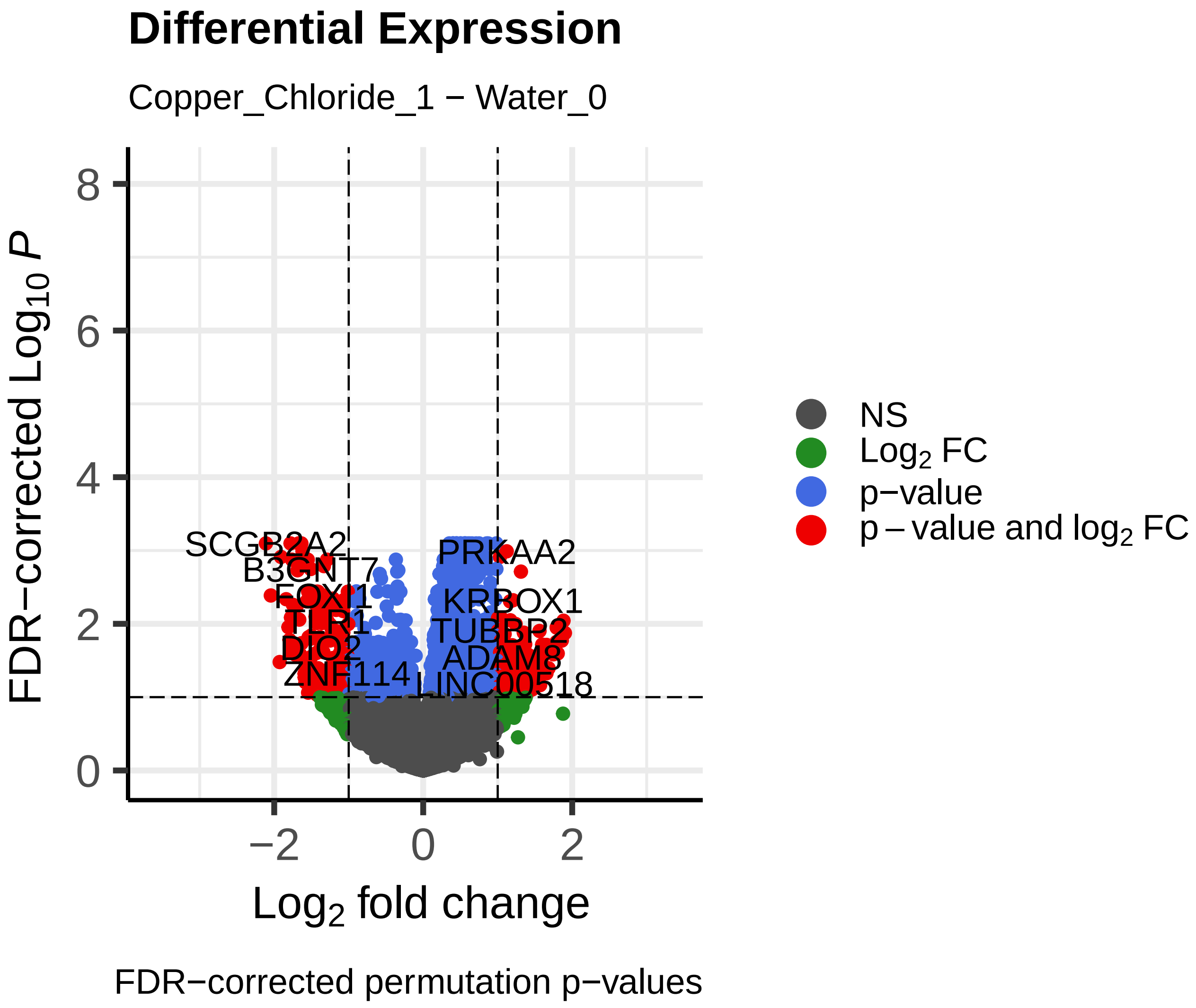
Differential Expression
Copper_Chloride_1 − Water_0
FDR−corrected Log10 P
Log2 fold change
NS
Log2 FC
p−value
p − value and log2 FC
FDR−corrected permutation p−values

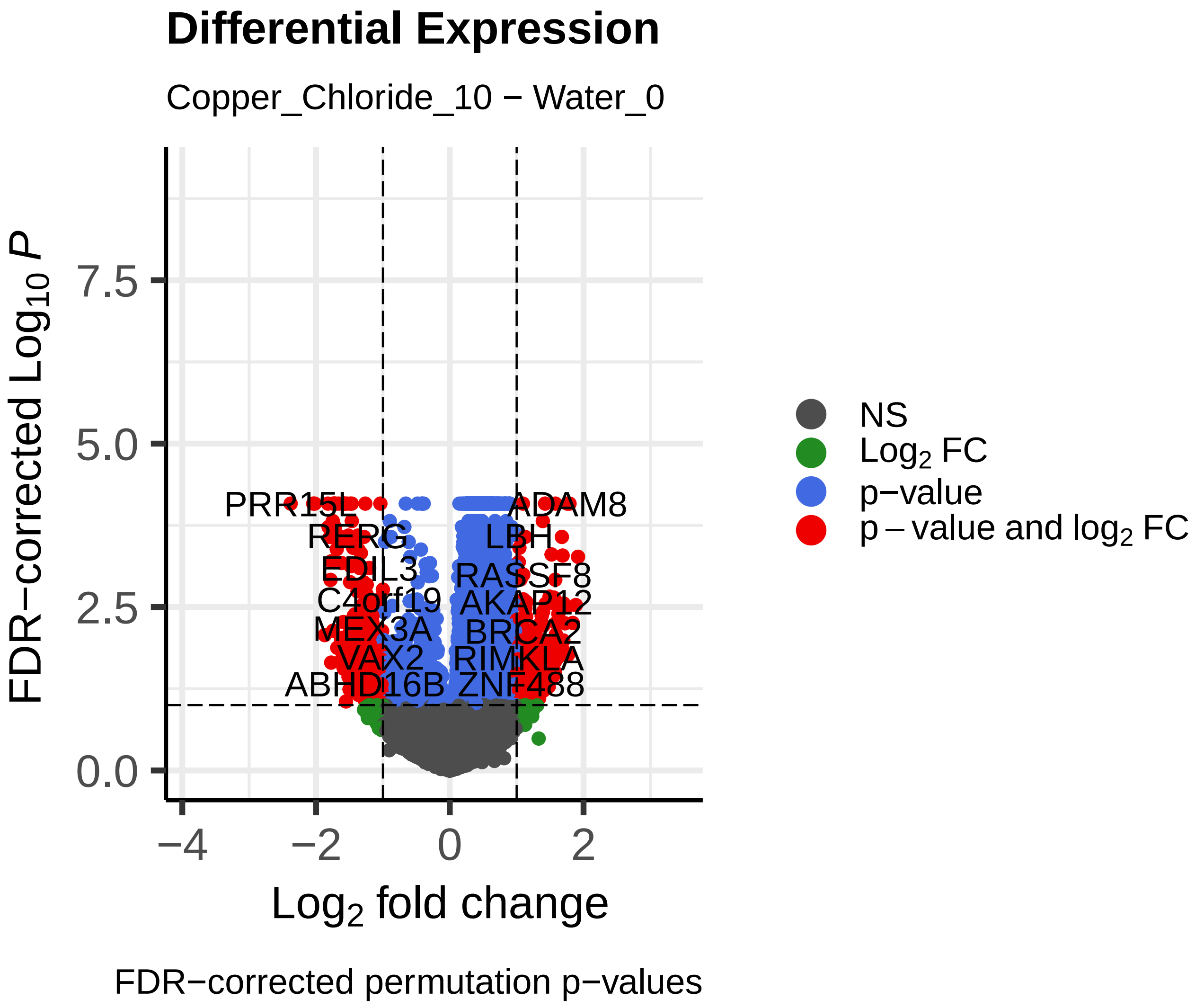
Differential Expression
Copper_Chloride_10 − Water_0
FDR−corrected Log10 P
Log2 fold change
NS
Log2 FC
p−value
p − value and log2 FC
FDR−corrected permutation p−values

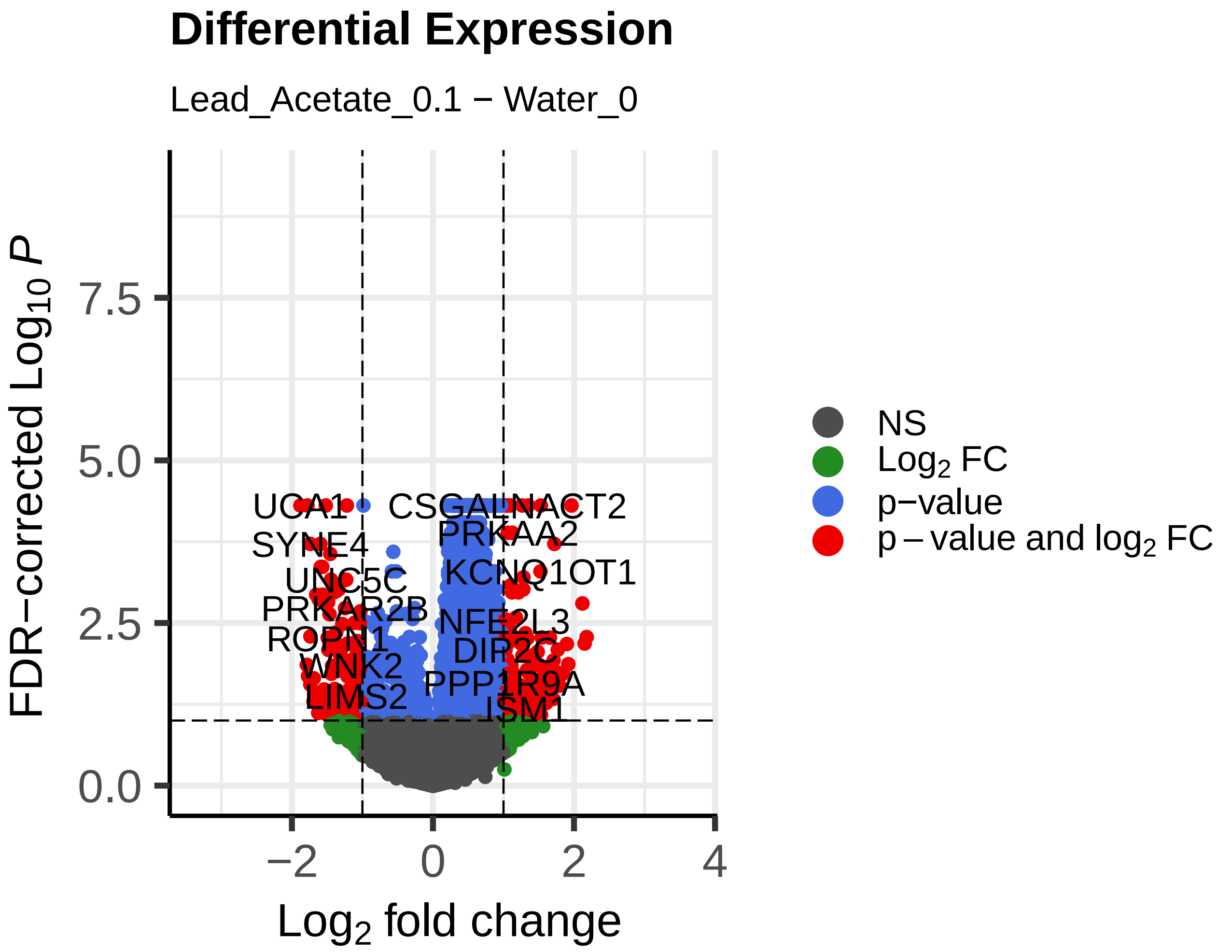


FDR−corrected permutation p−values

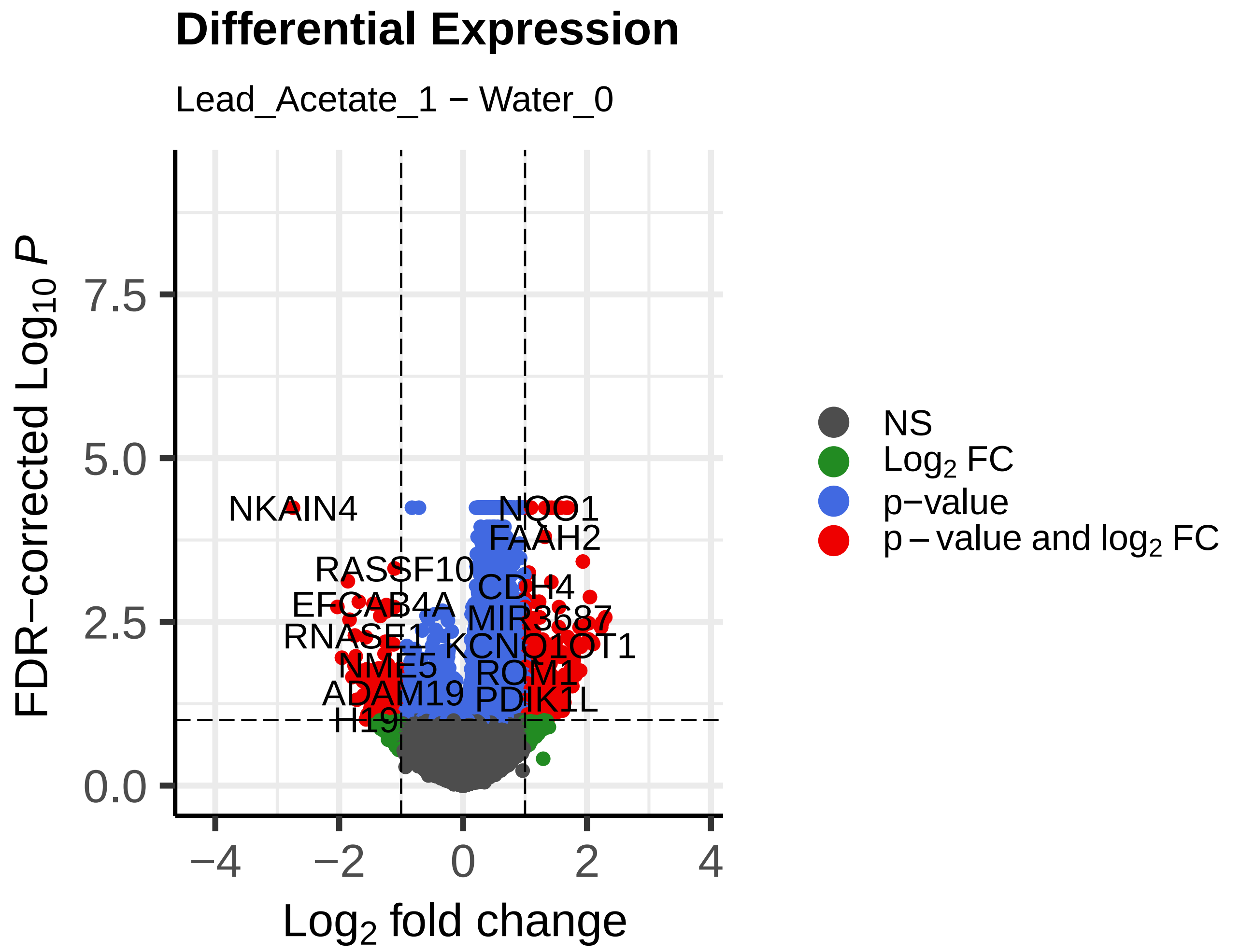


FDR−corrected permutation p−values

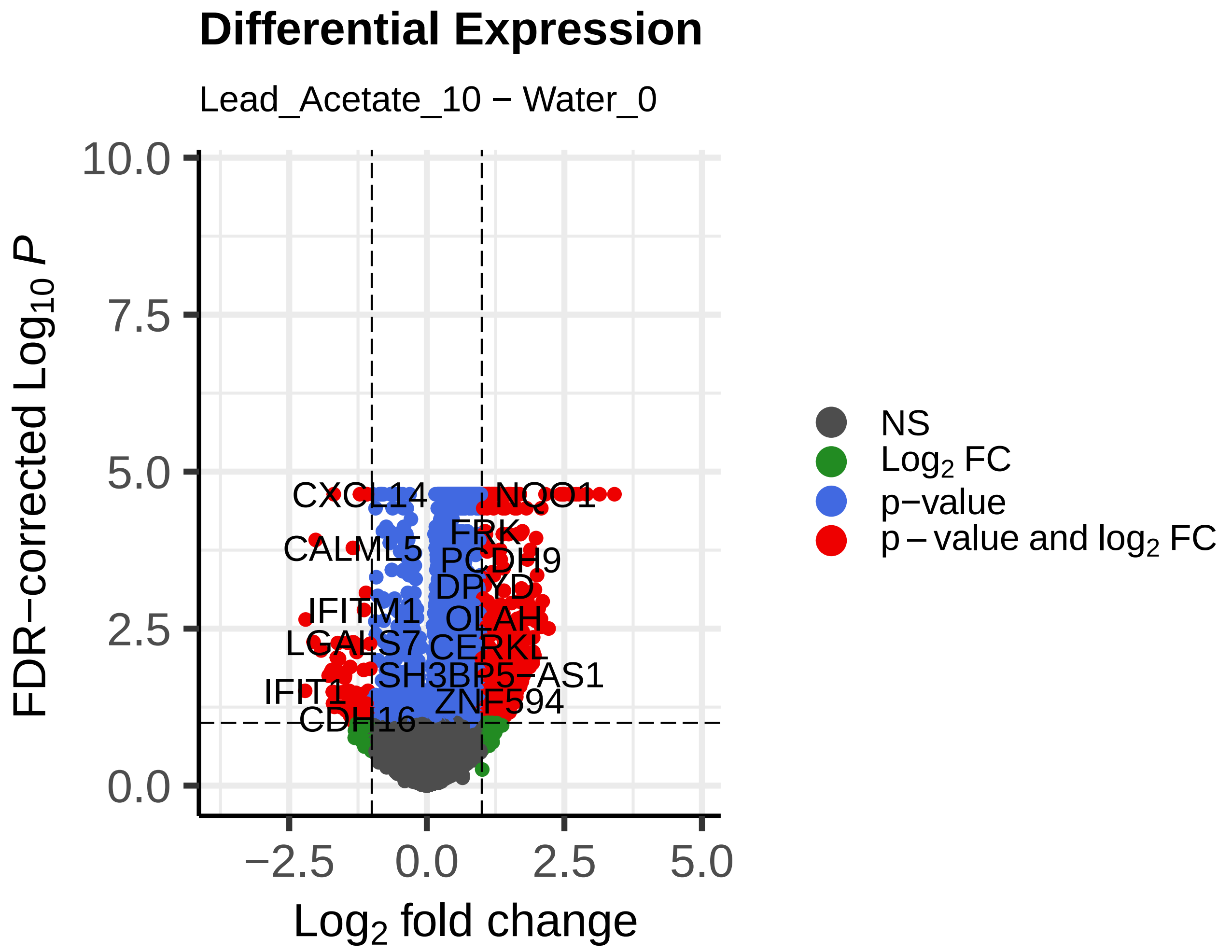


FDR−corrected permutation p−values

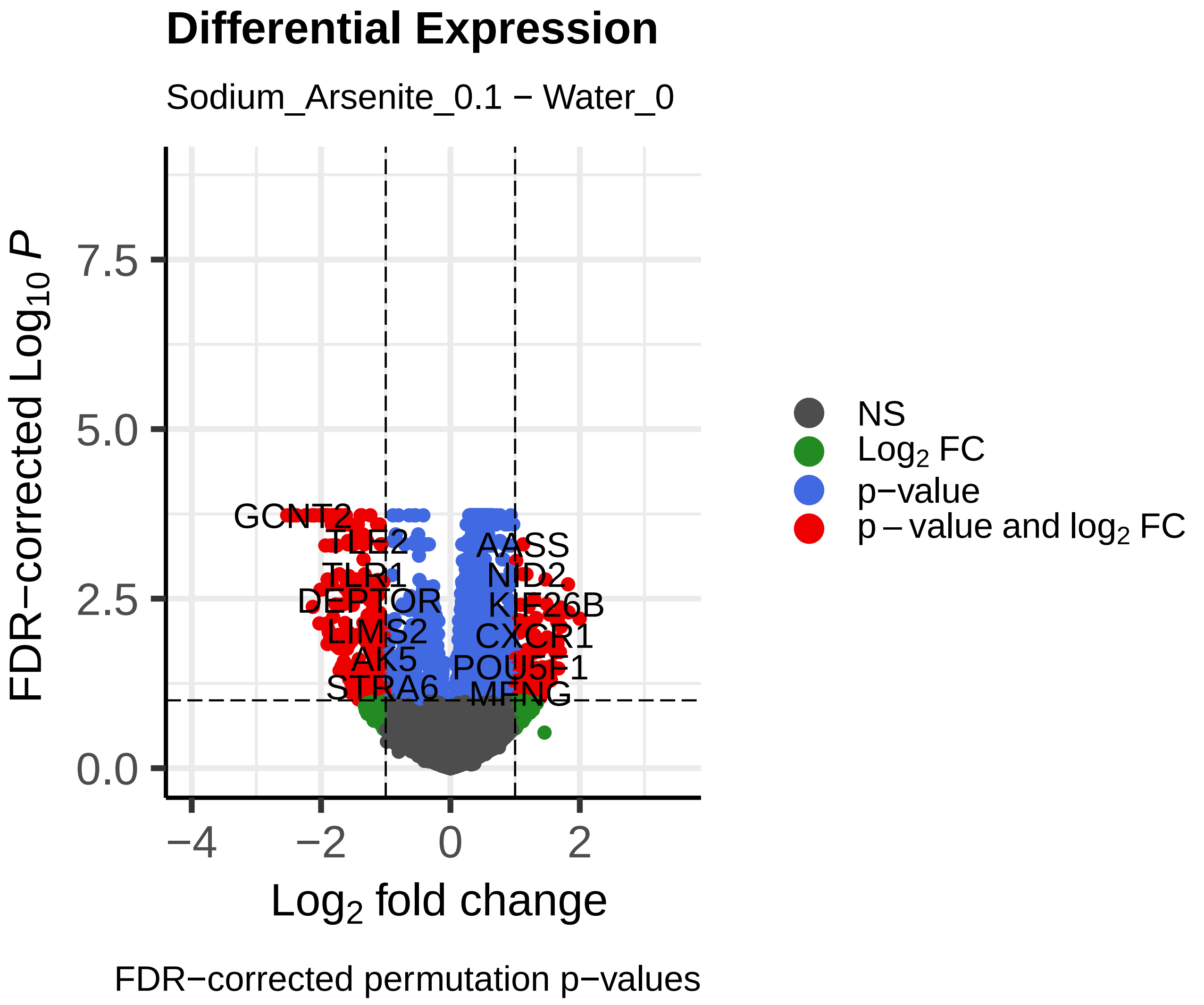
Differential Expression
Sodium_Arsenite_0.1 − Water_0
FDR−corrected $Log_{10}$ $P$
Log2 fold change
GCNT2
TLE2
AASS
TLR1
NID2
DEPTOR
KIF26B
LIMS2
CXCR1
AK5
POU5F1
STRA6
MFNG
NS
Log2 FC
p−value
p – value and log2 FC
FDR−corrected permutation p−values

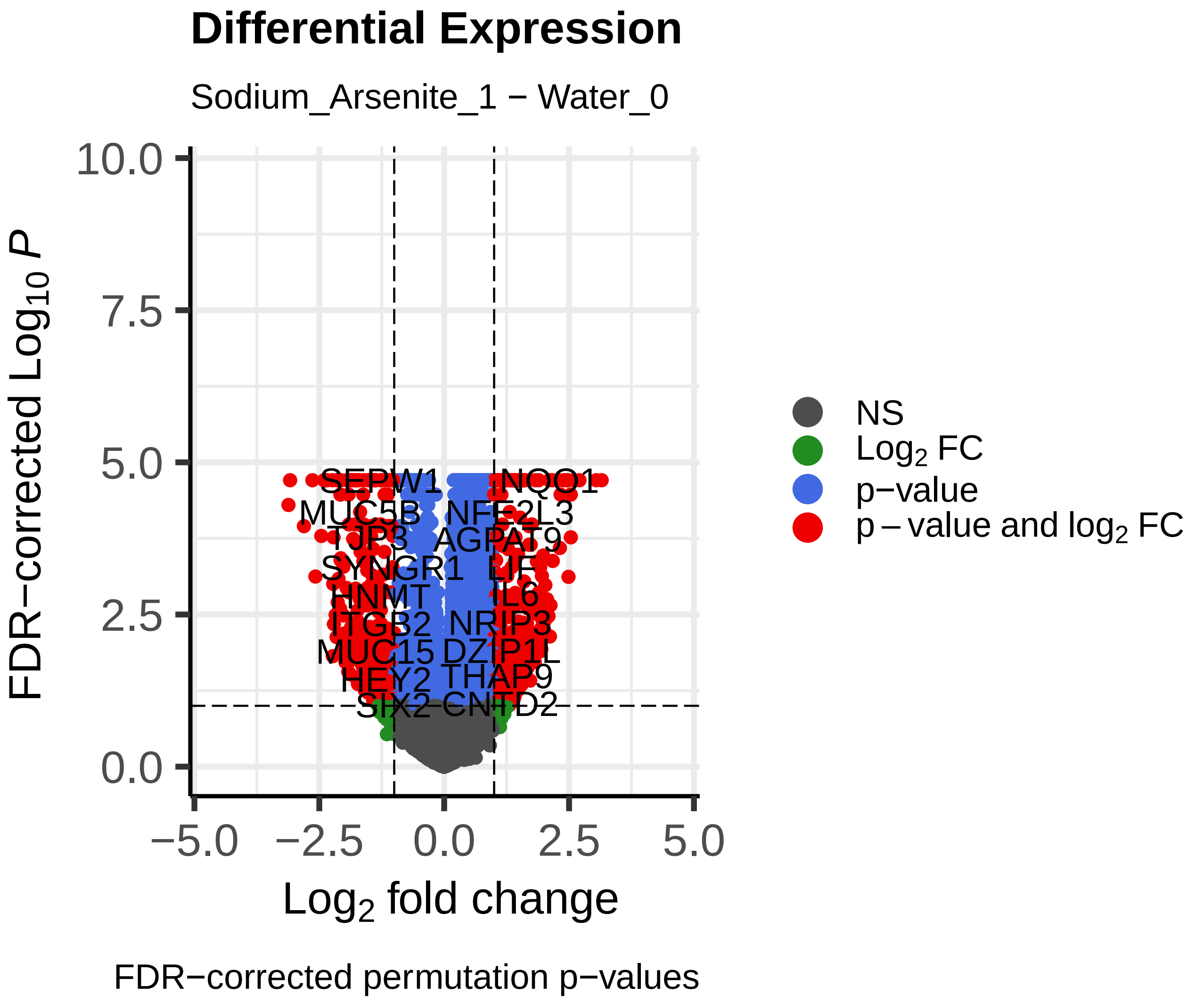
Differential Expression
Sodium_Arsenite_1 − Water_0
FDR−corrected $Log_{10}$ $P$
Log2 fold change
SEPW1
NQO1
MUC5B
NFE2L3
AGPAT9
SYNGR1
LIF
HNMT
IL6
NRIP3
MUC15
DZIP1L
HEY2
THAP9
SIX2
CNTD2
NS
Log2 FC
p−value
p – value and log2 FC
FDR−corrected permutation p−values

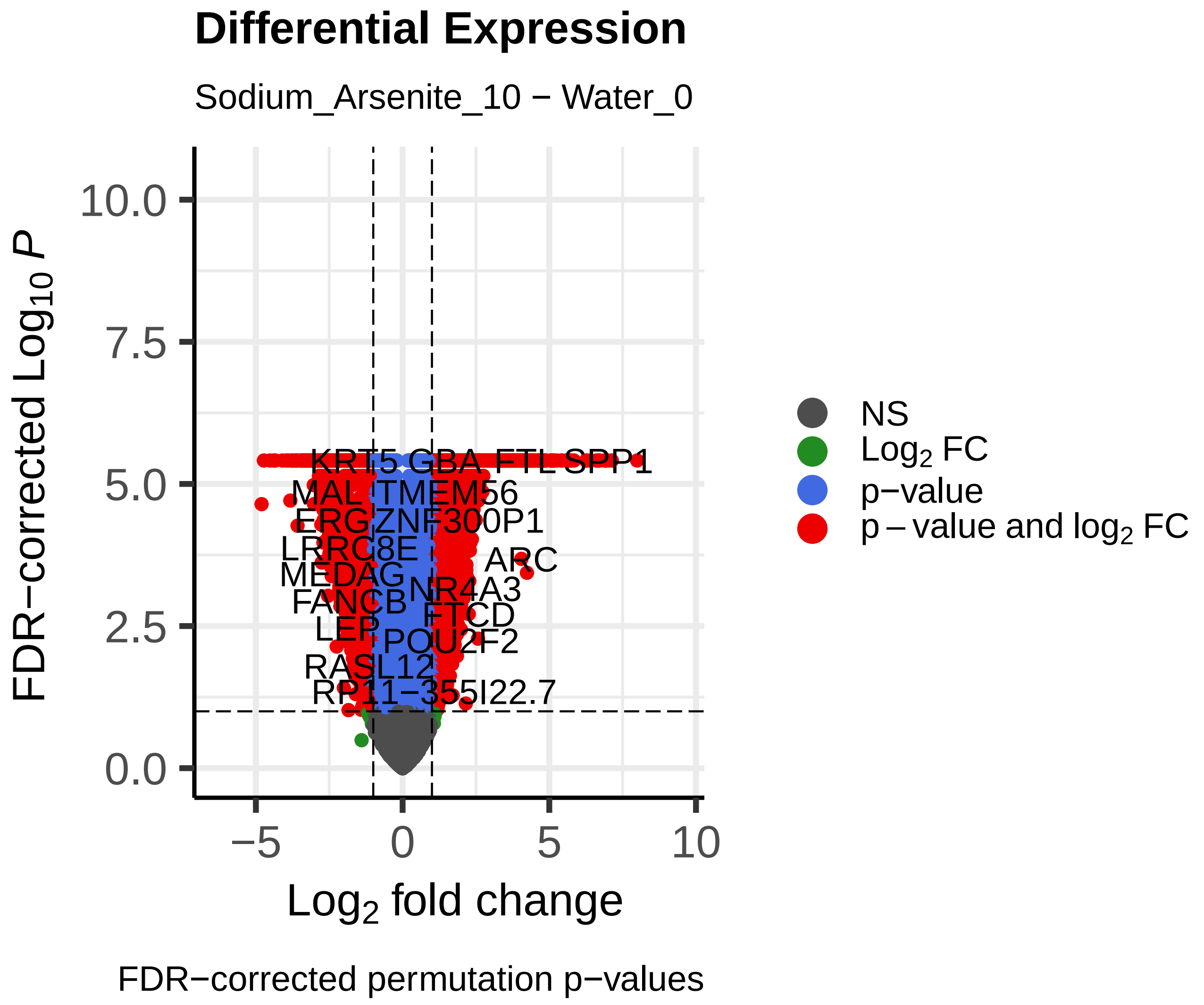
Differential Expression
Sodium_Arsenite_10 − Water_0
FDR−corrected $Log_{10}$ $P$
Log2 fold change
KRT5
GBA
FTL
SPP1
MAL
TMEM56
ERG
ZNF300P1
LRRC8E
ARC
MEDAG
NR4A3
FANCB
FTCD
LEP
POU2F2
RASL12
RP11−355I22.7
NS
Log2 FC
p−value
p – value and log2 FC
FDR−corrected permutation p−values
