## Supplementary material for "Exploring the Influence of Chemical Exposures in Breast Cancer Disparities: High-Throughput Transcriptomic Analysis in Normal Breast Cells from Diverse Donors": Figure S-3

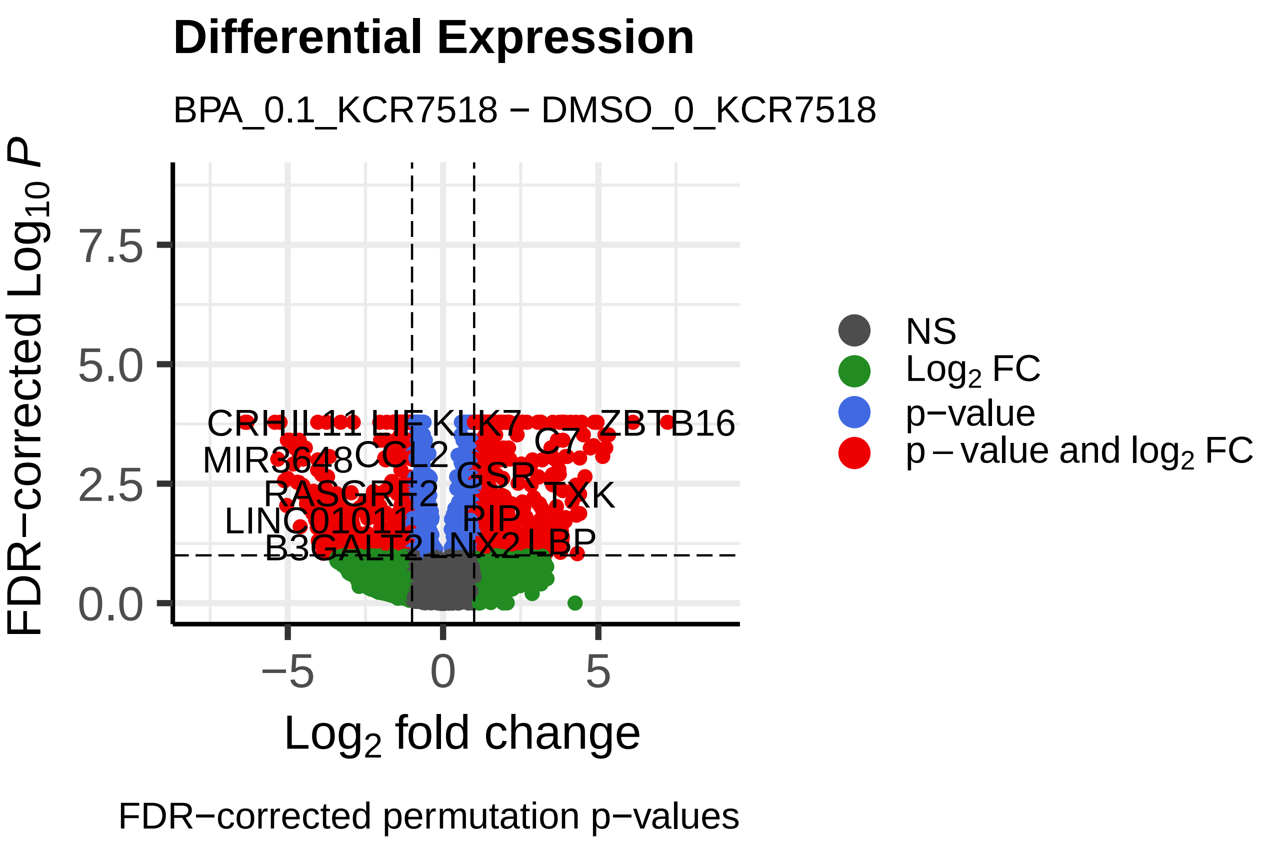
Differential Expression
BPA_0.1_KCR7518 − DMSO_0_KCR7518
FDR−corrected $Log_{10}$ $P$
Log₂ fold change
NS
Log₂ FC
p-value
p − value and log₂ FC
CRHR1
IL11
LIF
KLK7
ZBTB16
MIR3648
CCL2
C7
GSR
RASGRF2
TXK
LINC01011
PIP
B3GALT2
LNX2
LBP
FDR−corrected permutation p−values

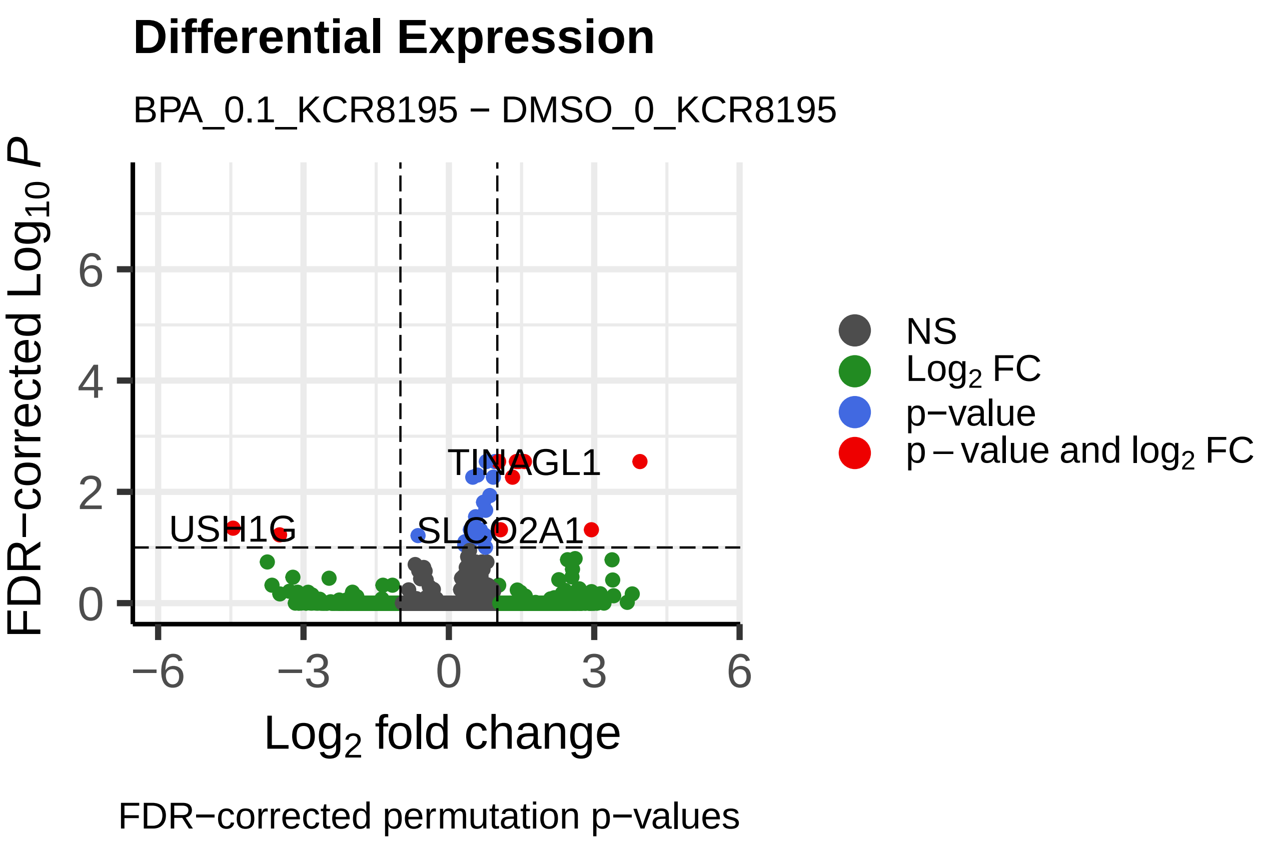
Differential Expression
BPA_0.1_KCR8195 − DMSO_0_KCR8195
FDR−corrected $Log_{10}$ $P$
Log₂ fold change
NS
Log₂ FC
p-value
p − value and log₂ FC
TINAGL1
USH1G
SLCO2A1
FDR−corrected permutation p−values

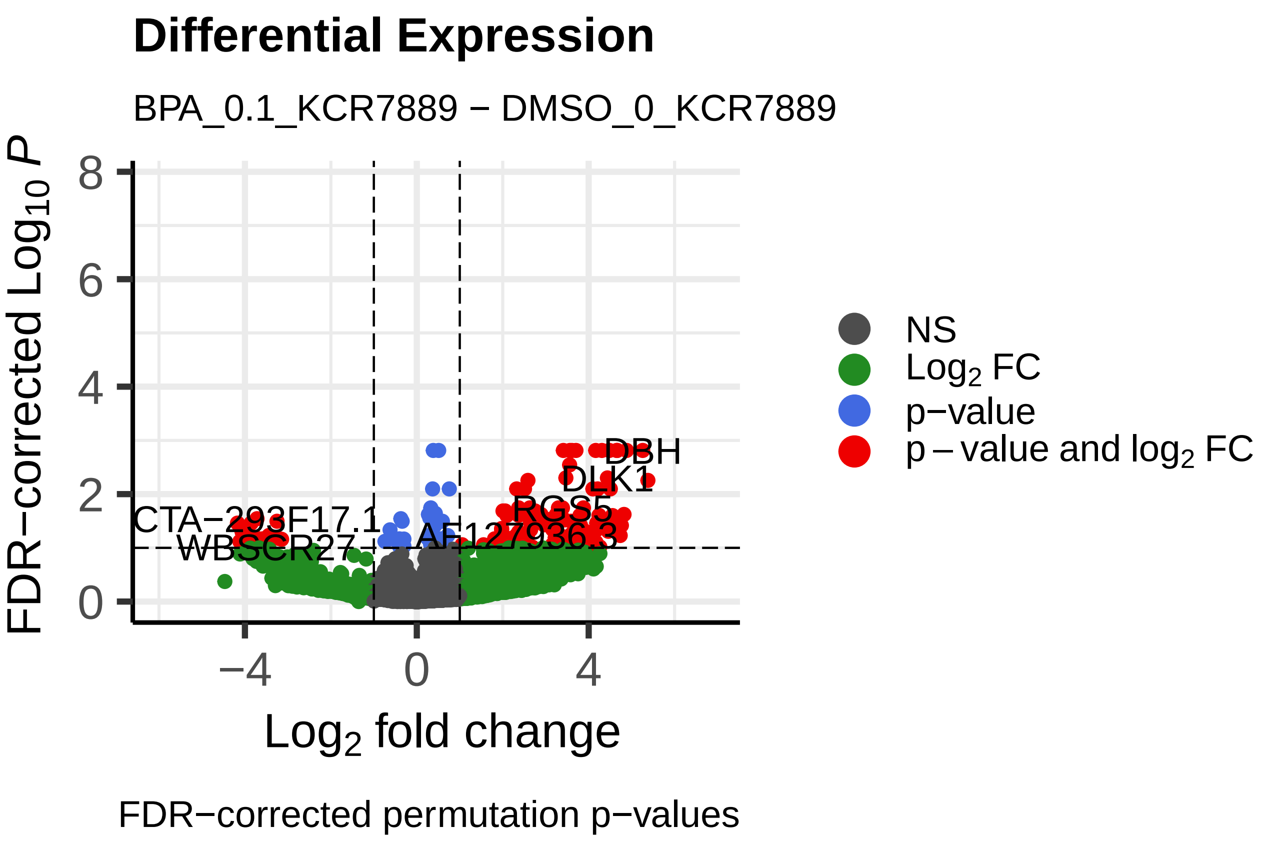
Differential Expression
BPA_0.1_KCR7889 − DMSO_0_KCR7889
FDR−corrected $Log_{10}$ $P$
Log₂ fold change
NS
Log₂ FC
p-value
p − value and log₂ FC
DBH
DLK1
RGS5
CTA−293F17.1
WBSCR27
AF127936.3
FDR−corrected permutation p−values

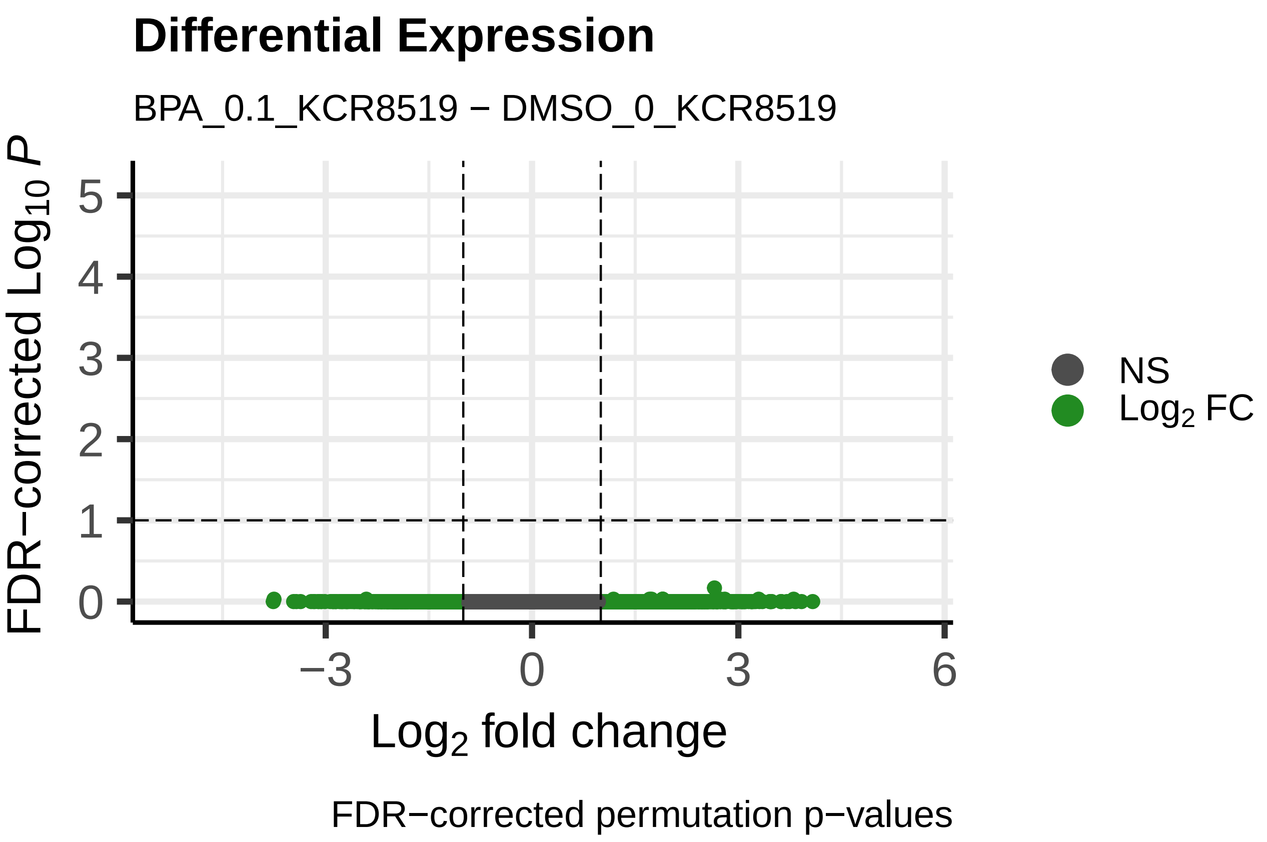
Differential Expression
BPA_0.1_KCR8519 − DMSO_0_KCR8519
FDR−corrected $Log_{10}$ $P$
Log₂ fold change
NS
Log₂ FC
FDR−corrected permutation p−values

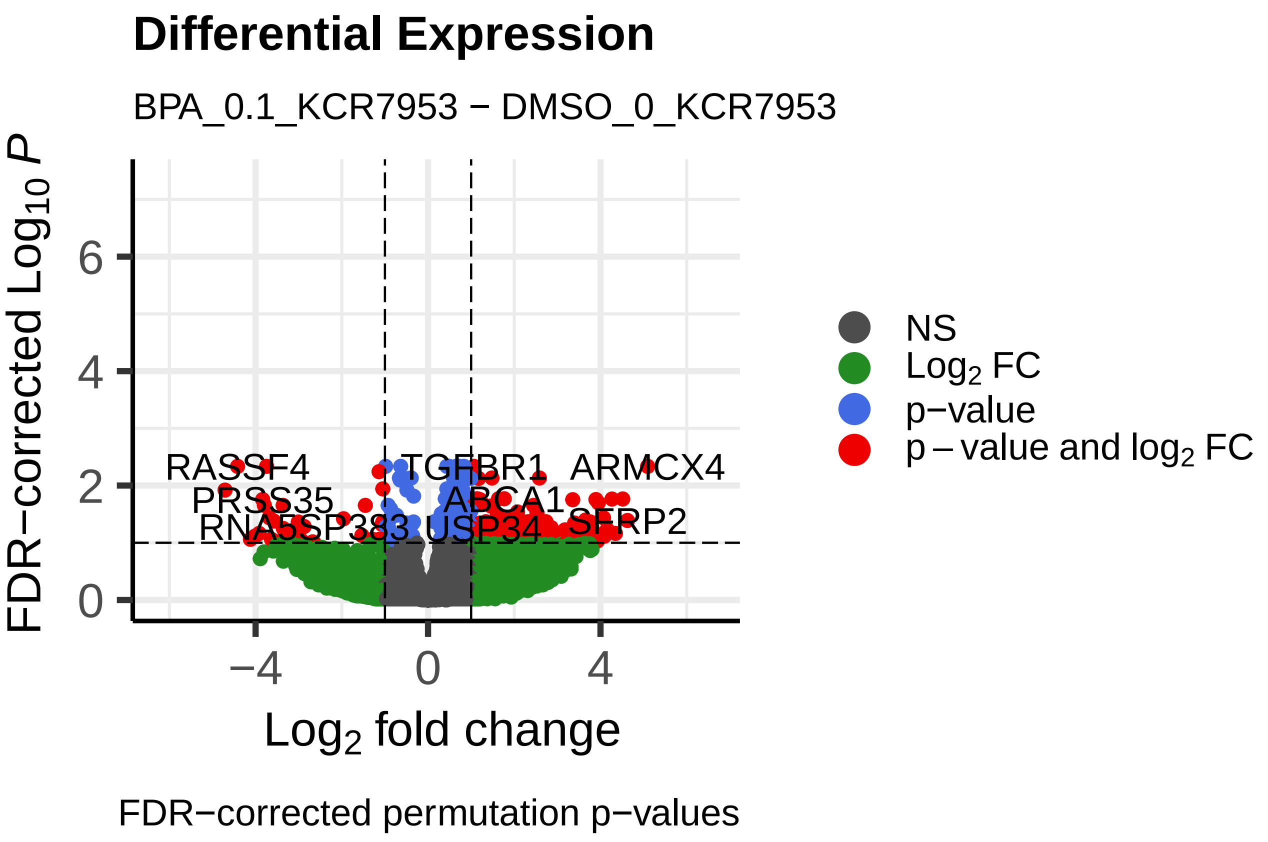
Differential Expression
BPA_0.1_KCR7953 − DMSO_0_KCR7953
FDR−corrected $Log_{10}$ $P$
Log₂ fold change
NS
Log₂ FC
p-value
p − value and log₂ FC
RASSF4
TGFBR1
ARMCX4
PRSS35
ABCA1
USP34
SFRP2
FDR−corrected permutation p−values

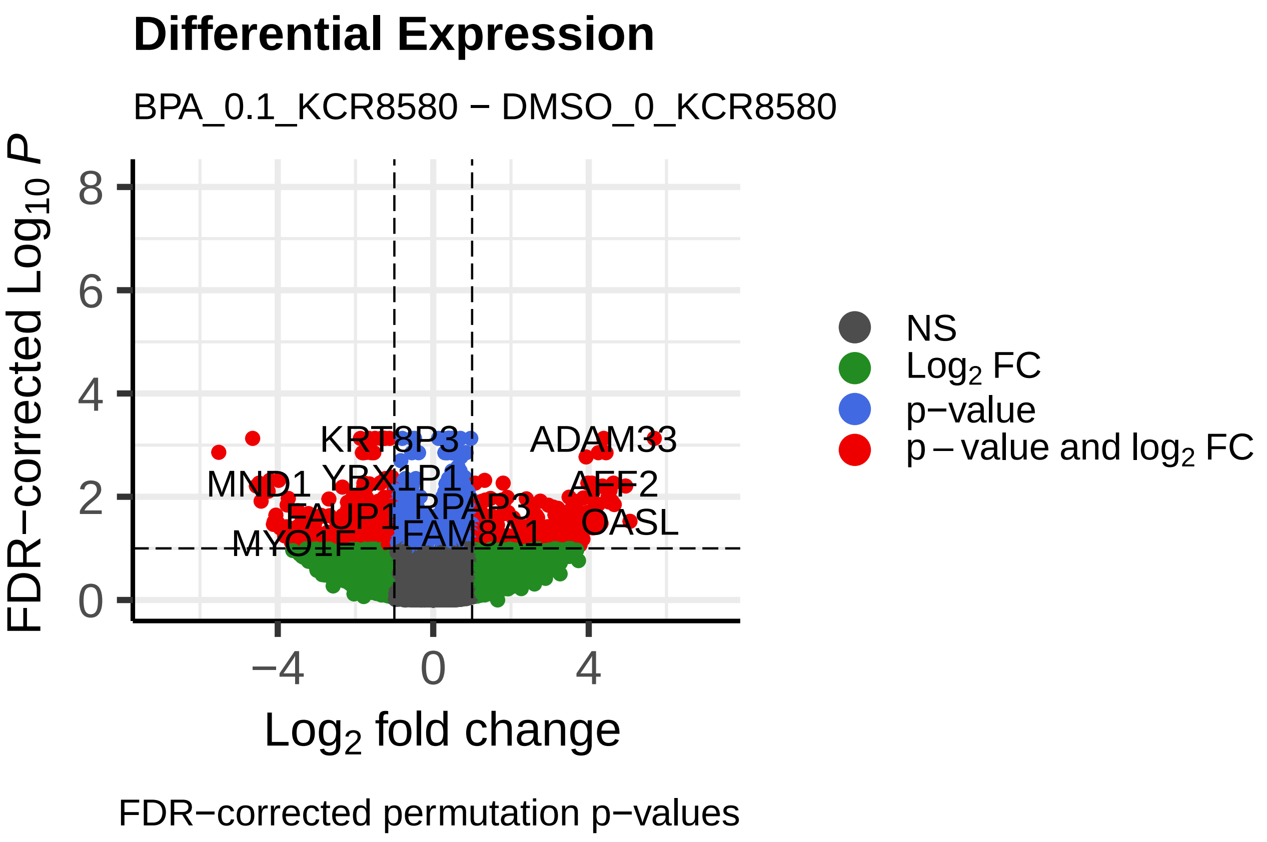
Differential Expression
BPA_0.1_KCR8580 − DMSO_0_KCR8580
FDR−corrected $Log_{10}$ $P$
Log₂ fold change
NS
Log₂ FC
p-value
p − value and log₂ FC
ADAM33
MND1
AFF2
FAUP1
OASL
MYO1F
FAM8A1
FDR−corrected permutation p−values

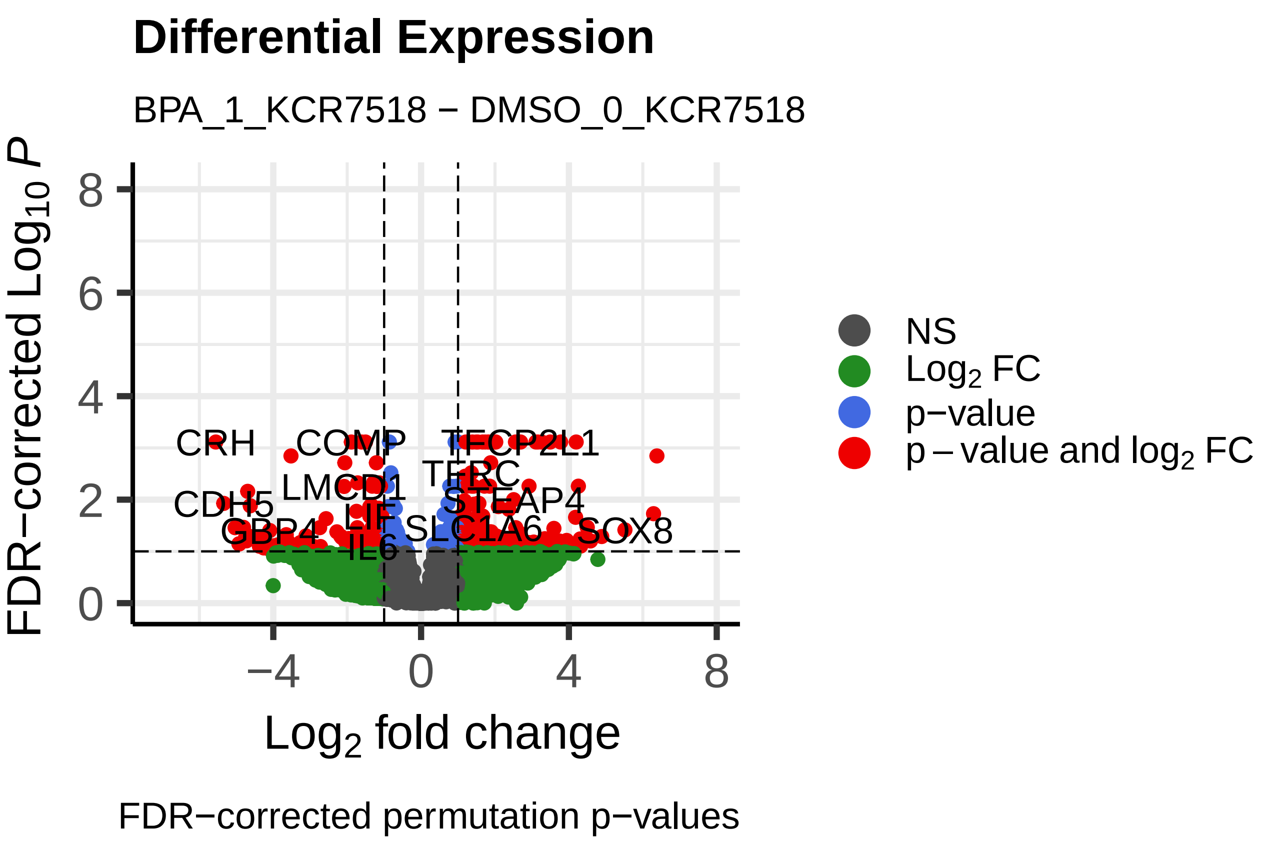
Differential Expression
BPA_1_KCR7518 − DMSO_0_KCR7518
FDR−corrected Log10 P
Log2 fold change
NS
Log2 FC
p−value
p − value and log2 FC
CRH
COMP
TFCP2L1
LMCD1
TERC
CDH5
STEAP4
GBP4
SLC1A6
SOX8
IL6
FDR−corrected permutation p−values

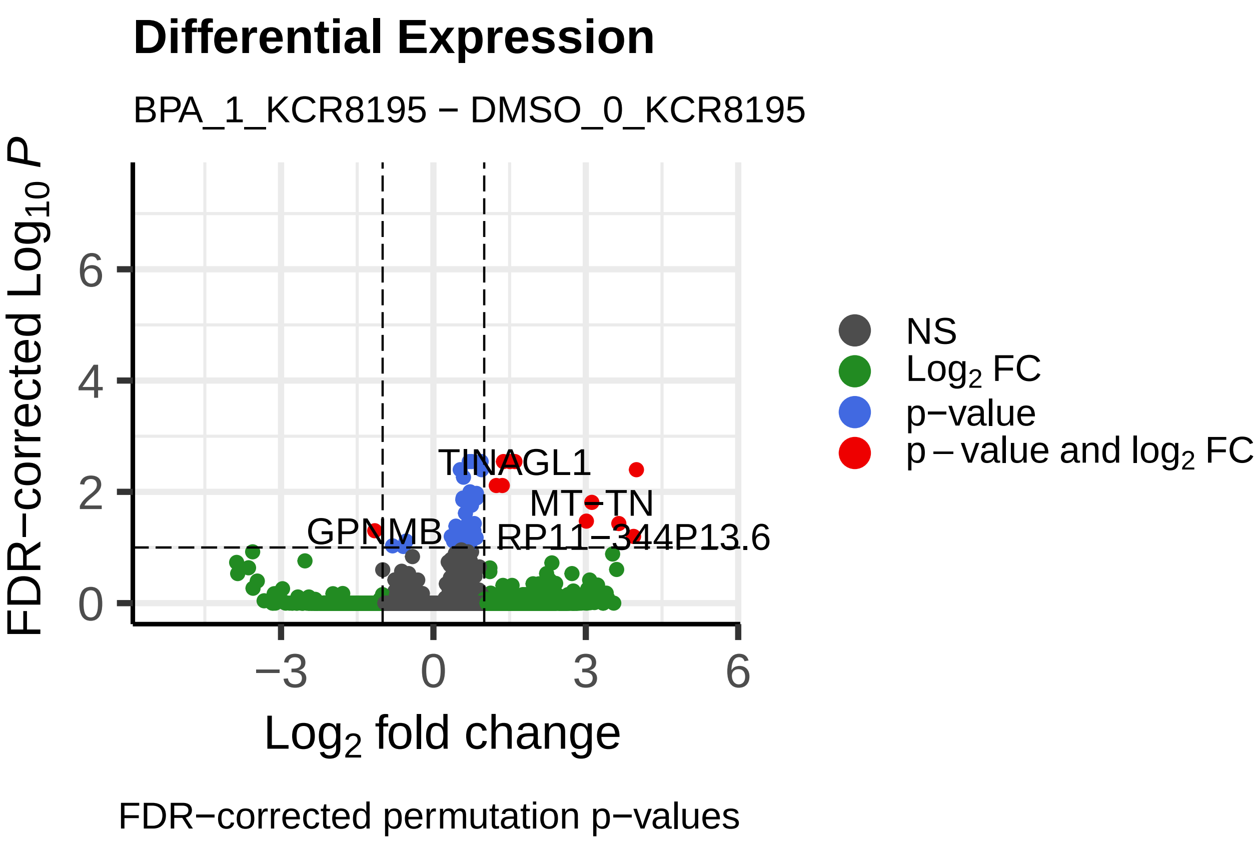
Differential Expression
BPA_1_KCR8195 − DMSO_0_KCR8195
FDR−corrected Log10 P
Log2 fold change
NS
Log2 FC
p−value
p − value and log2 FC
TINAGL1
MT−TN
GPNMB
RP11−344P13.6
FDR−corrected permutation p−values

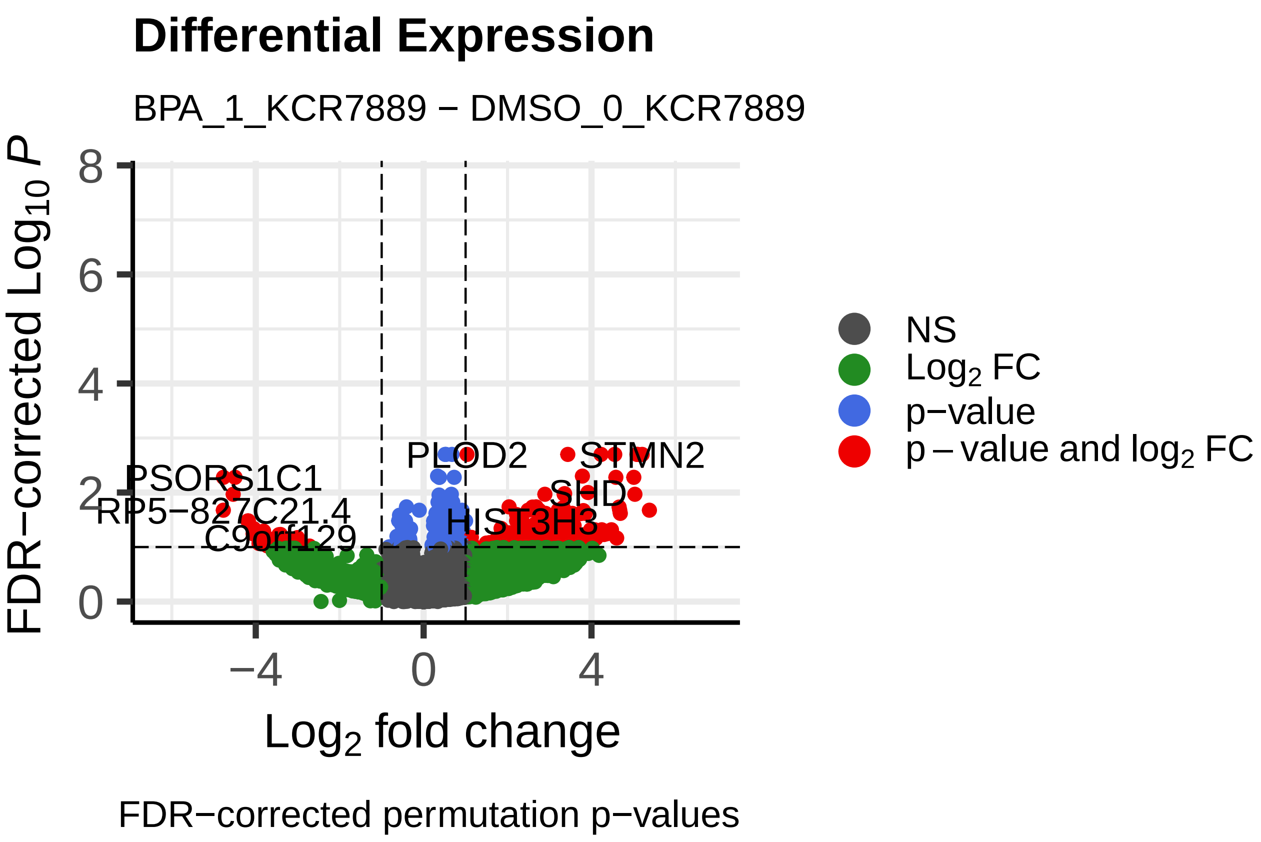
Differential Expression
BPA_1_KCR7889 − DMSO_0_KCR7889
FDR−corrected Log10 P
Log2 fold change
NS
Log2 FC
p−value
p − value and log2 FC
PLOD2
STMN2
PSORS1C1
SHD
RP5−827C21.4
HIST3H3
C9orf129
FDR−corrected permutation p−values

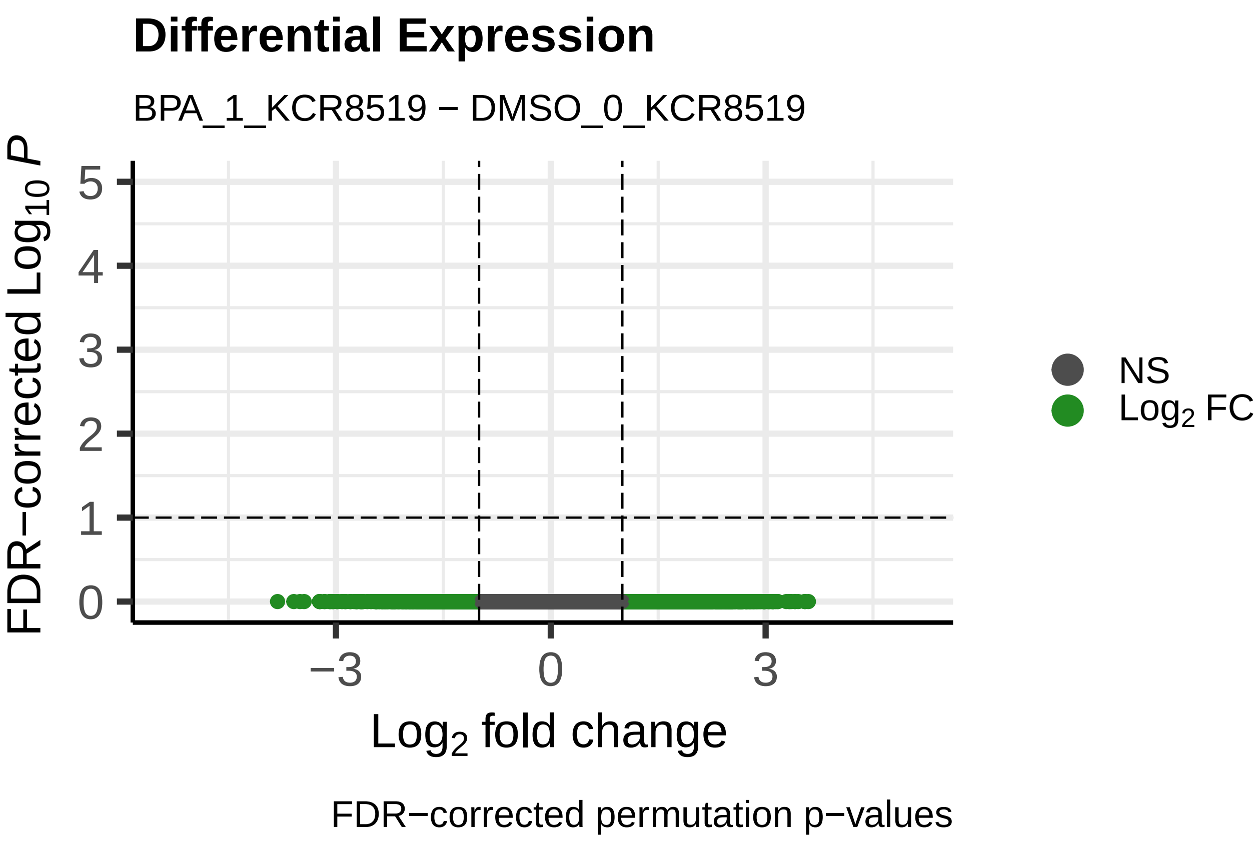
Differential Expression
BPA_1_KCR8519 − DMSO_0_KCR8519
FDR−corrected Log10 P
Log2 fold change
NS
Log2 FC
FDR−corrected permutation p−values

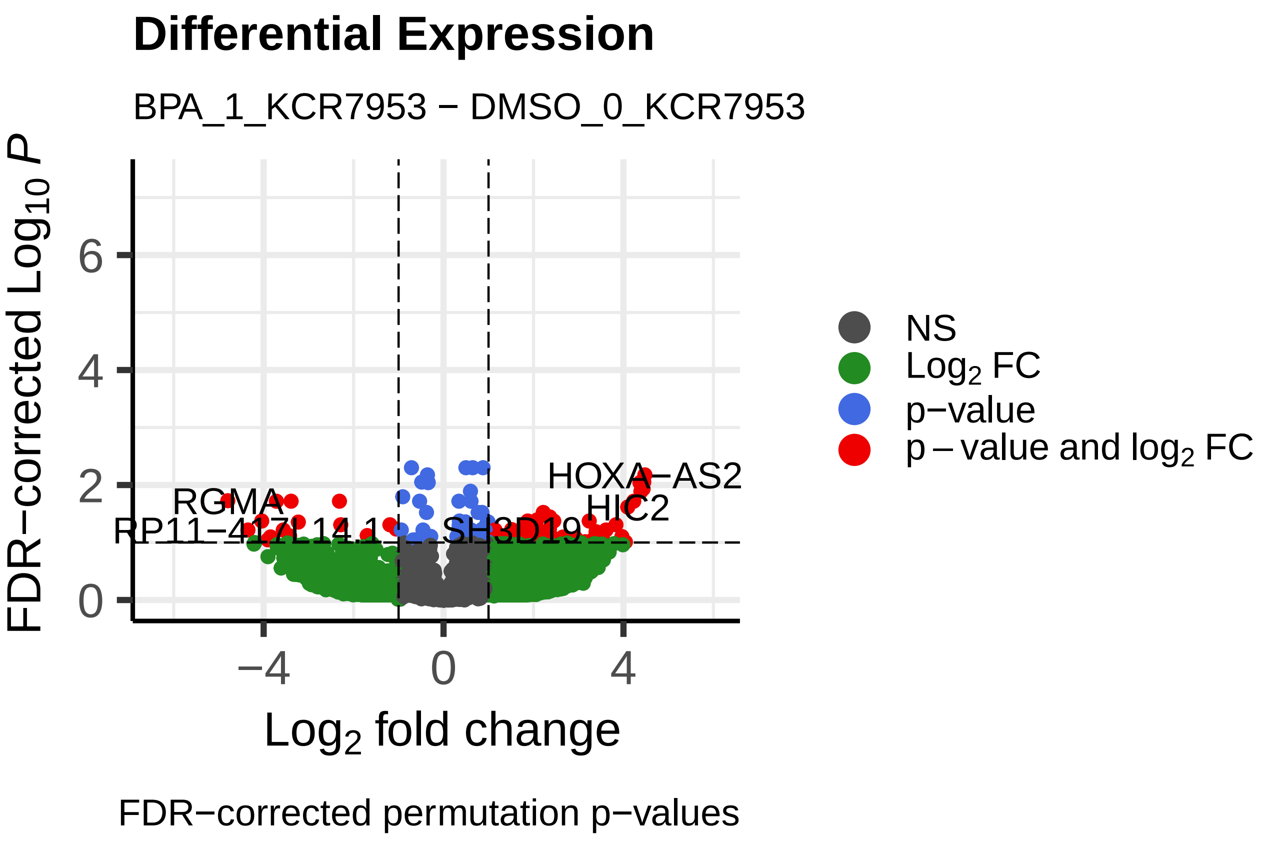
Differential Expression
BPA_1_KCR7953 − DMSO_0_KCR7953
FDR−corrected Log10 P
Log2 fold change
NS
Log2 FC
p−value
p − value and log2 FC
HOXA−AS2
RGMA
HIC2
SH3D19
FDR−corrected permutation p−values

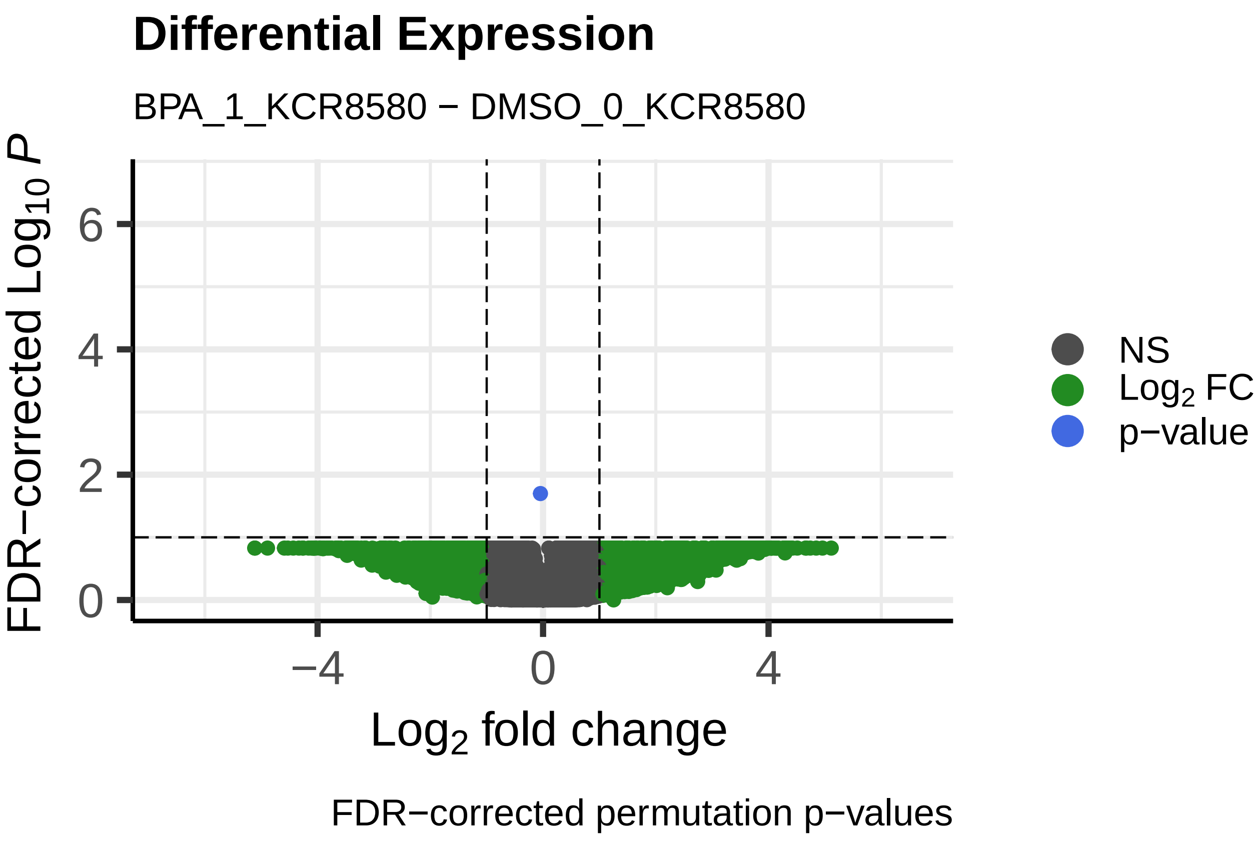
Differential Expression
BPA_1_KCR8580 − DMSO_0_KCR8580
FDR−corrected Log10 P
Log2 fold change
NS
Log2 FC
p−value
FDR−corrected permutation p−values

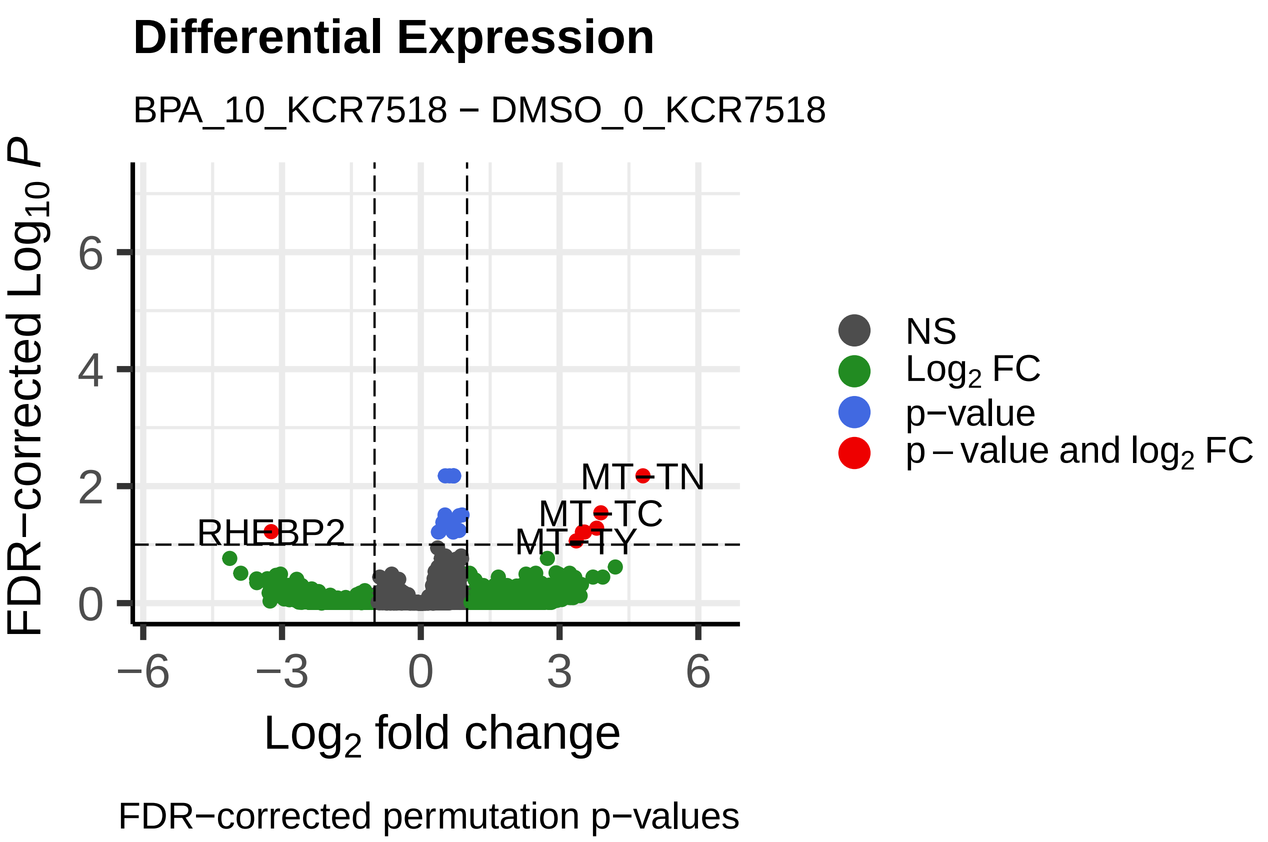
Differential Expression
BPA_10_KCR7518 − DMSO_0_KCR7518
FDR−corrected Log10 P
Log2 fold change
NS
Log2 FC
p−value
p − value and log2 FC
MT−TN
MT−TC
MT−TY
RHEBP2
FDR−corrected permutation p−values

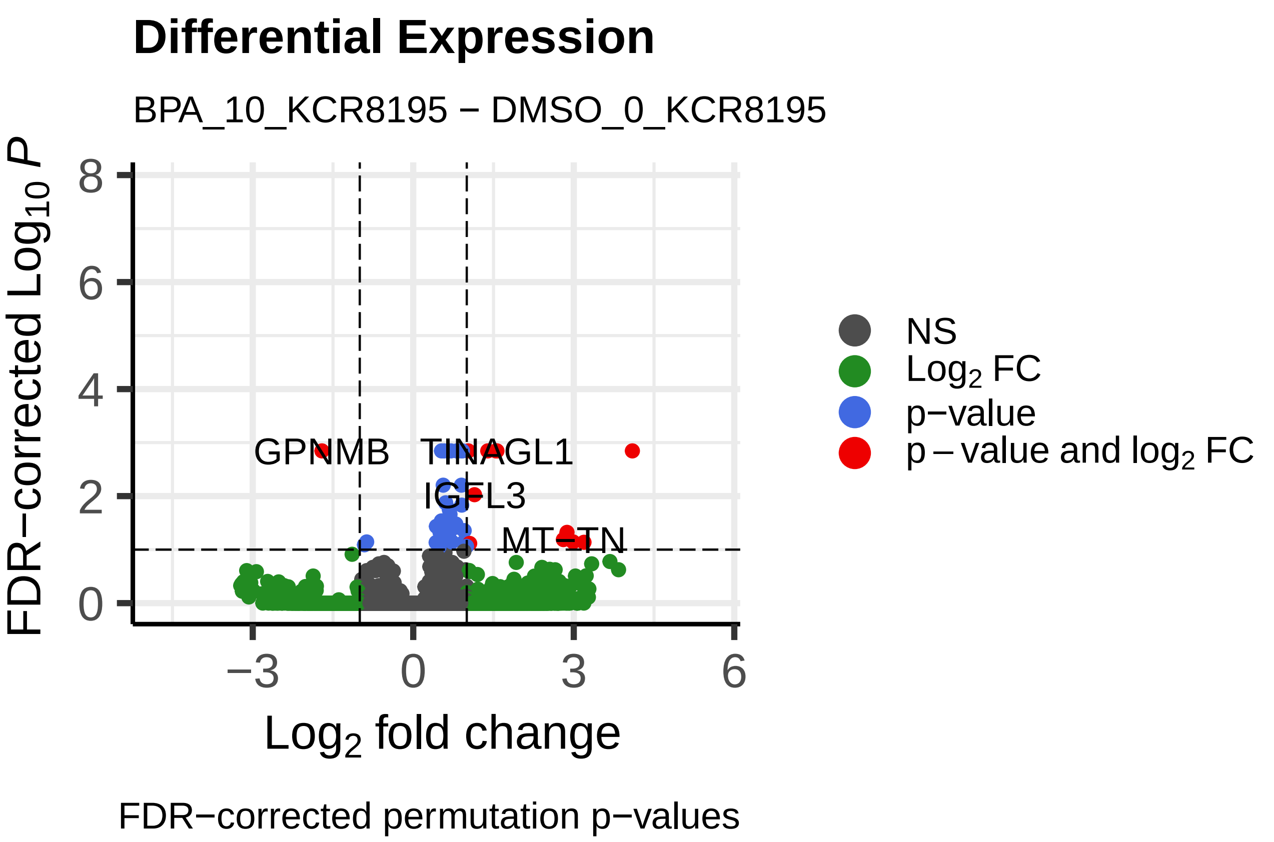
Differential Expression
BPA_10_KCR8195 − DMSO_0_KCR8195
FDR−corrected Log10 P
Log2 fold change
NS
Log2 FC
p−value
p − value and log2 FC
GPNMB
TINAGL1
IGFL3
MT−TN
FDR−corrected permutation p−values

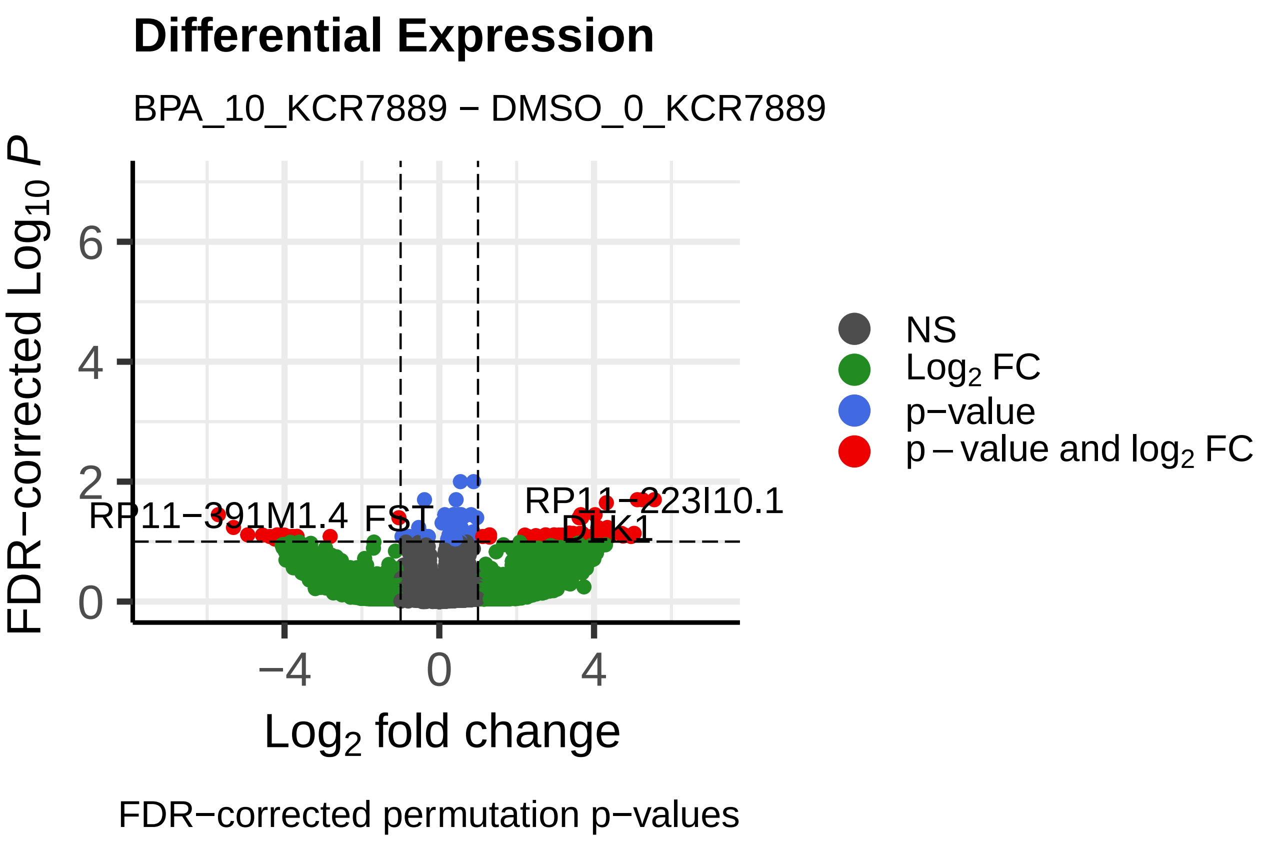
Differential Expression
BPA_10_KCR7889 − DMSO_0_KCR7889
FDR−corrected Log10 P
Log2 fold change
NS
Log2 FC
p−value
p − value and log2 FC
RP11−391M1.4
FST
RP11−223I10.1
DLK1
FDR−corrected permutation p−values

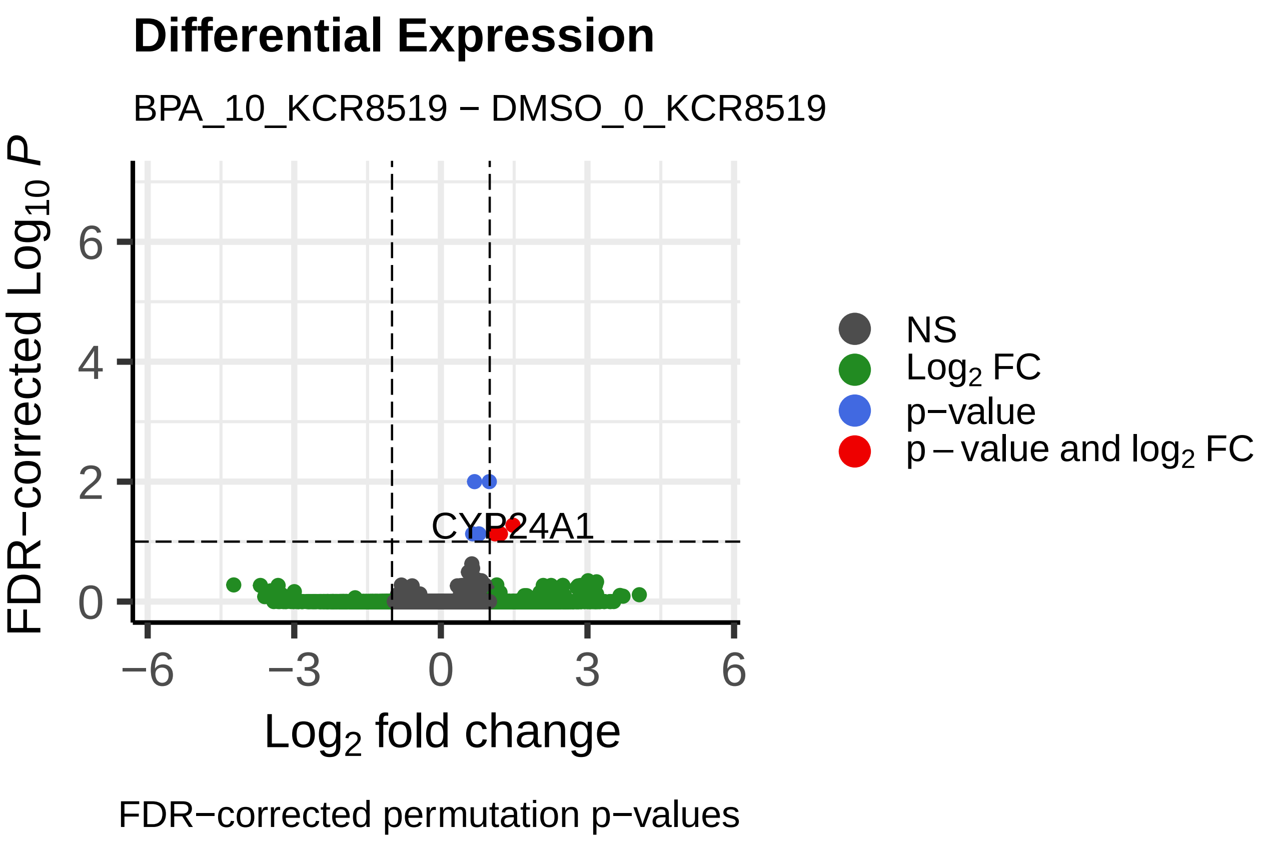
Differential Expression
BPA_10_KCR8519 − DMSO_0_KCR8519
FDR−corrected Log10 P
Log2 fold change
NS
Log2 FC
p−value
p − value and log2 FC
CYP24A1
FDR−corrected permutation p−values

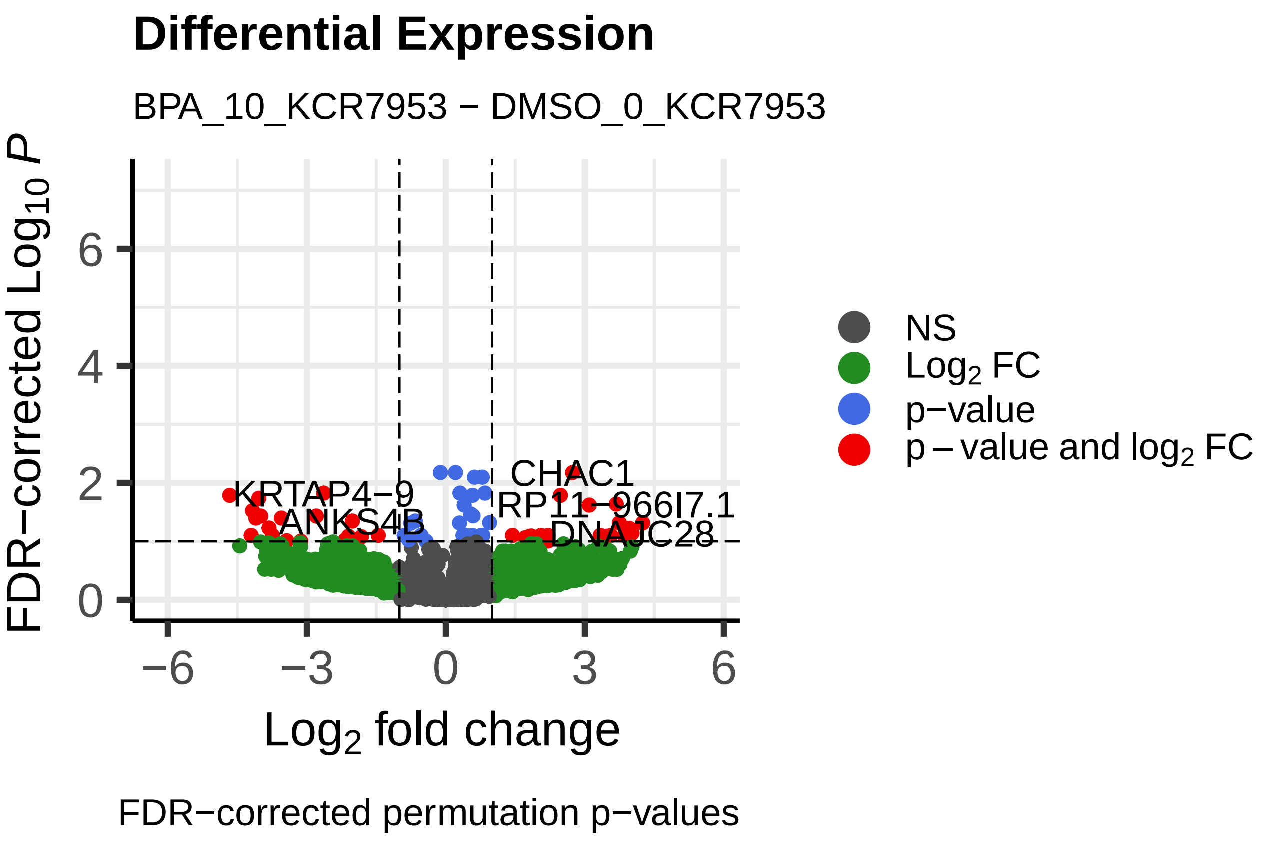
Differential Expression
BPA_10_KCR7953 − DMSO_0_KCR7953
FDR−corrected Log10 P
Log2 fold change
NS
Log2 FC
p−value
p − value and log2 FC
CHAC1
KRTAP4−9
RP11−966I7.1
ANKS4B
DNAJC28
FDR−corrected permutation p−values

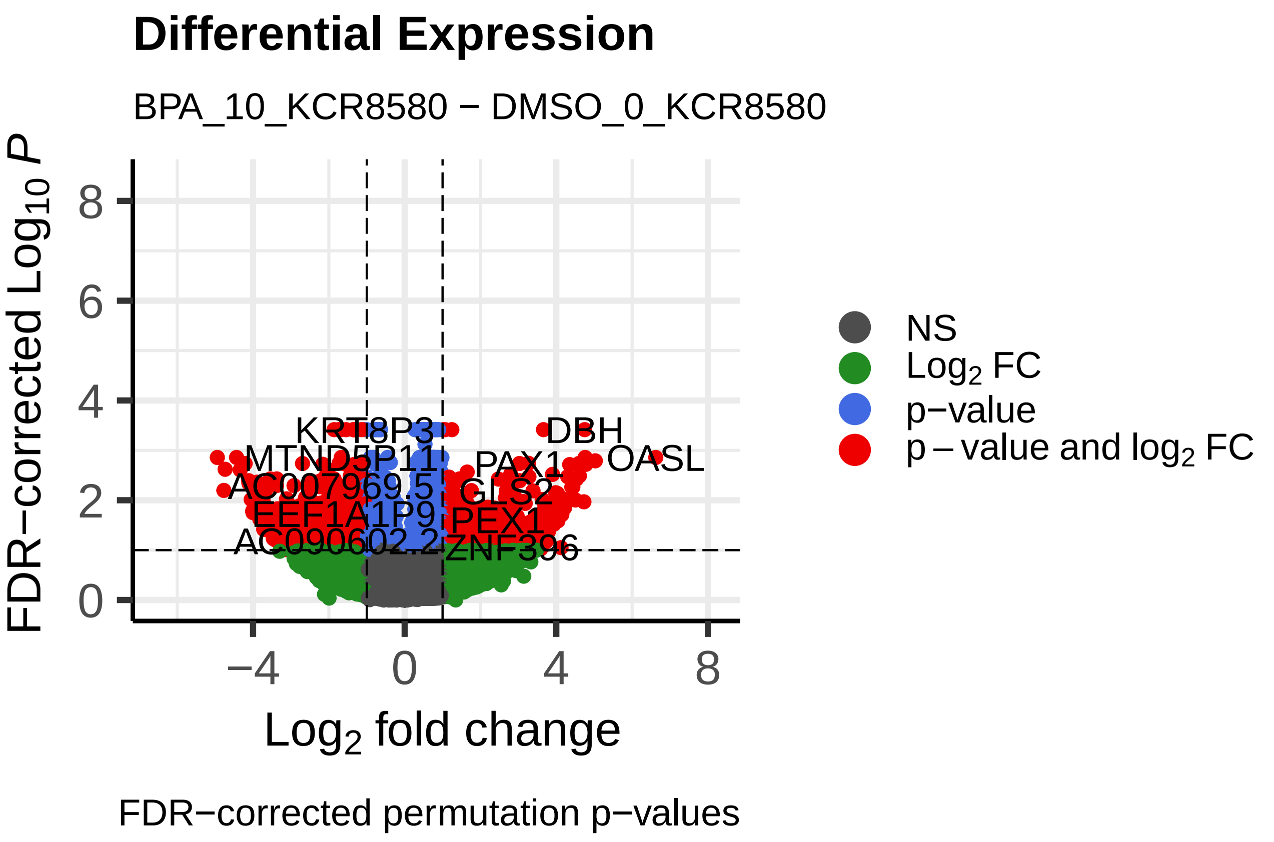
Differential Expression
BPA_10_KCR8580 − DMSO_0_KCR8580
FDR−corrected Log10 P
Log2 fold change
NS
Log2 FC
p−value
p − value and log2 FC
KRT8P3
DBH
PAX1
OASL
CTS2
PEX1
ZNF396
FDR−corrected permutation p−values

### Differential Expression

BPS_0.1_KCR7518 − DMSO_0_KCR7518

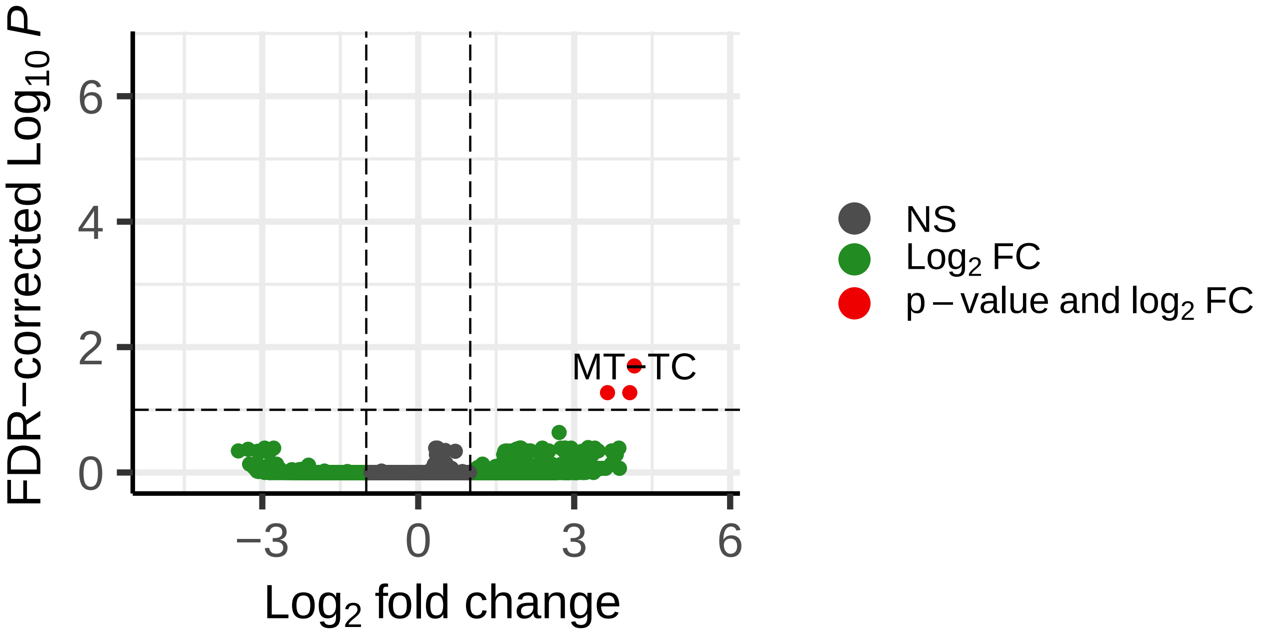


FDR−corrected permutation p−values

### Differential Expression

BPS_0.1_KCR8195 − DMSO_0_KCR8195

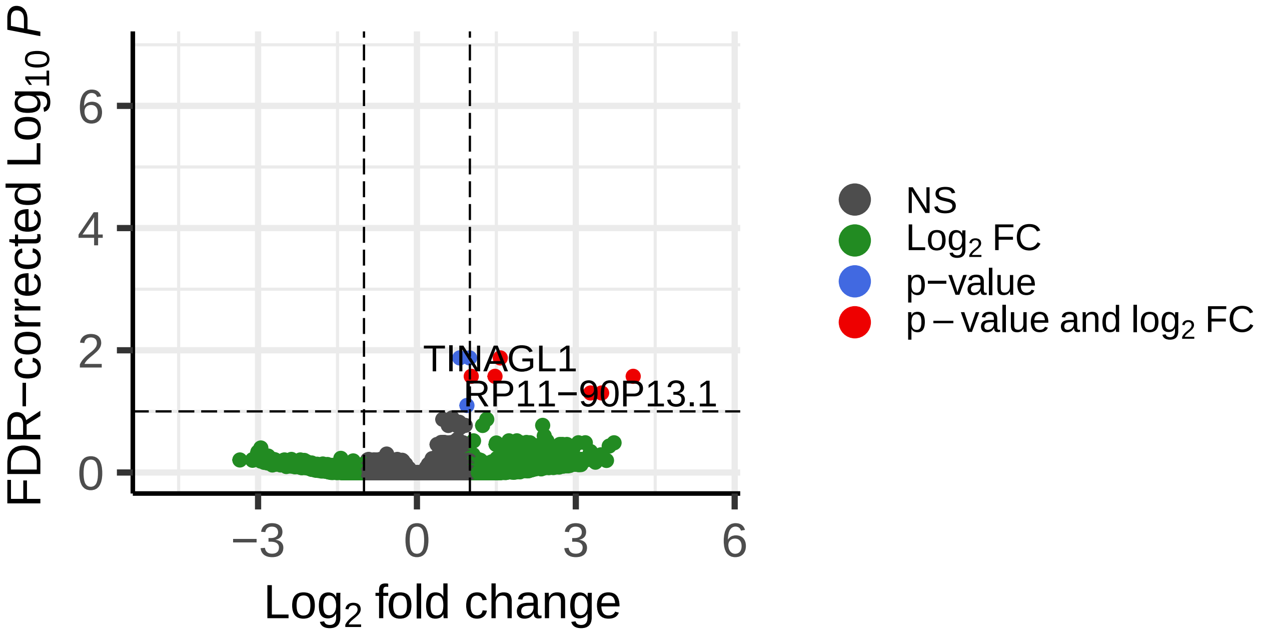


FDR−corrected permutation p−values

### Differential Expression

BPS_0.1_KCR7889 − DMSO_0_KCR7889

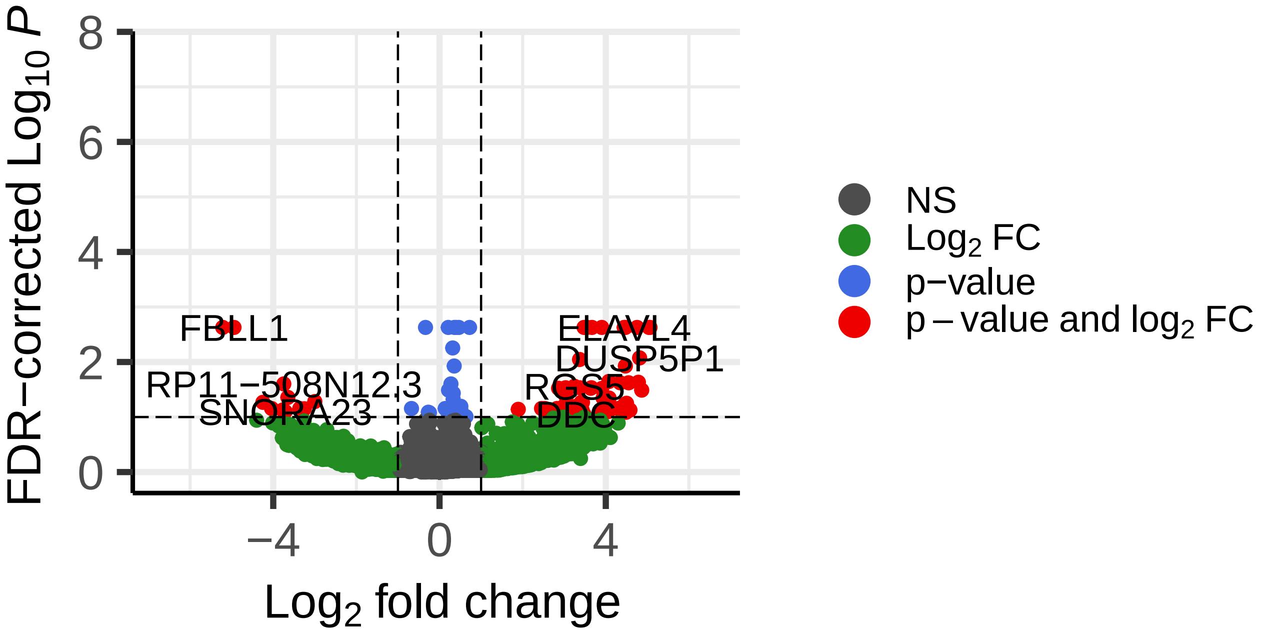


FDR−corrected permutation p−values

### Differential Expression

BPS_0.1_KCR8519 − DMSO_0_KCR8519

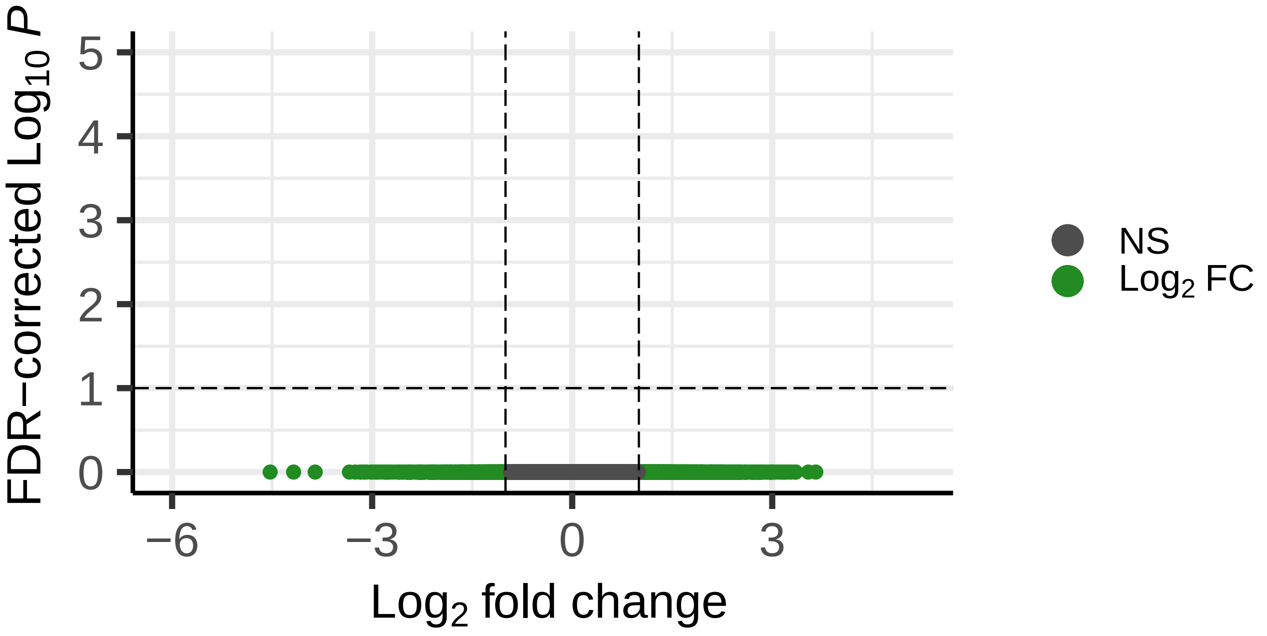


FDR−corrected permutation p−values

### Differential Expression

BPS_0.1_KCR7953 − DMSO_0_KCR7953

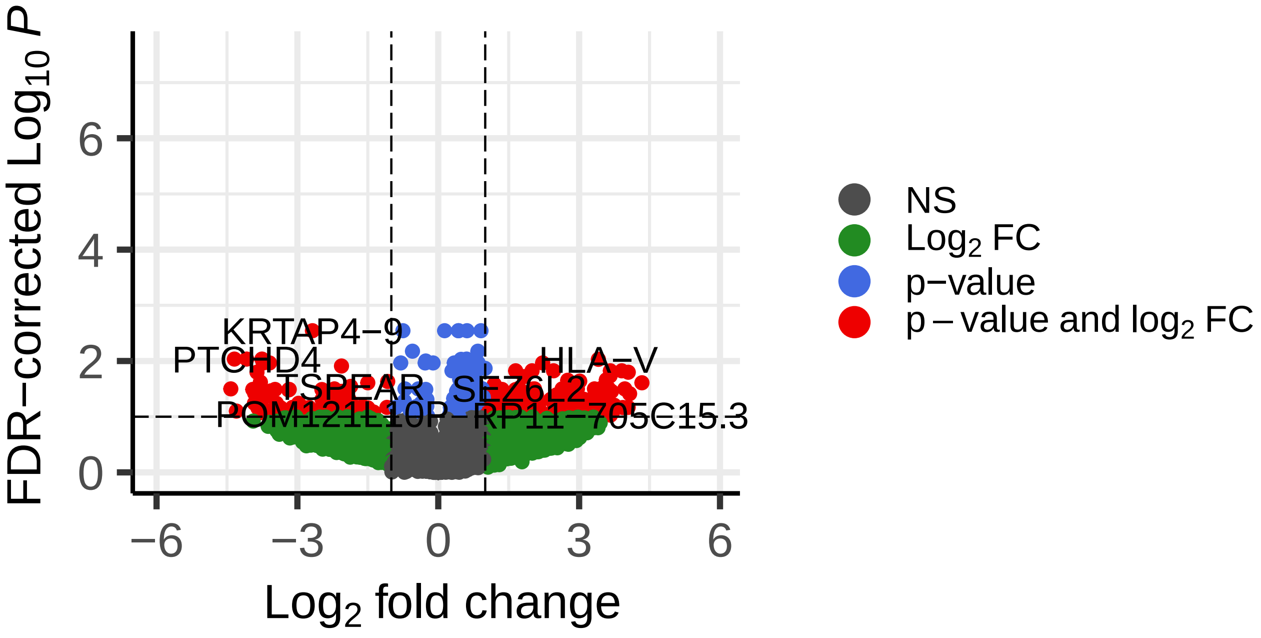


FDR−corrected permutation p−values

### Differential Expression

BPS_0.1_KCR8580 − DMSO_0_KCR8580

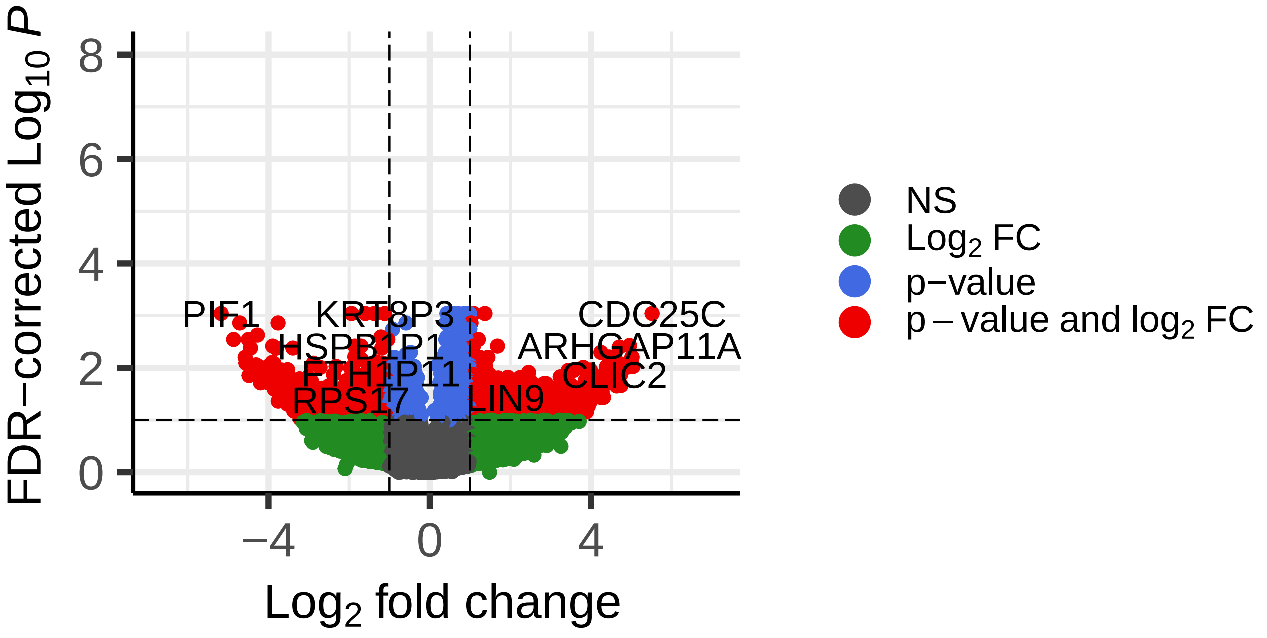


FDR−corrected permutation p−values

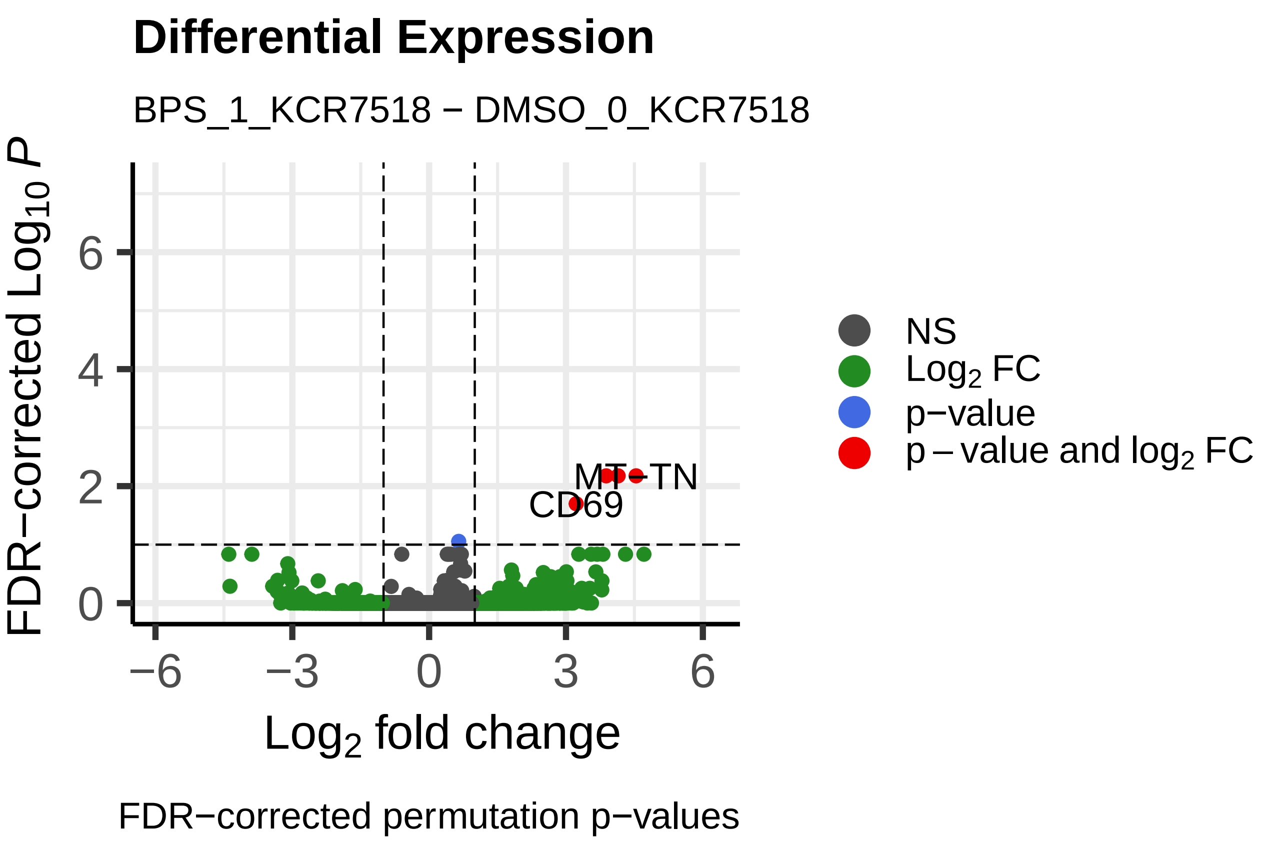
Differential Expression
BPS_1_KCR7518 − DMSO_0_KCR7518
FDR−corrected $Log_{10}$ $P$
Log₂ fold change
MT−TN
CD69
NS
$Log_2$ FC
p−value
p − value and $log_2$ FC
FDR−corrected permutation p−values

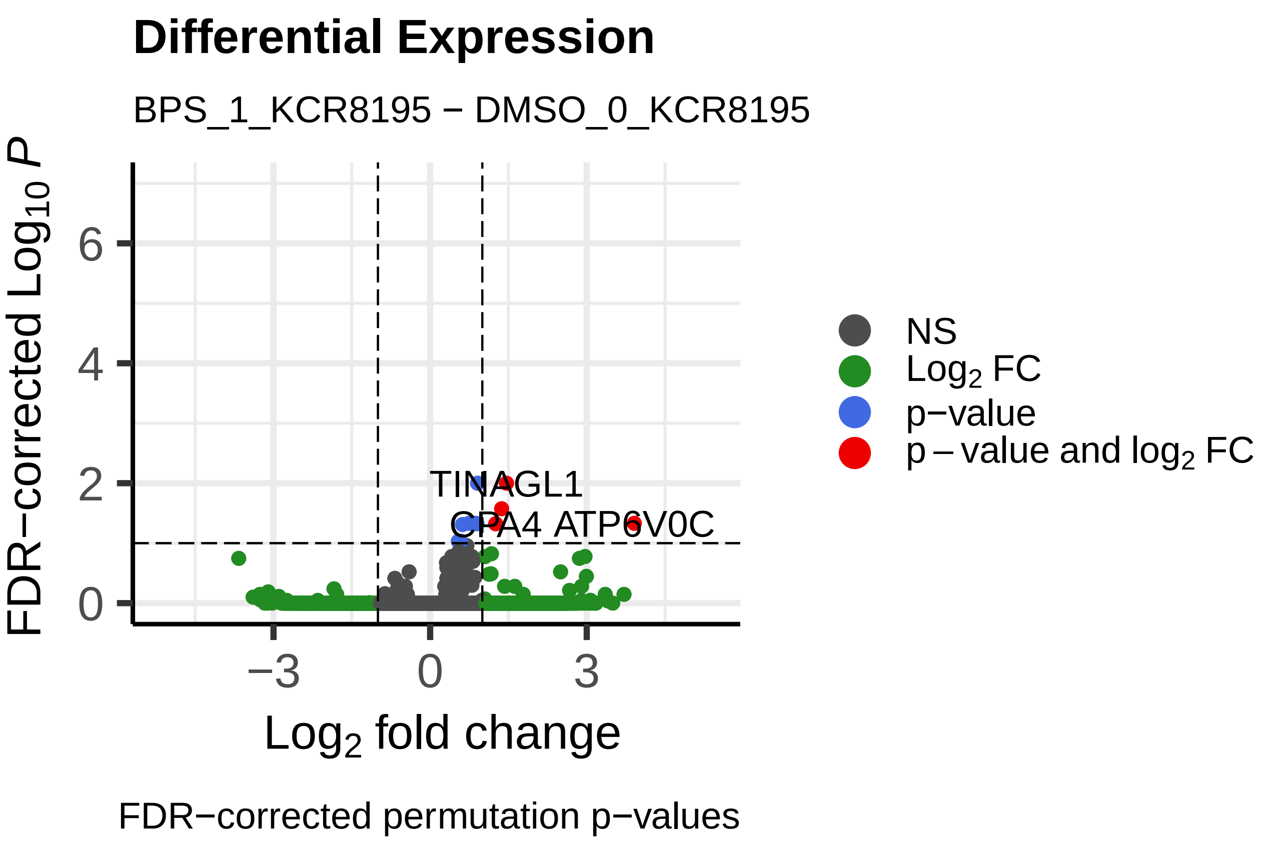
Differential Expression
BPS_1_KCR8195 − DMSO_0_KCR8195
FDR−corrected $Log_{10}$ $P$
$Log_2$ fold change
TINAGL1
CPA4
ATP6V0C
NS
$Log_2$ FC
p−value
p − value and $log_2$ FC
FDR−corrected permutation p−values

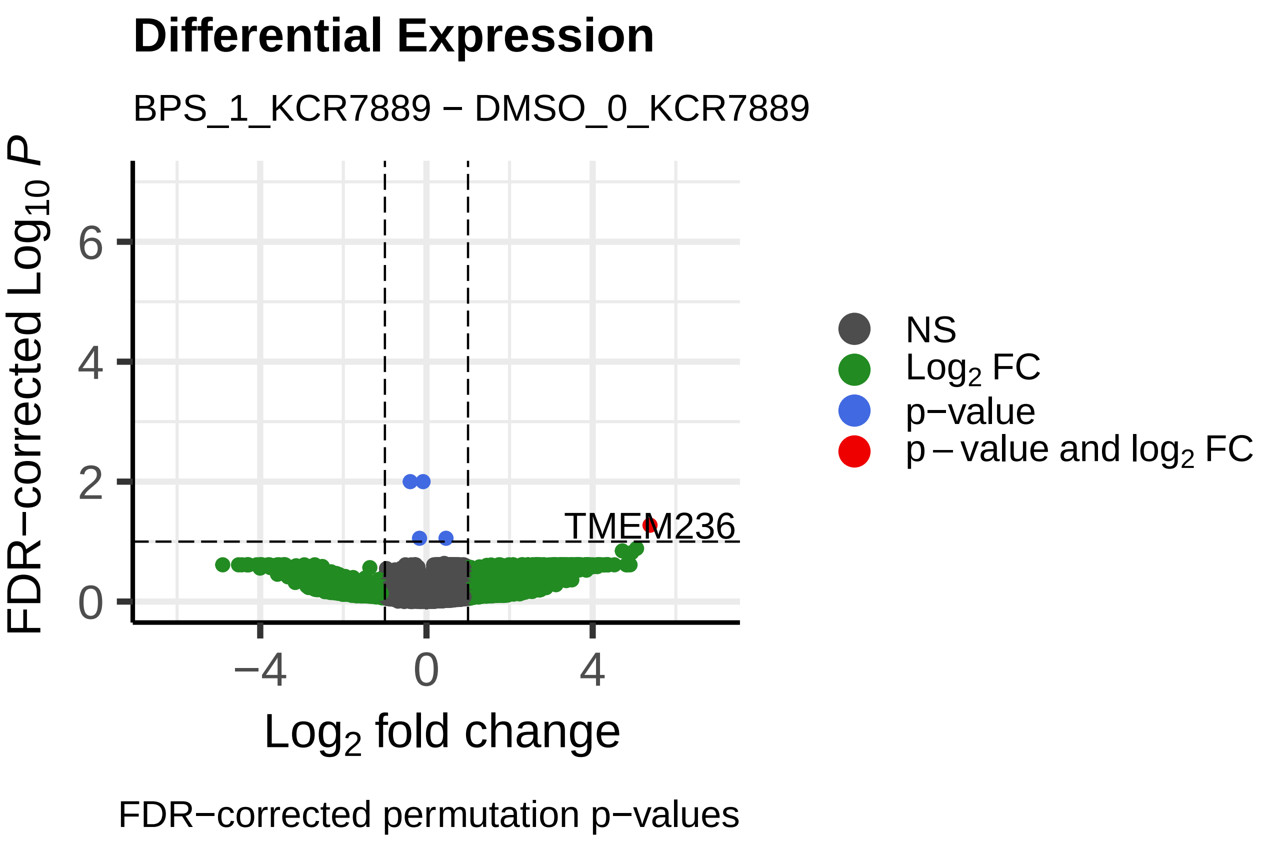
Differential Expression
BPS_1_KCR7889 − DMSO_0_KCR7889
FDR−corrected $Log_{10}$ $P$
$Log_2$ fold change
TMEM236
NS
$Log_2$ FC
p−value
p − value and $log_2$ FC
FDR−corrected permutation p−values

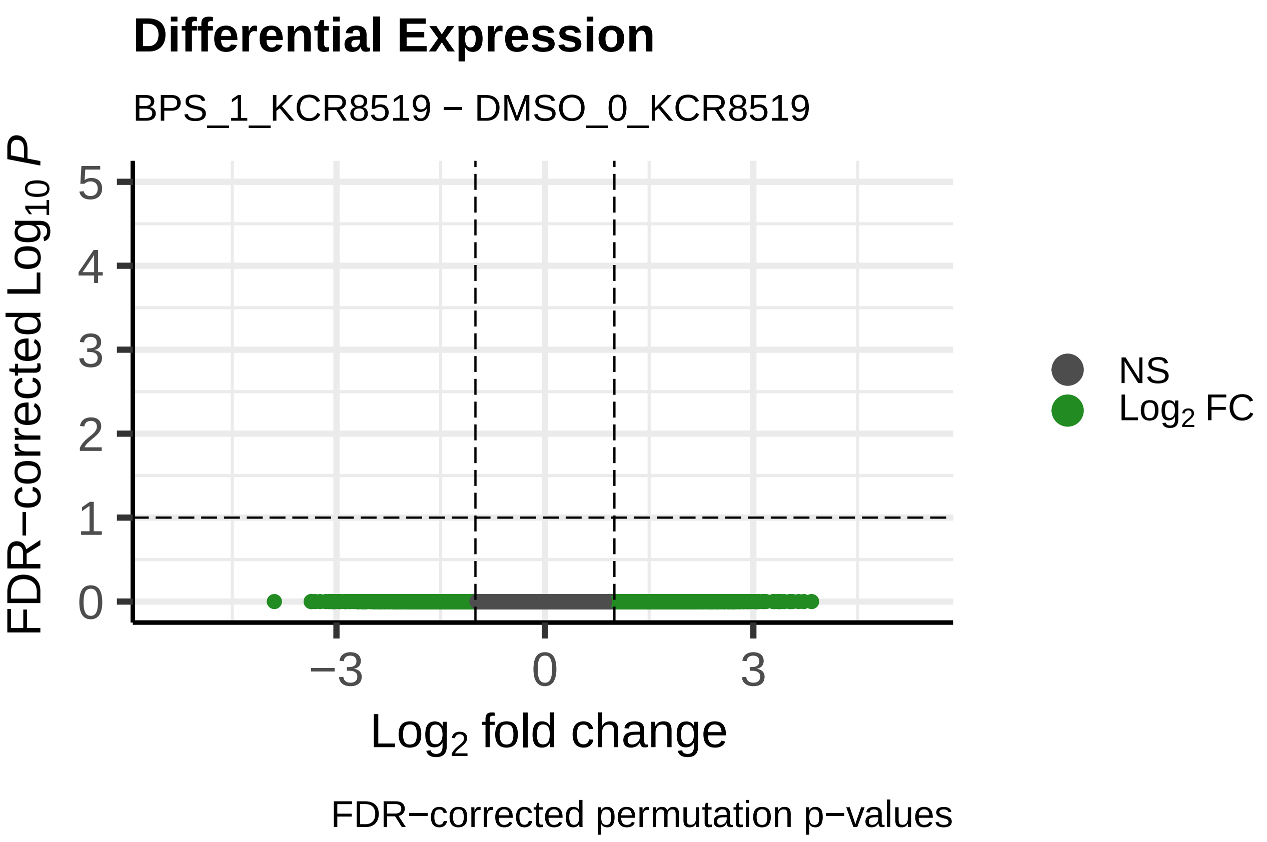
Differential Expression
BPS_1_KCR8519 − DMSO_0_KCR8519
FDR−corrected $Log_{10}$ $P$
$Log_2$ fold change
NS
$Log_2$ FC
FDR−corrected permutation p−values

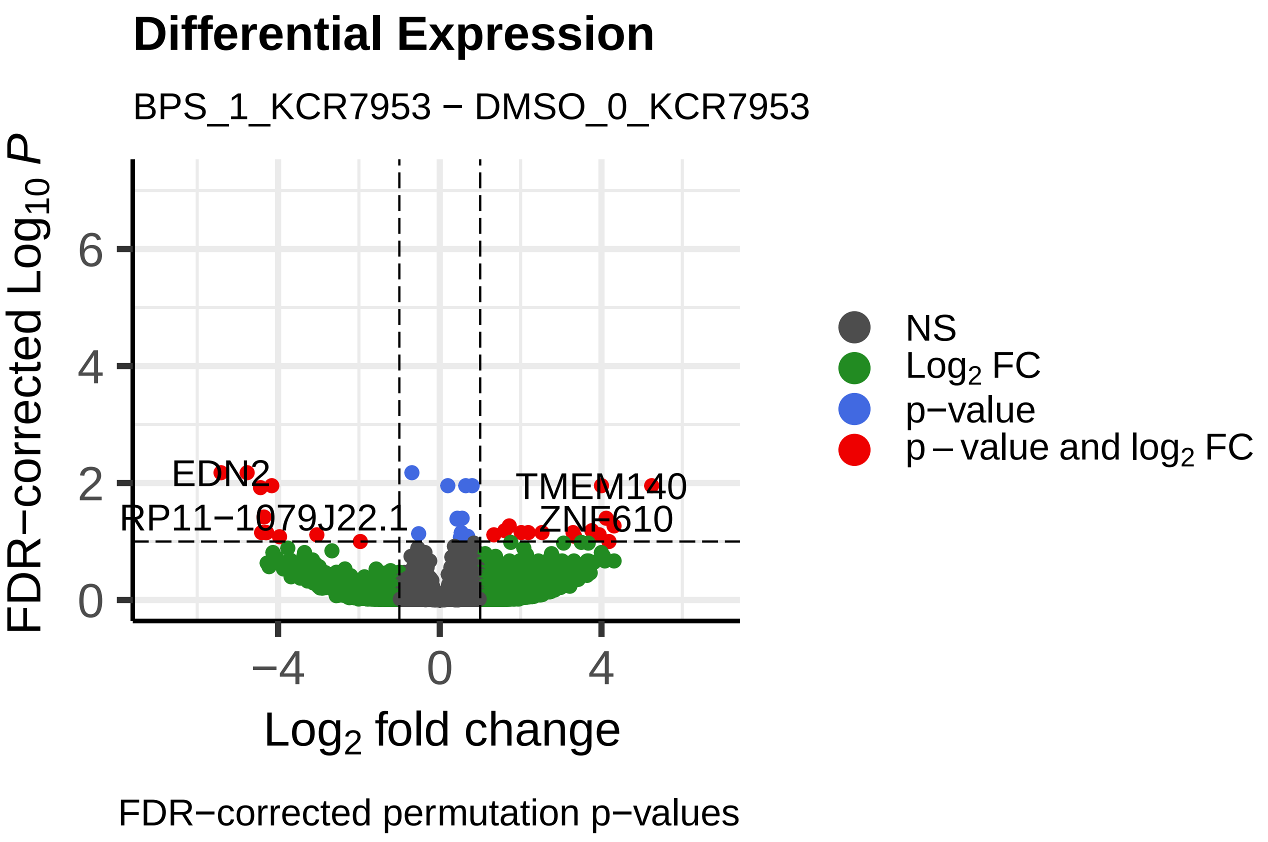
Differential Expression
BPS_1_KCR7953 − DMSO_0_KCR7953
FDR−corrected $Log_{10}$ $P$
$Log_2$ fold change
EDN2
RP11−1079J22.1
TMEM140
ZNF610
NS
$Log_2$ FC
p−value
p − value and $log_2$ FC
FDR−corrected permutation p−values

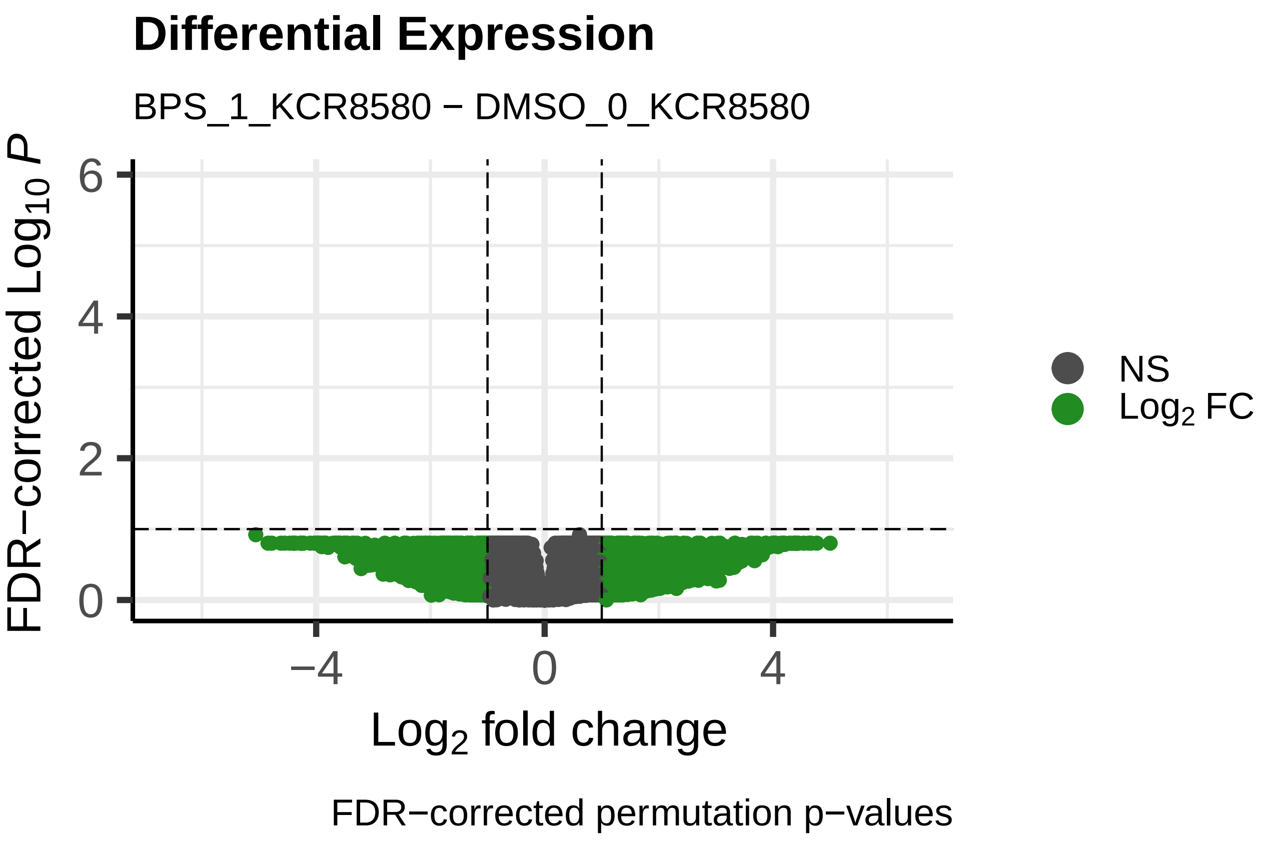
Differential Expression
BPS_1_KCR8580 − DMSO_0_KCR8580
FDR−corrected $Log_{10}$ $P$
$Log_2$ fold change
NS
$Log_2$ FC
FDR−corrected permutation p−values

### Differential Expression

BPS_10_KCR7518 − DMSO_0_KCR7518

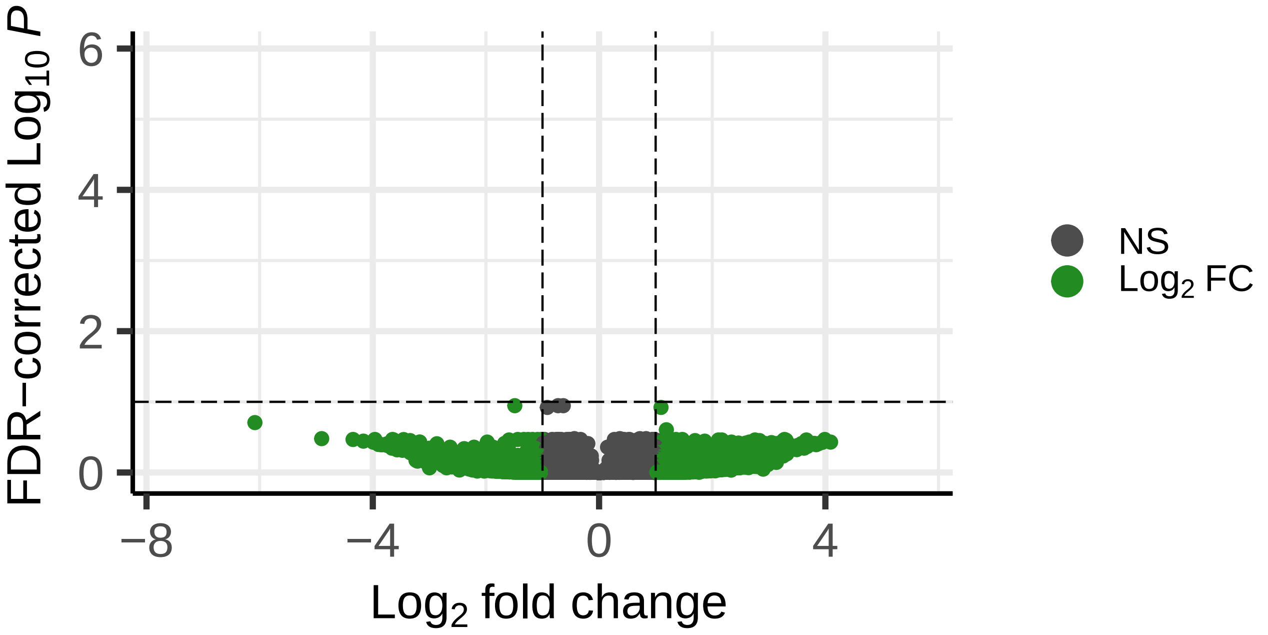


FDR−corrected permutation p−values

### Differential Expression

BPS_10_KCR8195 − DMSO_0_KCR8195

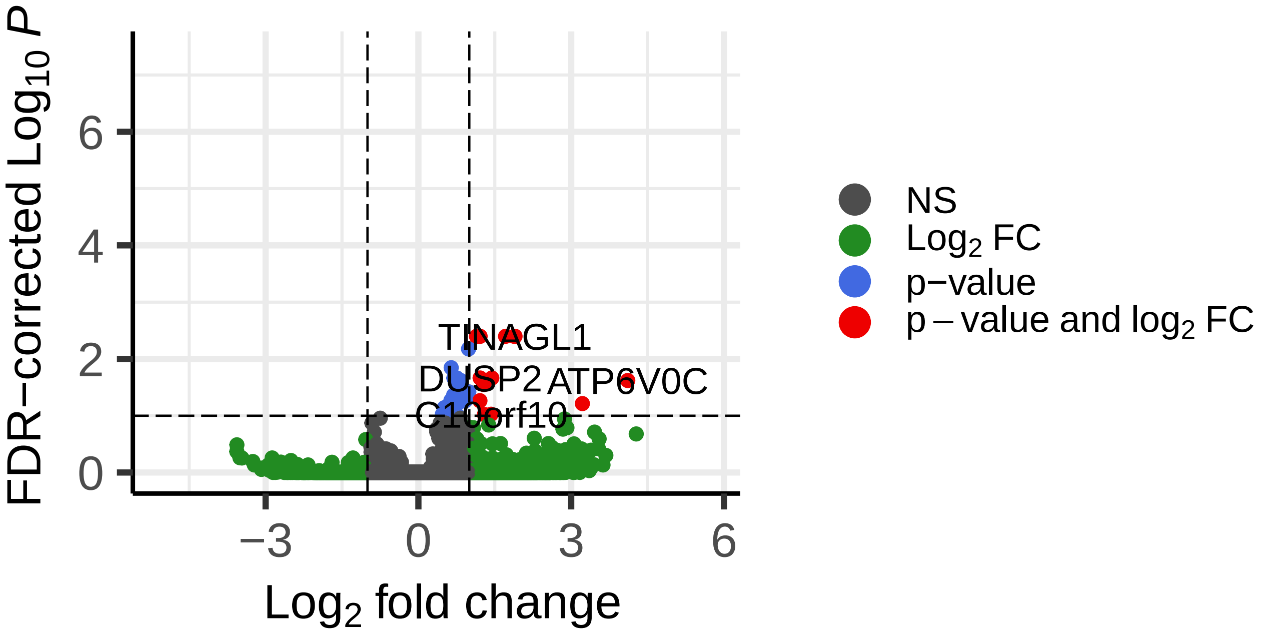


FDR−corrected permutation p−values

### Differential Expression

BPS_10_KCR7889 − DMSO_0_KCR7889

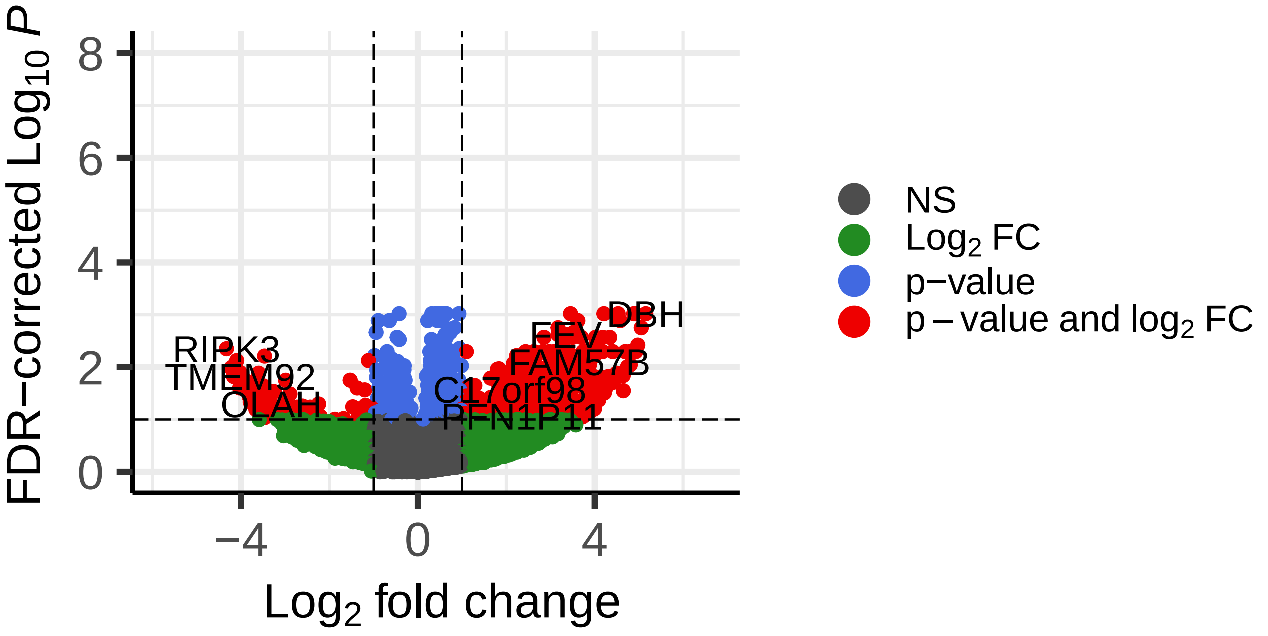


FDR−corrected permutation p−values

### Differential Expression

BPS_10_KCR8519 − DMSO_0_KCR8519

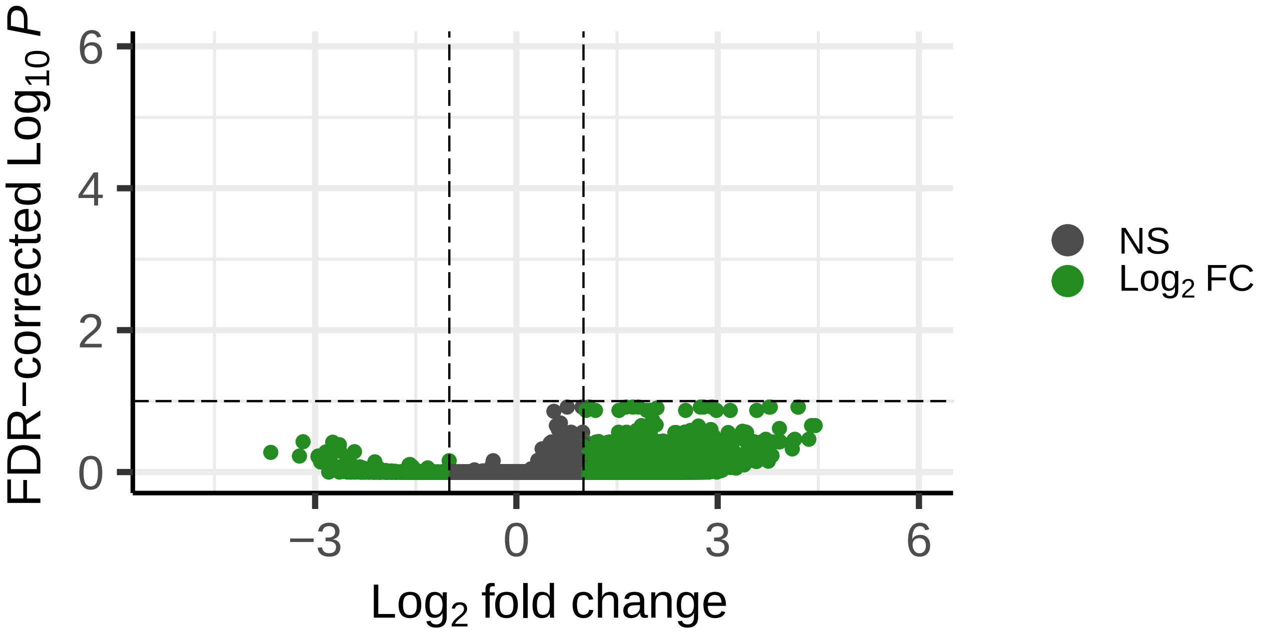


FDR−corrected permutation p−values

### Differential Expression

BPS_10_KCR7953 − DMSO_0_KCR7953

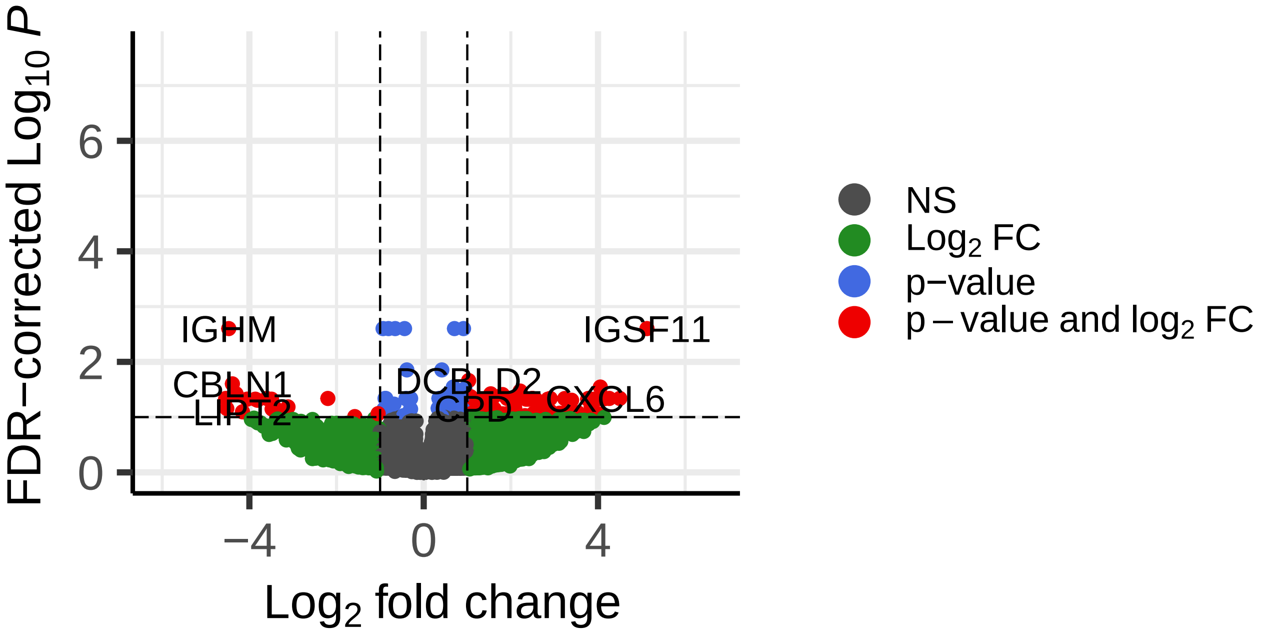


FDR−corrected permutation p−values

### Differential Expression

BPS_10_KCR8580 − DMSO_0_KCR8580

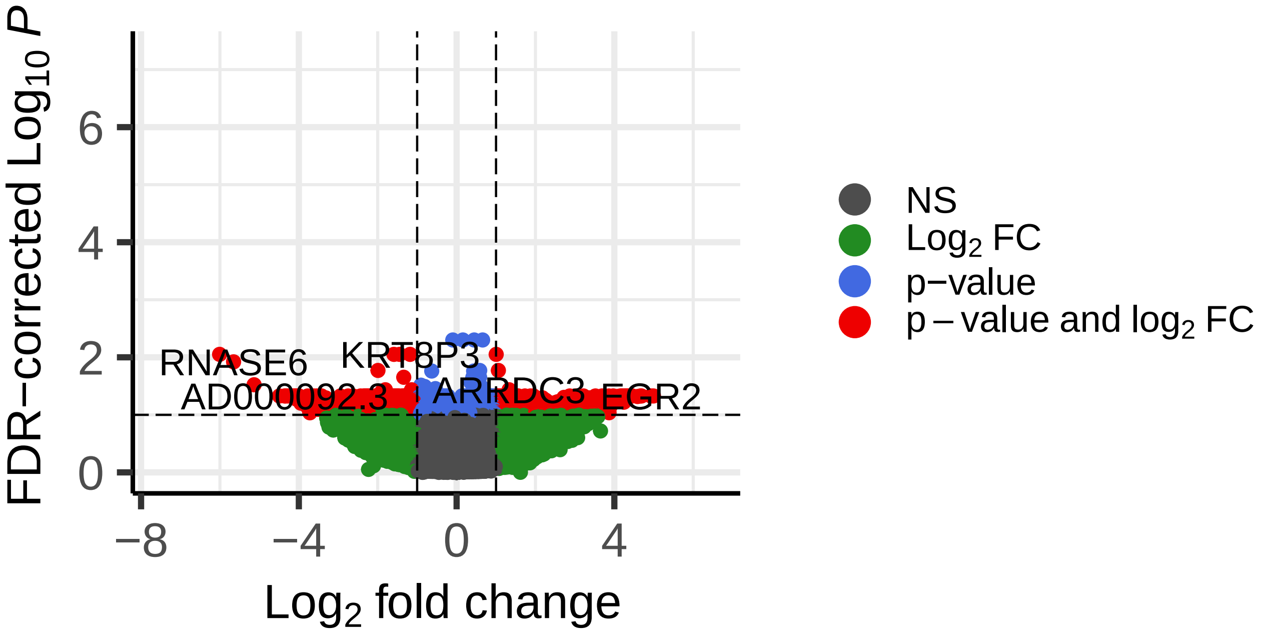


FDR−corrected permutation p−values

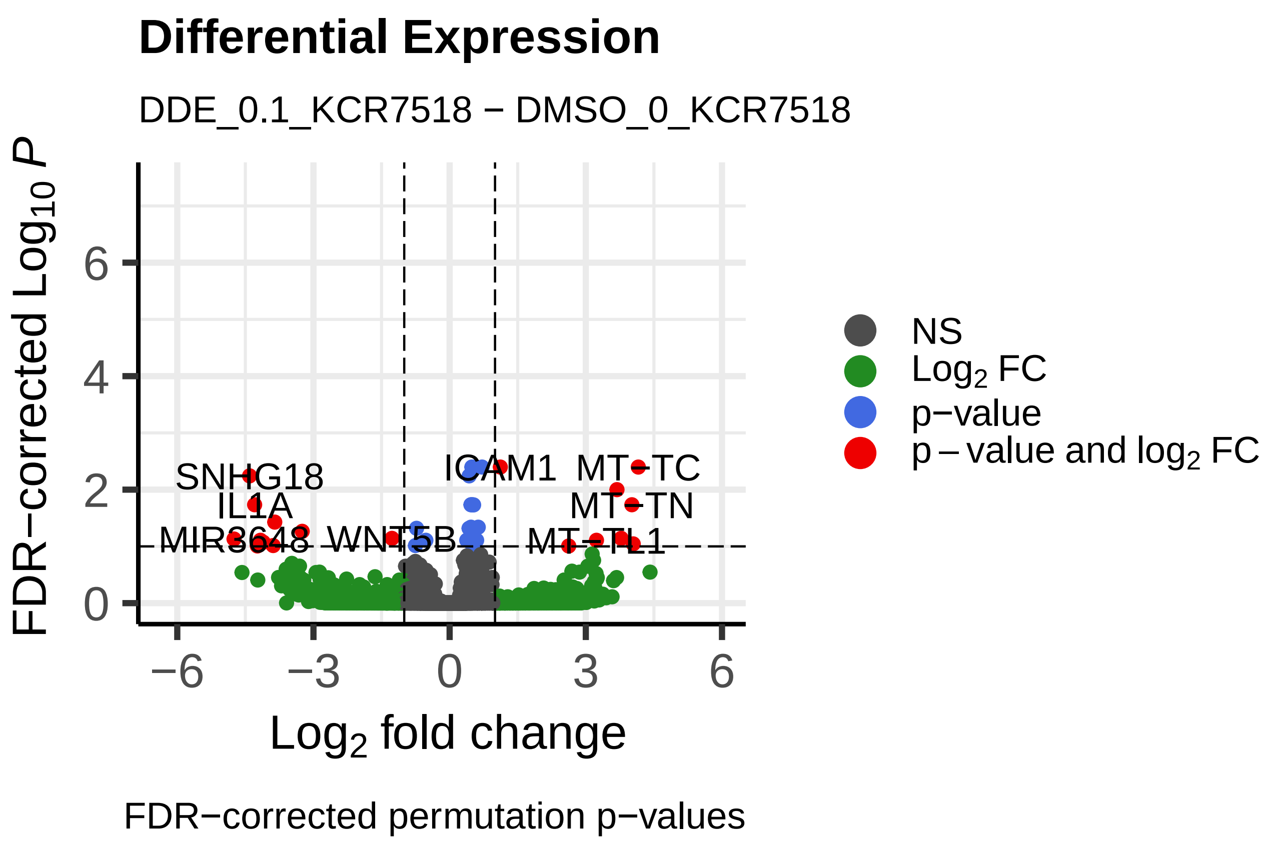
Differential Expression
DDE_0.1_KCR7518 − DMSO_0_KCR7518
FDR−corrected $Log_{10}$ P
Log2 fold change
NS
Log2 FC
p−value
p − value and log2 FC
SNHG18
IL1A
MIR3648
WNT5B
ICAM1
MT−TC
MT−TN
MT−TL1
FDR−corrected permutation p−values

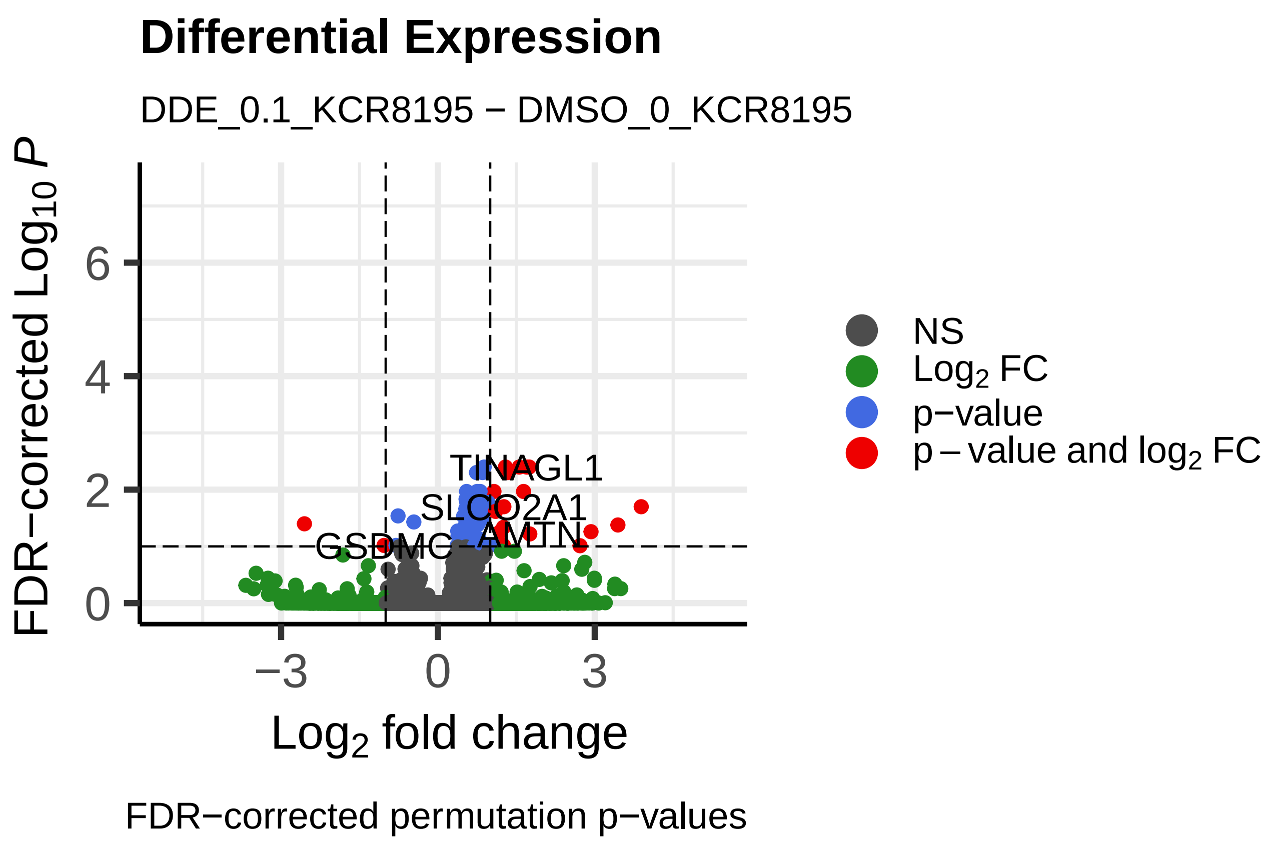
Differential Expression
DDE_0.1_KCR8195 − DMSO_0_KCR8195
FDR−corrected $Log_{10}$ P
Log2 fold change
NS
Log2 FC
p−value
p − value and log2 FC
TINAGL1
SLCO2A1
GSDMC
AMTN
FDR−corrected permutation p−values

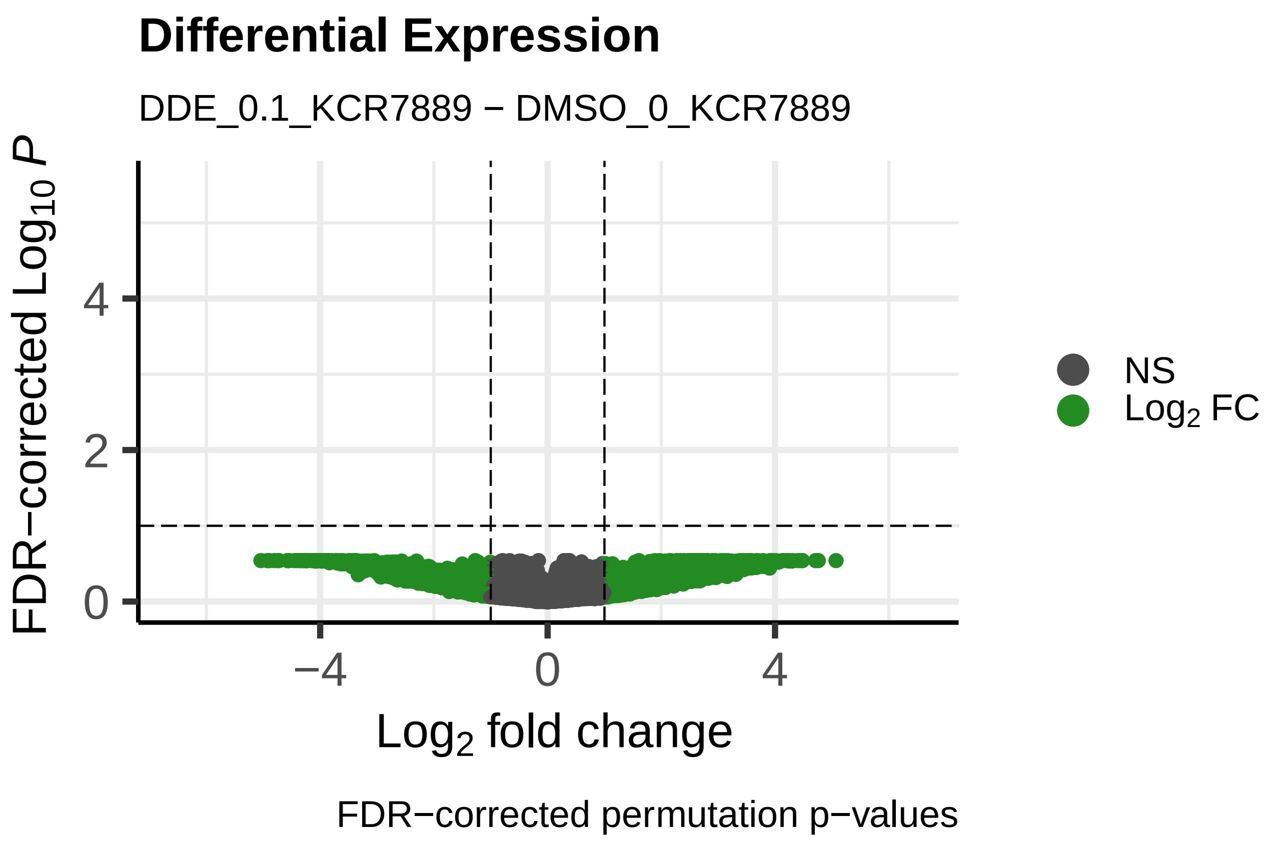
Differential Expression
DDE_0.1_KCR7889 − DMSO_0_KCR7889
FDR−corrected $Log_{10}$ P
Log2 fold change
NS
Log2 FC
FDR−corrected permutation p−values

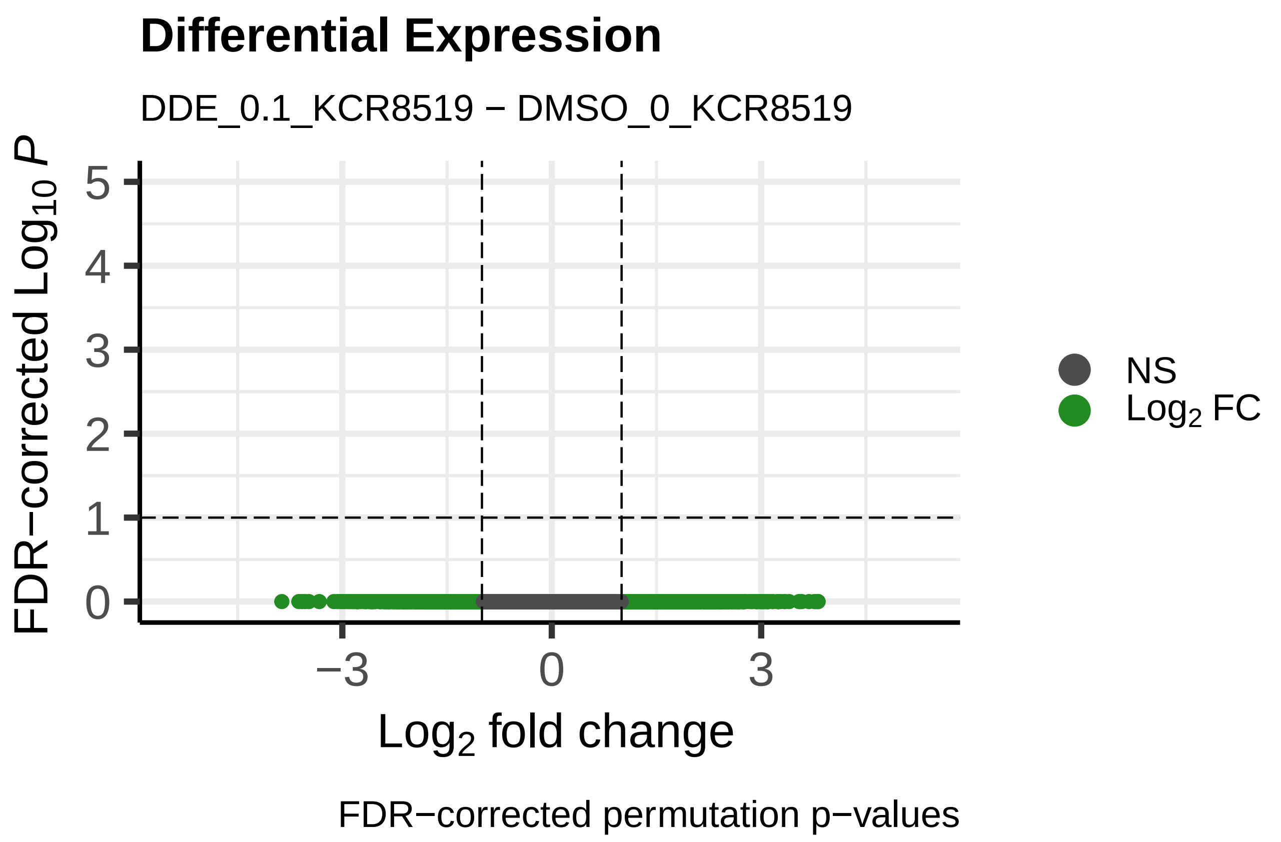
Differential Expression
DDE_0.1_KCR8519 − DMSO_0_KCR8519
FDR−corrected $Log_{10}$ P
Log2 fold change
NS
Log2 FC
FDR−corrected permutation p−values

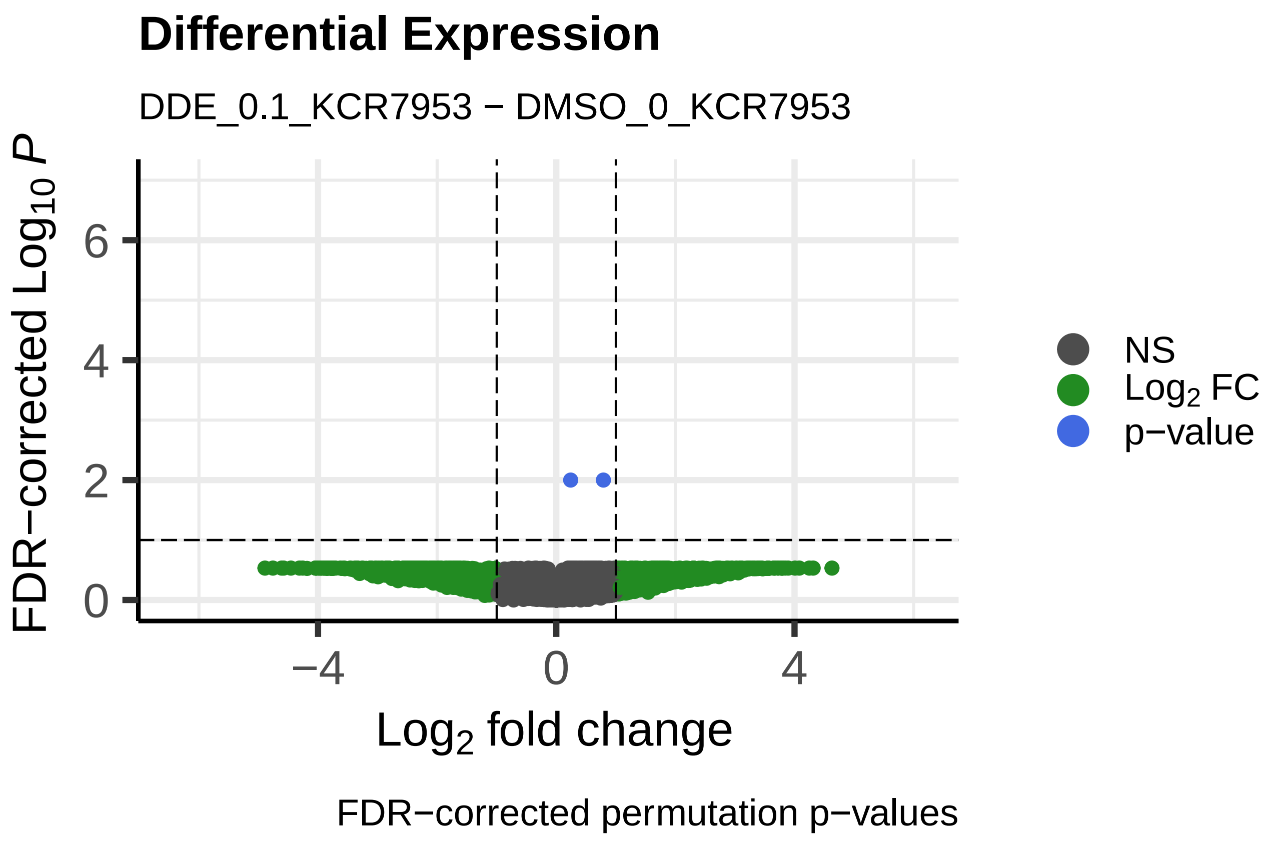
Differential Expression
DDE_0.1_KCR7953 − DMSO_0_KCR7953
FDR−corrected $Log_{10}$ P
Log2 fold change
NS
Log2 FC
p−value
FDR−corrected permutation p−values

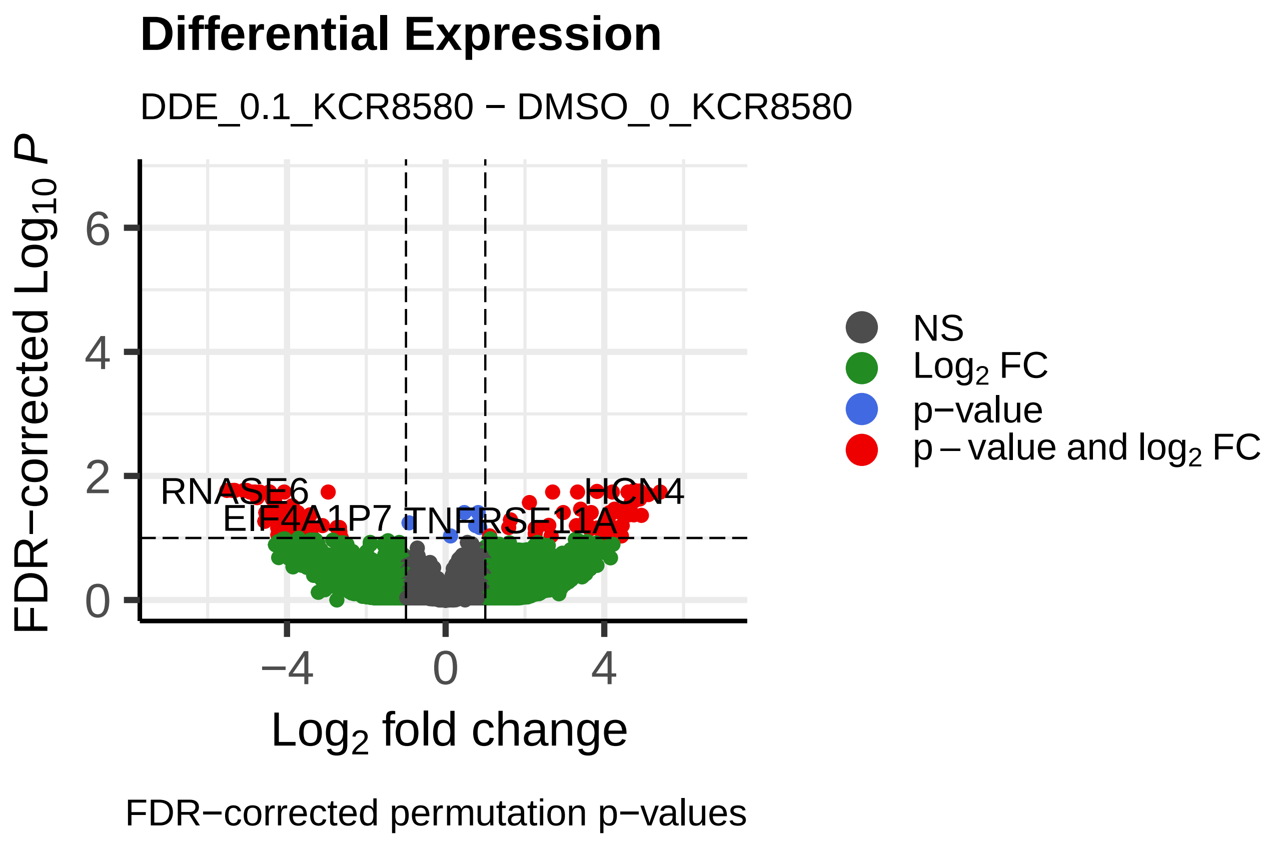
Differential Expression
DDE_0.1_KCR8580 − DMSO_0_KCR8580
FDR−corrected $Log_{10}$ P
Log2 fold change
NS
Log2 FC
p−value
p − value and log2 FC
RNASE6
EIF4A1P7
TNFRSF11A
HCN4
FDR−corrected permutation p−values

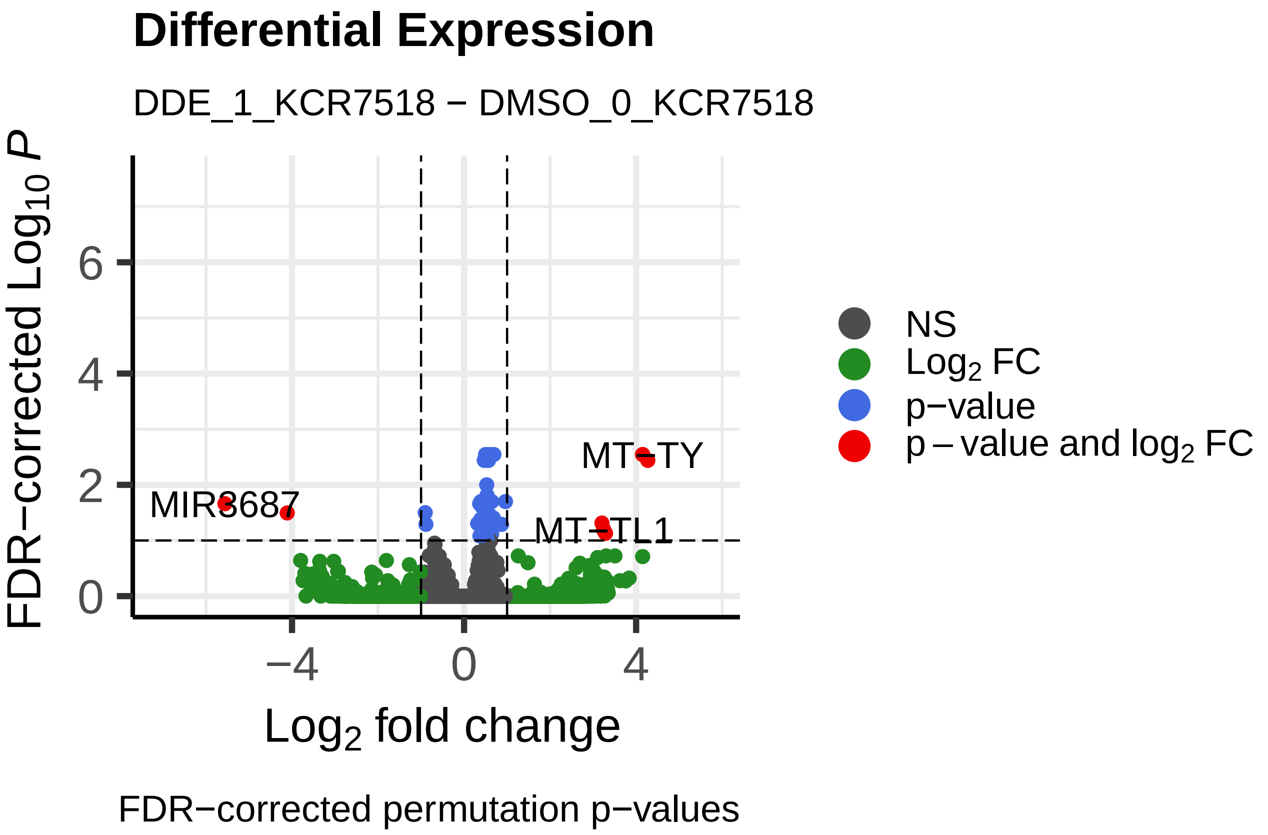
Differential Expression
DDE_1_KCR7518 − DMSO_0_KCR7518
FDR−corrected Log10 P
Log2 fold change
MIR3687
MT−TY
MT−TL1
NS
Log2 FC
p−value
p − value and log2 FC
FDR−corrected permutation p−values

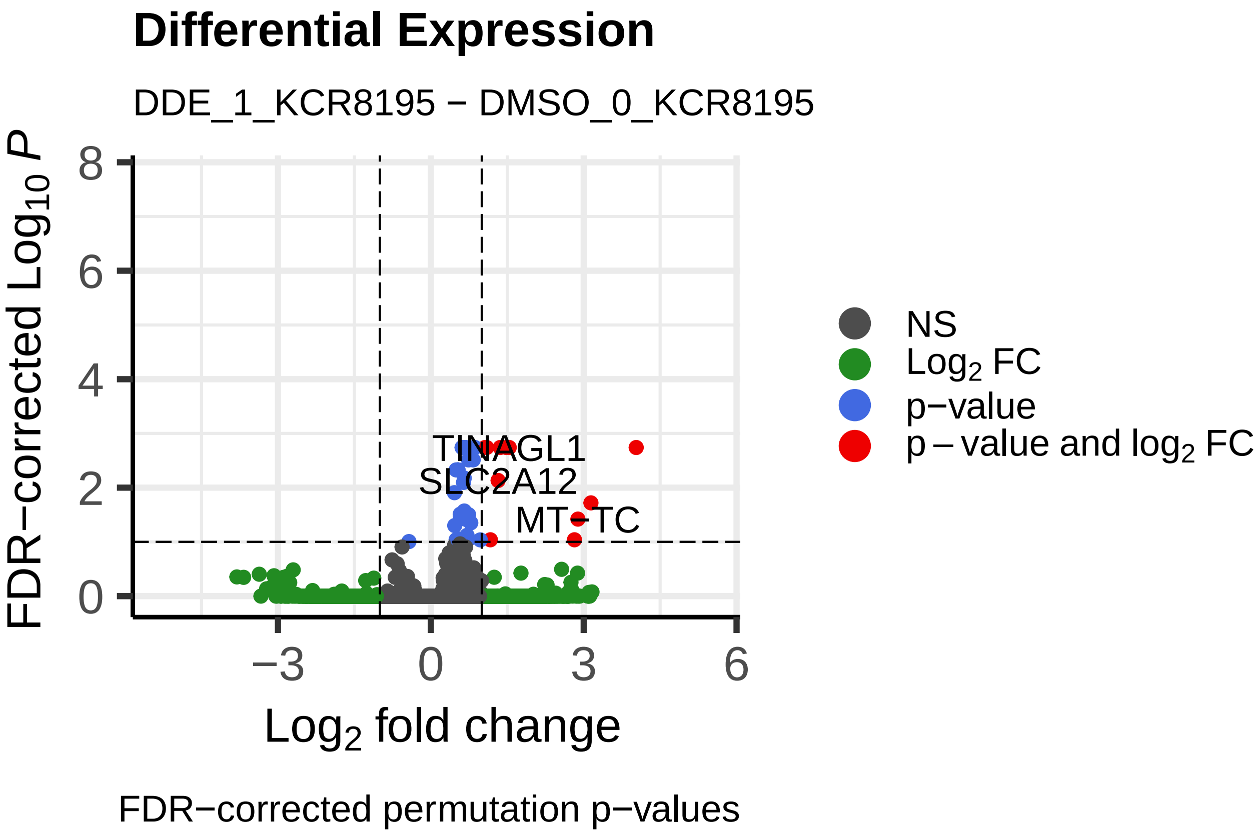
Differential Expression
DDE_1_KCR8195 − DMSO_0_KCR8195
FDR−corrected Log10 P
Log2 fold change
TINAGL1
SLC2A12
MT−TC
NS
Log2 FC
p−value
p − value and log2 FC
FDR−corrected permutation p−values

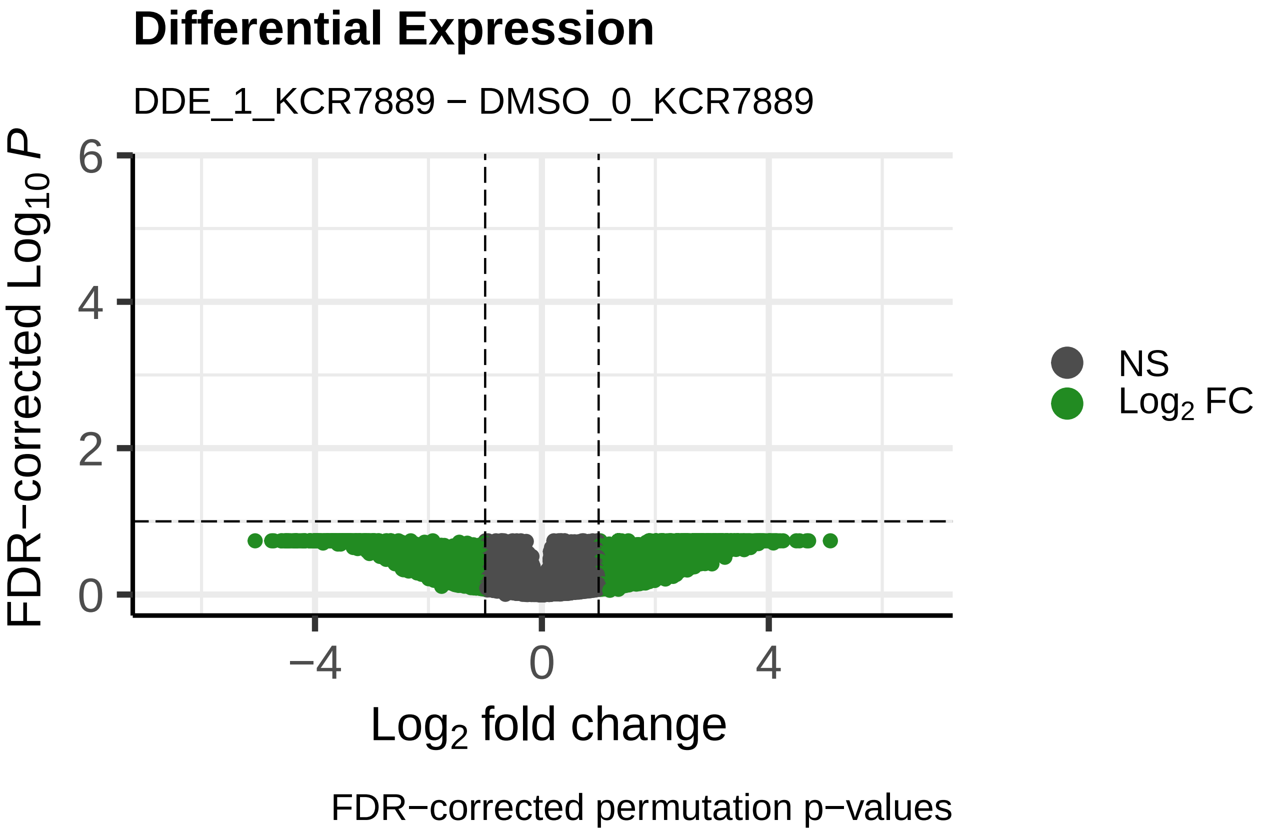
Differential Expression
DDE_1_KCR7889 − DMSO_0_KCR7889
FDR−corrected Log10 P
Log2 fold change
NS
Log2 FC
FDR−corrected permutation p−values

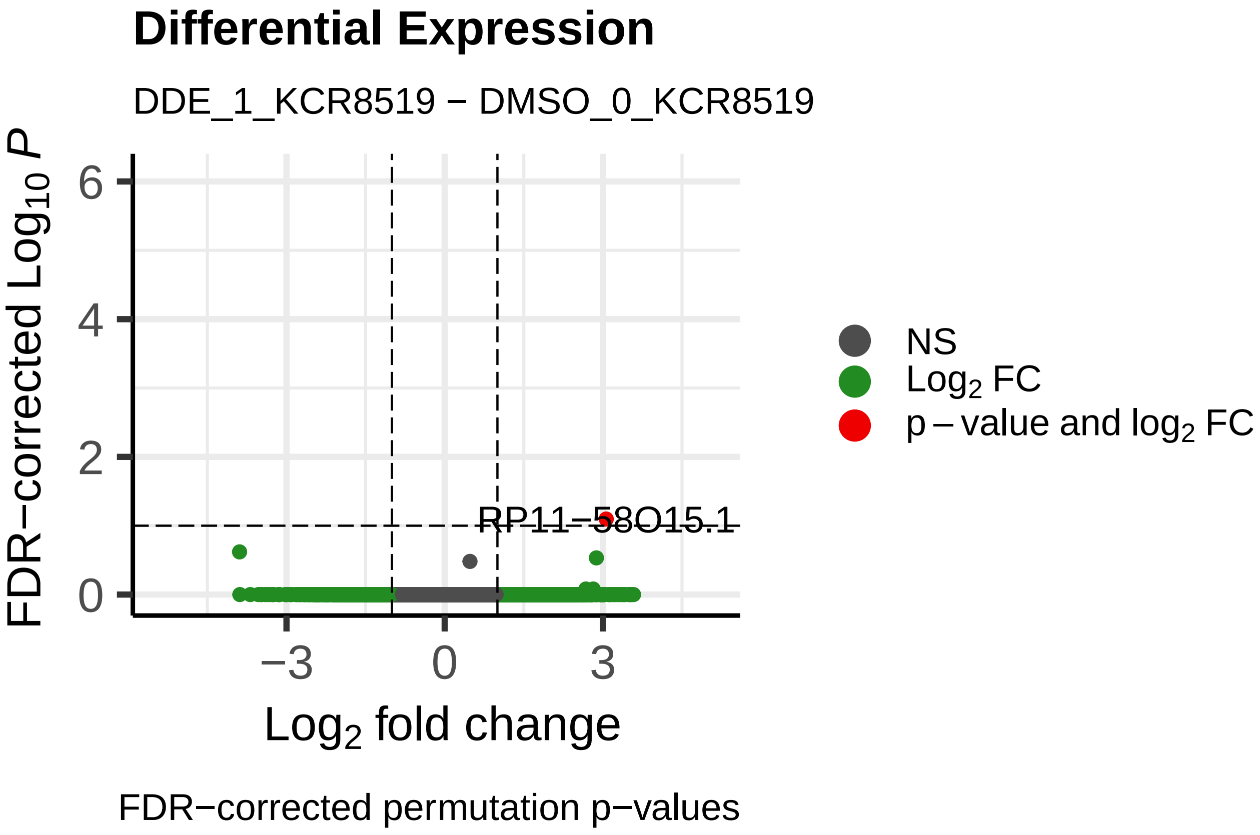
Differential Expression
DDE_1_KCR8519 − DMSO_0_KCR8519
FDR−corrected Log10 P
Log2 fold change
RP11−58O15.1
NS
Log2 FC
p − value and log2 FC
FDR−corrected permutation p−values

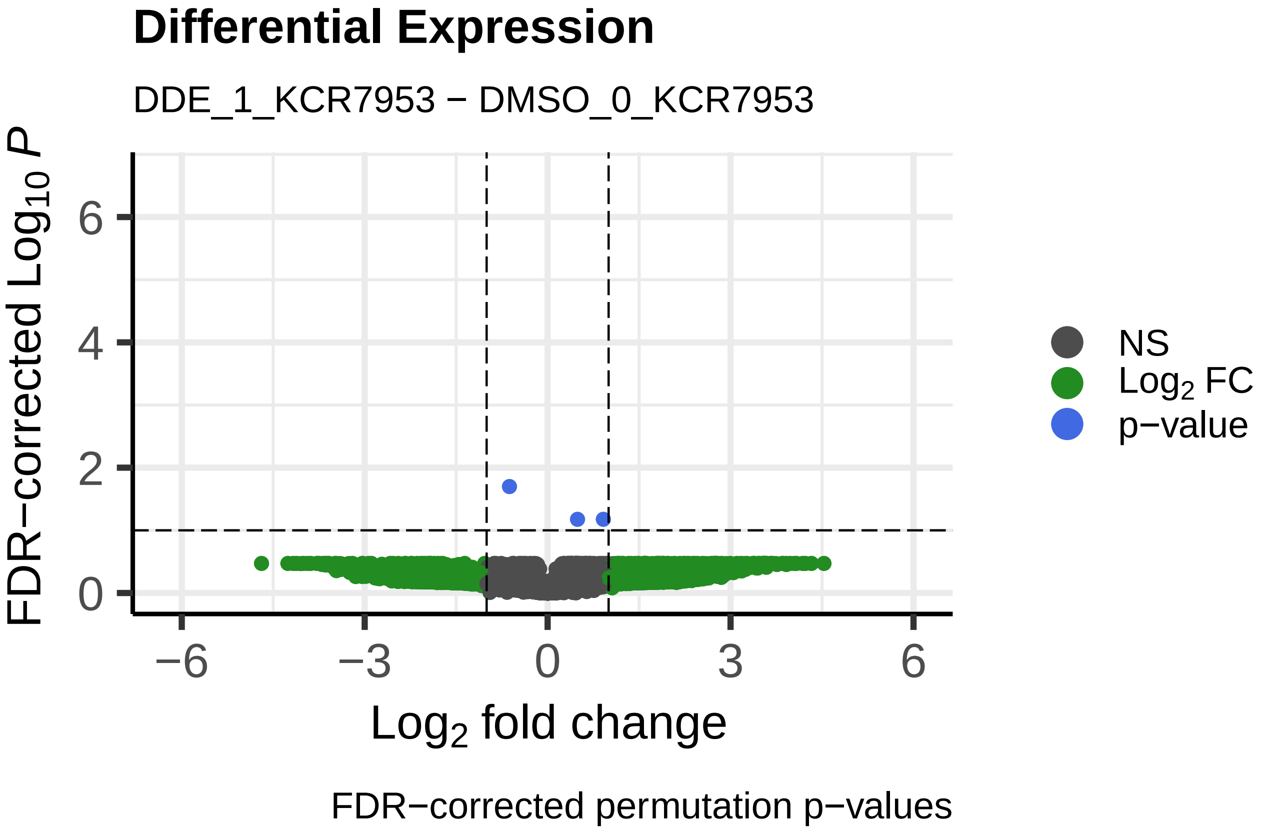
Differential Expression
DDE_1_KCR7953 − DMSO_0_KCR7953
FDR−corrected Log10 P
Log2 fold change
NS
Log2 FC
p−value
FDR−corrected permutation p−values

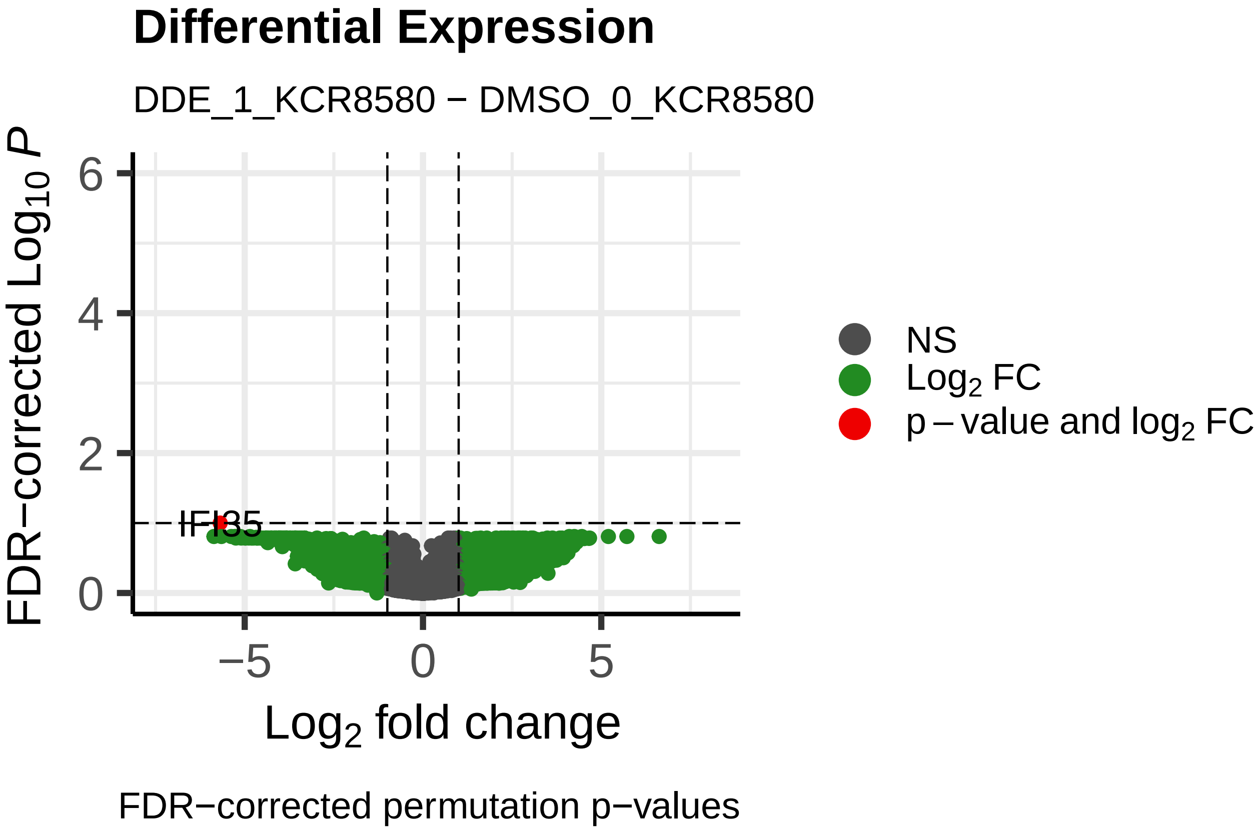
Differential Expression
DDE_1_KCR8580 − DMSO_0_KCR8580
FDR−corrected Log10 P
Log2 fold change
IFI35
NS
Log2 FC
p − value and log2 FC
FDR−corrected permutation p−values

**Differential Expression**

DDE_10_KCR7518 − DMSO_0_KCR7518

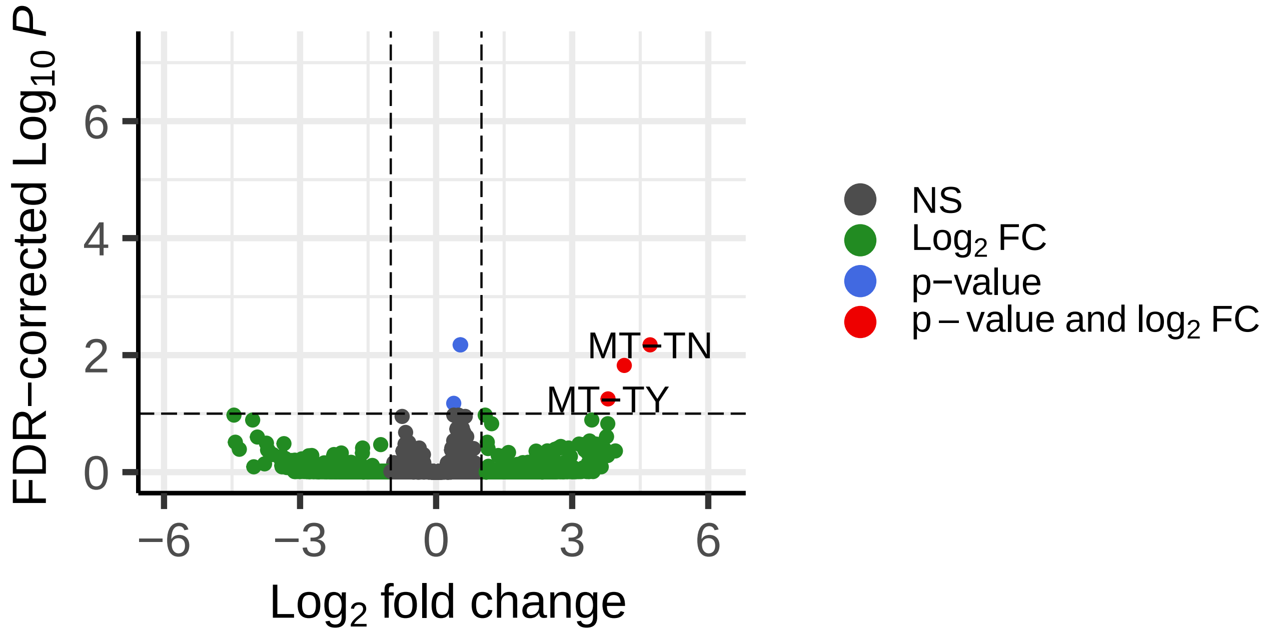


FDR−corrected permutation p−values

**Differential Expression**

DDE_10_KCR8195 − DMSO_0_KCR8195

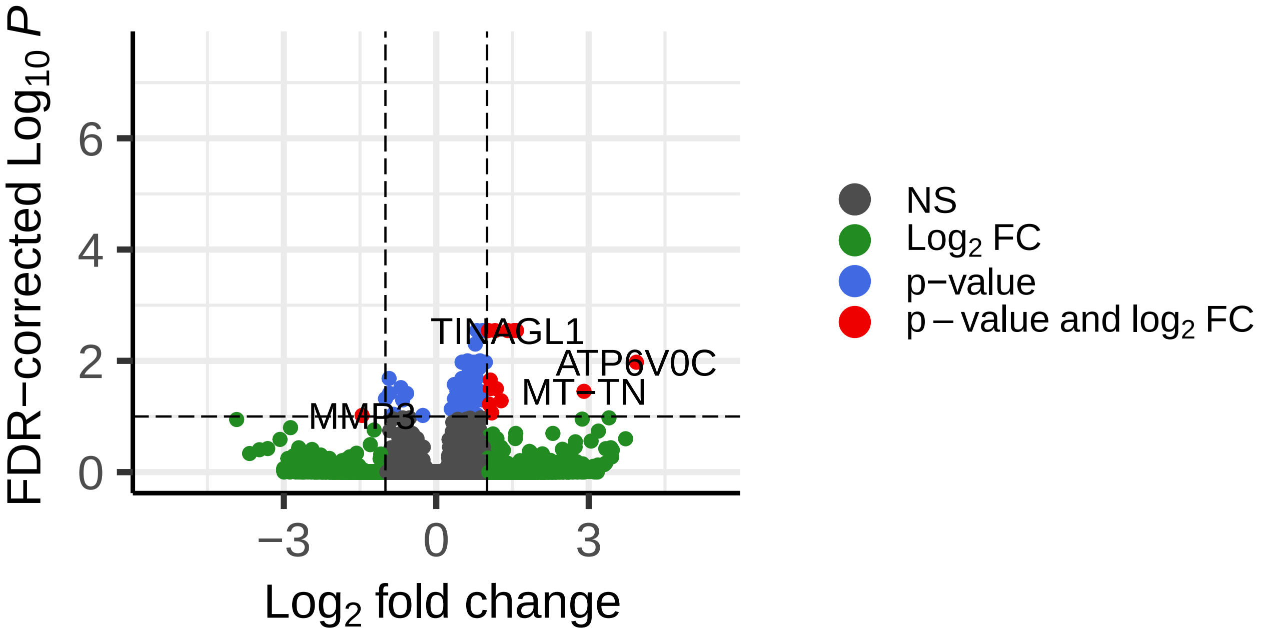


FDR−corrected permutation p−values

**Differential Expression**

DDE_10_KCR7889 − DMSO_0_KCR7889

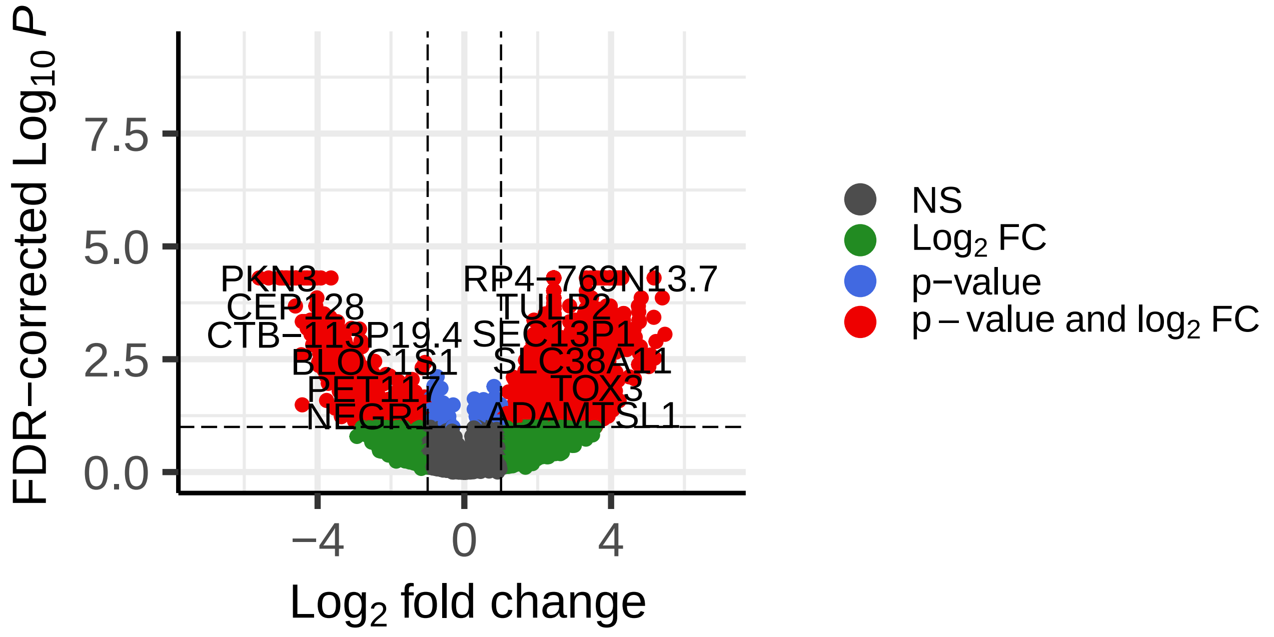


FDR−corrected permutation p−values

**Differential Expression**

DDE_10_KCR8519 − DMSO_0_KCR8519

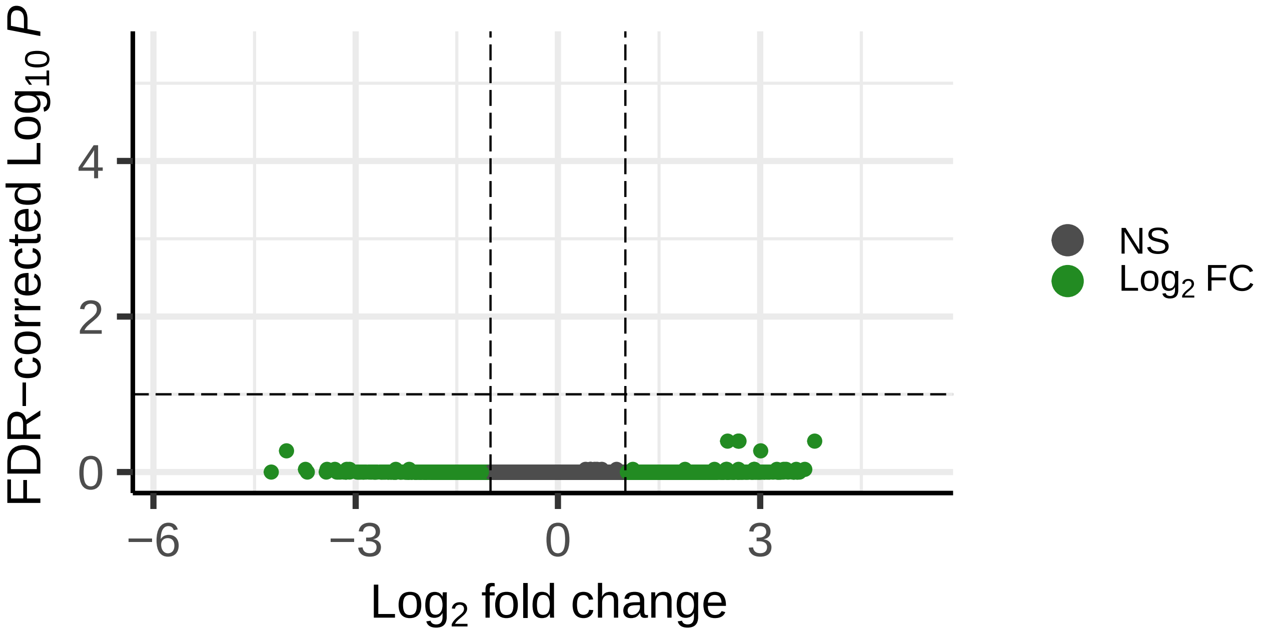


FDR−corrected permutation p−values

**Differential Expression**

DDE_10_KCR7953 − DMSO_0_KCR7953

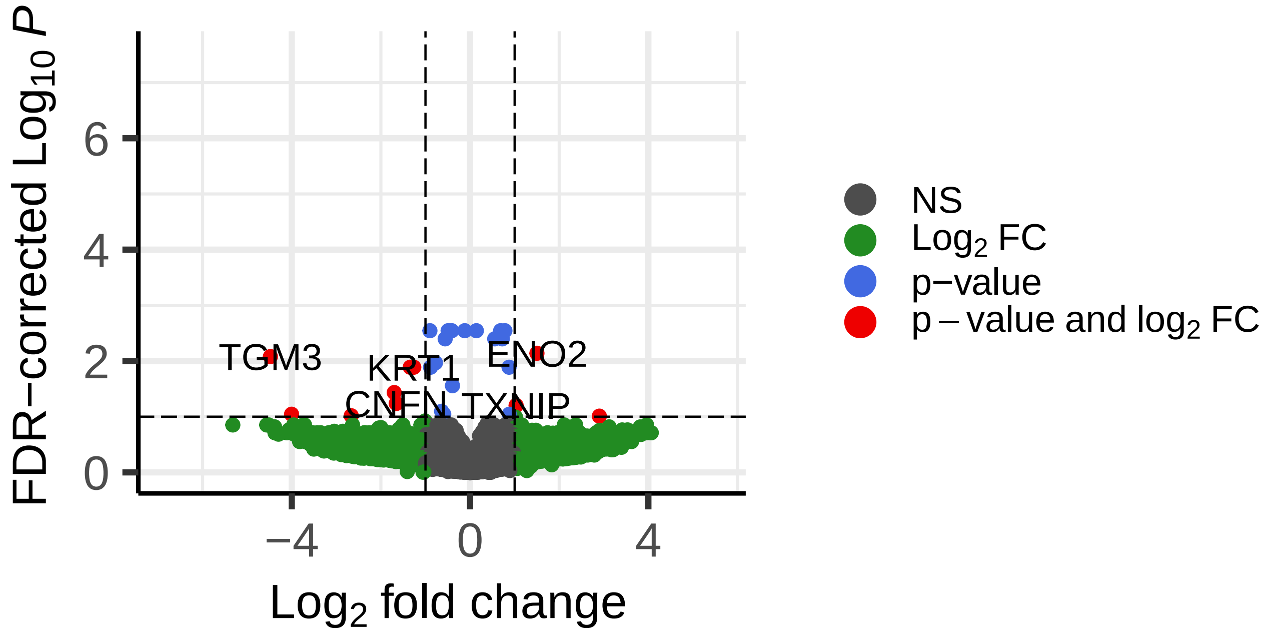


FDR−corrected permutation p−values

**Differential Expression**

DDE_10_KCR8580 − DMSO_0_KCR8580

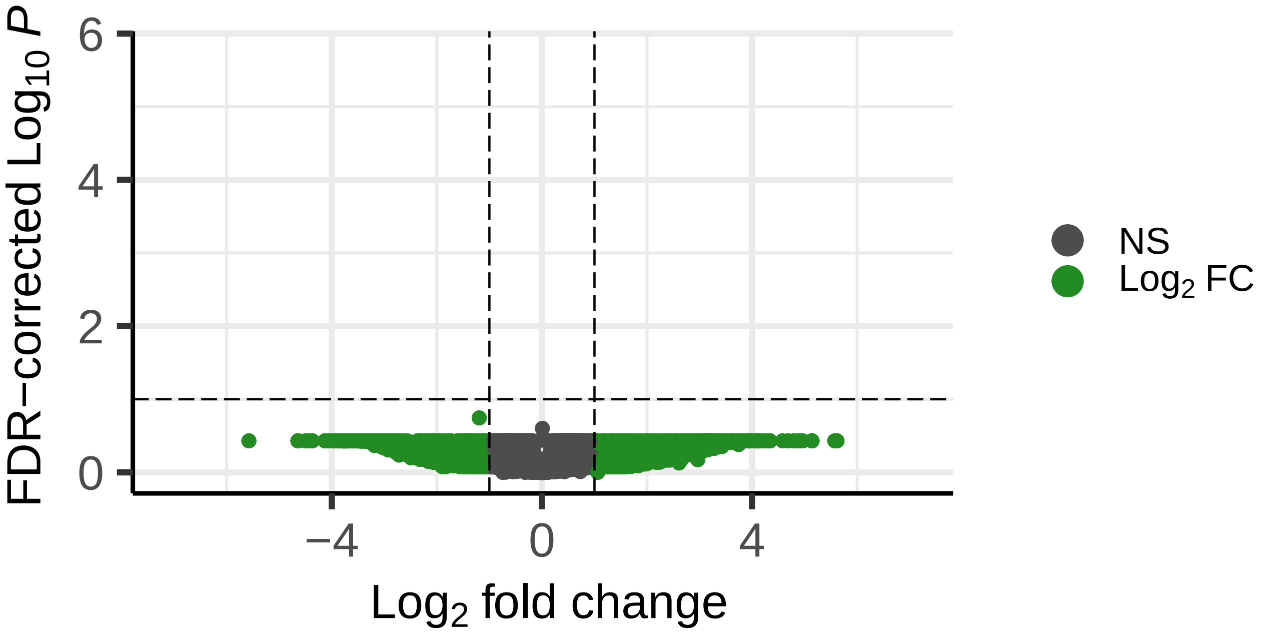


FDR−corrected permutation p−values

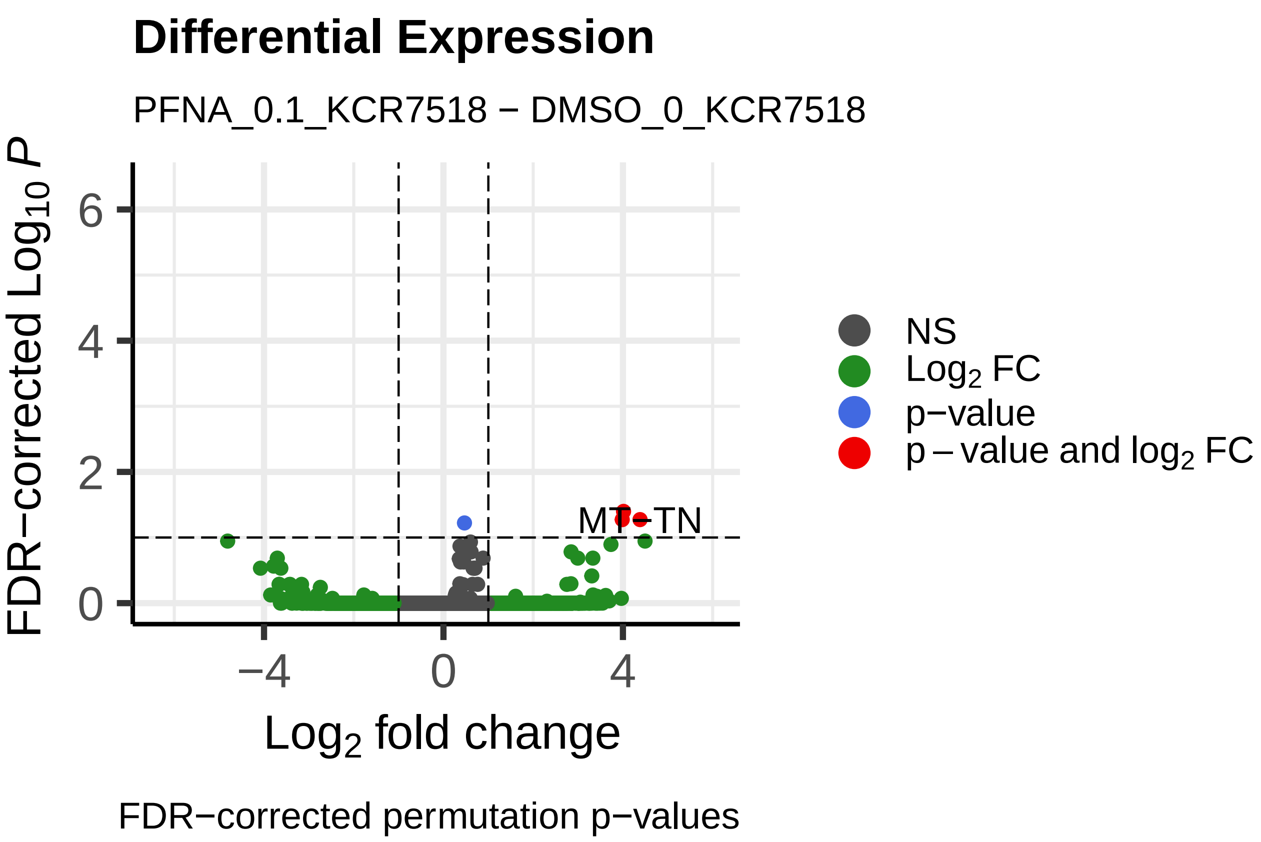
Differential Expression
PFNA_0.1_KCR7518 − DMSO_0_KCR7518
FDR−corrected $Log_{10}$ $P$
0
2
4
6
−4
0
4
$Log_2$ fold change
MT-TN
NS
$Log_2$ FC
p−value
p − value and $log_2$ FC
FDR−corrected permutation p−values

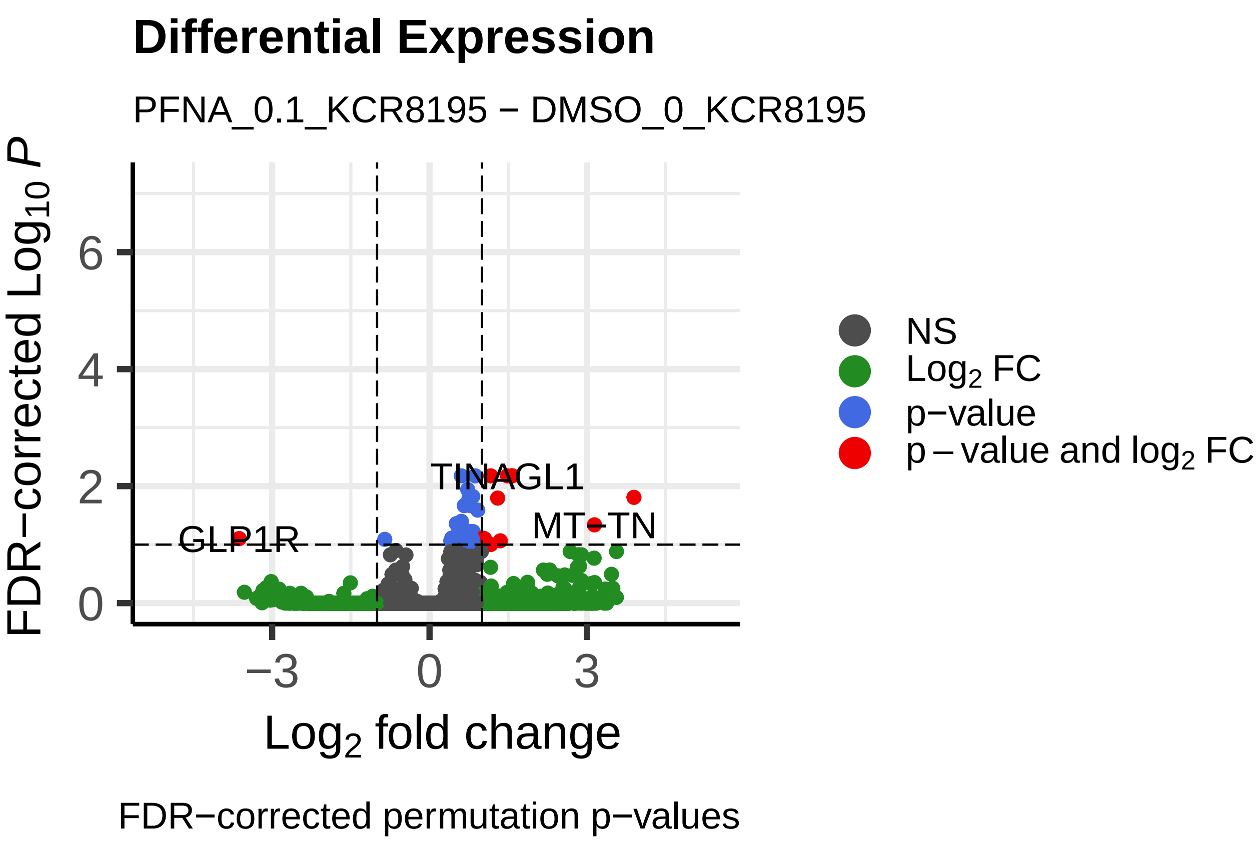
Differential Expression
PFNA_0.1_KCR8195 − DMSO_0_KCR8195
FDR−corrected $Log_{10}$ $P$
0
2
4
6
−3
0
3
$Log_2$ fold change
TINAGL1
GLP1R
MT-TN
NS
$Log_2$ FC
p−value
p − value and $log_2$ FC
FDR−corrected permutation p−values

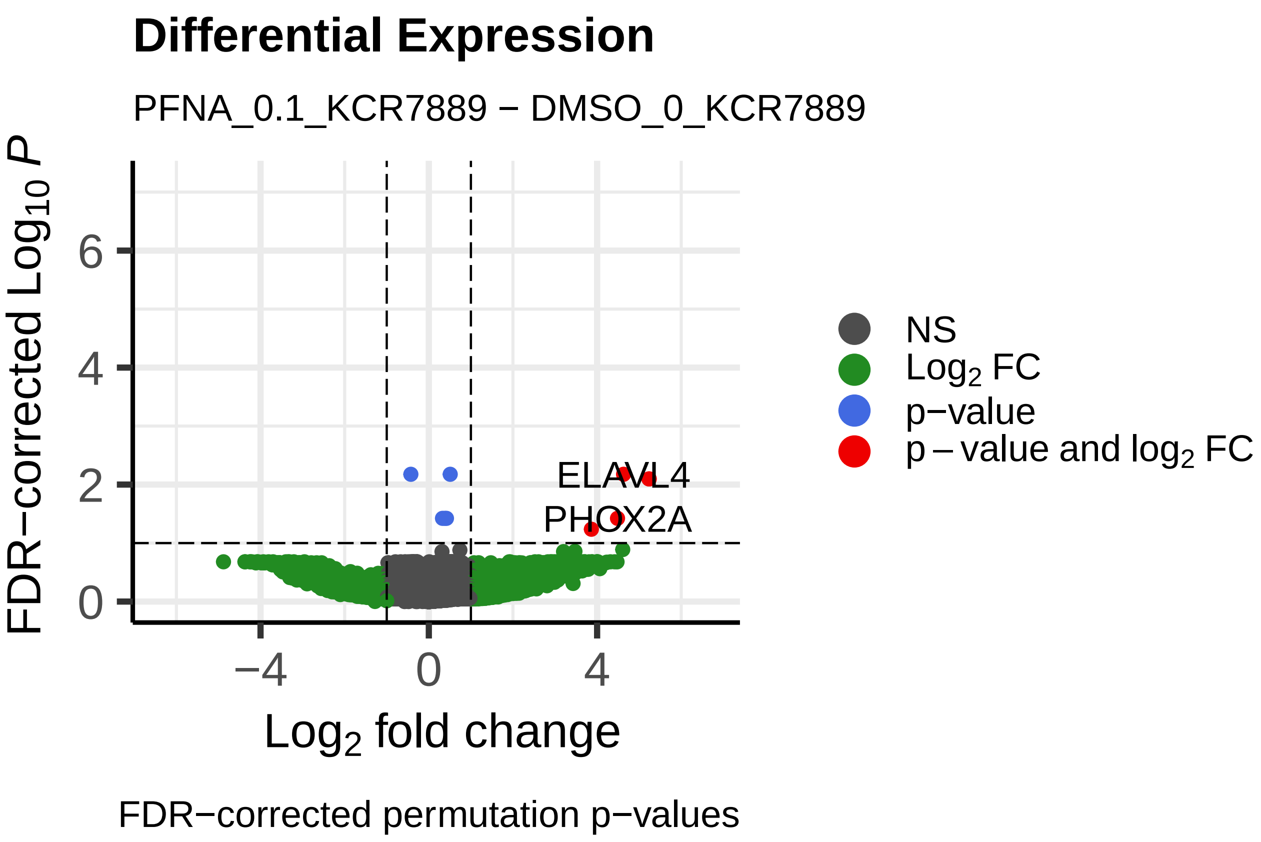
Differential Expression
PFNA_0.1_KCR7889 − DMSO_0_KCR7889
FDR−corrected $Log_{10}$ $P$
0
2
4
6
−4
0
4
$Log_2$ fold change
ELAVL4
PHOX2A
NS
$Log_2$ FC
p−value
p − value and $log_2$ FC
FDR−corrected permutation p−values

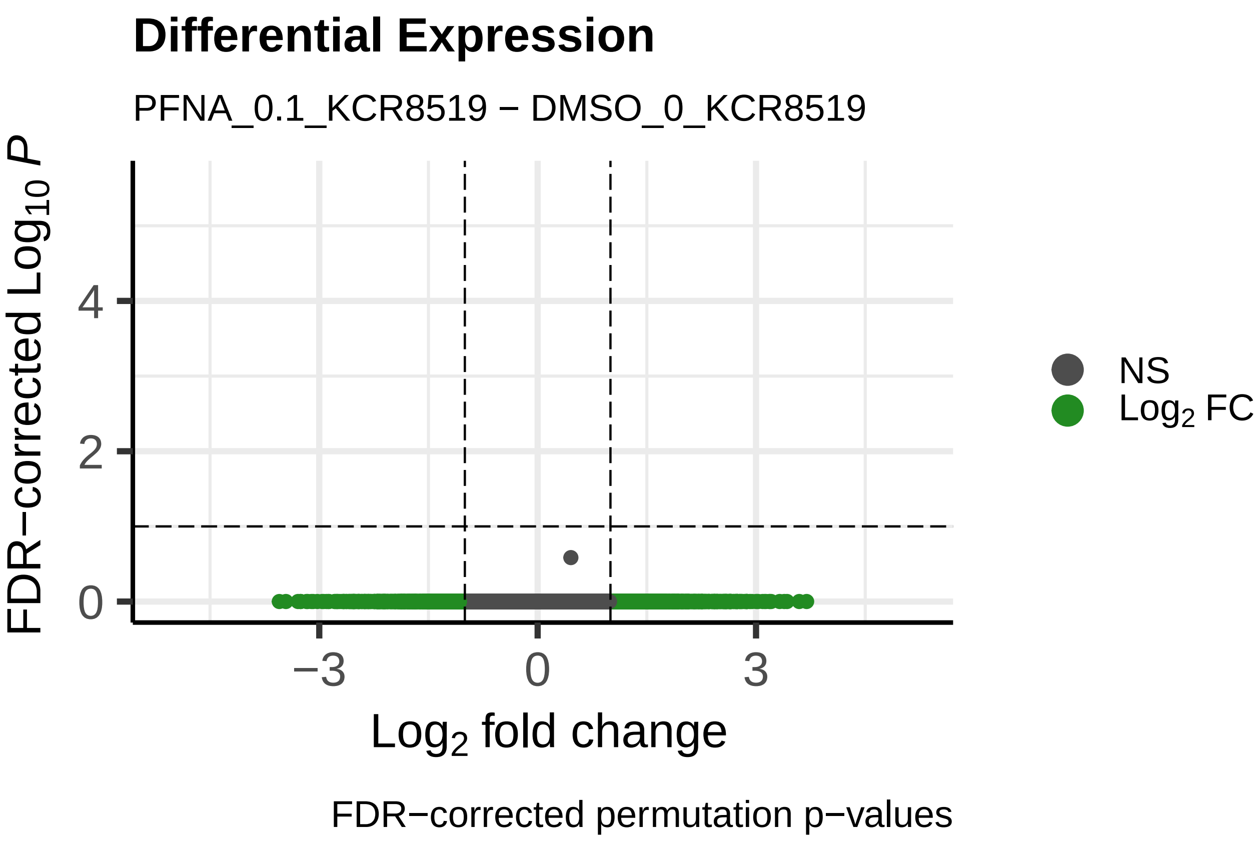
Differential Expression
PFNA_0.1_KCR8519 − DMSO_0_KCR8519
FDR−corrected $Log_{10}$ $P$
0
2
4
−3
0
3
$Log_2$ fold change
NS
$Log_2$ FC
FDR−corrected permutation p−values

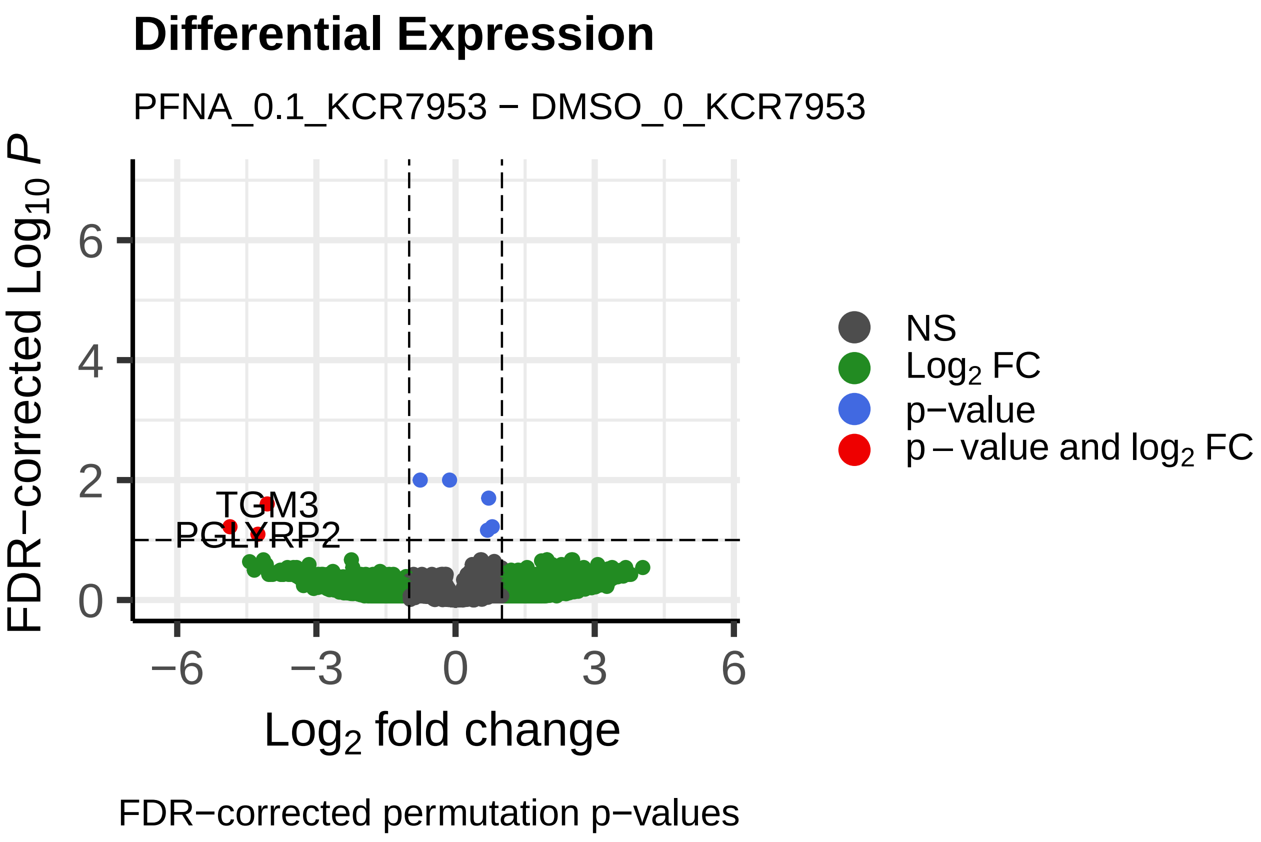
Differential Expression
PFNA_0.1_KCR7953 − DMSO_0_KCR7953
FDR−corrected $Log_{10}$ $P$
0
2
4
6
−6
−3
0
3
6
$Log_2$ fold change
TGM3
PGLYRP2
NS
$Log_2$ FC
p−value
p − value and $log_2$ FC
FDR−corrected permutation p−values

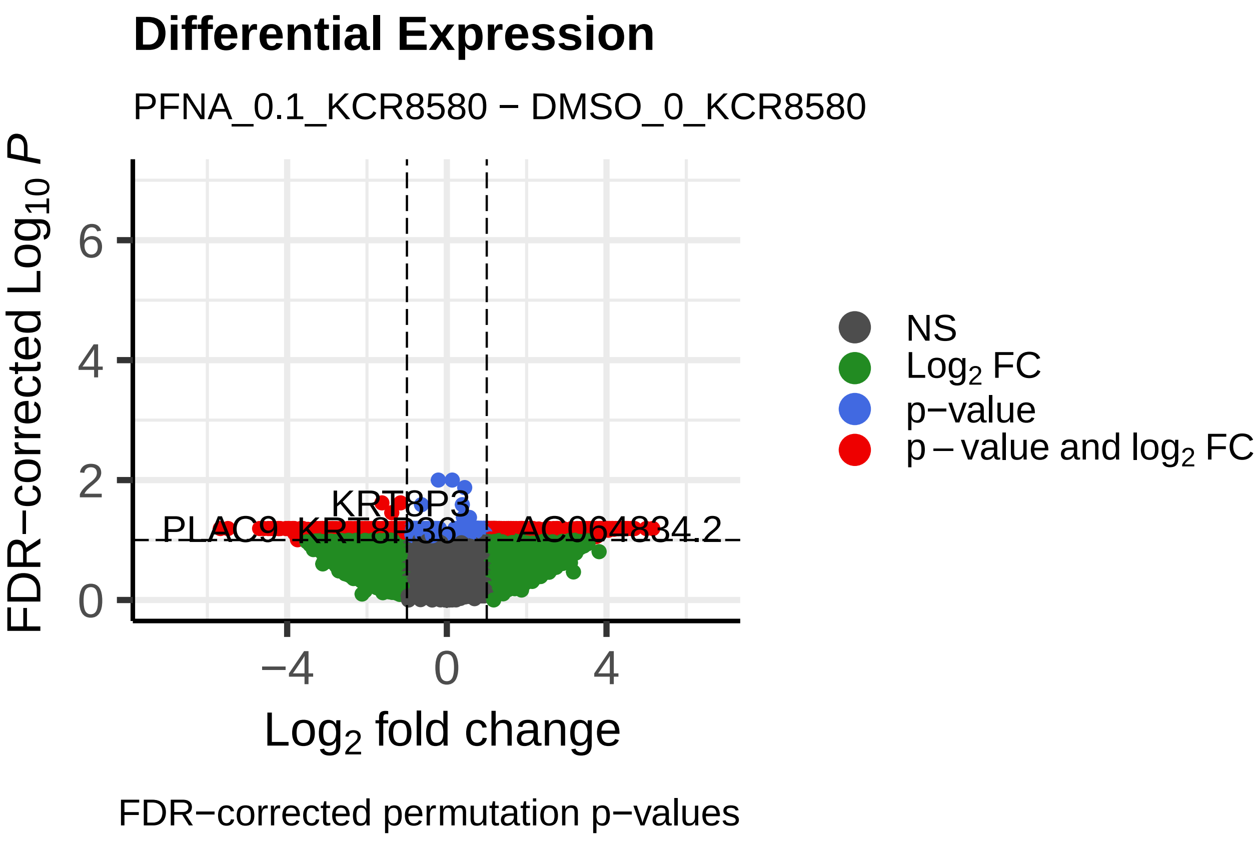
Differential Expression
PFNA_0.1_KCR8580 − DMSO_0_KCR8580
FDR−corrected $Log_{10}$ $P$
0
2
4
6
−4
0
4
$Log_2$ fold change
KRT8P3
PLAC9
KRT8P36
NS
$Log_2$ FC
p−value
p − value and $log_2$ FC
FDR−corrected permutation p−values

**Differential Expression**

PFNA_1_KCR7518 − DMSO_0_KCR7518

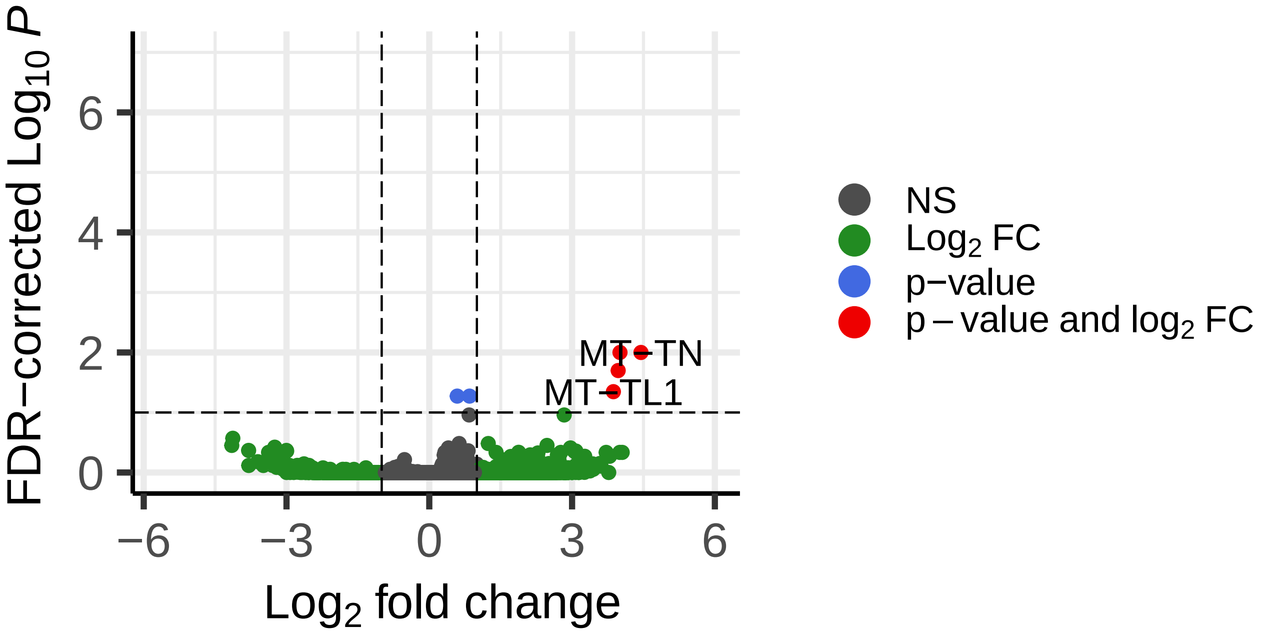


FDR−corrected permutation p−values

**Differential Expression**

PFNA_1_KCR8195 − DMSO_0_KCR8195

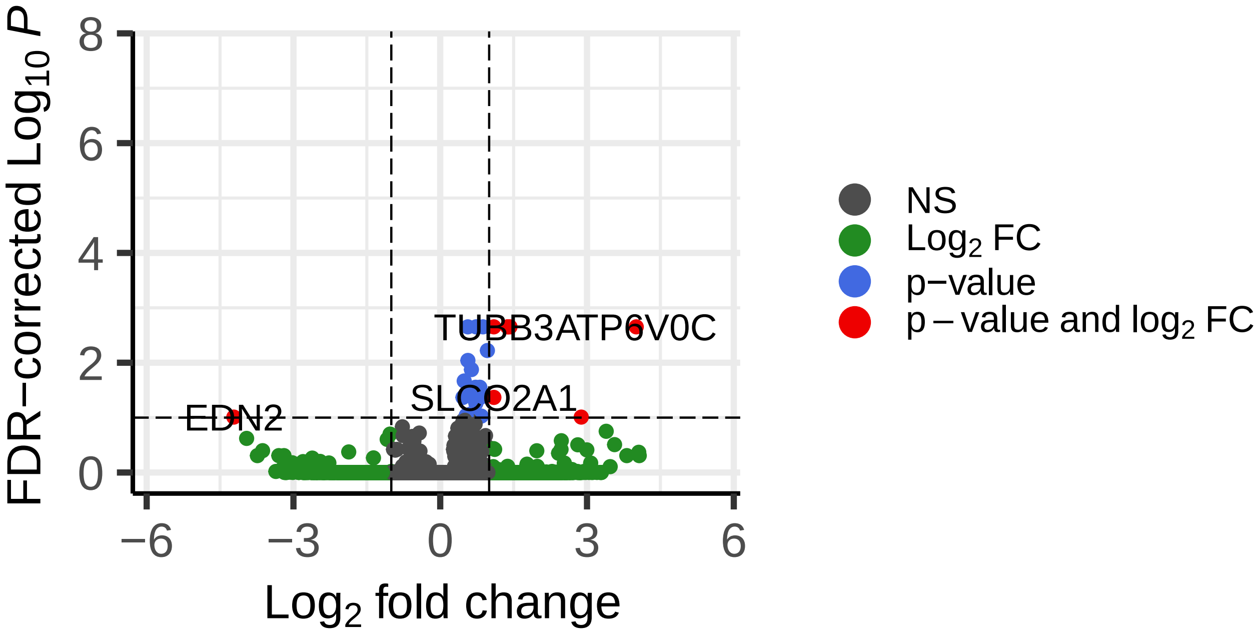


FDR−corrected permutation p−values

**Differential Expression**

PFNA_1_KCR7889 − DMSO_0_KCR7889

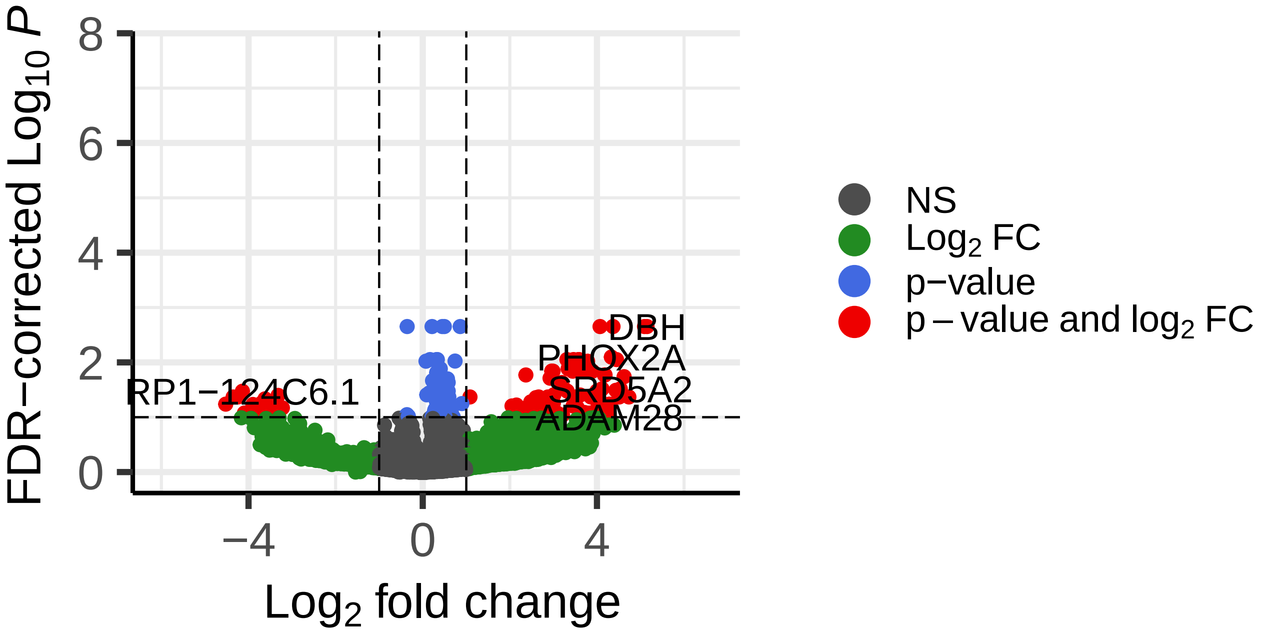


FDR−corrected permutation p−values

**Differential Expression**

PFNA_1_KCR8519 − DMSO_0_KCR8519

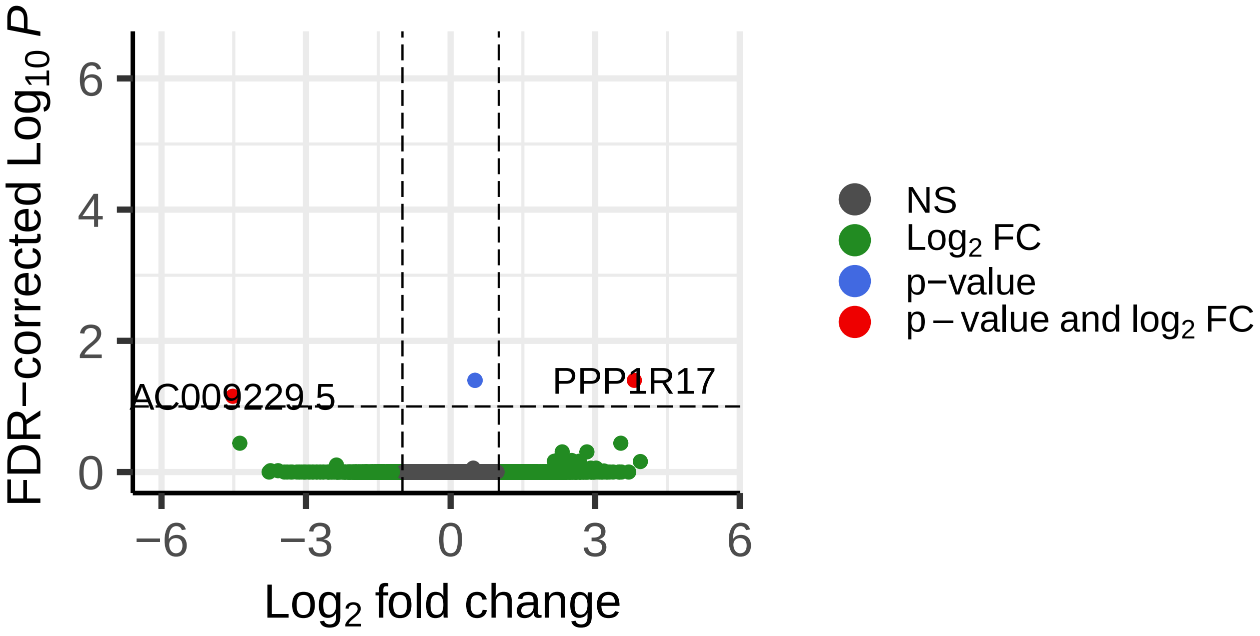


FDR−corrected permutation p−values

**Differential Expression**

PFNA_1_KCR7953 − DMSO_0_KCR7953

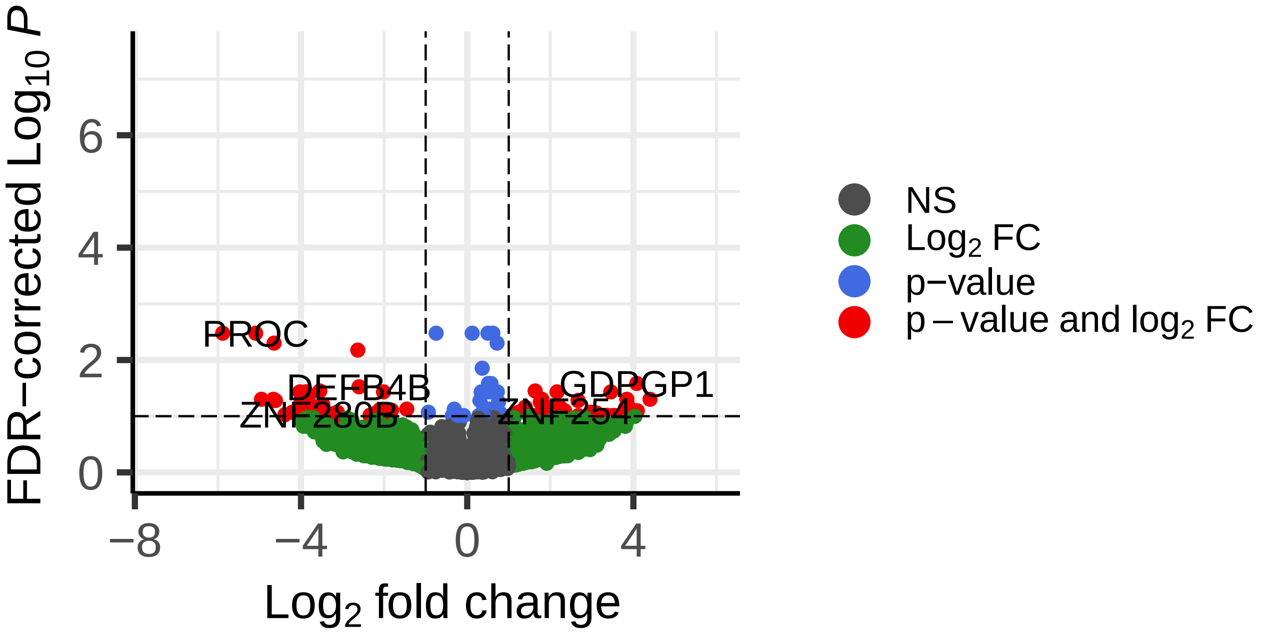


FDR−corrected permutation p−values

**Differential Expression**

PFNA_1_KCR8580 − DMSO_0_KCR8580

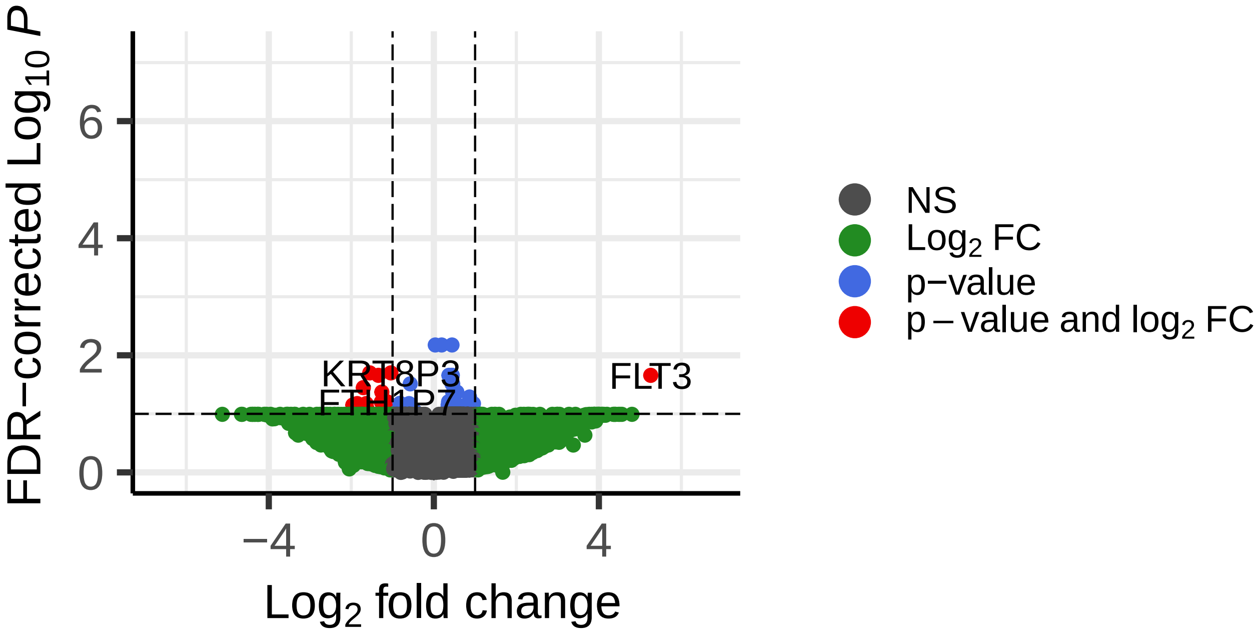


FDR−corrected permutation p−values

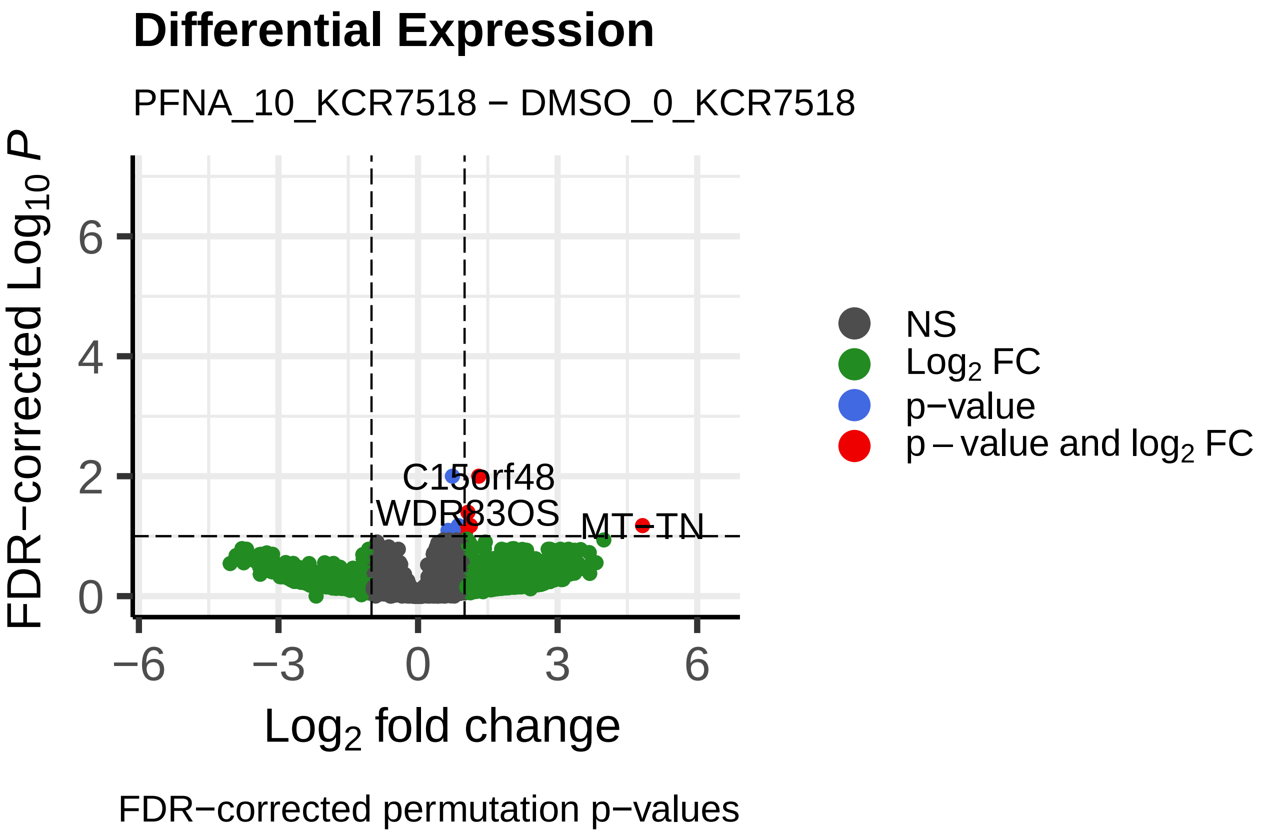
Differential Expression
PFNA_10_KCR7518 − DMSO_0_KCR7518
FDR−corrected $Log_{10}$ P
Log2 fold change
C15orf48
WDR83OS
MT−TN
NS
$Log_2$ FC
p−value
p − value and $log_2$ FC
FDR−corrected permutation p−values

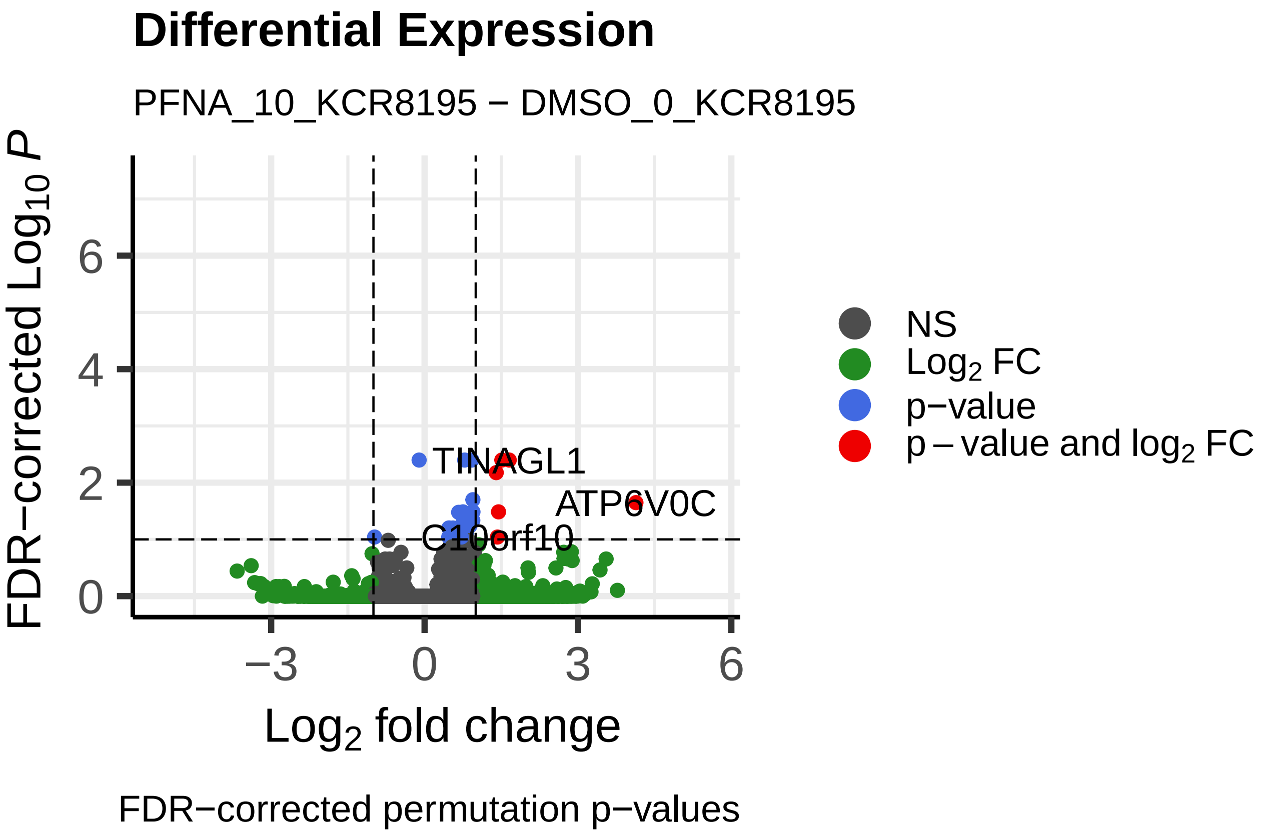
Differential Expression
PFNA_10_KCR8195 − DMSO_0_KCR8195
FDR−corrected $Log_{10}$ P
Log2 fold change
TINAGL1
ATP6V0C
C10orf10
NS
$Log_2$ FC
p−value
p − value and $log_2$ FC
FDR−corrected permutation p−values

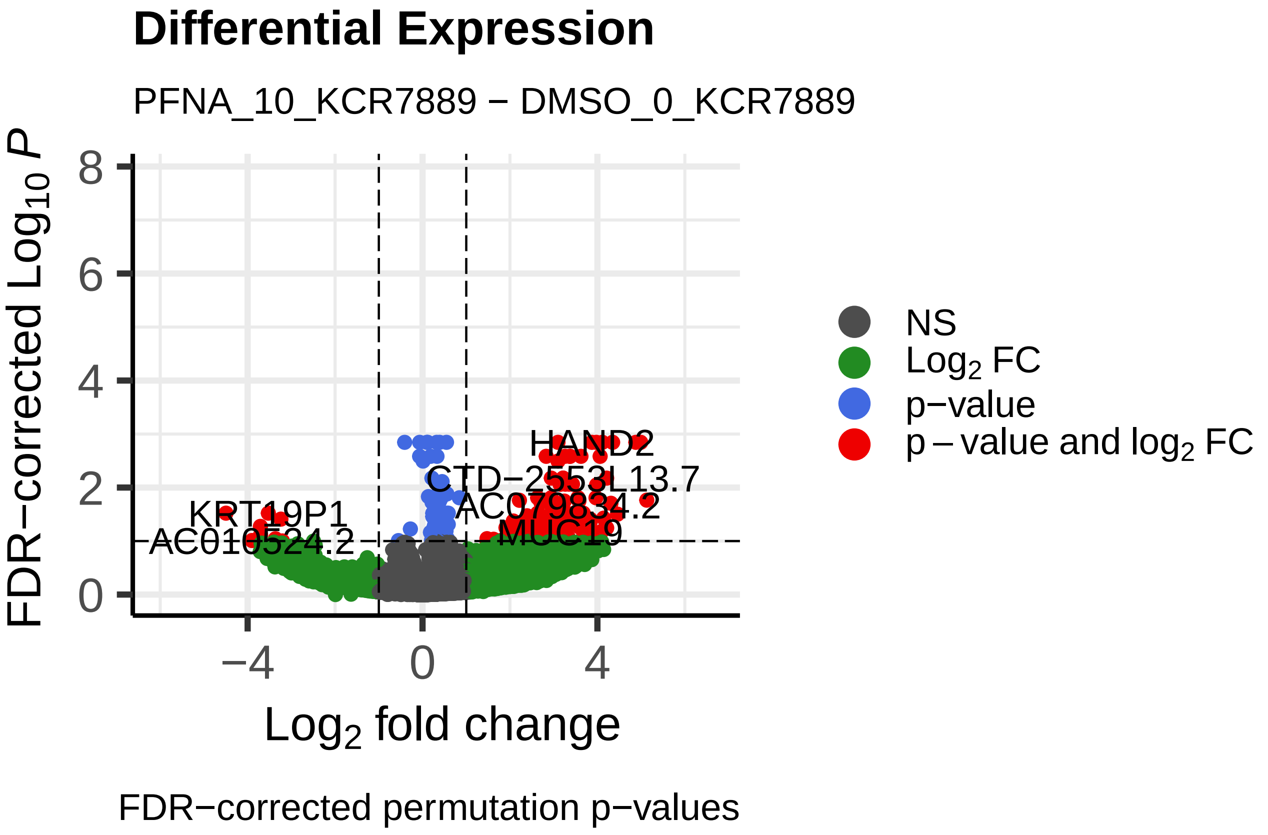
Differential Expression
PFNA_10_KCR7889 − DMSO_0_KCR7889
FDR−corrected $Log_{10}$ P
Log2 fold change
HAND2
CTD−2553L13.7
KRT19P1
AC010524.2
MUC19
NS
$Log_2$ FC
p−value
p − value and $log_2$ FC
FDR−corrected permutation p−values

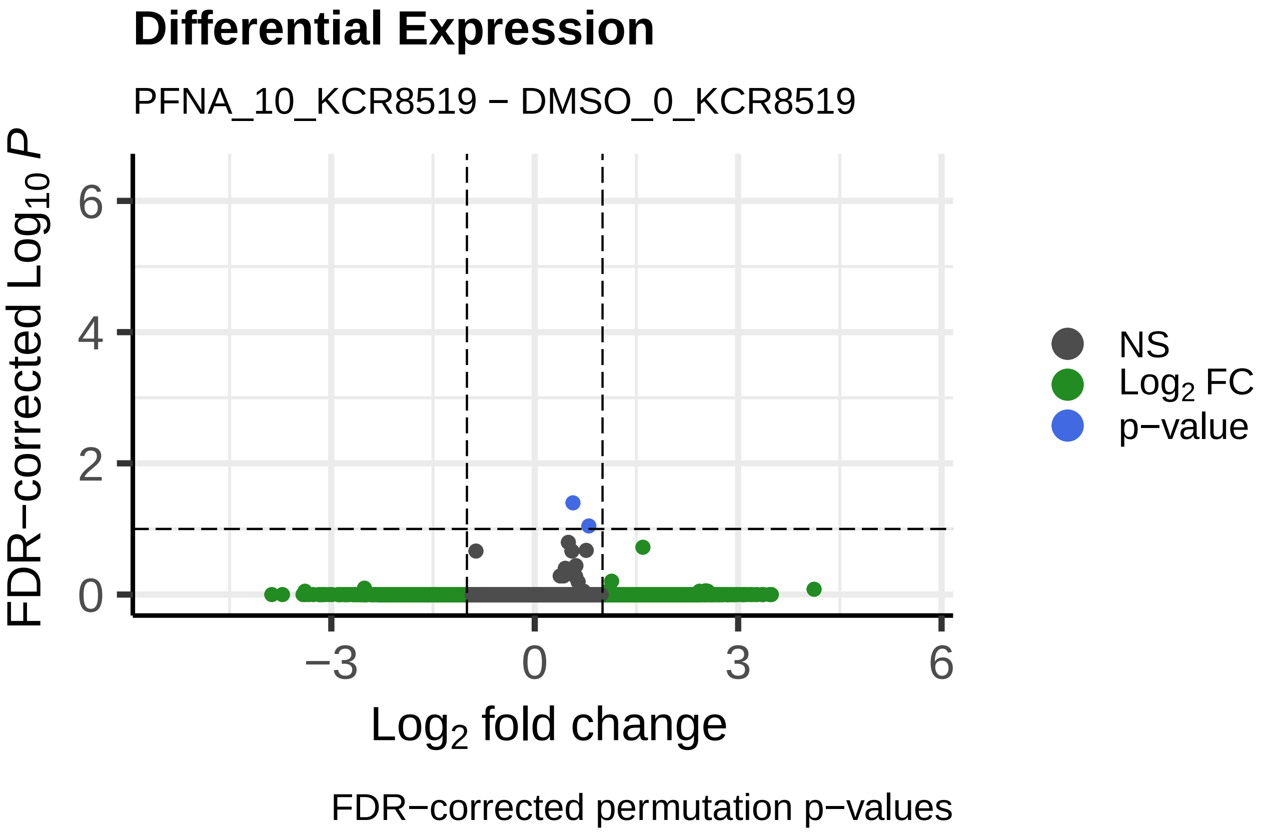
Differential Expression
PFNA_10_KCR8519 − DMSO_0_KCR8519
FDR−corrected $Log_{10}$ P
Log2 fold change
NS
$Log_2$ FC
p−value
FDR−corrected permutation p−values

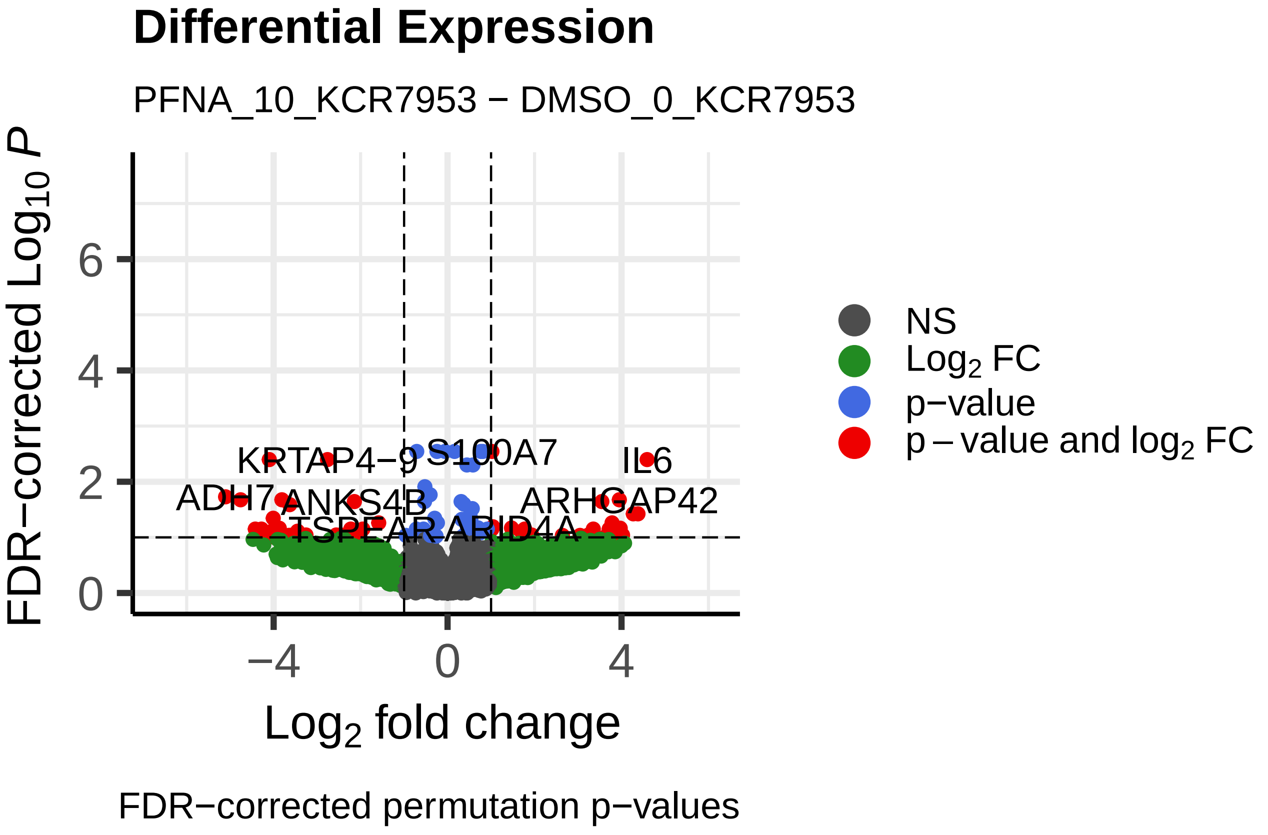
Differential Expression
PFNA_10_KCR7953 − DMSO_0_KCR7953
FDR−corrected $Log_{10}$ P
Log2 fold change
KRTAP4−9
S100A7
IL6
ADH7
ANKS4B
ARHGAP42
NS
$Log_2$ FC
p−value
p − value and $log_2$ FC
FDR−corrected permutation p−values

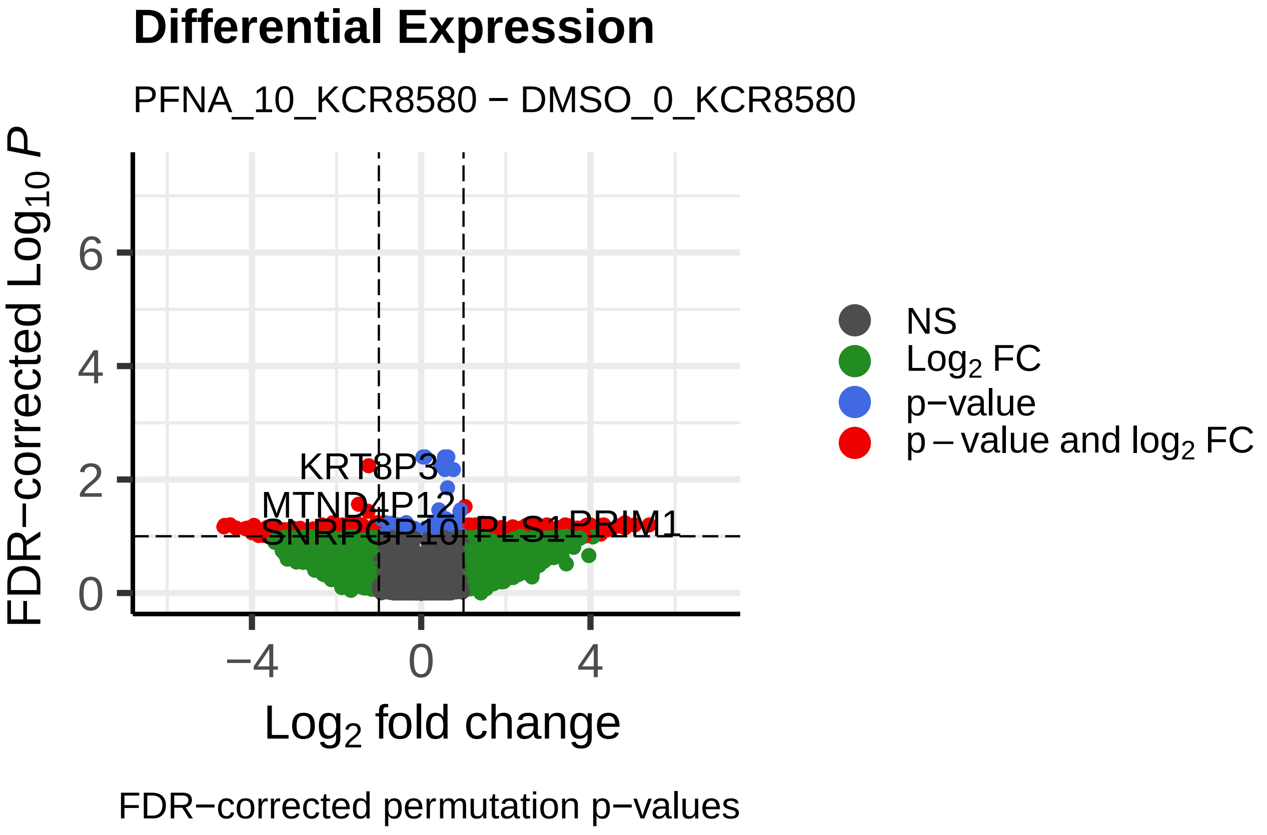
Differential Expression
PFNA_10_KCR8580 − DMSO_0_KCR8580
FDR−corrected $Log_{10}$ P
Log2 fold change
KRT8P3
MTND4P12
PRIM1
NS
$Log_2$ FC
p−value
p − value and $log_2$ FC
FDR−corrected permutation p−values

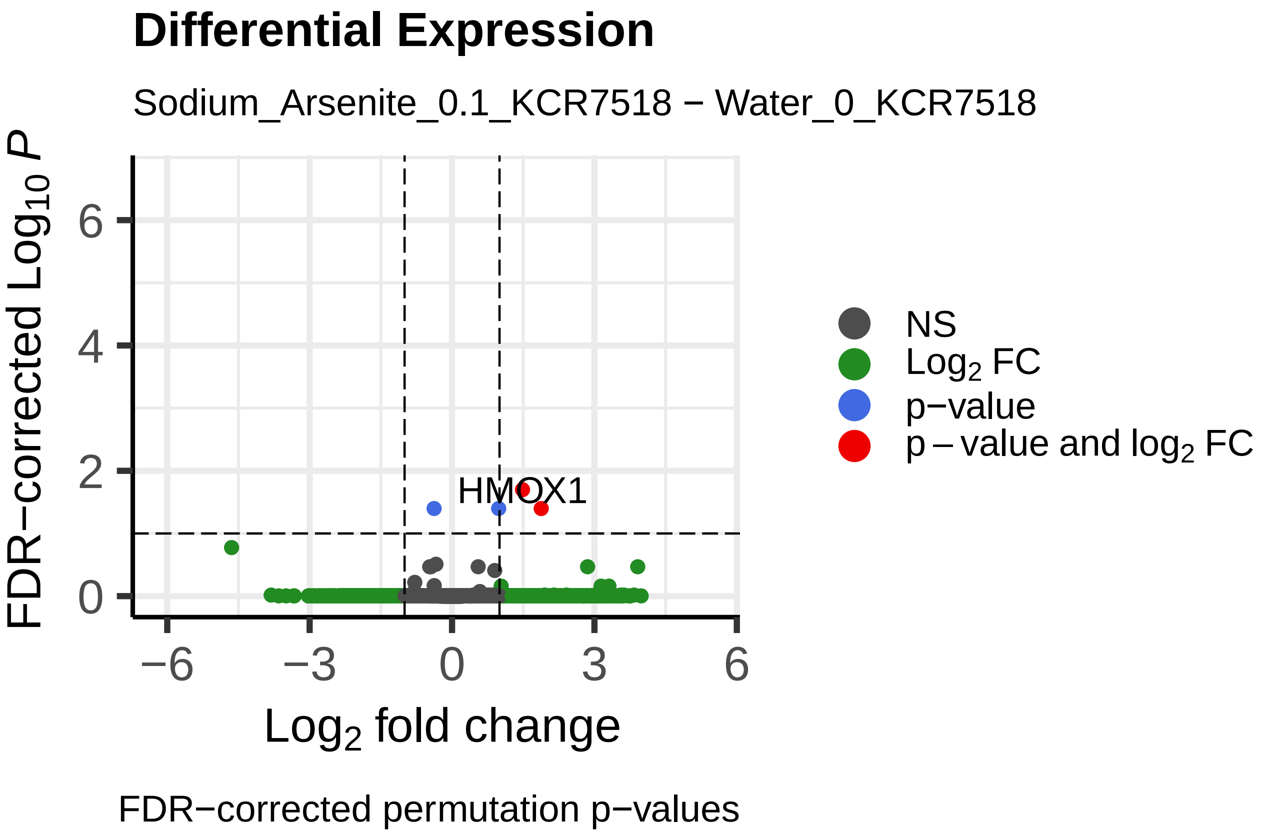
Differential Expression
Sodium_Arsenite_0.1_KCR7518 − Water_0_KCR7518
FDR−corrected Log10 P
Log2 fold change
HMOX1
NS
Log2 FC
p−value
p − value and log2 FC
FDR−corrected permutation p−values

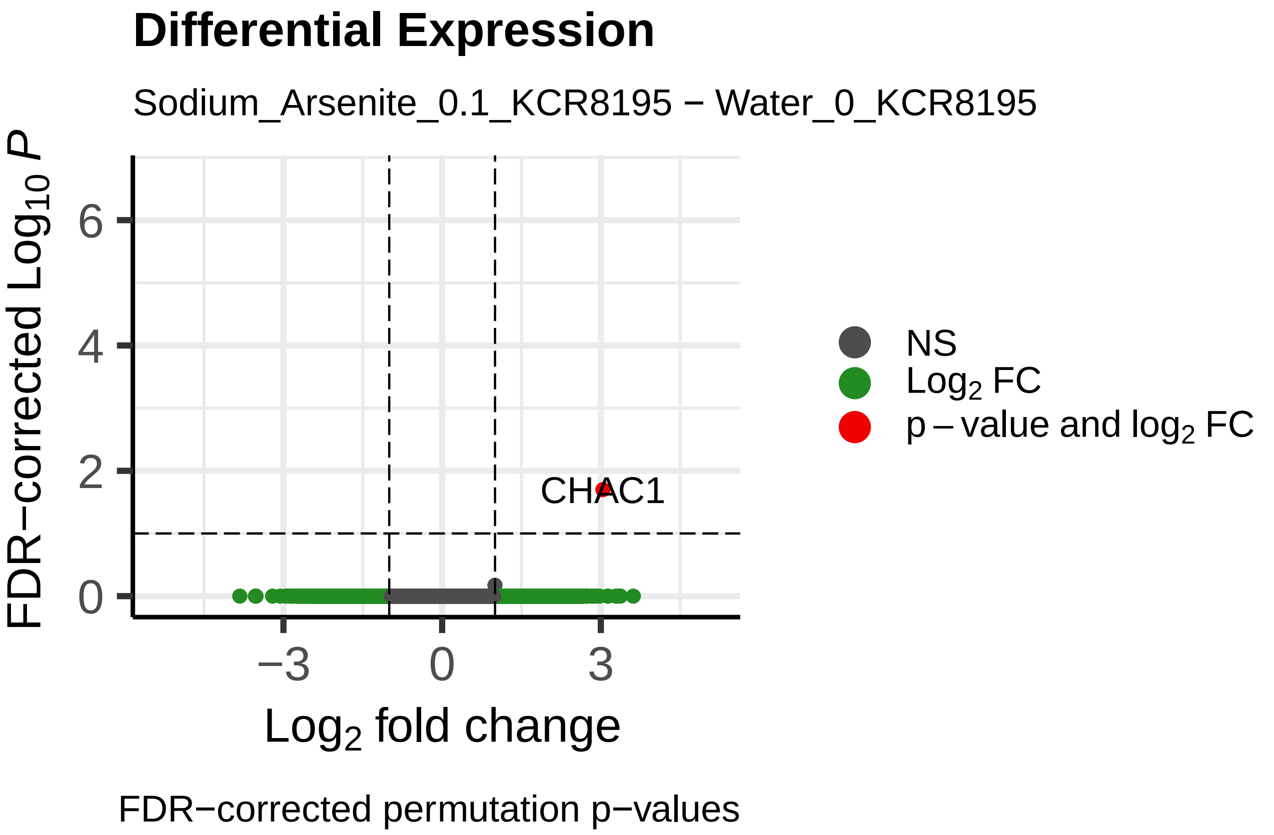
Differential Expression
Sodium_Arsenite_0.1_KCR8195 − Water_0_KCR8195
FDR−corrected Log10 P
Log2 fold change
CHAC1
NS
Log2 FC
p − value and log2 FC
FDR−corrected permutation p−values

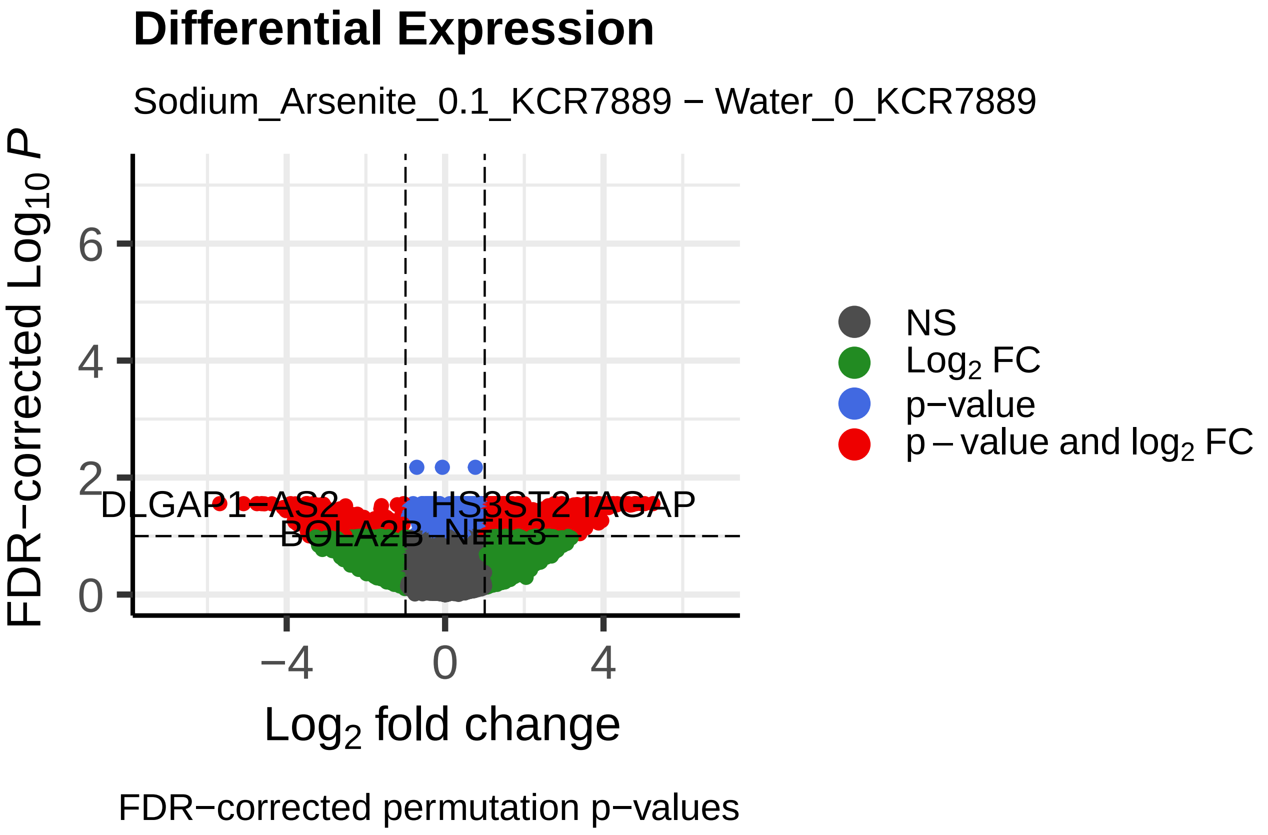
Differential Expression
Sodium_Arsenite_0.1_KCR7889 − Water_0_KCR7889
FDR−corrected Log10 P
Log2 fold change
DLGAP1−AS2
BOLA2B
HS3ST2
TAGAP
NEIL3
NS
Log2 FC
p−value
p − value and log2 FC
FDR−corrected permutation p−values

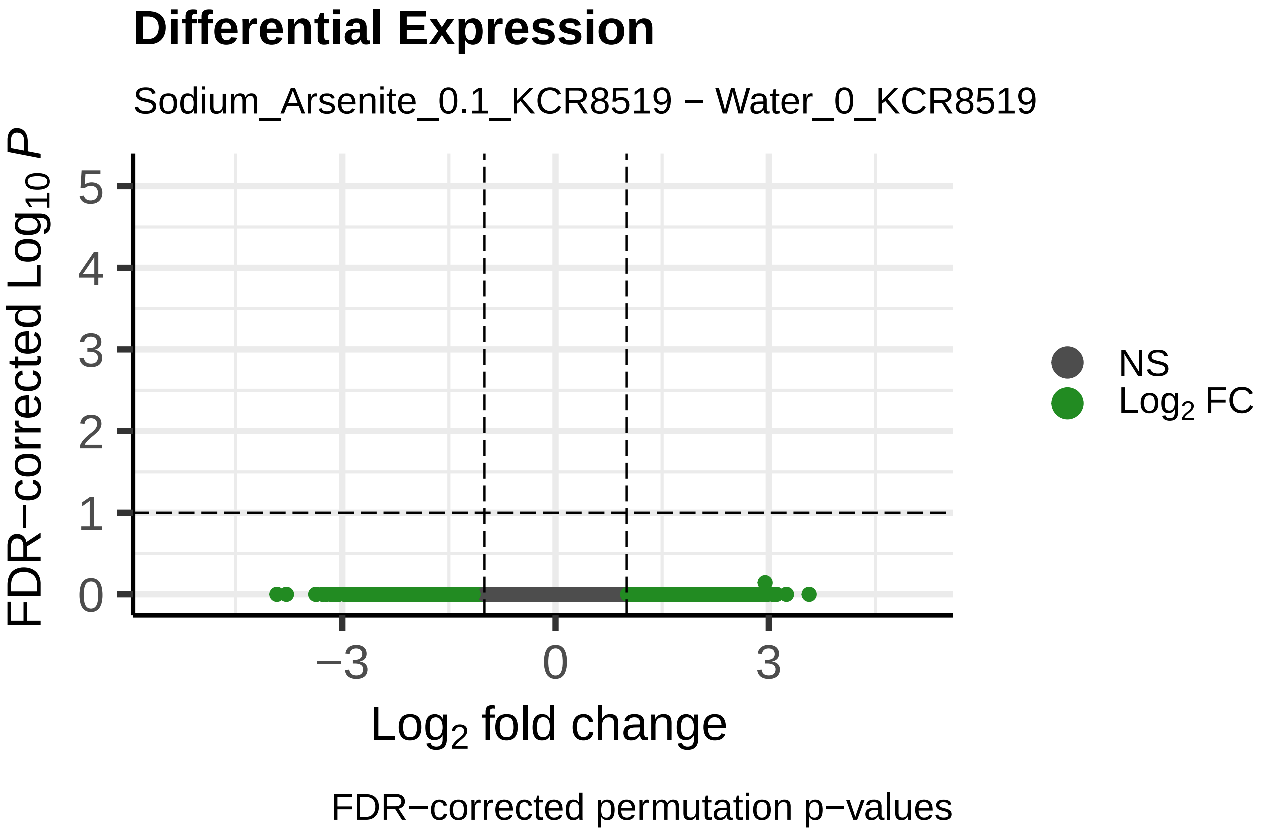
Differential Expression
Sodium_Arsenite_0.1_KCR8519 − Water_0_KCR8519
FDR−corrected Log10 P
Log2 fold change
NS
Log2 FC
FDR−corrected permutation p−values

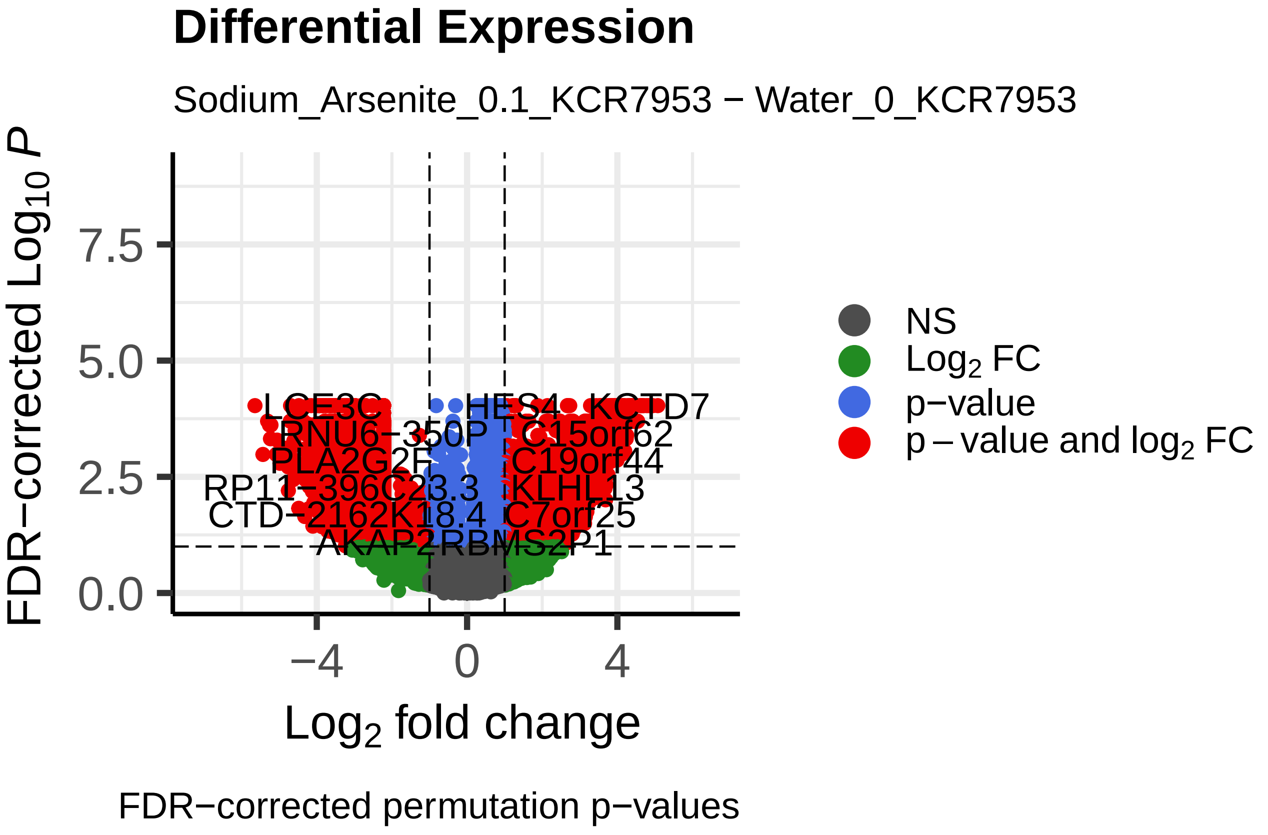
Differential Expression
Sodium_Arsenite_0.1_KCR7953 − Water_0_KCR7953
FDR−corrected Log10 P
Log2 fold change
LCE3C
HES4
KCTD7
C15orf62
C19orf44
KLHL13
C7orf25
AKAP2
RBMS2P1
NS
Log2 FC
p−value
p − value and log2 FC
FDR−corrected permutation p−values

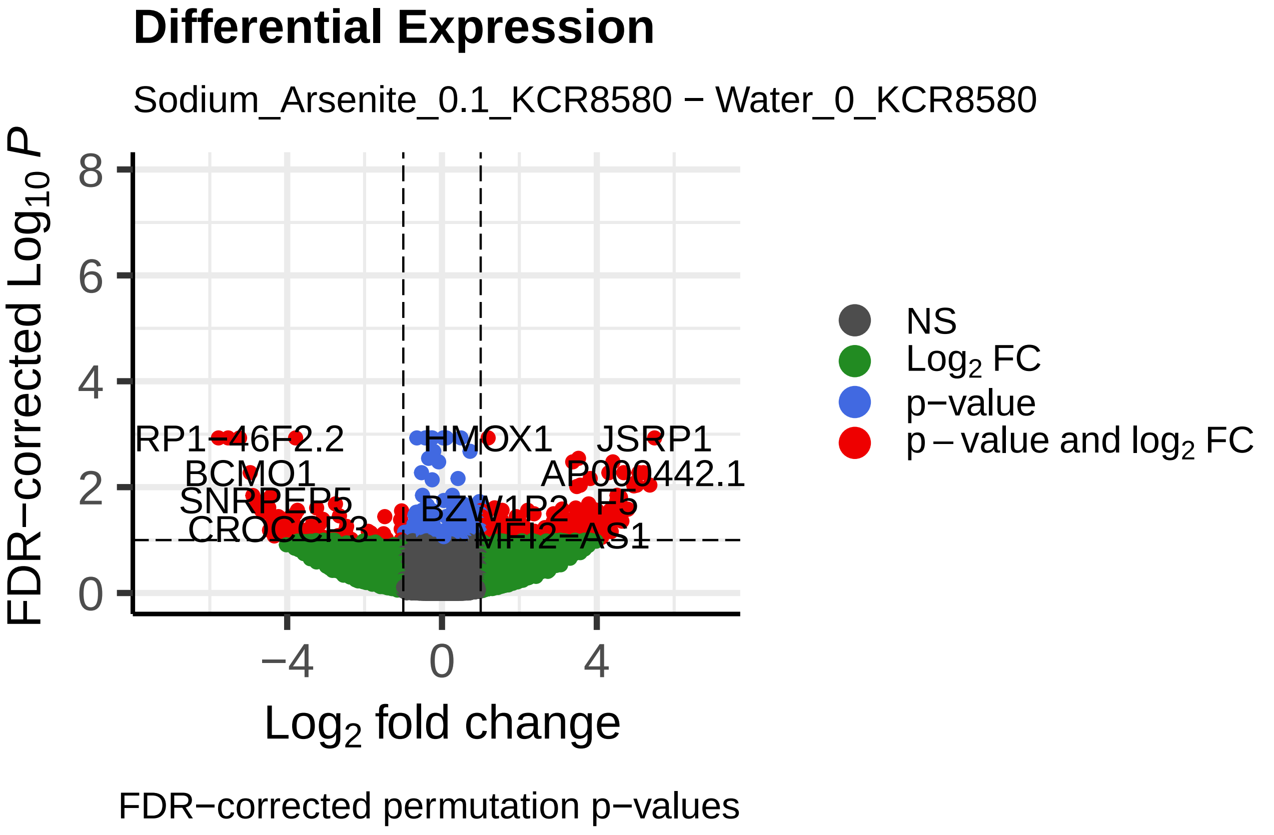
Differential Expression
Sodium_Arsenite_0.1_KCR8580 − Water_0_KCR8580
FDR−corrected Log10 P
Log2 fold change
RP1−46F2.2
HMOX1
JSRP1
BCMO1
AP000442.1
MFI2−AS1
NS
Log2 FC
p−value
p − value and log2 FC
FDR−corrected permutation p−values

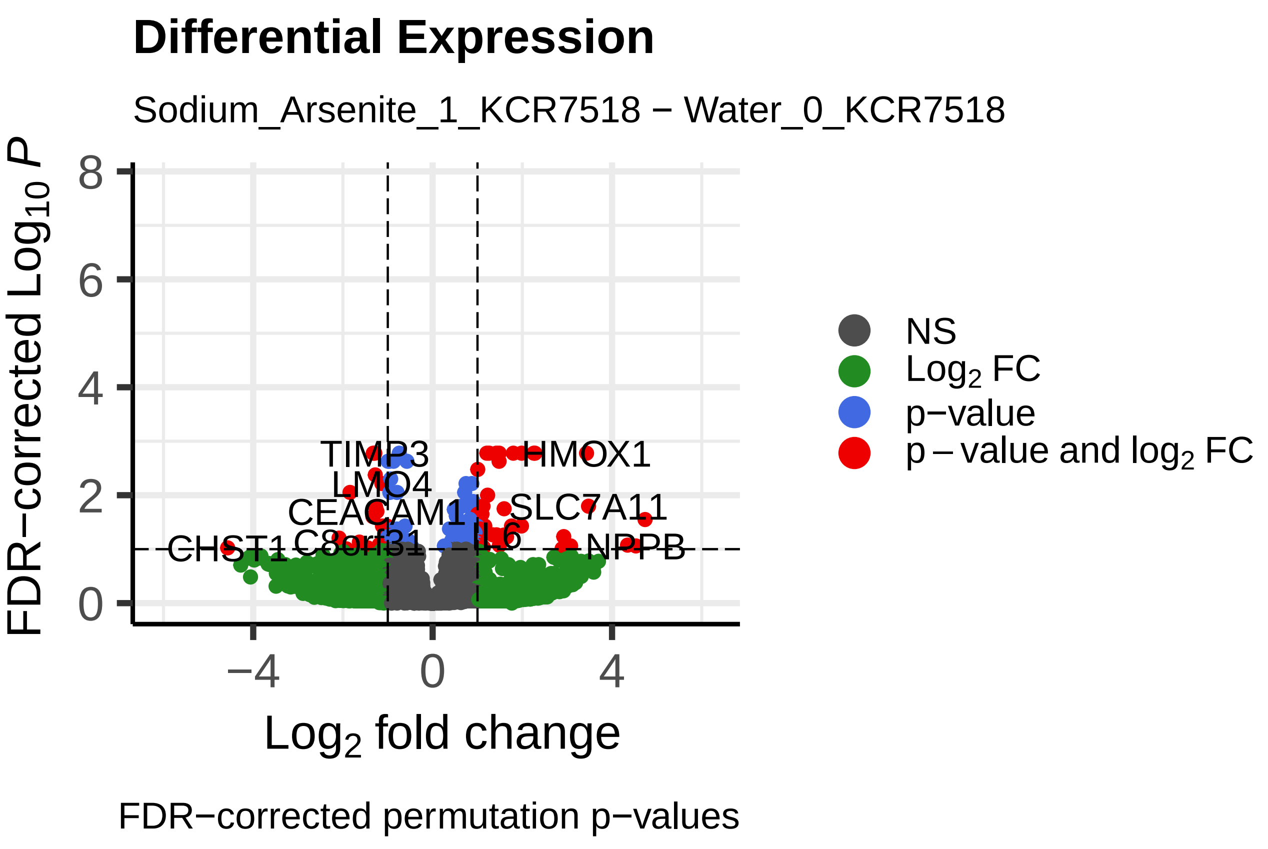
Differential Expression
Sodium_Arsenite_1_KCR7518 − Water_0_KCR7518
FDR−corrected $Log_{10}$ $P$
Log$_2$ fold change
TIMP3
LMO4
CEACAM1
CHST1
C8orf31
IL6
HMOX1
SLC7A11
NPPB
NS
Log$_2$ FC
p−value
p − value and log$_2$ FC
FDR−corrected permutation p−values

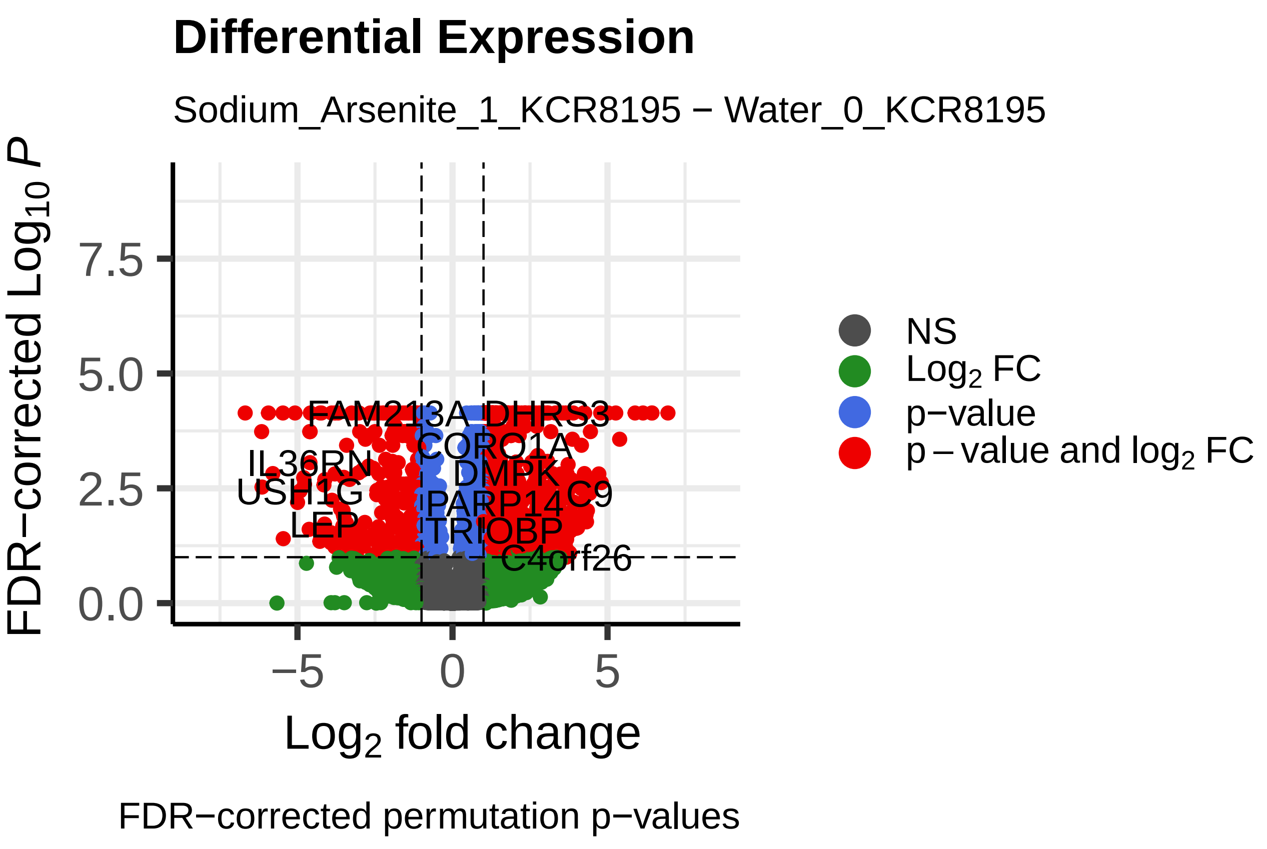
Differential Expression
Sodium_Arsenite_1_KCR8195 − Water_0_KCR8195
FDR−corrected $Log_{10}$ $P$
Log$_2$ fold change
FAM213A
DHRS3
CORO1A
IL36RN
DMPK
USH1G
PARP14
C9
LEP
TRIOBP
C4orf26
NS
Log$_2$ FC
p−value
p − value and log$_2$ FC
FDR−corrected permutation p−values

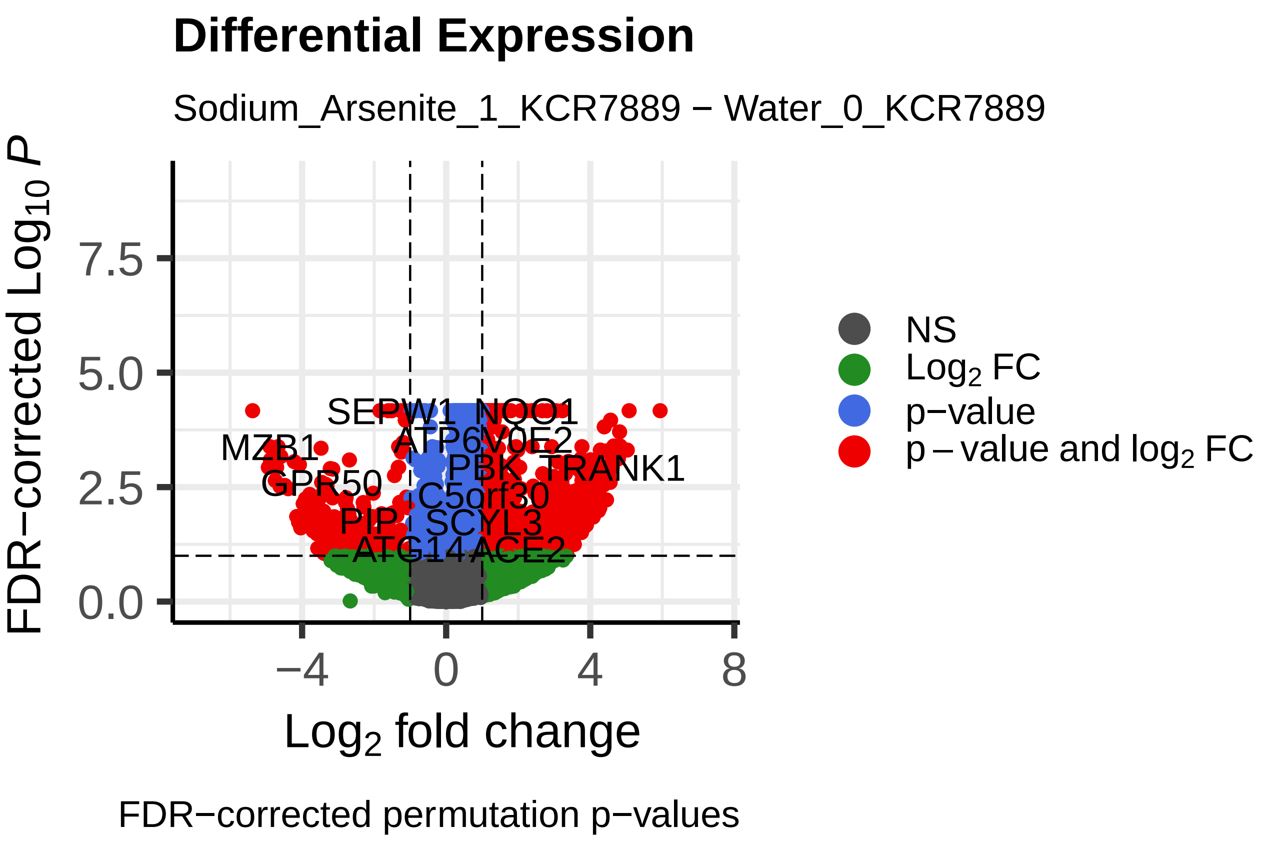
Differential Expression
Sodium_Arsenite_1_KCR7889 − Water_0_KCR7889
FDR−corrected $Log_{10}$ $P$
Log$_2$ fold change
SEPW1
NQO1
MZB1
ATP6V0E2
PBK
TRANK1
GPR50
C5orf30
PIP
SCYL3
ATG14
ACE2
NS
Log$_2$ FC
p−value
p − value and log$_2$ FC
FDR−corrected permutation p−values

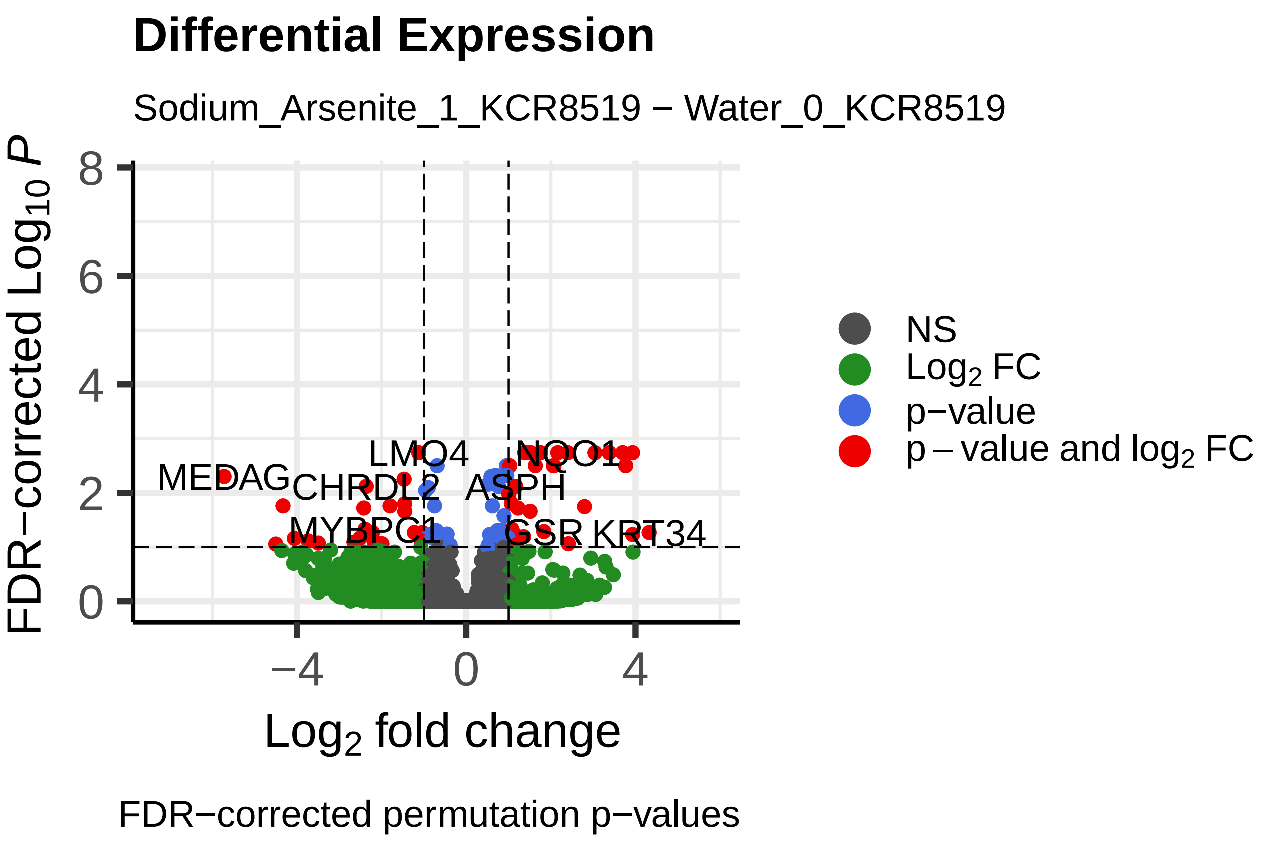
Differential Expression
Sodium_Arsenite_1_KCR8519 − Water_0_KCR8519
FDR−corrected $Log_{10}$ $P$
Log$_2$ fold change
LMO4
NQO1
MEDAG
CHRDL2
ASPH
MYBPC1
GSR
KRT34
NS
Log$_2$ FC
p−value
p − value and log$_2$ FC
FDR−corrected permutation p−values

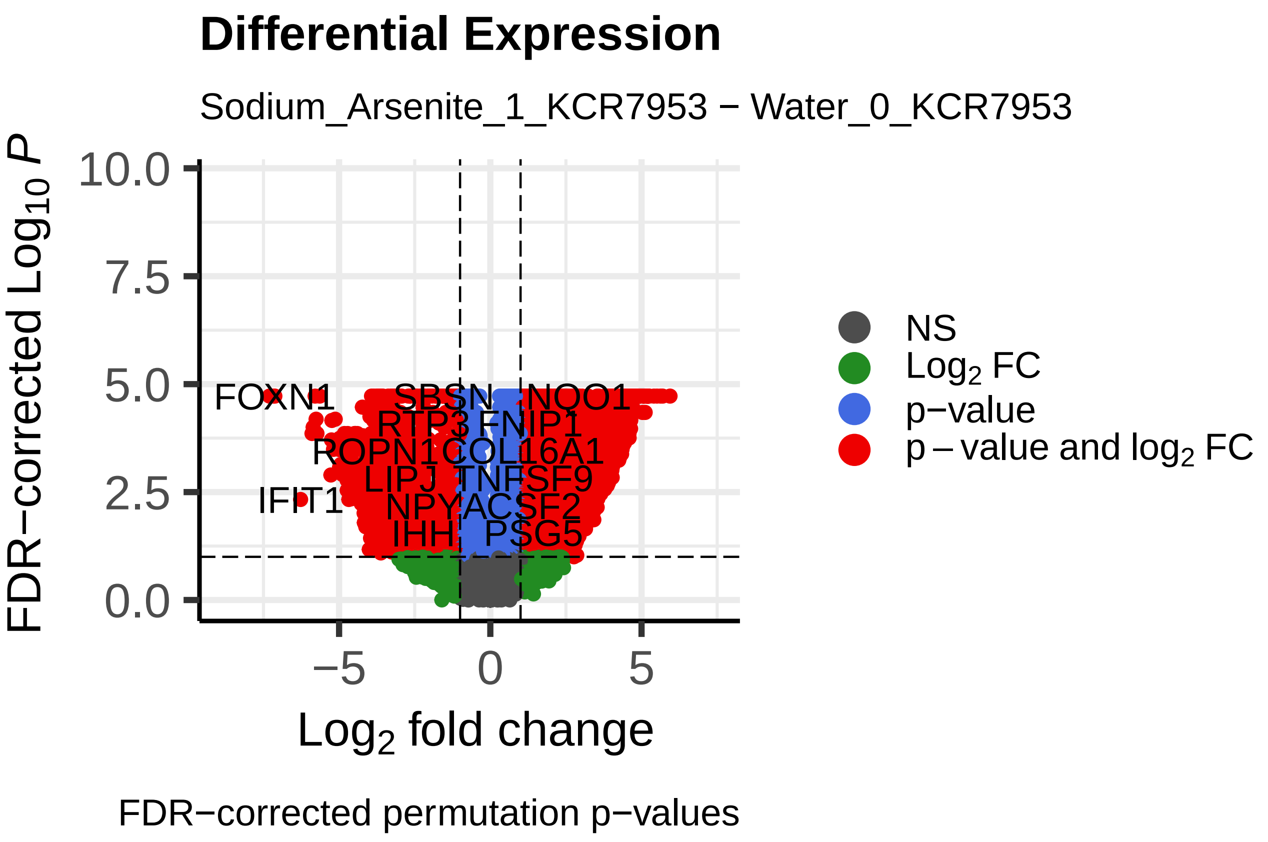
Differential Expression
Sodium_Arsenite_1_KCR7953 − Water_0_KCR7953
FDR−corrected $Log_{10}$ $P$
Log$_2$ fold change
FOXN1
SBSN
NQO1
RTP3
FNIP1
ROPN1
COL16A1
LIPJ
TNFSF9
IFIT1
NPY
ACSF2
IHH
PSG5
NS
Log$_2$ FC
p−value
p − value and log$_2$ FC
FDR−corrected permutation p−values

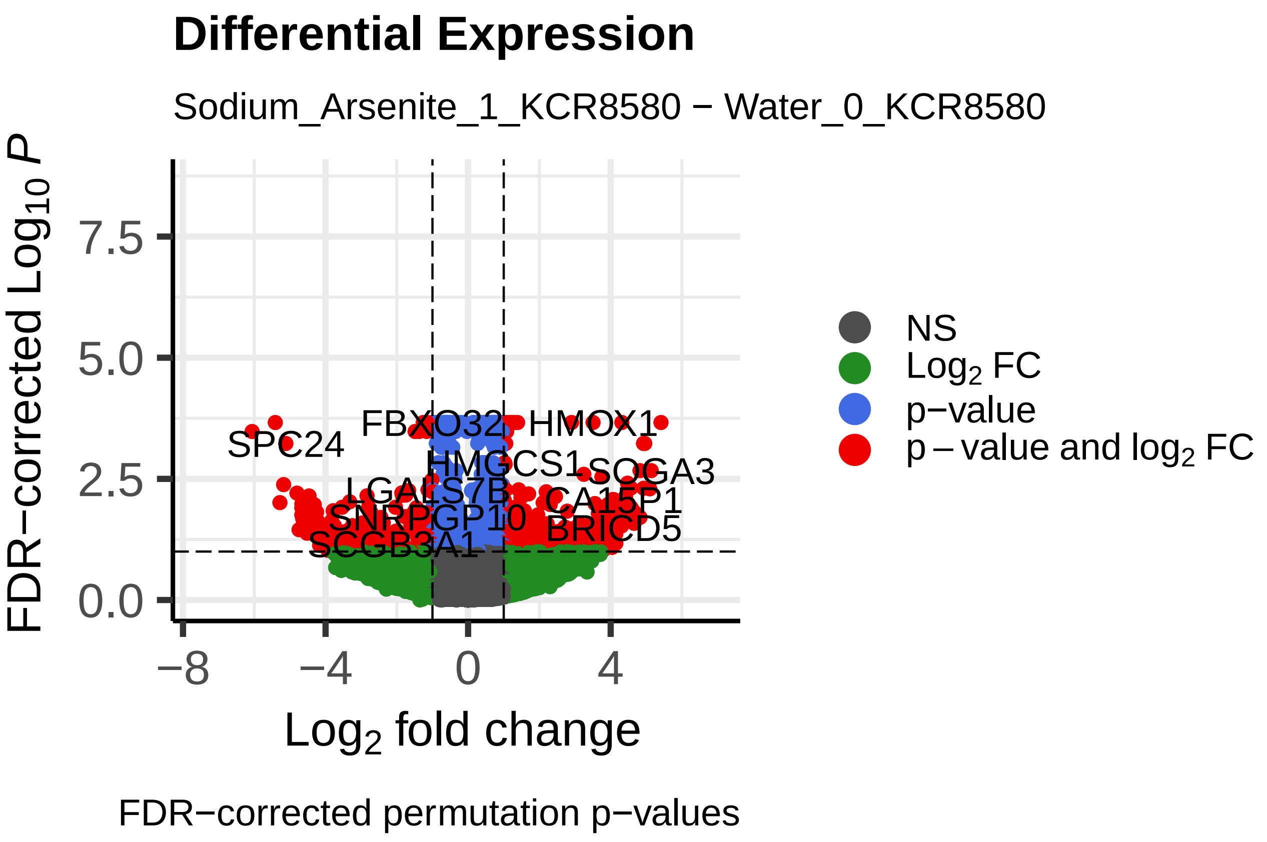
Differential Expression
Sodium_Arsenite_1_KCR8580 − Water_0_KCR8580
FDR−corrected $Log_{10}$ $P$
Log$_2$ fold change
FBXO32
HMOX1
SPC24
HMGCS1
SOGA3
LGALS7B
CA15P1
SNRPGP10
BRICD5
SCGB3A1
NS
Log$_2$ FC
p−value
p − value and log$_2$ FC
FDR−corrected permutation p−values

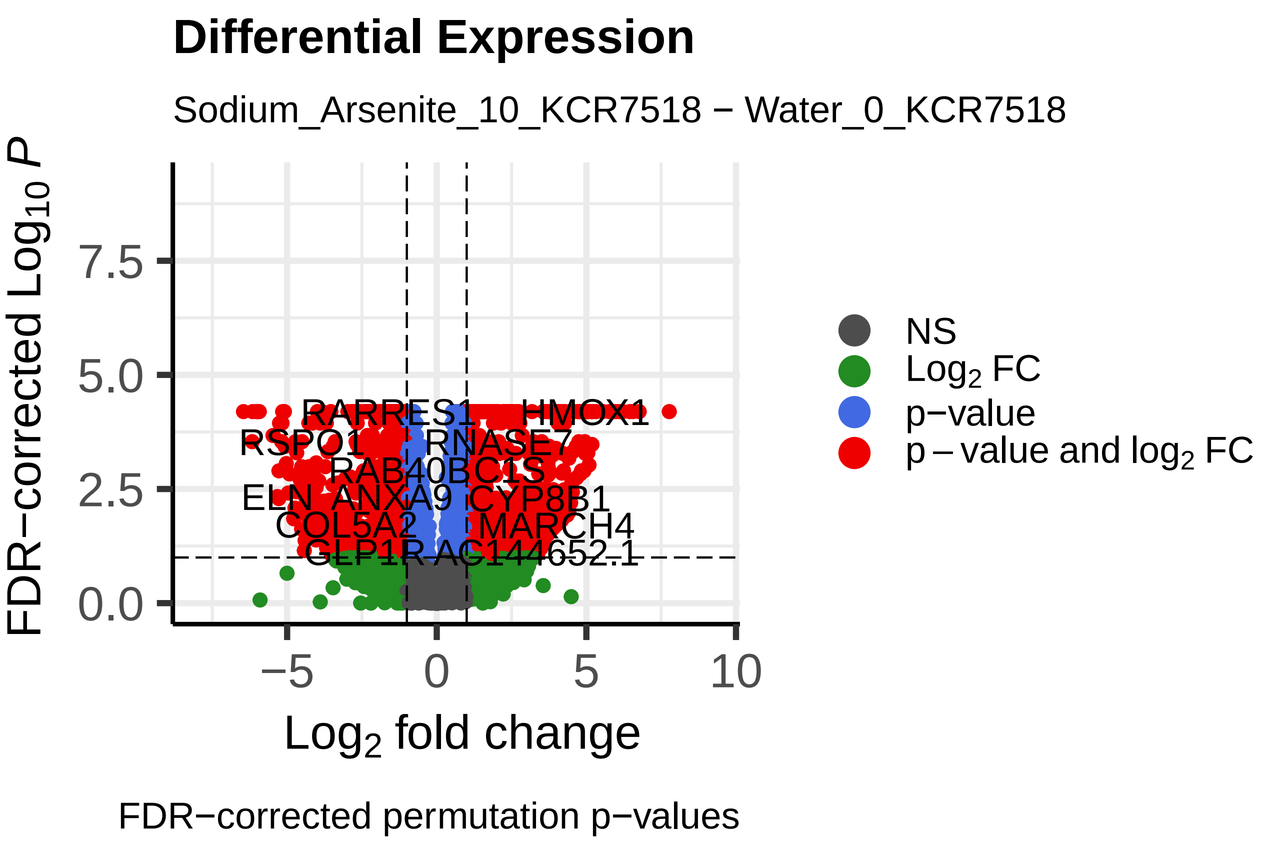
Differential Expression
Sodium_Arsenite_10_KCR7518 − Water_0_KCR7518
FDR−corrected Log10 P
Log2 fold change
NS
Log2 FC
p−value
p − value and log2 FC
RARRES1
HMOX1
RSPO1
RNASE7
RAB40B
C1S
ELN
ANXA9
CYP8B1
COL5A2
MARCH4
GLP1R
AC144652.1
FDR−corrected permutation p−values

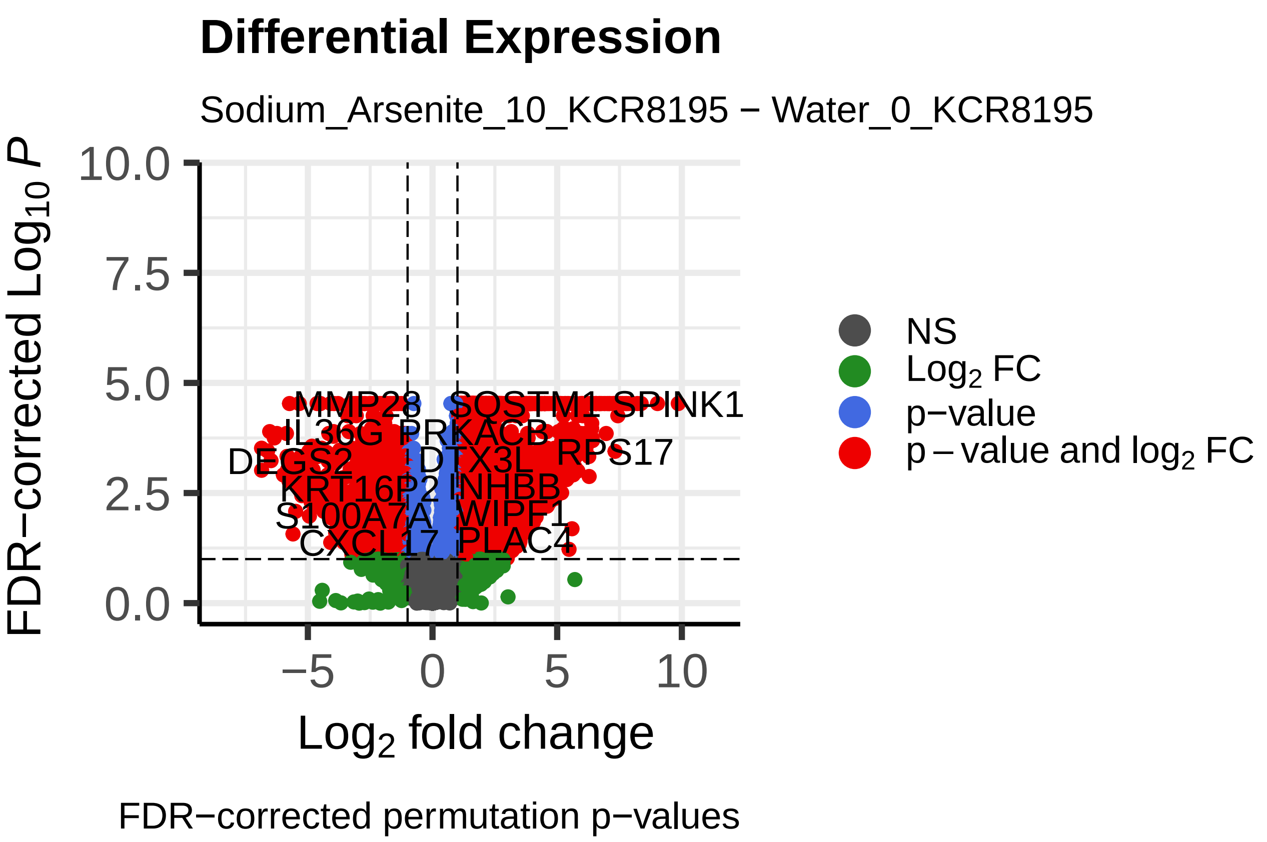
Differential Expression
Sodium_Arsenite_10_KCR8195 − Water_0_KCR8195
FDR−corrected Log10 P
Log2 fold change
NS
Log2 FC
p−value
p − value and log2 FC
MMP28
SQSTM1
SPINK1
IL36G
PRKACB
DEGS2
DTX3L
RPS17
KRT16P2
INHBB
S100A7A
WIPF1
CXCL17
PLAC4
FDR−corrected permutation p−values

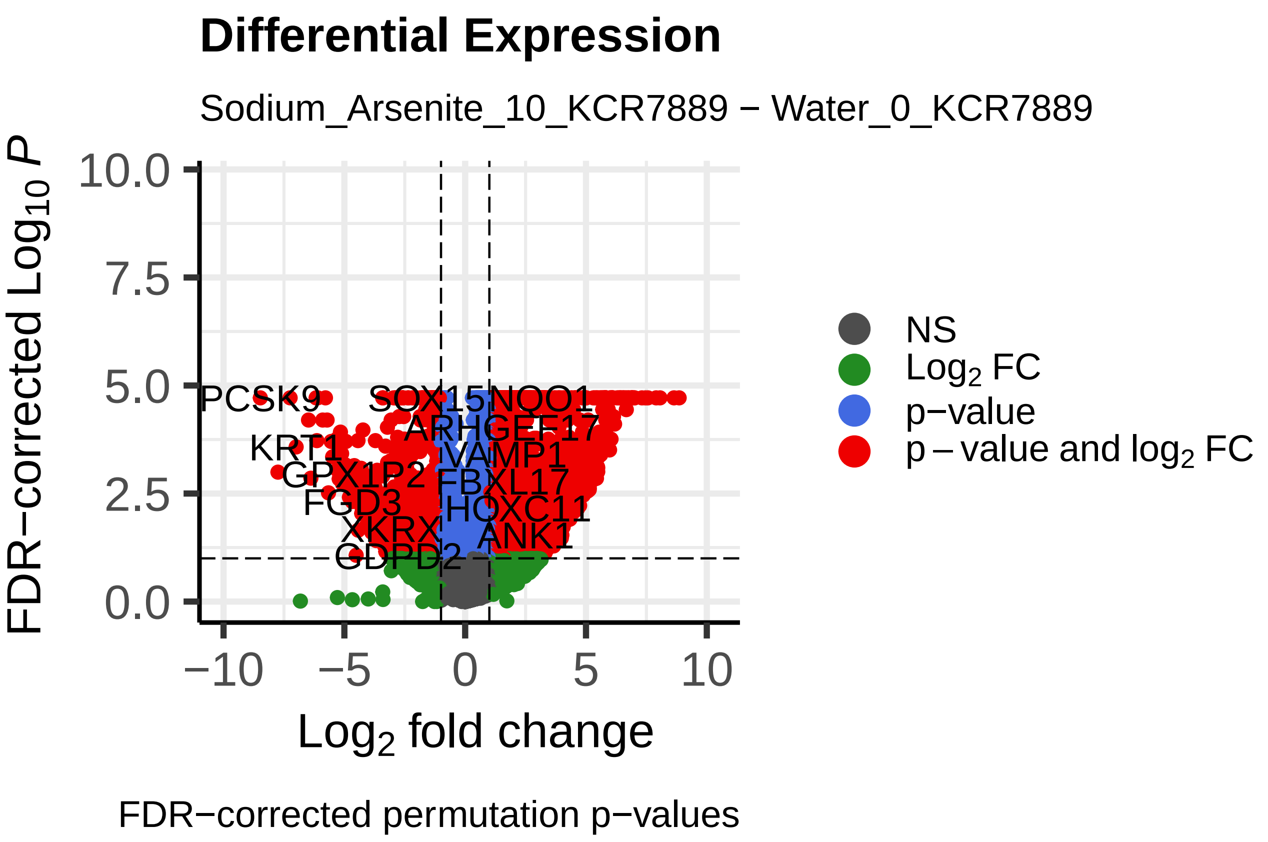
Differential Expression
Sodium_Arsenite_10_KCR7889 − Water_0_KCR7889
FDR−corrected Log10 P
Log2 fold change
NS
Log2 FC
p−value
p − value and log2 FC
PCSK9
SOX15
NQO1
ARHGEF17
KRT1
VAMP1
GPX1P2
FBXL17
FGD3
HOXC11
XKRX
ANK1
GDPD2
FDR−corrected permutation p−values

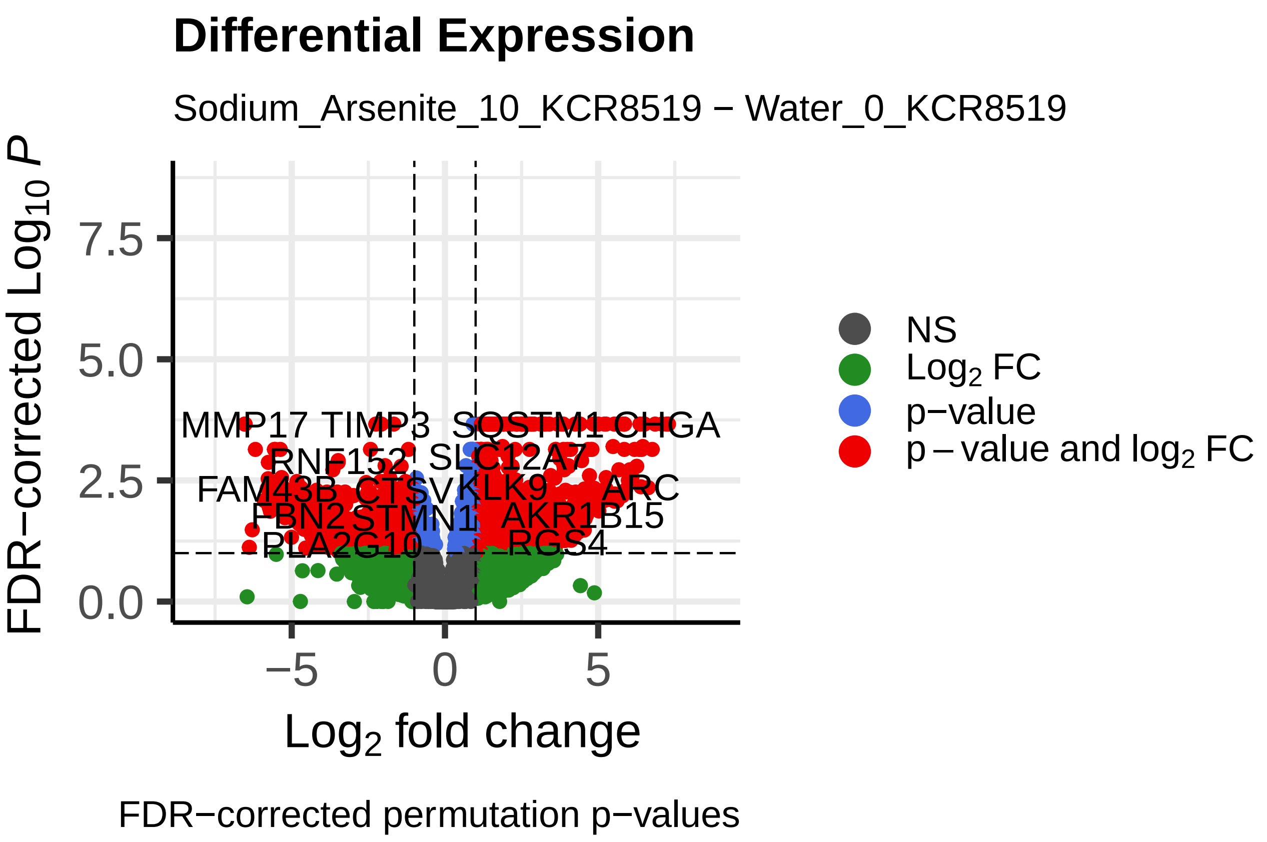
Differential Expression
Sodium_Arsenite_10_KCR8519 − Water_0_KCR8519
FDR−corrected Log10 P
Log2 fold change
NS
Log2 FC
p−value
p − value and log2 FC
MMP17
TIMP3
SQSTM1
CHGA
RNF152
SLC12A7
FAM43B
CTSV
KLK9
ARC
FBN2
STMN1
AKR1B15
PLA2G10
RGS4
FDR−corrected permutation p−values

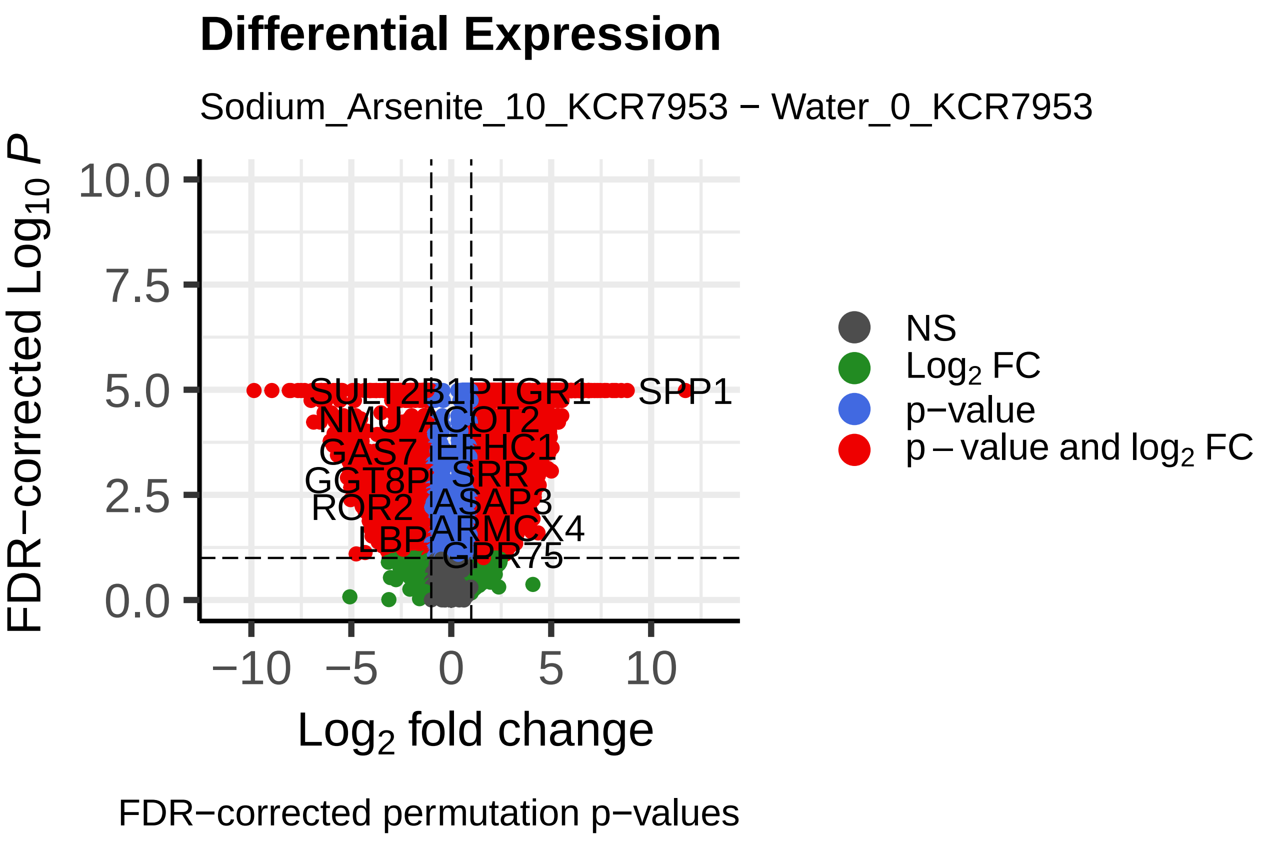
Differential Expression
Sodium_Arsenite_10_KCR7953 − Water_0_KCR7953
FDR−corrected Log10 P
Log2 fold change
NS
Log2 FC
p−value
p − value and log2 FC
SULT2B1
PTGR1
SPP1
NMU
ACOT2
GAS7
EFHC1
GGT8P
SRR
ROR2
ASAP3
BPI
ARMCX4
GPR75
FDR−corrected permutation p−values

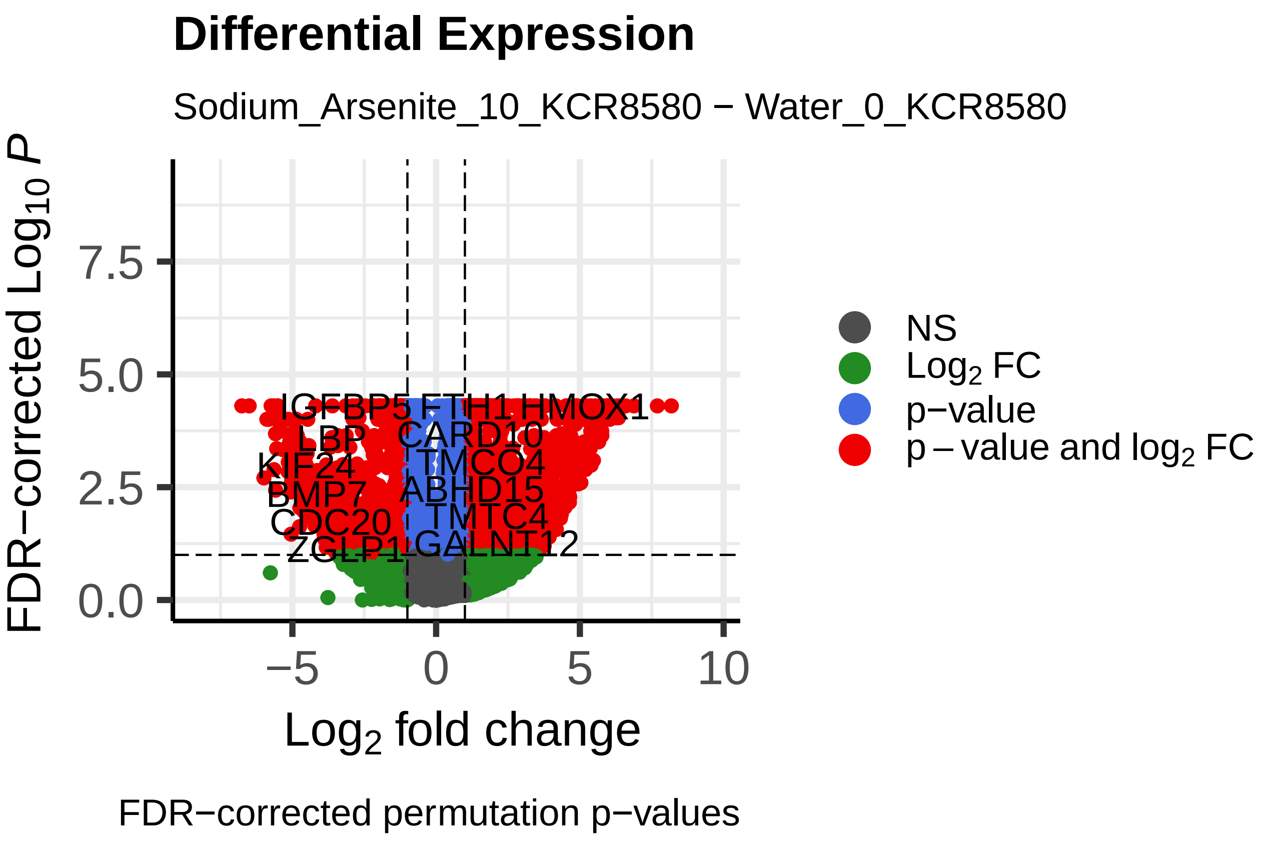
Differential Expression
Sodium_Arsenite_10_KCR8580 − Water_0_KCR8580
FDR−corrected Log10 P
Log2 fold change
NS
Log2 FC
p−value
p − value and log2 FC
IGFBP5
FTH1
HMOX1
LBP
CARD10
KIF24
TMCO4
BMP7
ABHD15
CDC20
TMTC4
ZGLP1
GALNT12
FDR−corrected permutation p−values

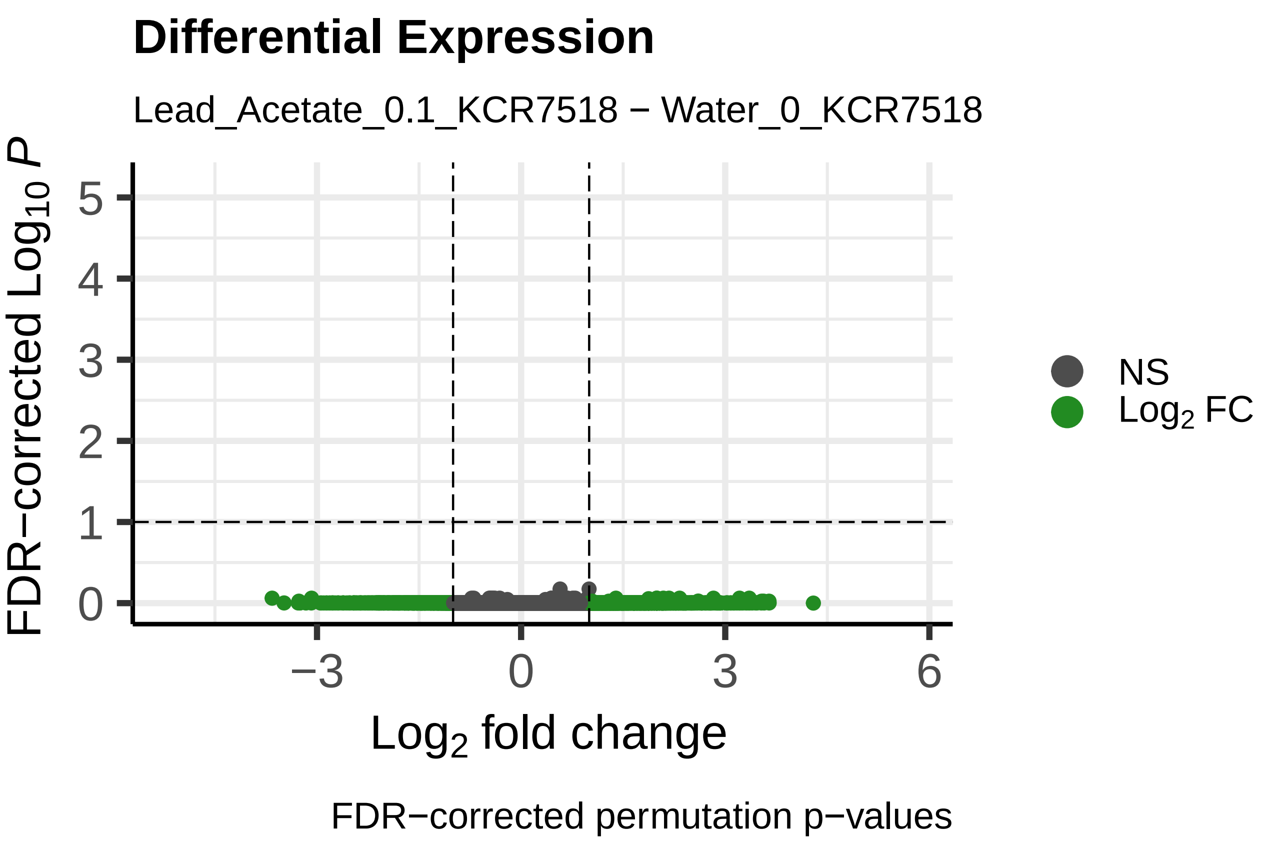
Differential Expression
Lead_Acetate_0.1_KCR7518 − Water_0_KCR7518
FDR−corrected Log$_{10}$ $P$
Log$_2$ fold change
NS
Log$_2$ FC
FDR−corrected permutation p−values

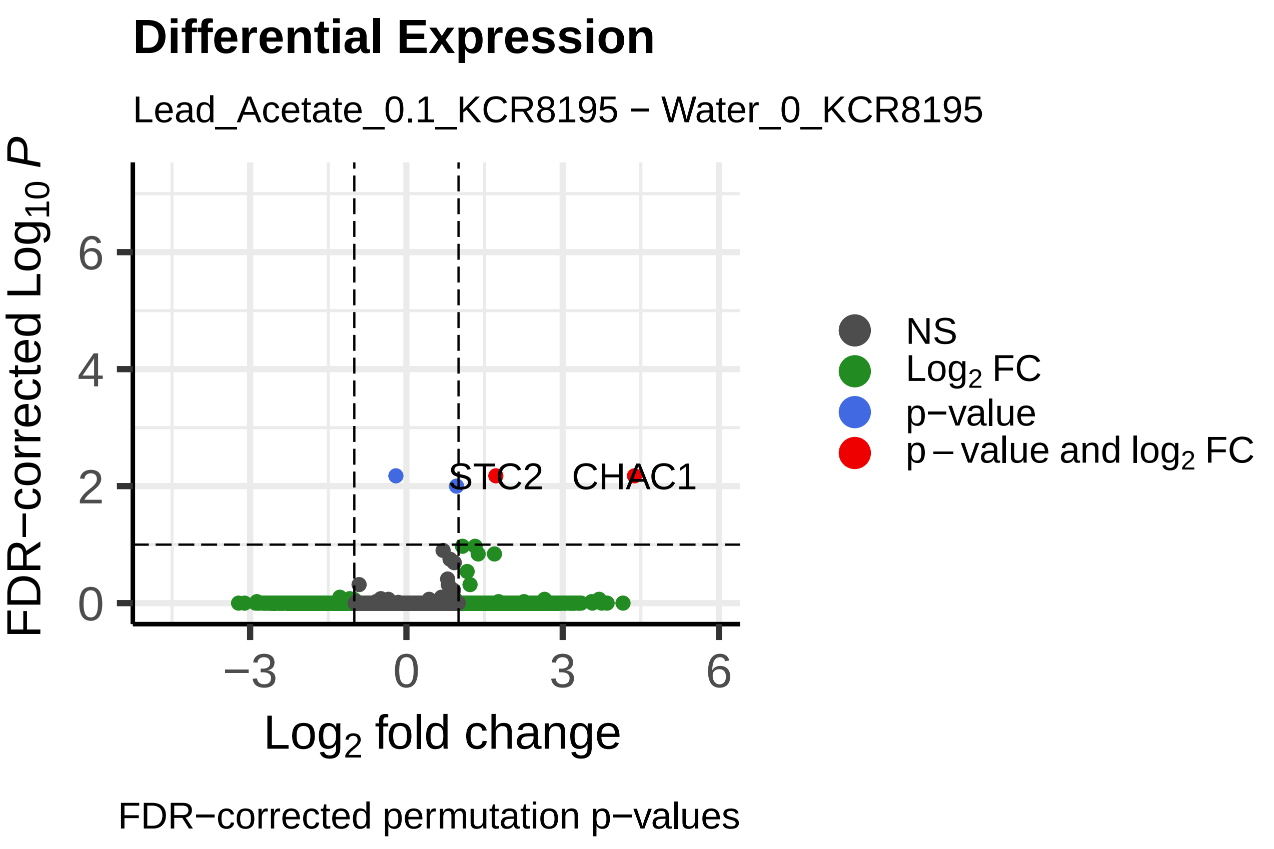
Differential Expression
Lead_Acetate_0.1_KCR8195 − Water_0_KCR8195
FDR−corrected Log$_{10}$ $P$
Log$_2$ fold change
STC2
CHAC1
NS
Log$_2$ FC
p−value
p − value and log$_2$ FC
FDR−corrected permutation p−values

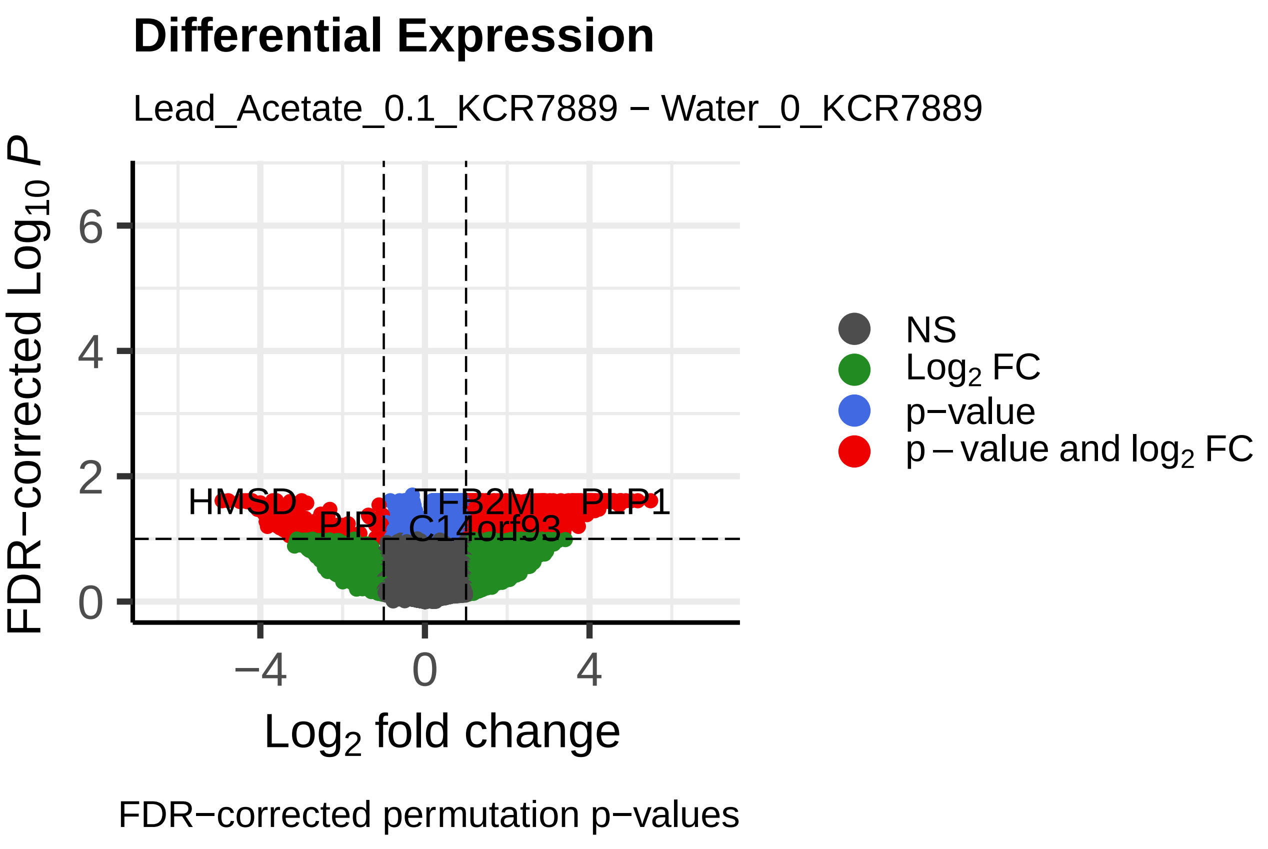
Differential Expression
Lead_Acetate_0.1_KCR7889 − Water_0_KCR7889
FDR−corrected Log$_{10}$ $P$
Log$_2$ fold change
HMSD
TFB2M
PLP1
PIP
C14orf93
NS
Log$_2$ FC
p−value
p − value and log$_2$ FC
FDR−corrected permutation p−values

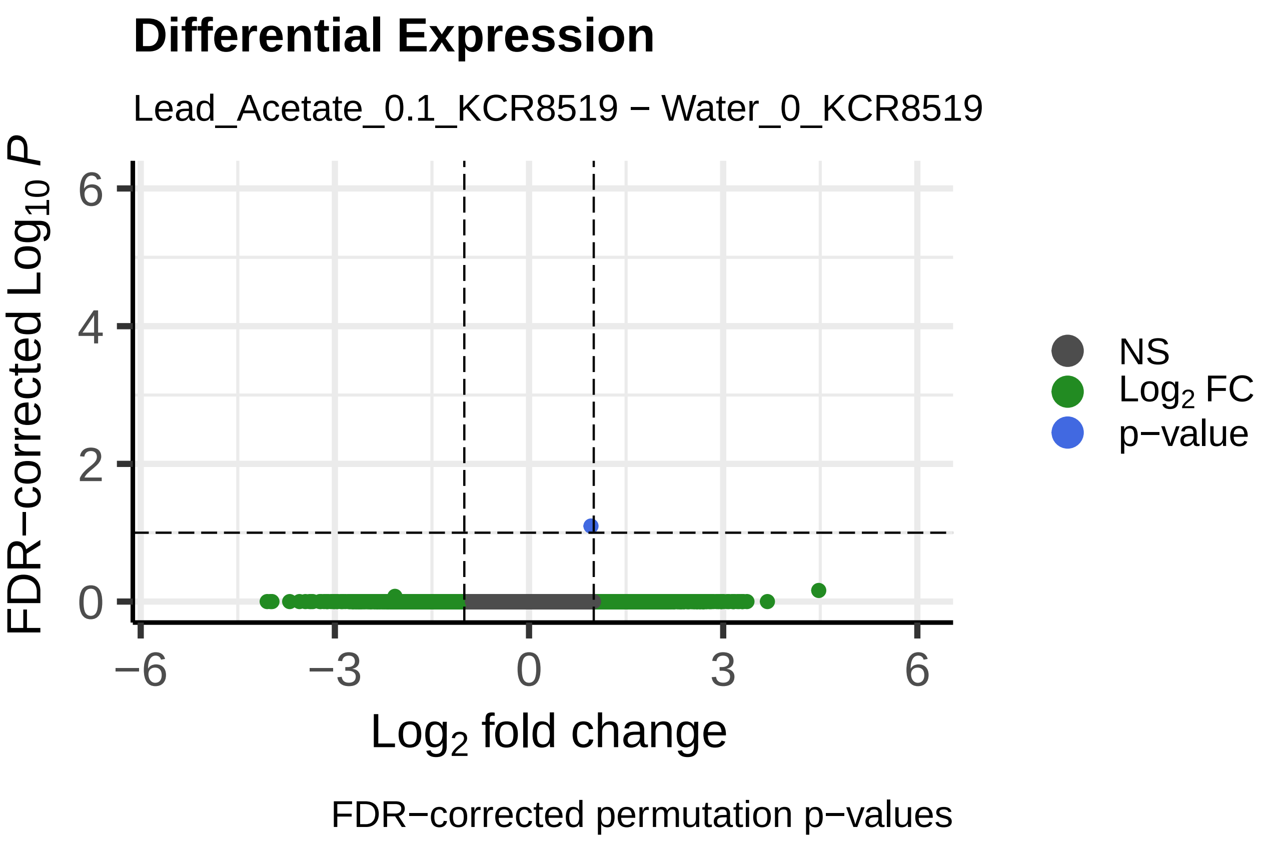
Differential Expression
Lead_Acetate_0.1_KCR8519 − Water_0_KCR8519
FDR−corrected Log$_{10}$ $P$
Log$_2$ fold change
NS
Log$_2$ FC
p−value
FDR−corrected permutation p−values

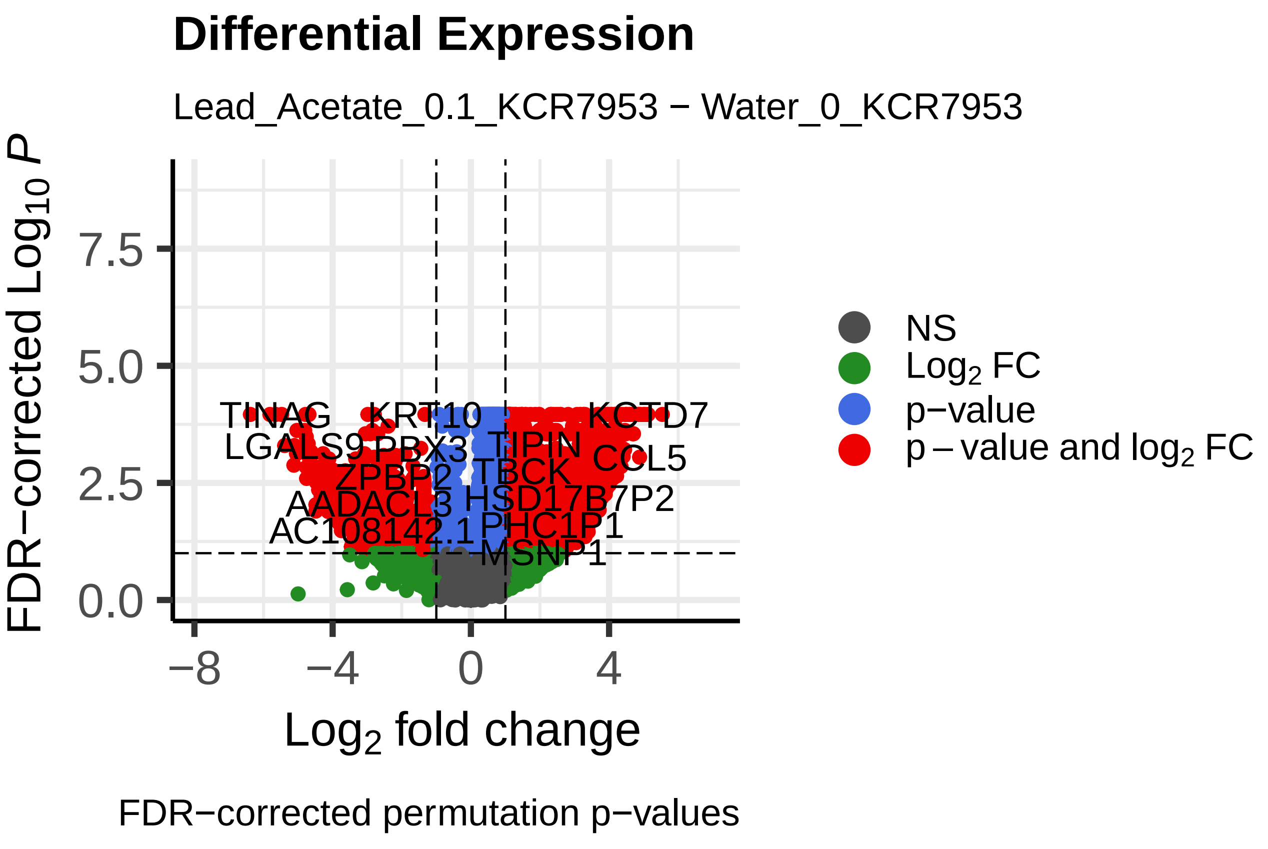
Differential Expression
Lead_Acetate_0.1_KCR7953 − Water_0_KCR7953
FDR−corrected Log$_{10}$ $P$
Log$_2$ fold change
TINAG
KRT10
KCTD7
TIPIN
COL5
TBCK
AADACL3
HSD17B7P2
PHC1P1
MSNP1
NS
Log$_2$ FC
p−value
p − value and log$_2$ FC
FDR−corrected permutation p−values

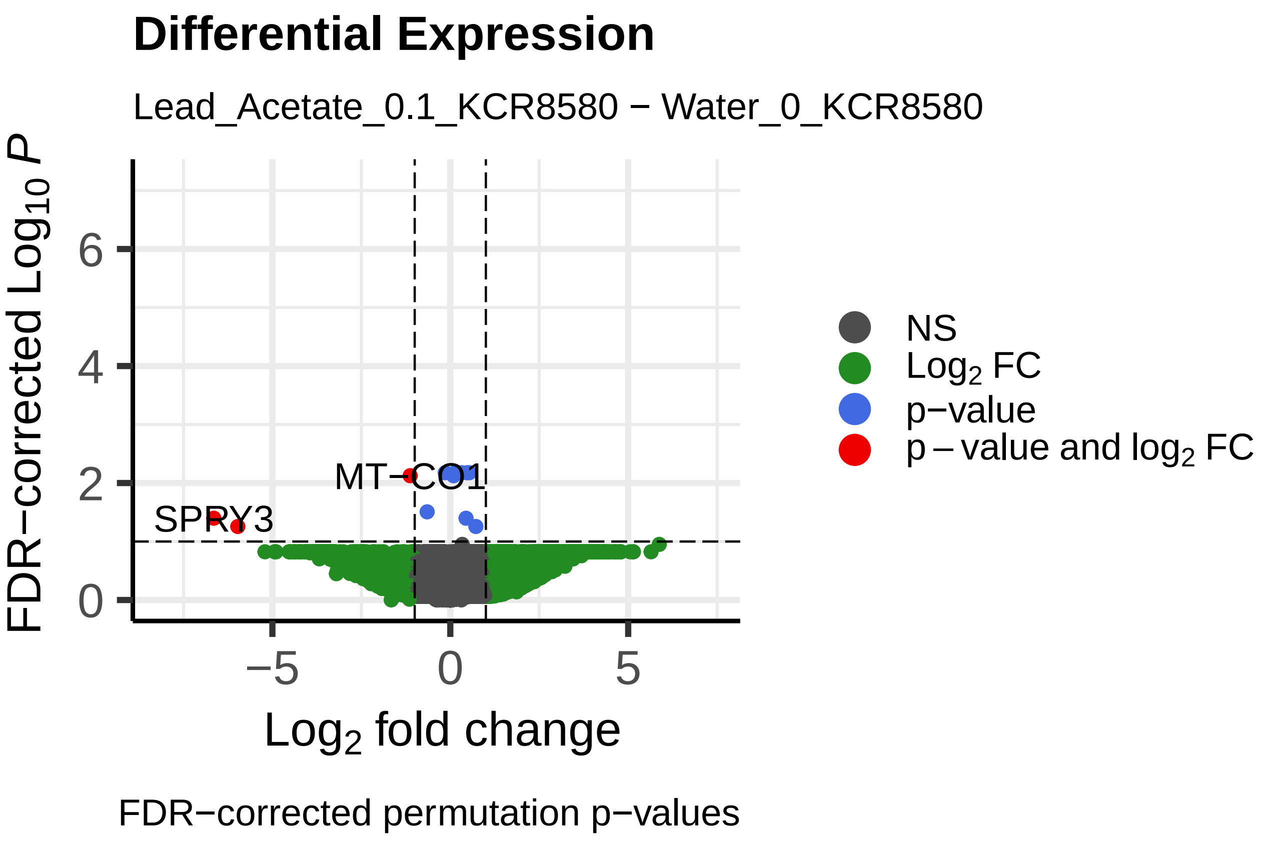
Differential Expression
Lead_Acetate_0.1_KCR8580 − Water_0_KCR8580
FDR−corrected Log$_{10}$ $P$
Log$_2$ fold change
MT−CO1
SPRY3
NS
Log$_2$ FC
p−value
p − value and log$_2$ FC
FDR−corrected permutation p−values

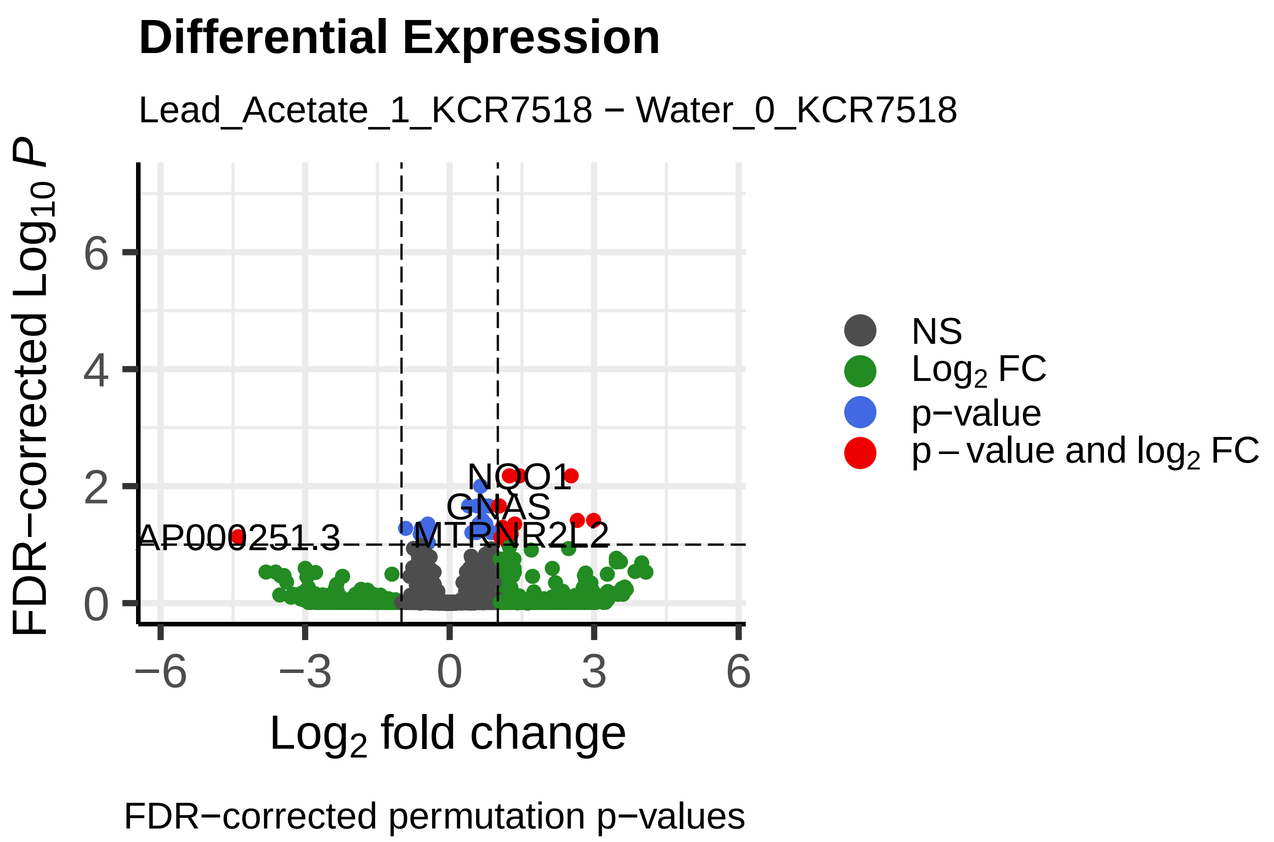
Differential Expression
Lead_Acetate_1_KCR7518 − Water_0_KCR7518
FDR−corrected Log10 P
Log2 fold change
NS
Log2 FC
p−value
p − value and log2 FC
NQO1
GNAS
AP000251.3
MTRNR2L2
FDR−corrected permutation p−values

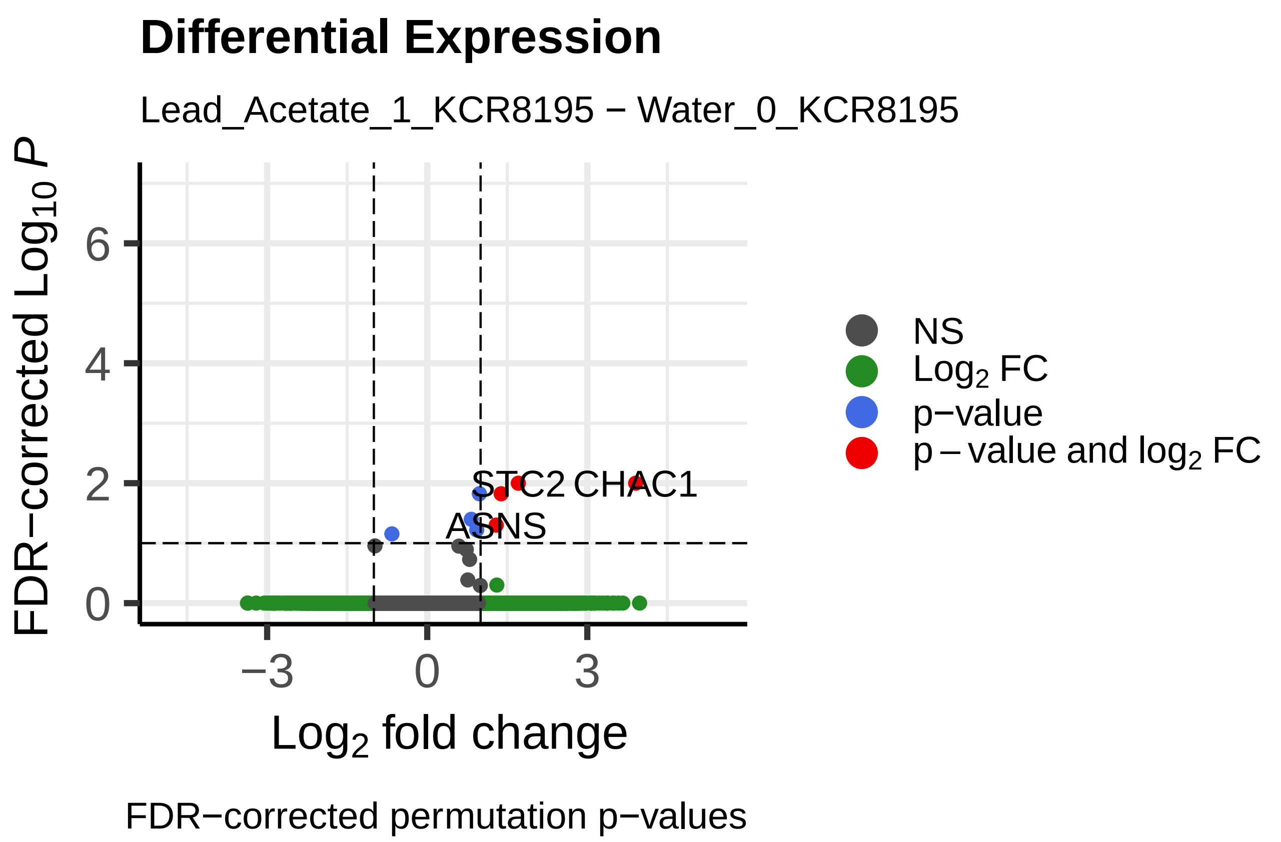
Differential Expression
Lead_Acetate_1_KCR8195 − Water_0_KCR8195
FDR−corrected Log10 P
Log2 fold change
NS
Log2 FC
p−value
p − value and log2 FC
STC2
CHAC1
ASNS
FDR−corrected permutation p−values

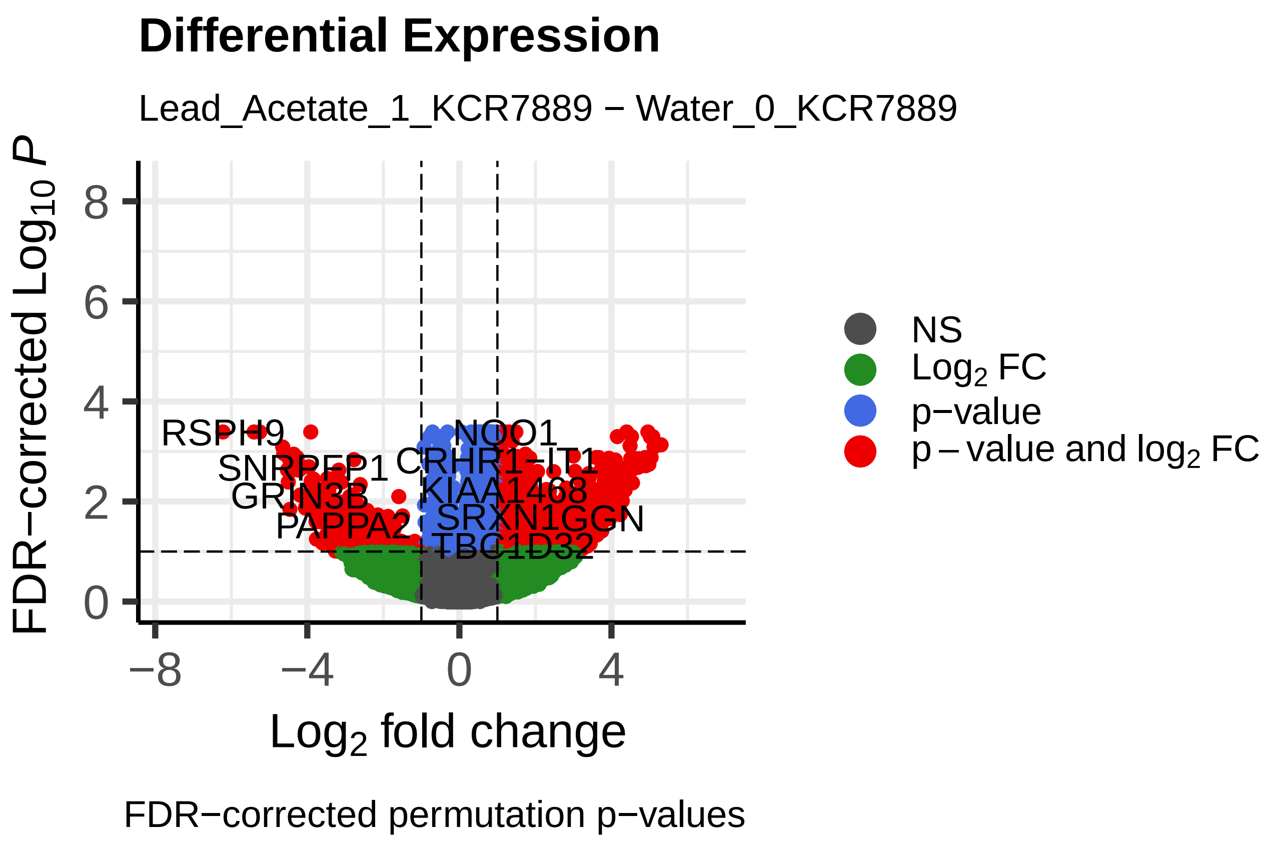
Differential Expression
Lead_Acetate_1_KCR7889 − Water_0_KCR7889
FDR−corrected Log10 P
Log2 fold change
NS
Log2 FC
p−value
p − value and log2 FC
RSPH9
NQO1
SNRPEP1
CRHR1−IT1
GRIN3B
KIAA1468
PAPPA2
SRXN1
GGN
TBC1D32
FDR−corrected permutation p−values

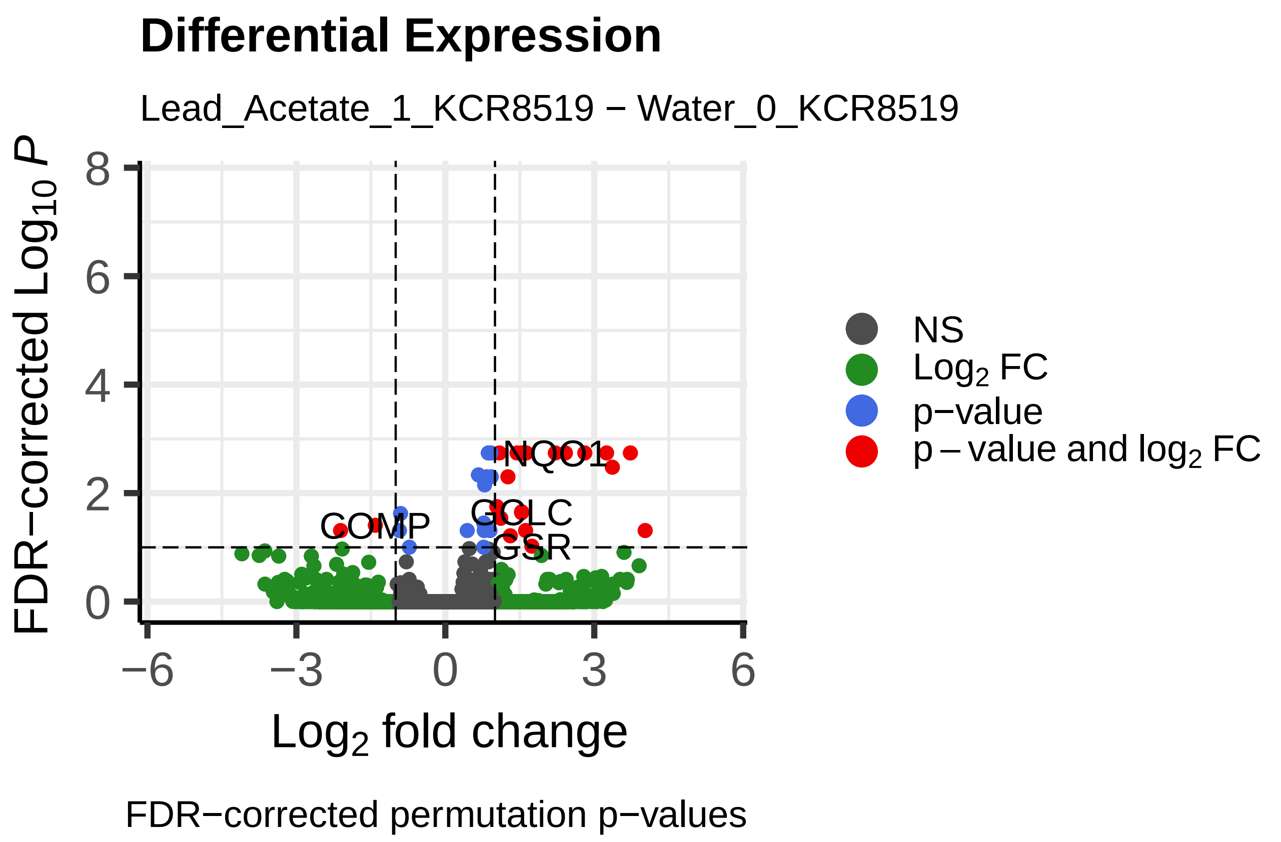
Differential Expression
Lead_Acetate_1_KCR8519 − Water_0_KCR8519
FDR−corrected Log10 P
Log2 fold change
NS
Log2 FC
p−value
p − value and log2 FC
NQO1
COMP
GCLC
GSR
FDR−corrected permutation p−values

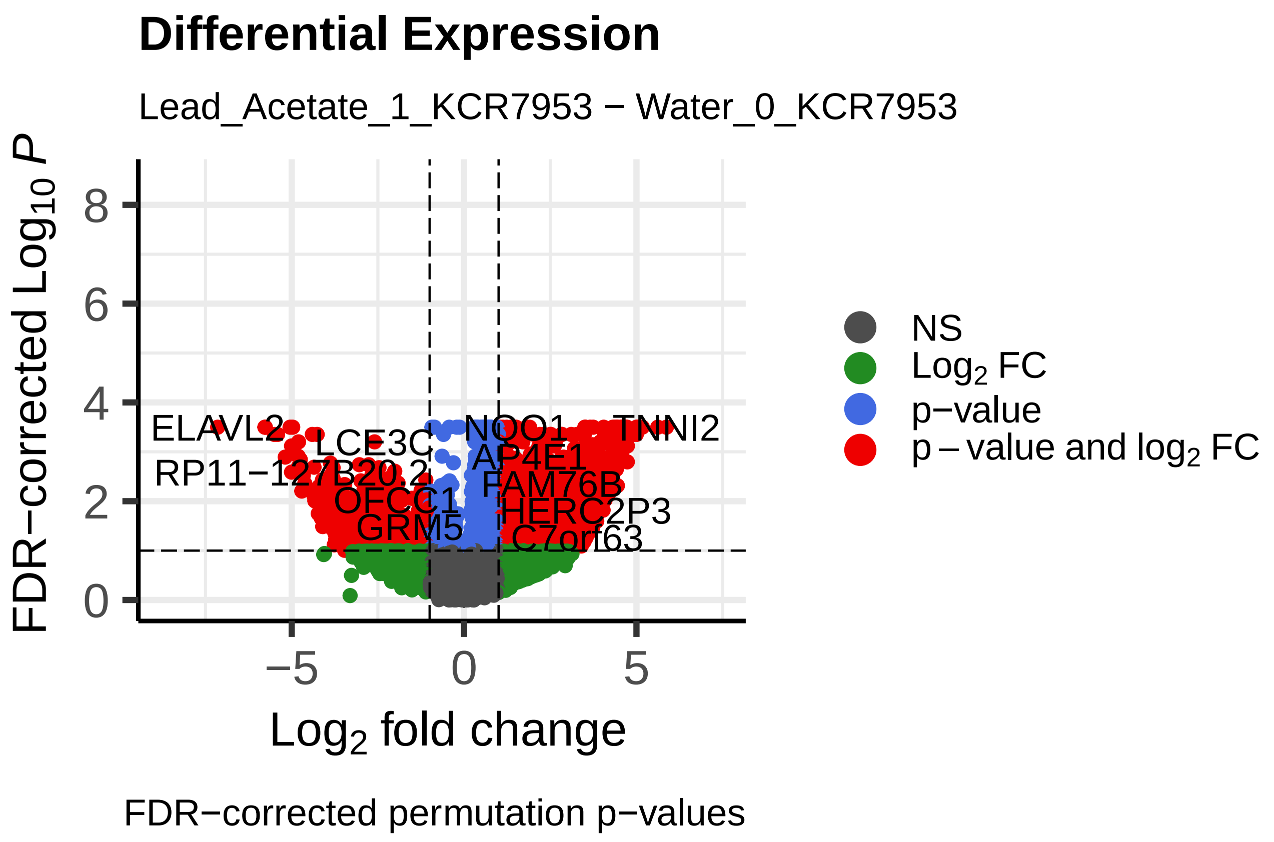
Differential Expression
Lead_Acetate_1_KCR7953 − Water_0_KCR7953
FDR−corrected Log10 P
Log2 fold change
NS
Log2 FC
p−value
p − value and log2 FC
ELAVL2
NQO1
TNNI2
LCE3C
AP4E1
RP11−127B20.2
FAM76B
OFCC1
HERC2P3
GRM5
C7orf63
FDR−corrected permutation p−values

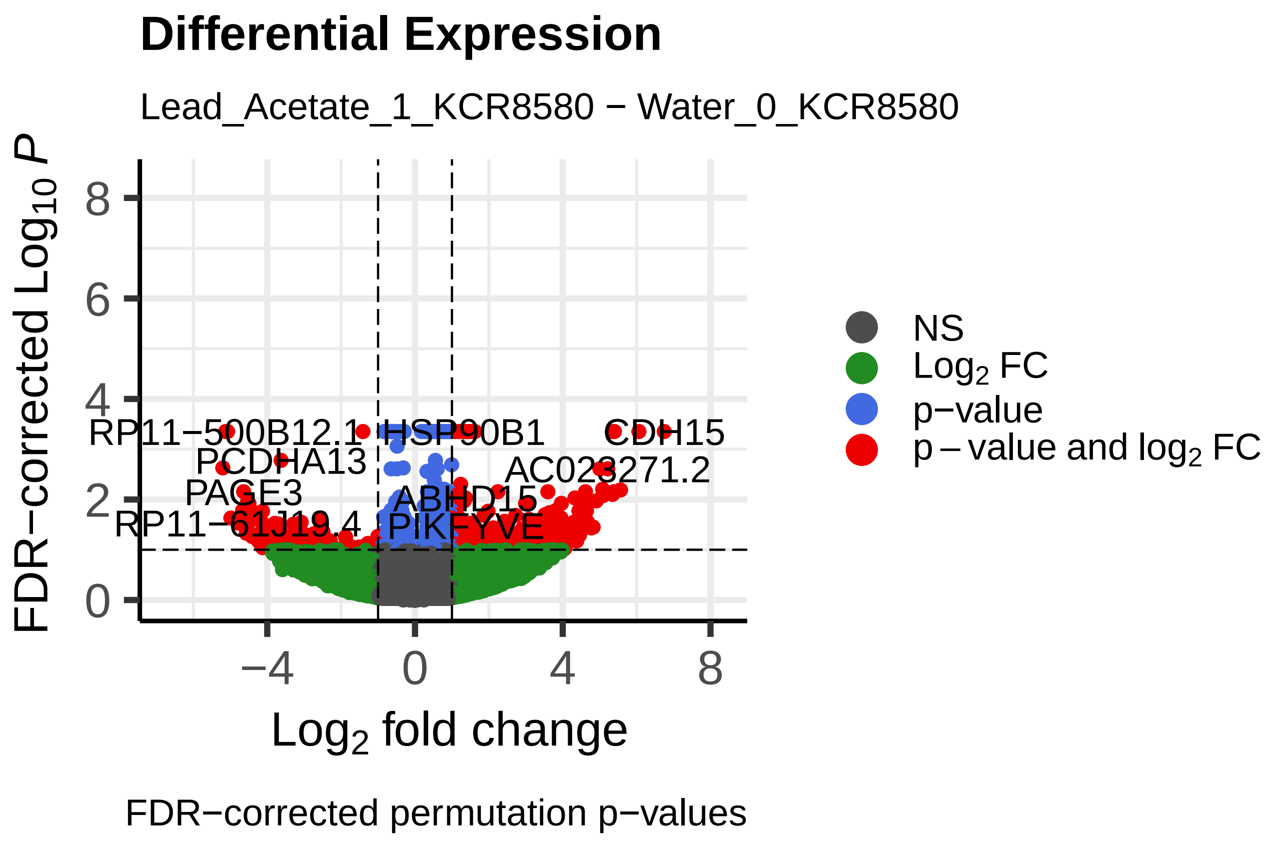
Differential Expression
Lead_Acetate_1_KCR8580 − Water_0_KCR8580
FDR−corrected Log10 P
Log2 fold change
NS
Log2 FC
p−value
p − value and log2 FC
RP11−500B12.1
HSP90B1
CDH15
PCDHA13
AC023271.2
PAGE3
ABHD15
RP11−61J19.4
PIKFYVE
FDR−corrected permutation p−values

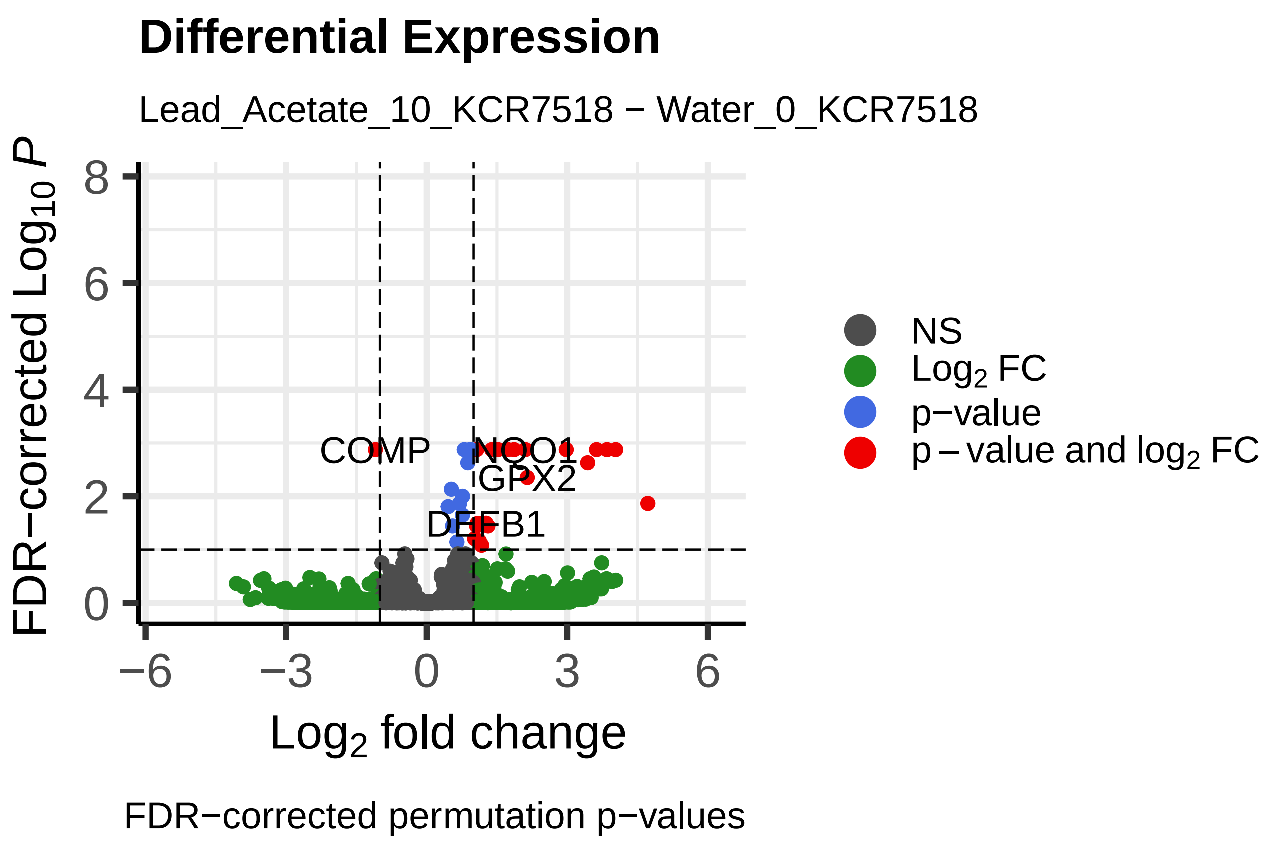
Differential Expression
Lead_Acetate_10_KCR7518 − Water_0_KCR7518
FDR−corrected Log10 P
Log2 fold change
NS
Log2 FC
p−value
p − value and log2 FC
COMP
NQO1
IGFX2
DEFB1
FDR−corrected permutation p−values

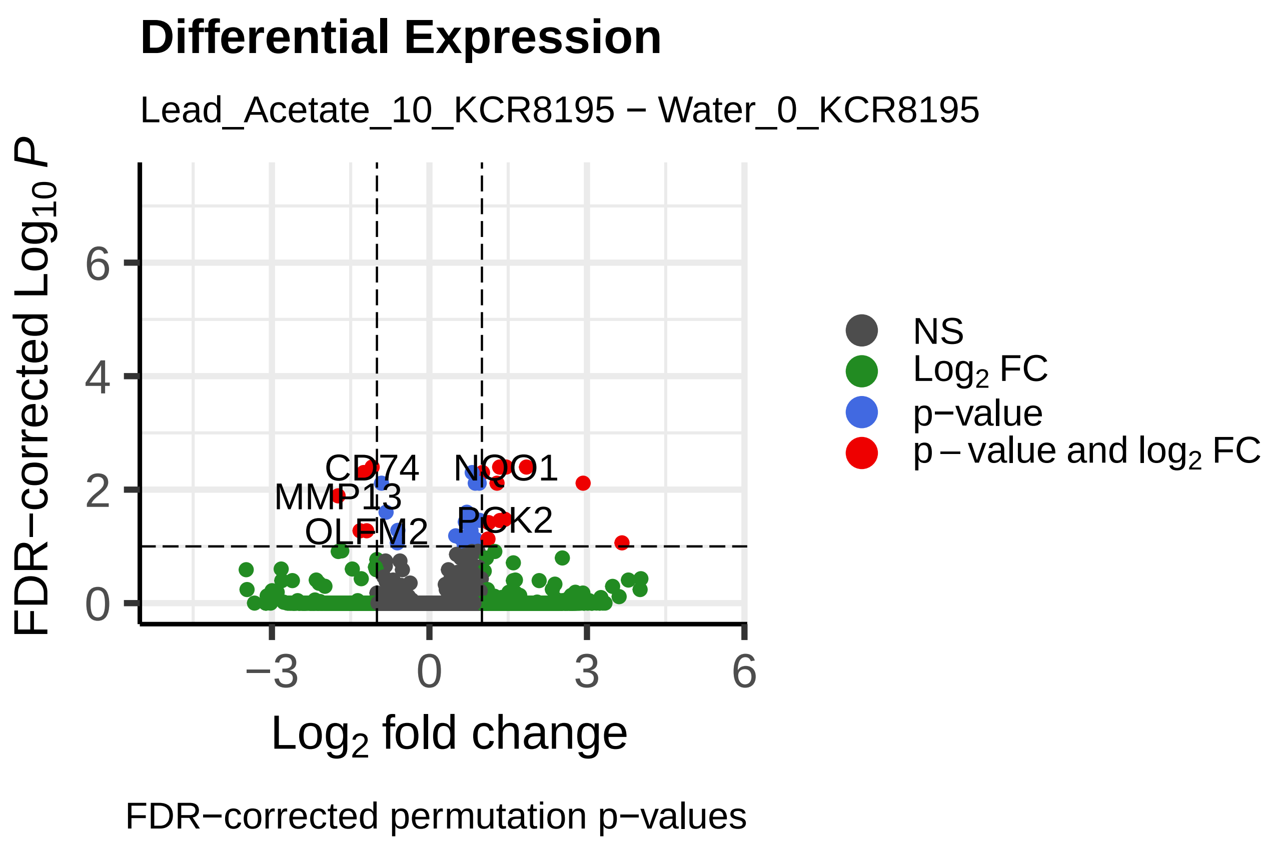
Differential Expression
Lead_Acetate_10_KCR8195 − Water_0_KCR8195
FDR−corrected Log10 P
Log2 fold change
NS
Log2 FC
p−value
p − value and log2 FC
CD74
NQO1
MMP13
OLFM2
PCK2
FDR−corrected permutation p−values

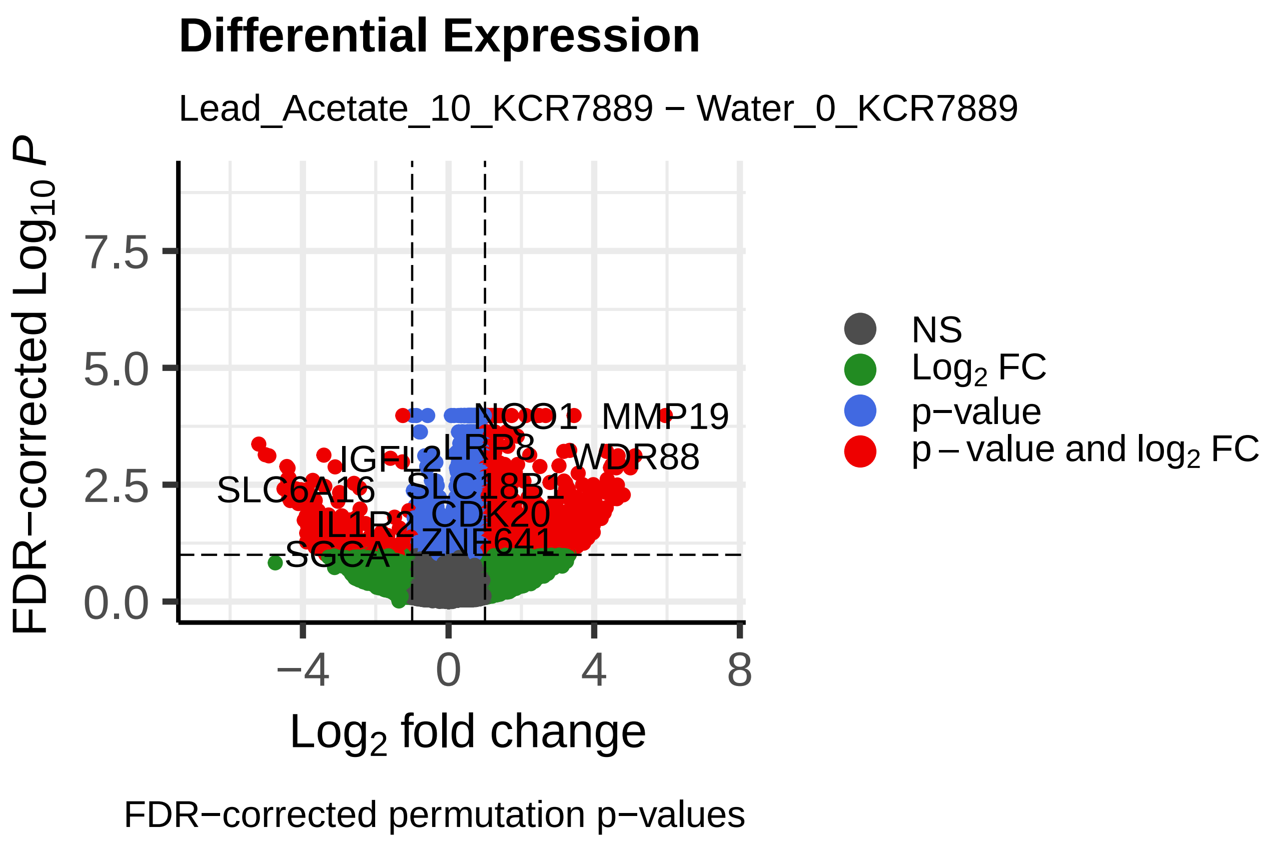
Differential Expression
Lead_Acetate_10_KCR7889 − Water_0_KCR7889
FDR−corrected Log10 P
Log2 fold change
NS
Log2 FC
p−value
p − value and log2 FC
NQO1
MMP19
LRP8
WDR88
SLC6A16
SLC18B1
CDK20
IL1R2
ZNF641
SGCA
FDR−corrected permutation p−values

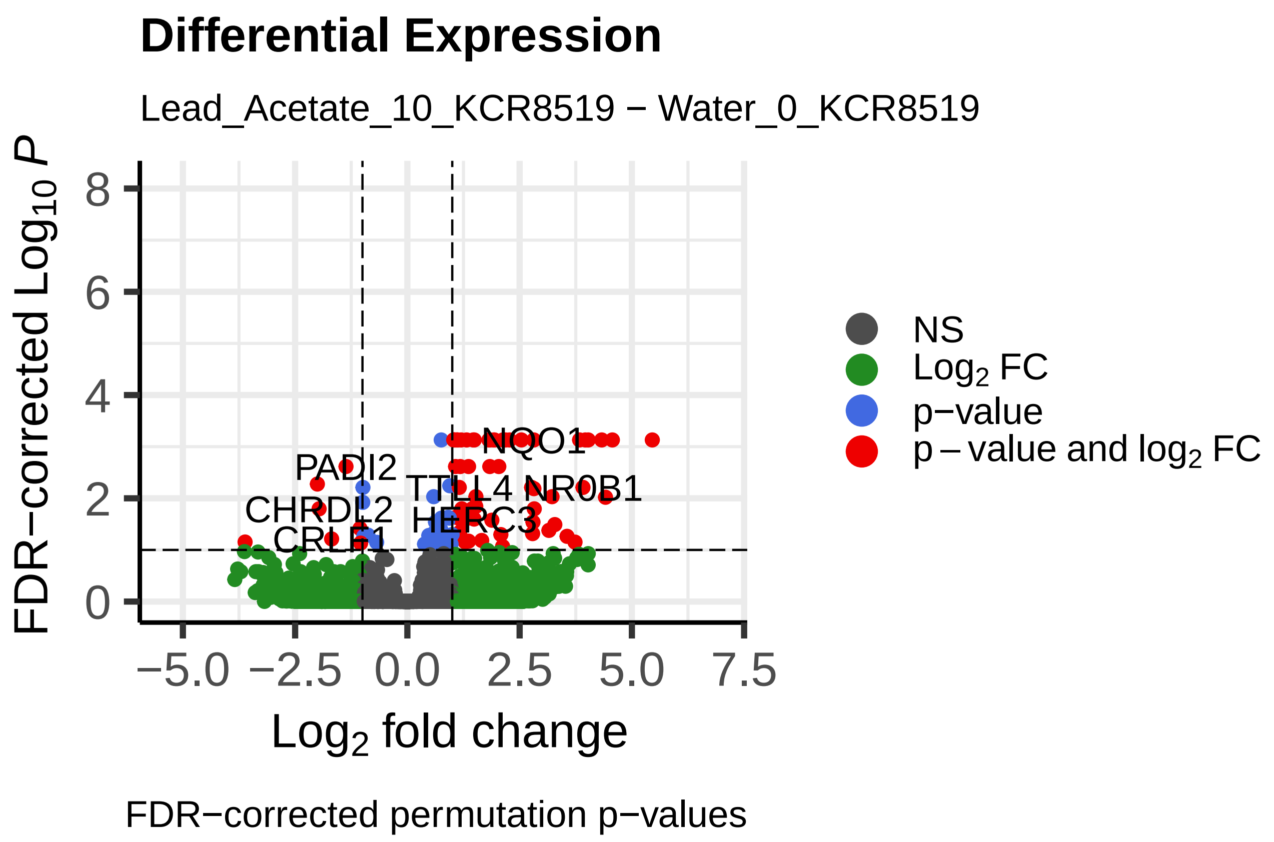
Differential Expression
Lead_Acetate_10_KCR8519 − Water_0_KCR8519
FDR−corrected Log10 P
Log2 fold change
NS
Log2 FC
p−value
p − value and log2 FC
NQO1
PADI2
NR0B1
CHRDL2
HERC3
FDR−corrected permutation p−values

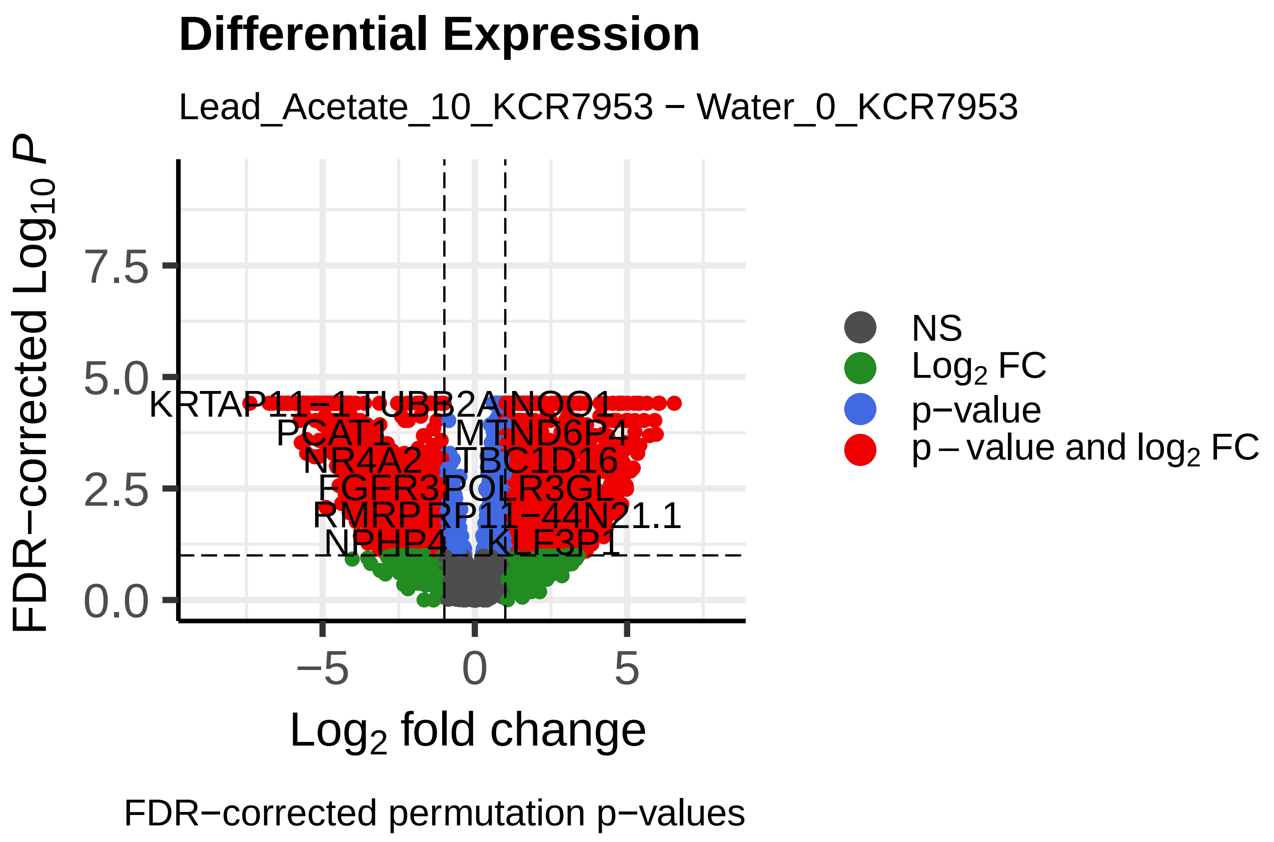
Differential Expression
Lead_Acetate_10_KCR7953 − Water_0_KCR7953
FDR−corrected Log10 P
Log2 fold change
NS
Log2 FC
p−value
p − value and log2 FC
KRTAP11−1
NQO1
PGAT1
NR4A2
TBC1D16
FGFR3
POLR3GL
RMRP
RP11−44N21.1
NPHP4
KLF3P1
FDR−corrected permutation p−values

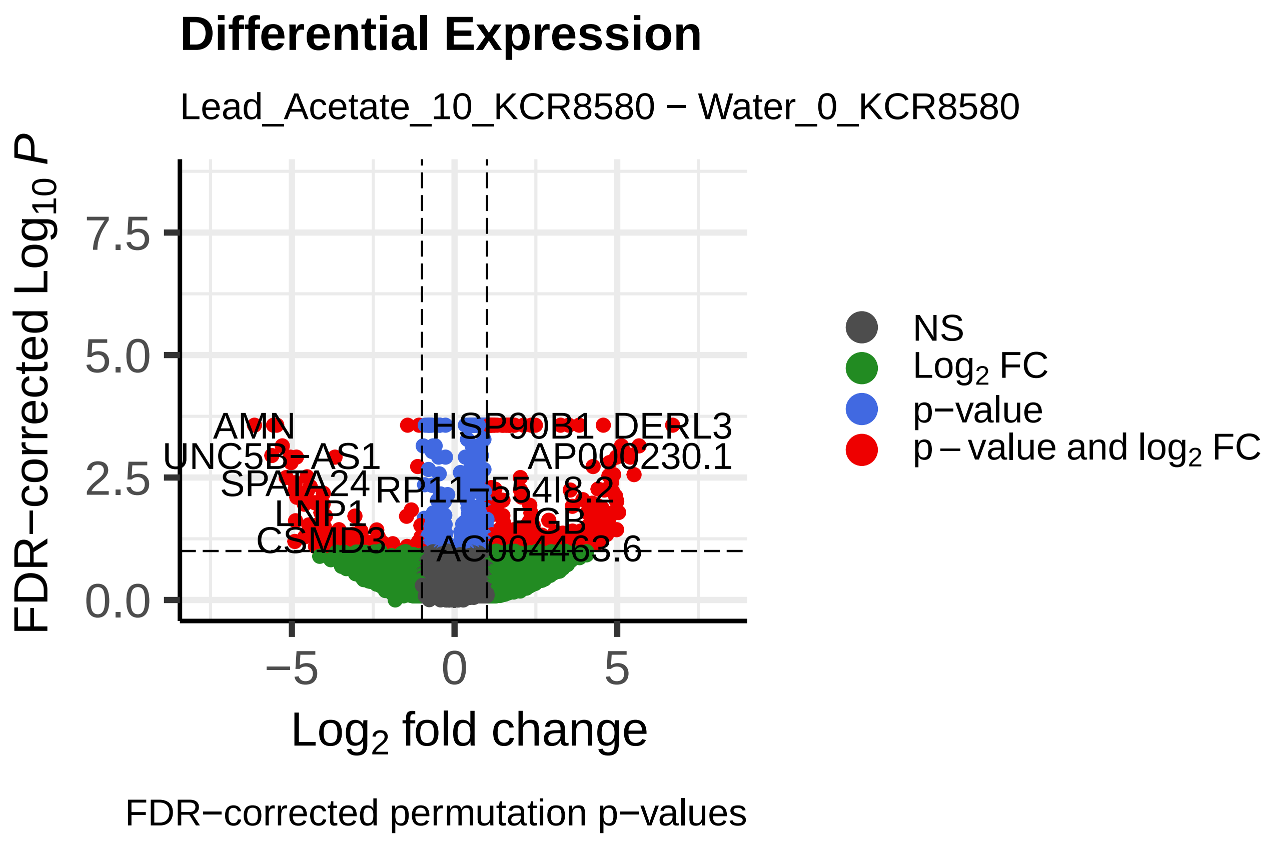
Differential Expression
Lead_Acetate_10_KCR8580 − Water_0_KCR8580
FDR−corrected Log10 P
Log2 fold change
NS
Log2 FC
p−value
p − value and log2 FC
AMN
HSP90B1
DERL3
UNC5B−AS1
AP000230.1
SPATA24
LNP1
CSMD3
AC004463.6
FDR−corrected permutation p−values

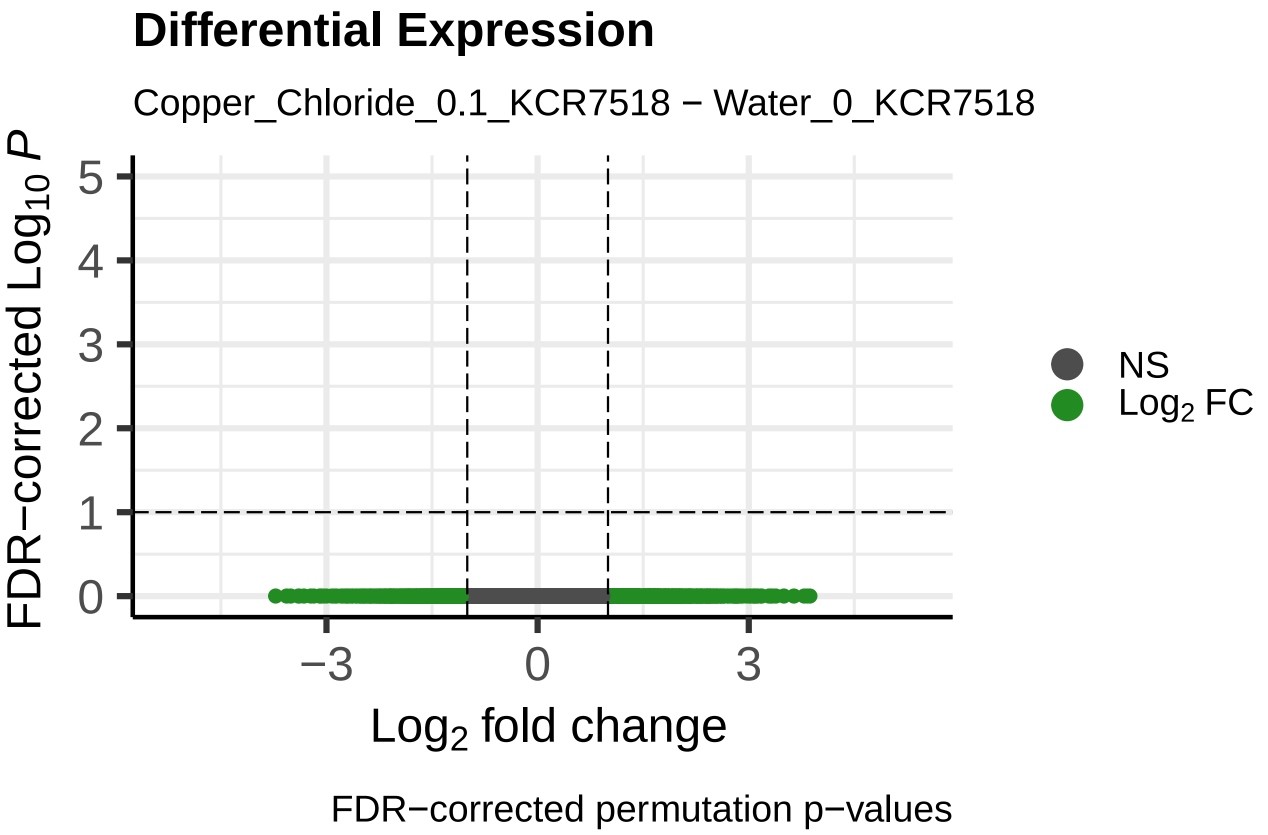
Differential Expression
Copper_Chloride_0.1_KCR7518 − Water_0_KCR7518
FDR−corrected Log10 P
Log2 fold change
NS
Log2 FC
FDR−corrected permutation p−values

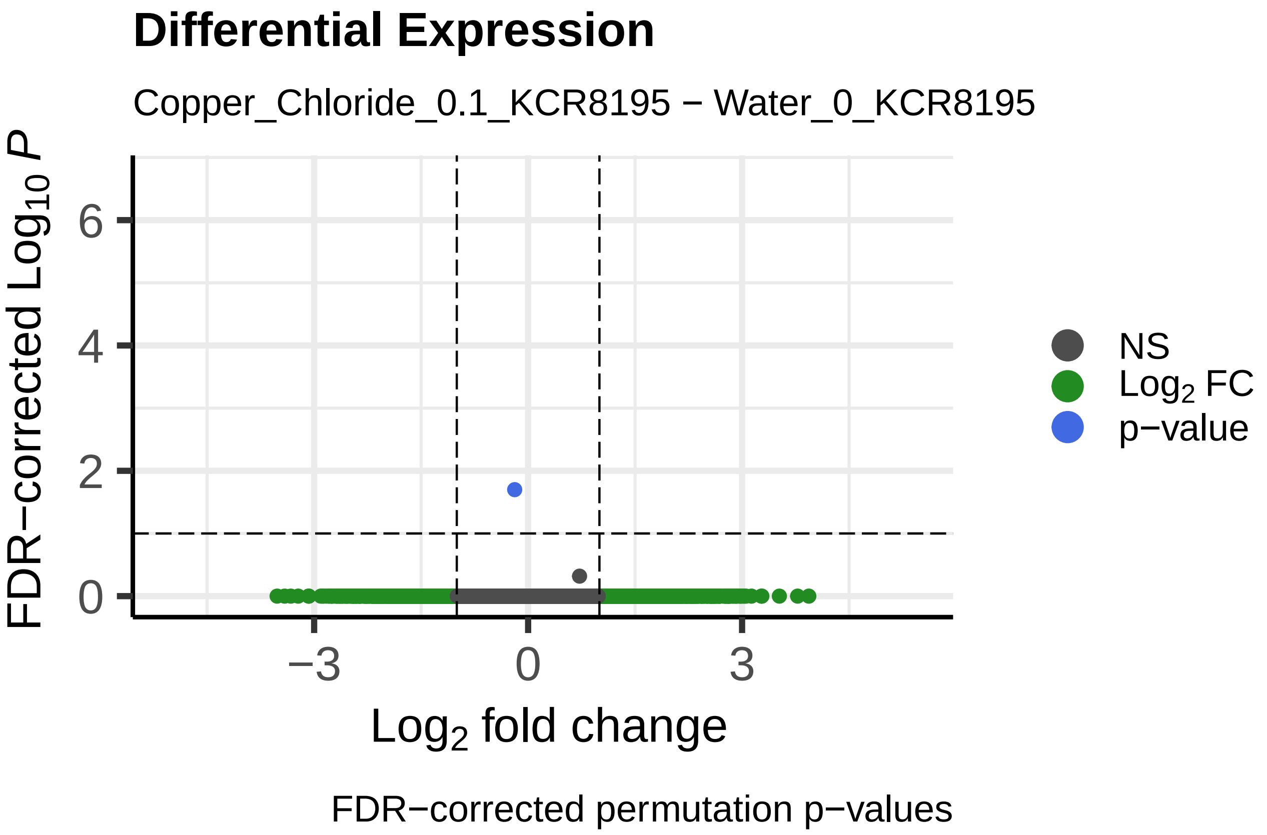
Differential Expression
Copper_Chloride_0.1_KCR8195 − Water_0_KCR8195
FDR−corrected Log10 P
Log2 fold change
NS
Log2 FC
p−value
FDR−corrected permutation p−values

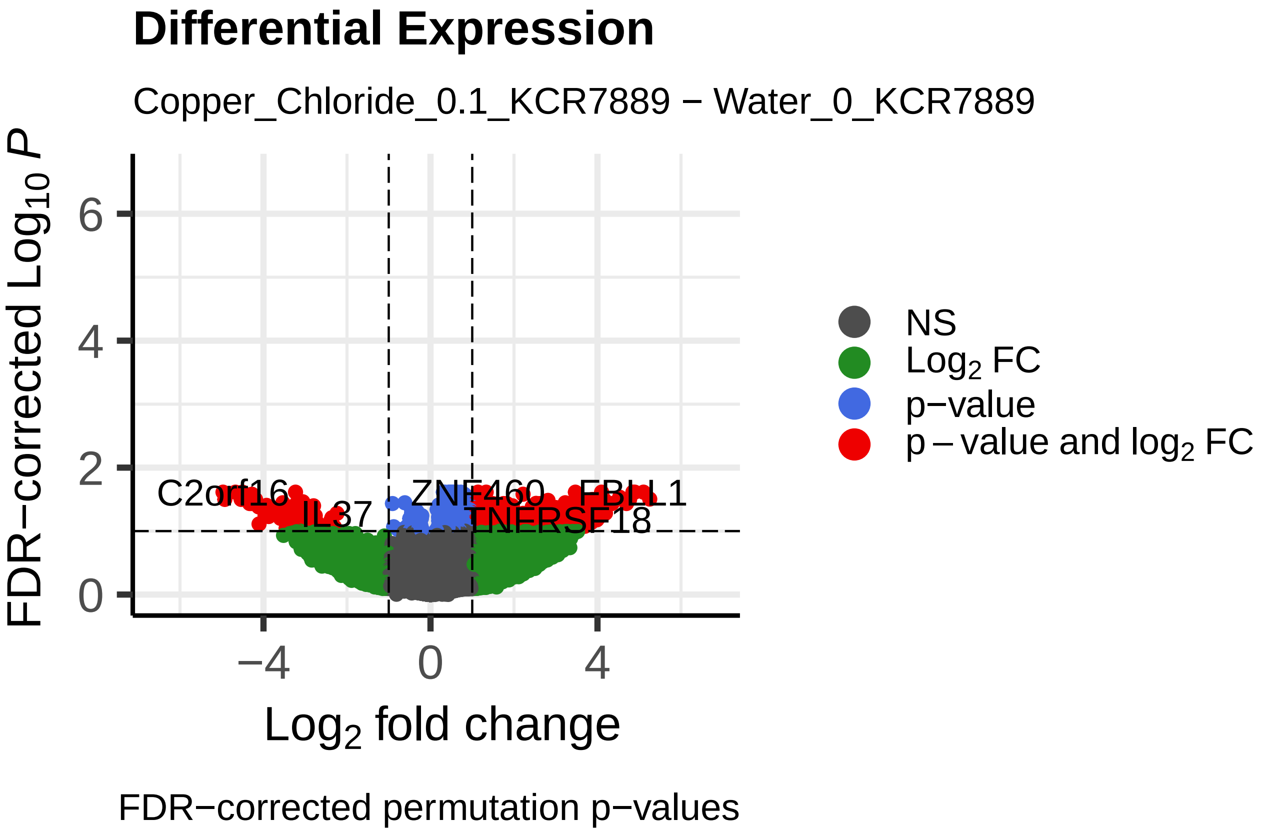
Differential Expression
Copper_Chloride_0.1_KCR7889 − Water_0_KCR7889
FDR−corrected Log10 P
Log2 fold change
C2orf16
IL37
ZNF460
FBLL1
TNFRSF18
NS
Log2 FC
p−value
p − value and log2 FC
FDR−corrected permutation p−values

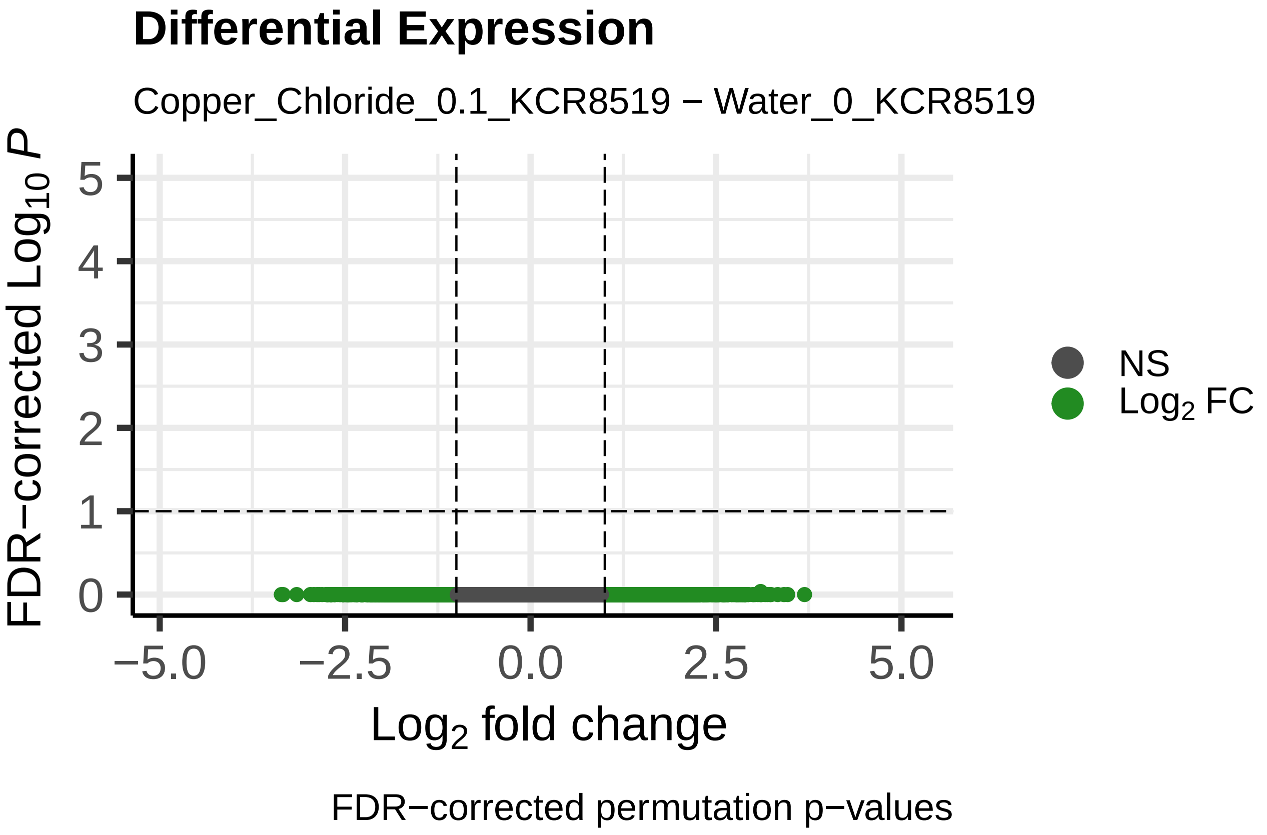
Differential Expression
Copper_Chloride_0.1_KCR8519 − Water_0_KCR8519
FDR−corrected Log10 P
Log2 fold change
NS
Log2 FC
FDR−corrected permutation p−values

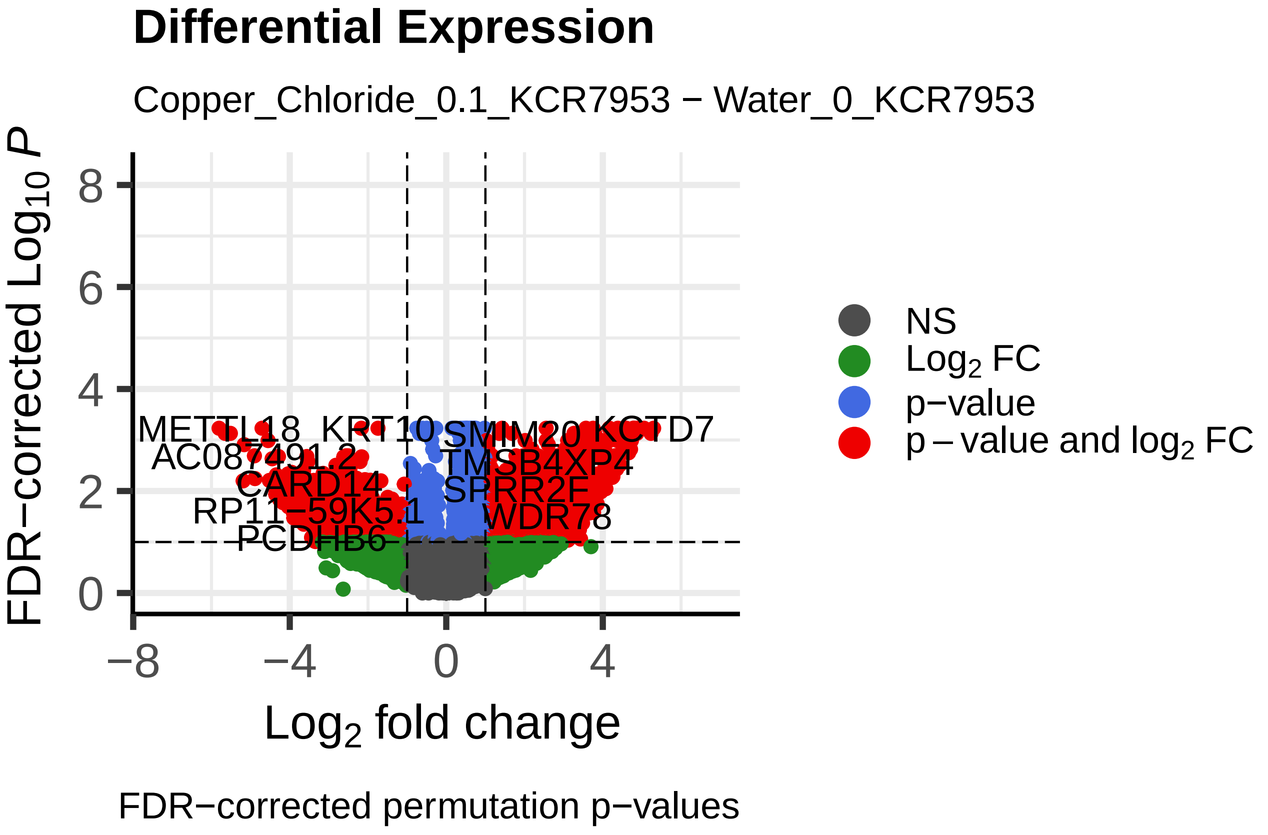
Differential Expression
Copper_Chloride_0.1_KCR7953 − Water_0_KCR7953
FDR−corrected Log10 P
Log2 fold change
METTL18
KRT10
SMIM20
KCTD7
TMSB4XP4
SPRR2F
CARD14
WDR78
PCDHB6
NS
Log2 FC
p−value
p − value and log2 FC
FDR−corrected permutation p−values

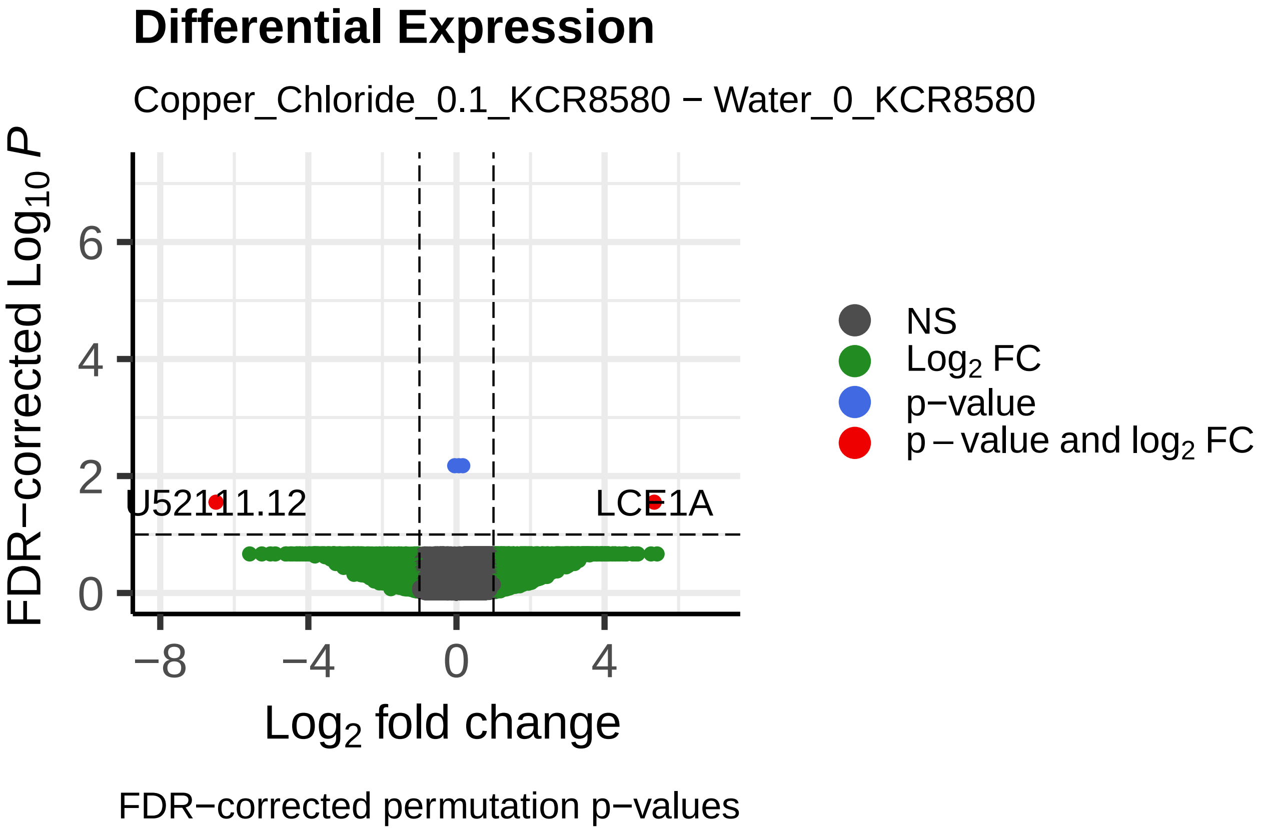
Differential Expression
Copper_Chloride_0.1_KCR8580 − Water_0_KCR8580
FDR−corrected Log10 P
Log2 fold change
U52111.12
LCE1A
NS
Log2 FC
p−value
p − value and log2 FC
FDR−corrected permutation p−values

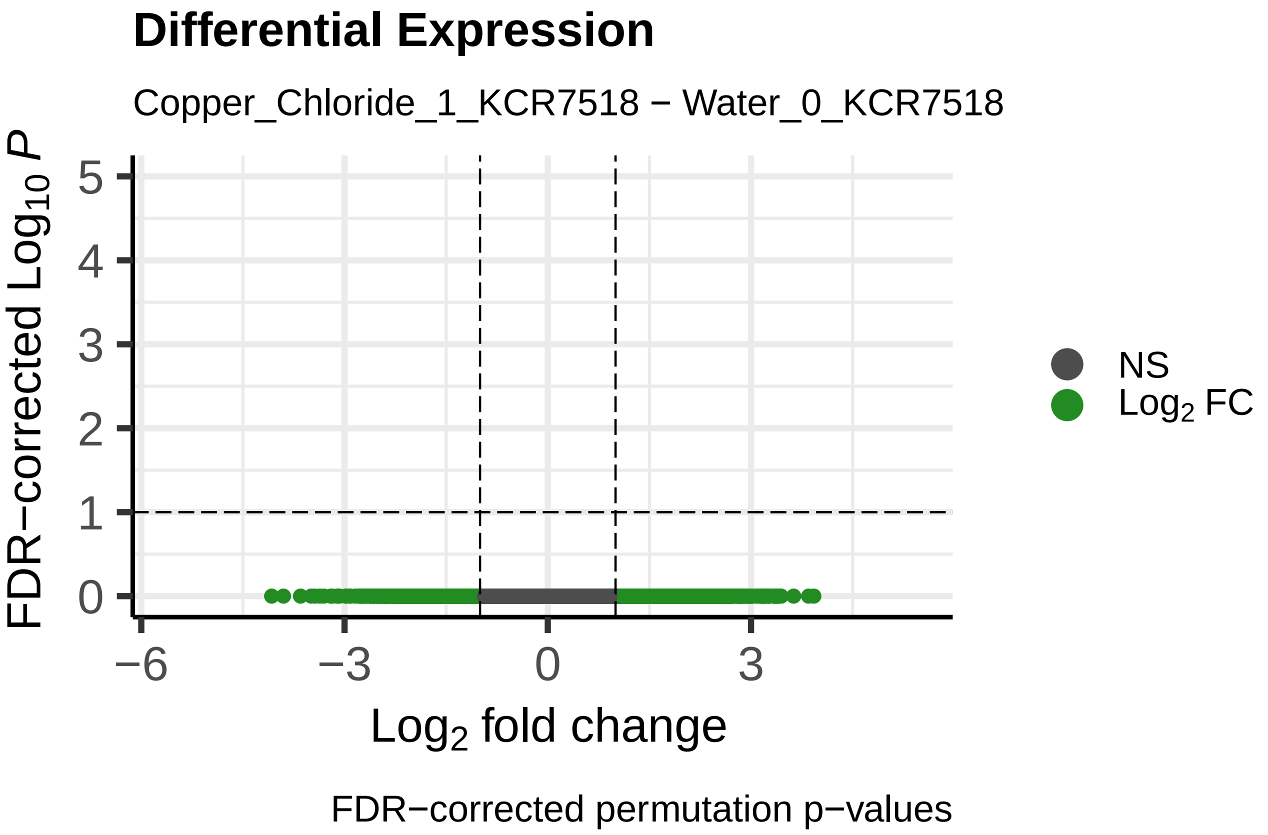
Differential Expression
Copper_Chloride_1_KCR7518 − Water_0_KCR7518
FDR−corrected Log10 P
Log2 fold change
NS
Log2 FC
FDR−corrected permutation p−values

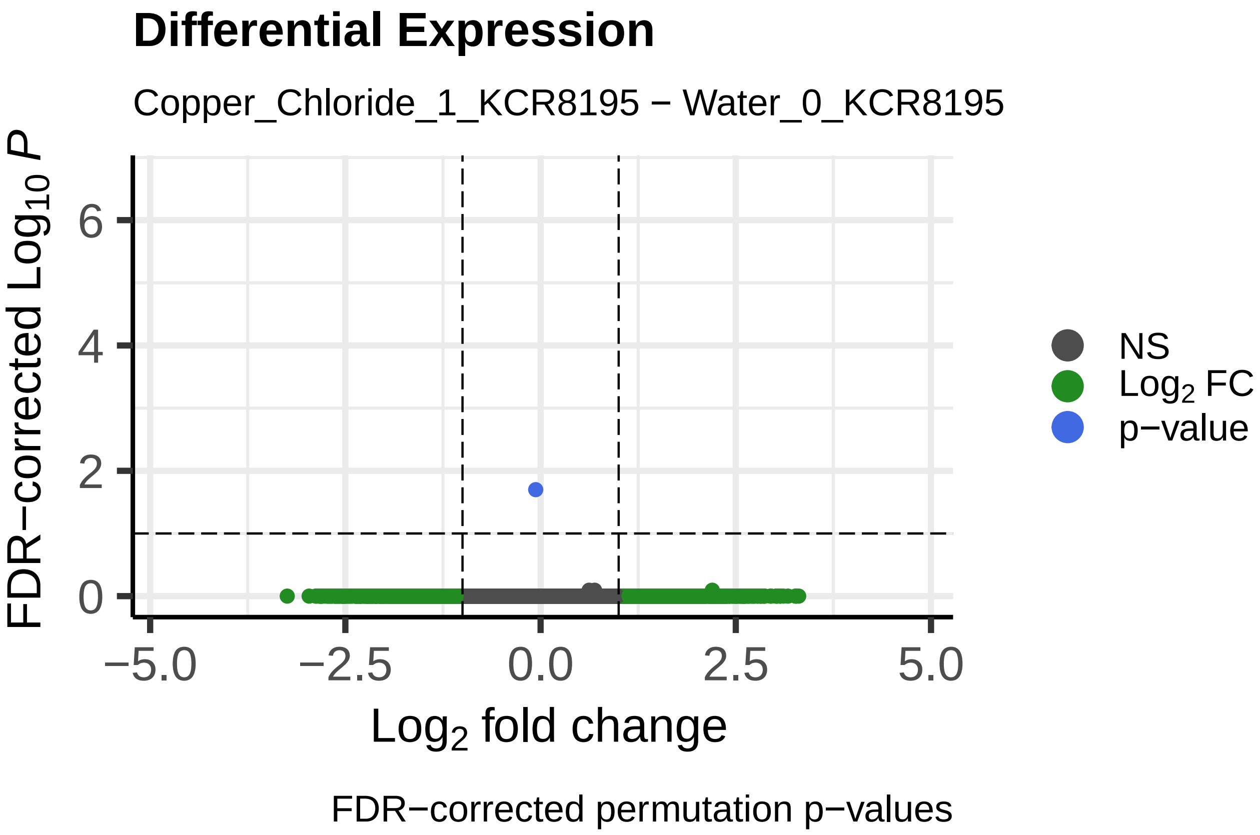
Differential Expression
Copper_Chloride_1_KCR8195 − Water_0_KCR8195
FDR−corrected Log10 P
Log2 fold change
NS
Log2 FC
p−value
FDR−corrected permutation p−values

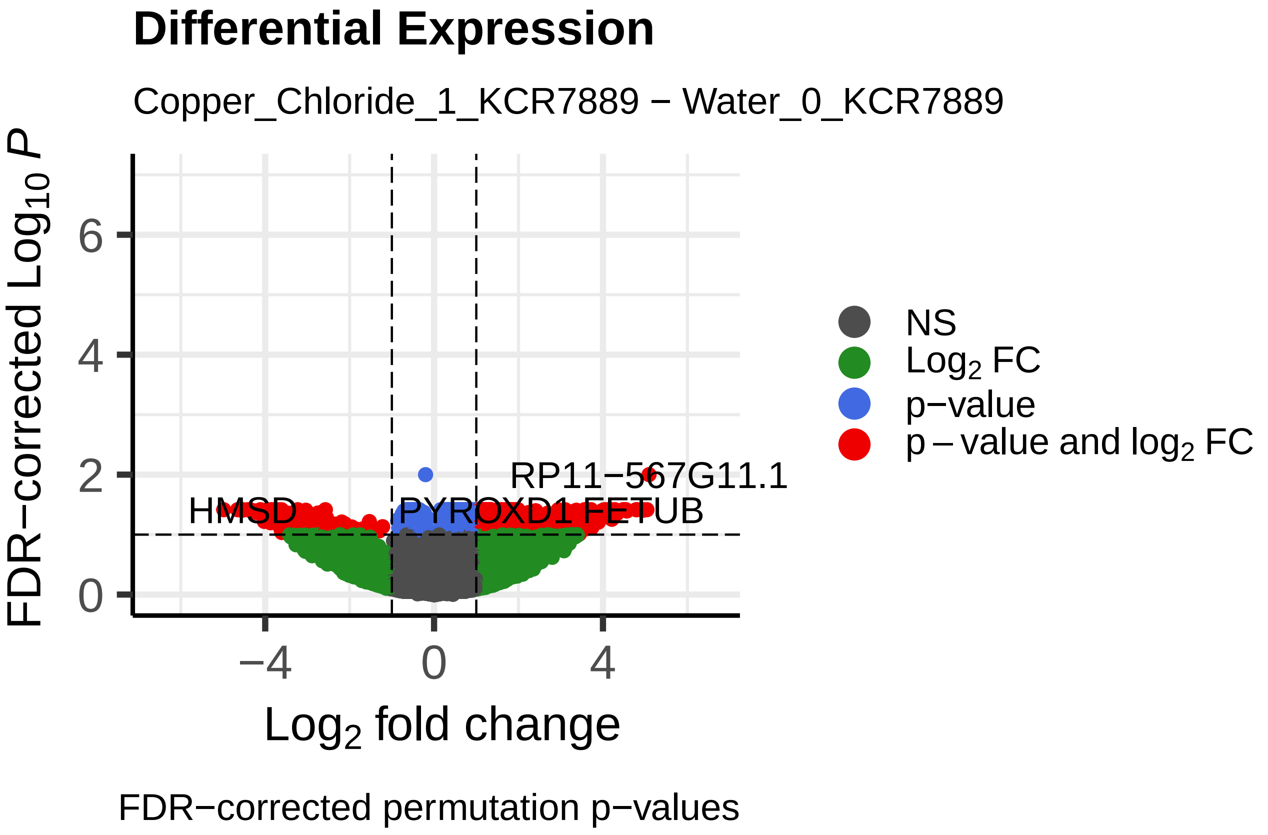
Differential Expression
Copper_Chloride_1_KCR7889 − Water_0_KCR7889
FDR−corrected Log10 P
RP11−567G11.1
HMSD
PYROXD1
FETUB
Log2 fold change
NS
Log2 FC
p−value
p − value and log2 FC
FDR−corrected permutation p−values

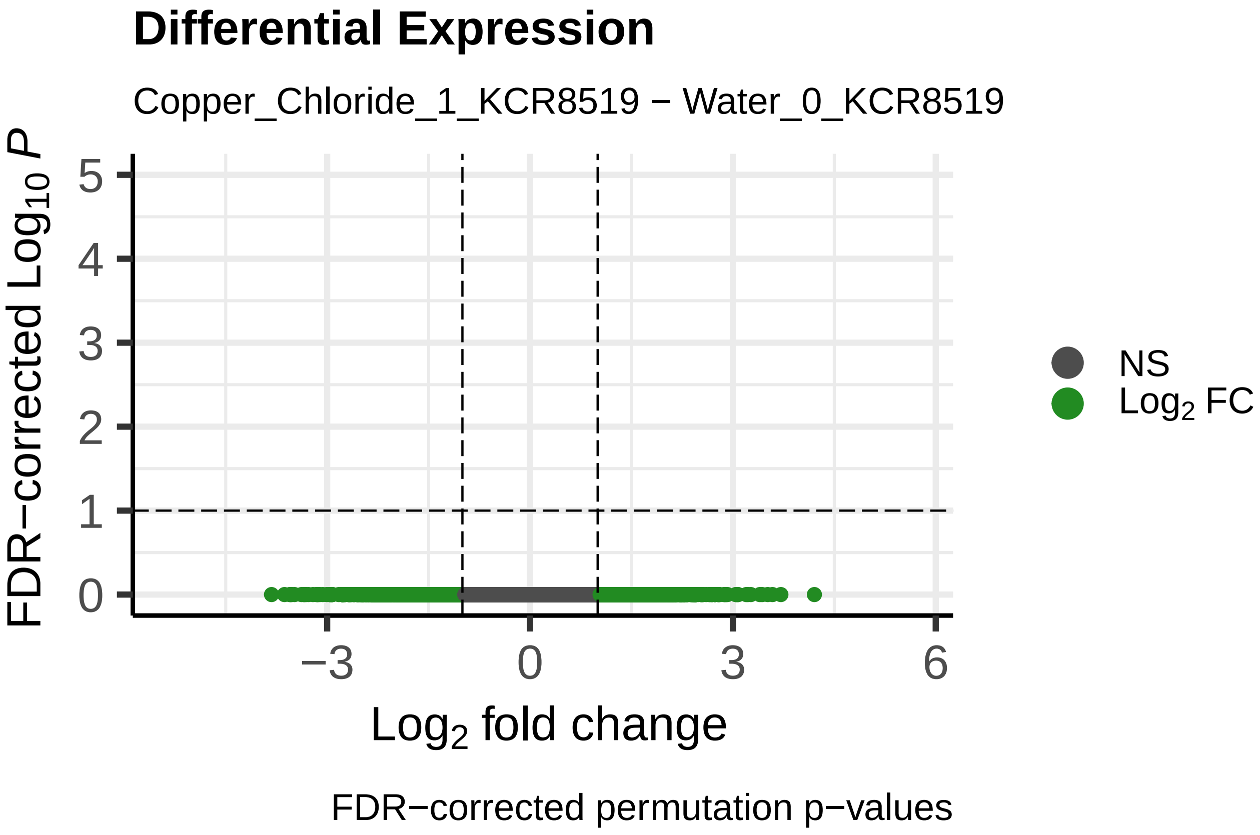
Differential Expression
Copper_Chloride_1_KCR8519 − Water_0_KCR8519
FDR−corrected Log10 P
Log2 fold change
NS
Log2 FC
FDR−corrected permutation p−values

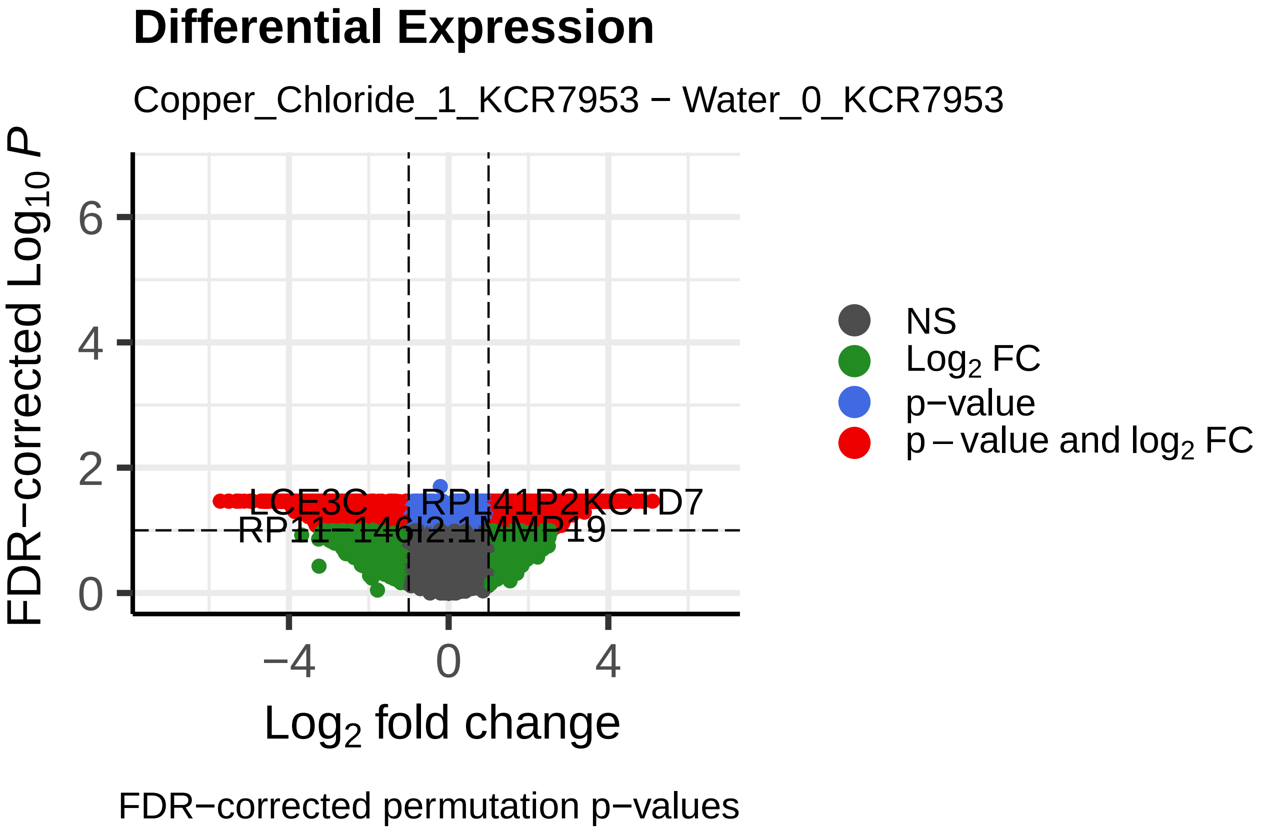
Differential Expression
Copper_Chloride_1_KCR7953 − Water_0_KCR7953
FDR−corrected Log10 P
LCE3C
RPL41P2
KCTD7
RP11−146I2.1
MMP19
Log2 fold change
NS
Log2 FC
p−value
p − value and log2 FC
FDR−corrected permutation p−values

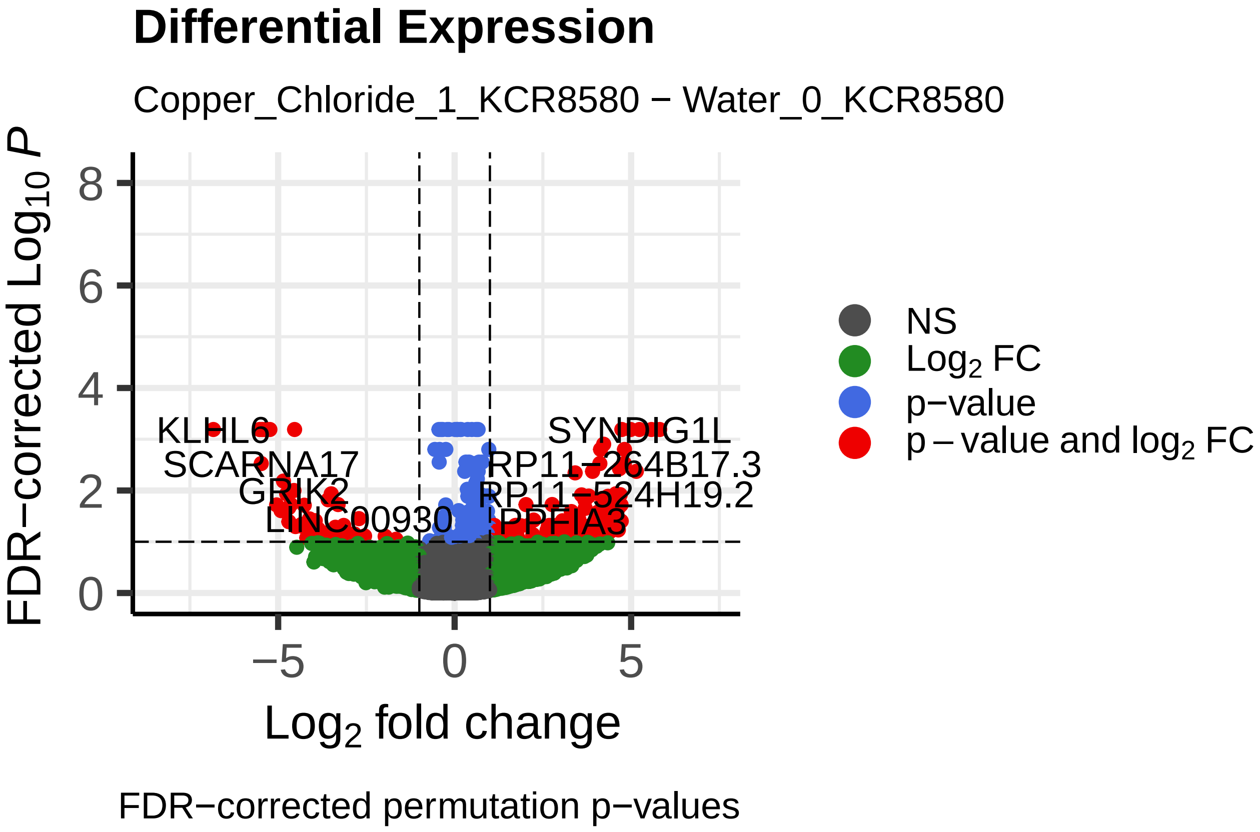
Differential Expression
Copper_Chloride_1_KCR8580 − Water_0_KCR8580
FDR−corrected Log10 P
KLHL6
SCARNA17
LINC00930
SYNDIG1L
RP11−264B17.3
RP11−524H19.2
PPFIA3
Log2 fold change
NS
Log2 FC
p−value
p − value and log2 FC
FDR−corrected permutation p−values

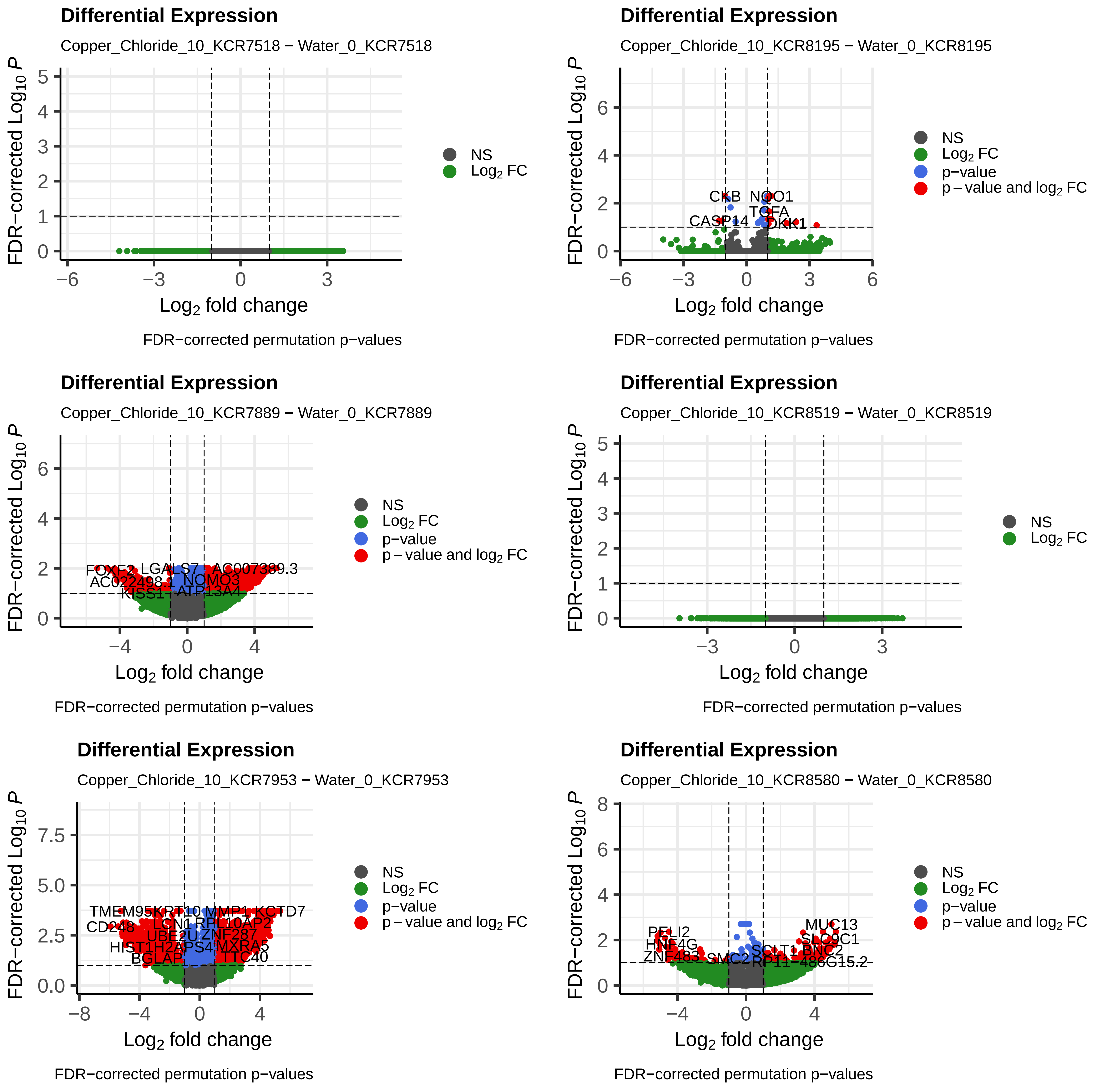

Differential Expression
Copper_Chloride_10_KCR7518 − Water_0_KCR7518
FDR−corrected Log10 P
Log2 fold change
NS
Log2 FC
FDR−corrected permutation p−values
Differential Expression
Copper_Chloride_10_KCR8195 − Water_0_KCR8195
CKB
NQO1
TGFA
CASP14
DKK1
FDR−corrected Log10 P
Log2 fold change
NS
Log2 FC
p−value
p − value and log2 FC
FDR−corrected permutation p−values
Differential Expression
Copper_Chloride_10_KCR7889 − Water_0_KCR7889
FOXF2
LGALS7
AC007389.3
AC022498.1
NOMO3
KISS1
ATP13A4
FDR−corrected Log10 P
Log2 fold change
NS
Log2 FC
p−value
p − value and log2 FC
FDR−corrected permutation p−values
Differential Expression
Copper_Chloride_10_KCR8519 − Water_0_KCR8519
FDR−corrected Log10 P
Log2 fold change
NS
Log2 FC
FDR−corrected permutation p−values
Differential Expression
Copper_Chloride_10_KCR7953 − Water_0_KCR7953
TMEM95
KRT10
MMP1
KCTD7
CD248
LCN1
RPL10AP2
UBE2U
ZNF287
HIST1H2APS4
MXRA5
BGLAP
TTC40
FDR−corrected Log10 P
Log2 fold change
NS
Log2 FC
p−value
p − value and log2 FC
FDR−corrected permutation p−values
Differential Expression
Copper_Chloride_10_KCR8580 − Water_0_KCR8580
PELI2
MUC13
HNF4G
SLC9C1
ZNF483
SMC2
SCLT1
BNC2
RP11−486G15.2
FDR−corrected Log10 P
Log2 fold change
NS
Log2 FC
p−value
p − value and log2 FC
FDR−corrected permutation p−values

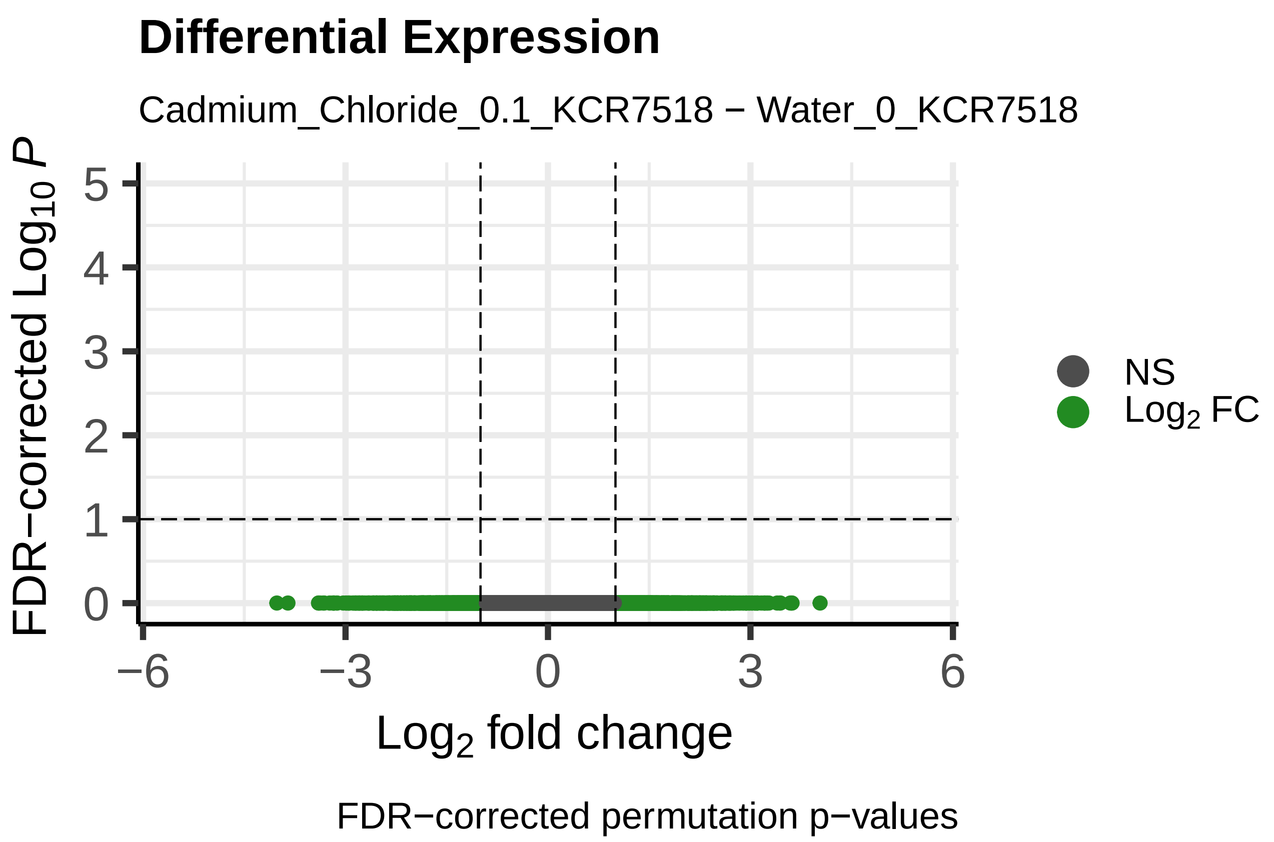
Differential Expression
Cadmium_Chloride_0.1_KCR7518 − Water_0_KCR7518
FDR−corrected Log10 P
Log2 fold change
NS
Log2 FC
FDR−corrected permutation p−values

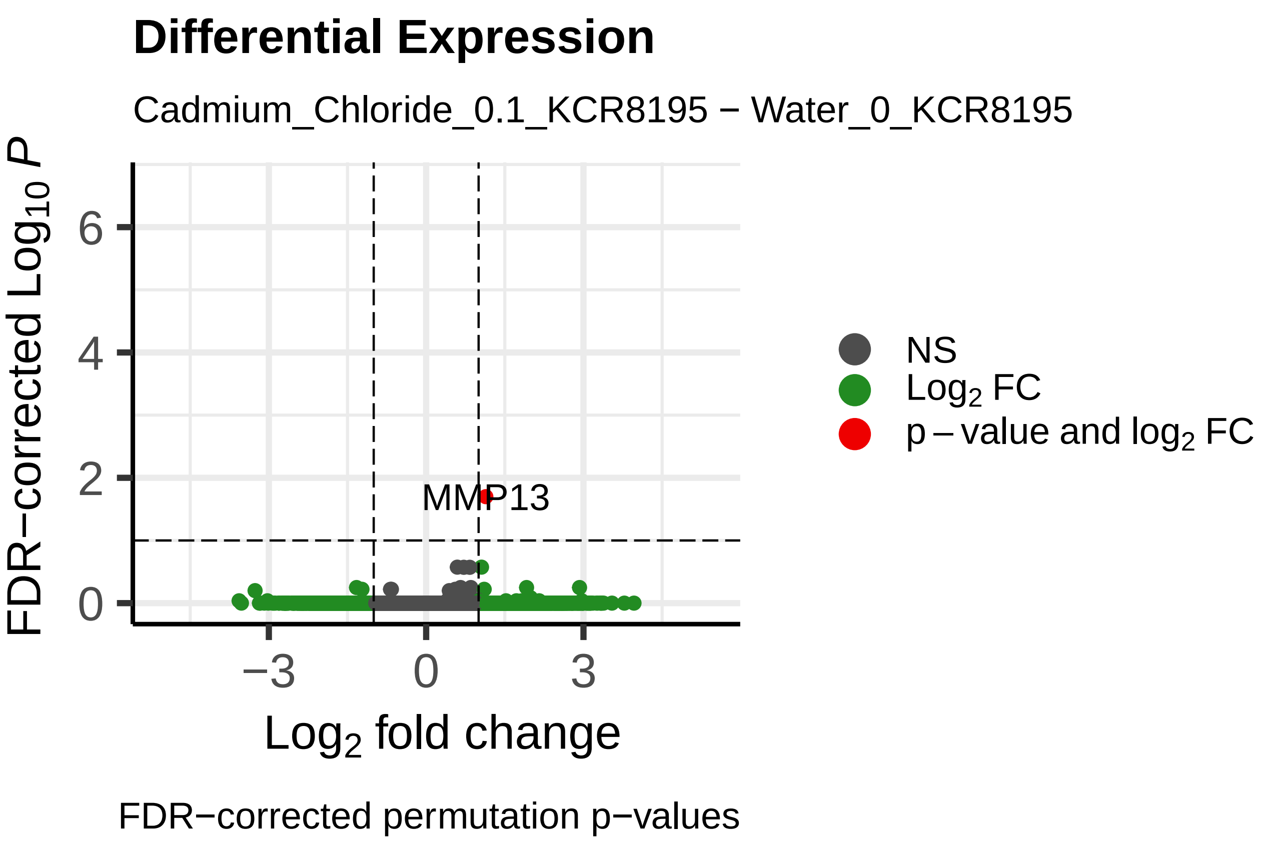
Differential Expression
Cadmium_Chloride_0.1_KCR8195 − Water_0_KCR8195
MMP13
FDR−corrected Log10 P
Log2 fold change
NS
Log2 FC
p − value and log2 FC
FDR−corrected permutation p−values

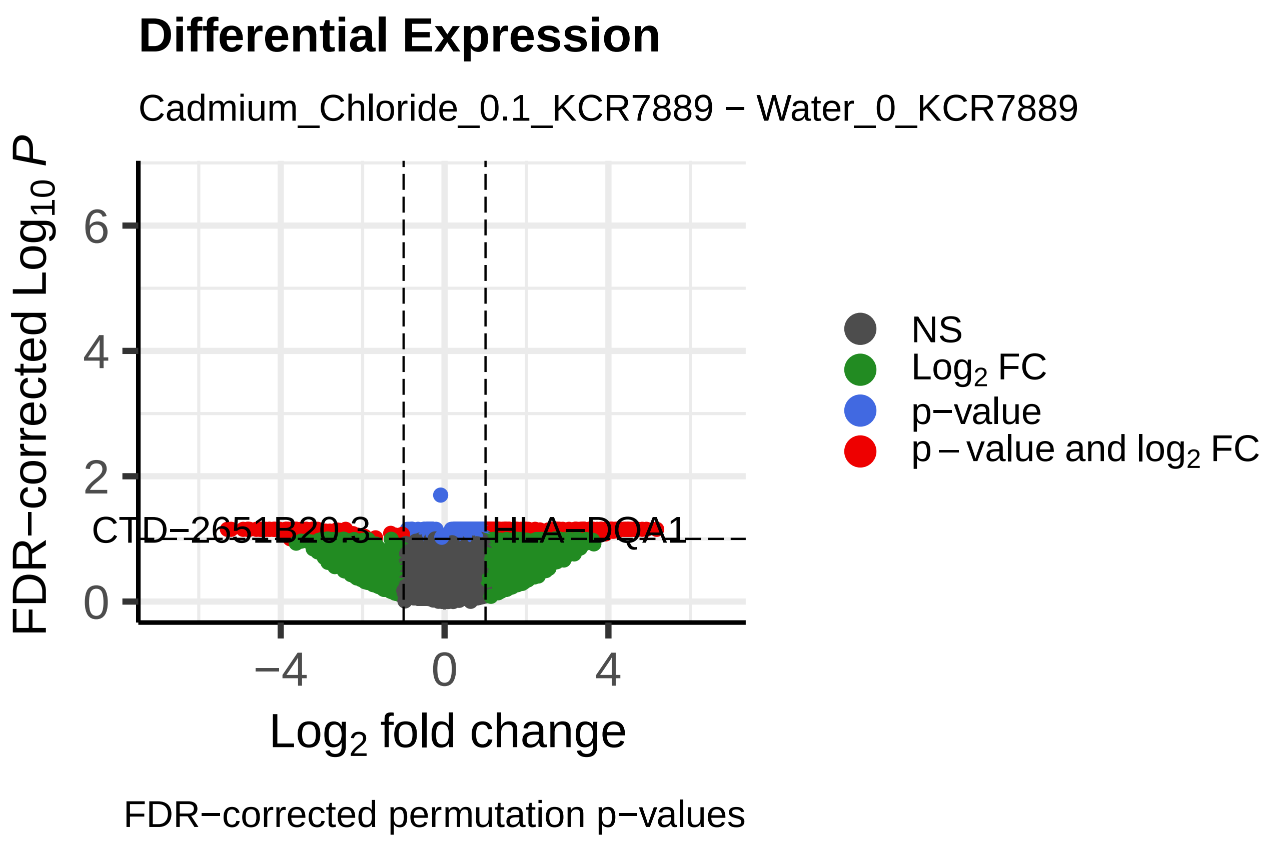
Differential Expression
Cadmium_Chloride_0.1_KCR7889 − Water_0_KCR7889
CTD−2651B20.3
HLA−DQA1
FDR−corrected Log10 P
Log2 fold change
NS
Log2 FC
p−value
p − value and log2 FC
FDR−corrected permutation p−values

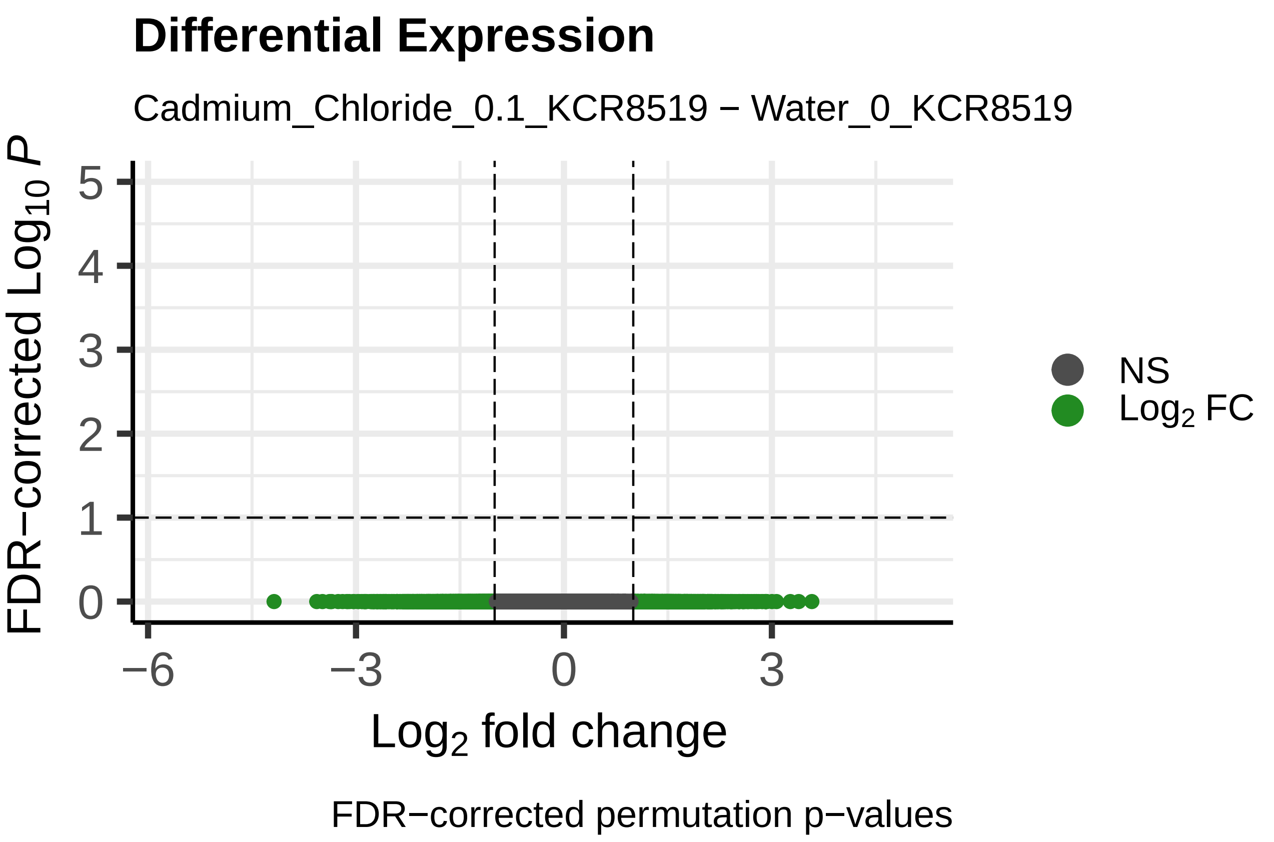
Differential Expression
Cadmium_Chloride_0.1_KCR8519 − Water_0_KCR8519
FDR−corrected Log10 P
Log2 fold change
NS
Log2 FC
FDR−corrected permutation p−values

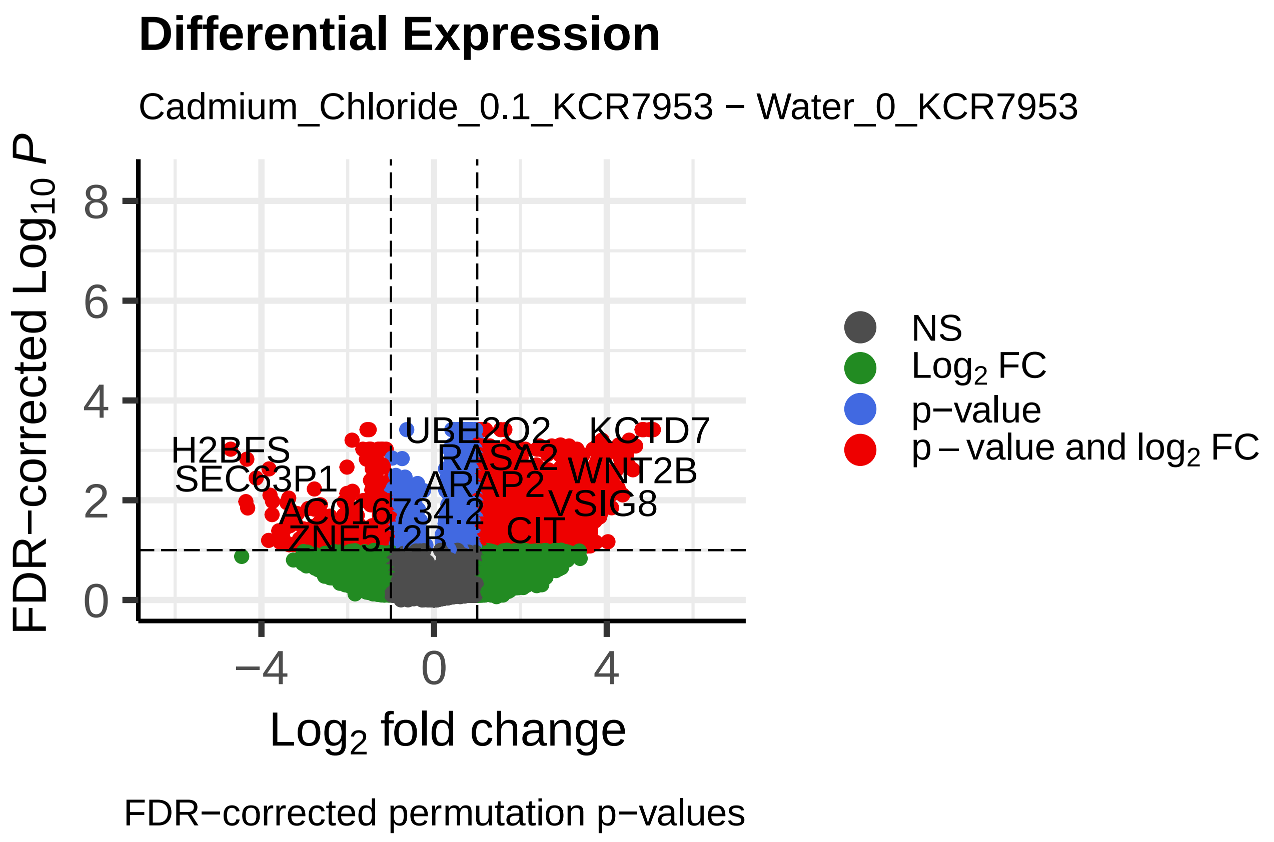
Differential Expression
Cadmium_Chloride_0.1_KCR7953 − Water_0_KCR7953
H2BFS
SEC63P1
AC016734.2
ZNF512B
UBE2Q2
RASA2
ARAP2
CIT
KCTD7
WNT2B
VSIG8
FDR−corrected Log10 P
Log2 fold change
NS
Log2 FC
p−value
p − value and log2 FC
FDR−corrected permutation p−values

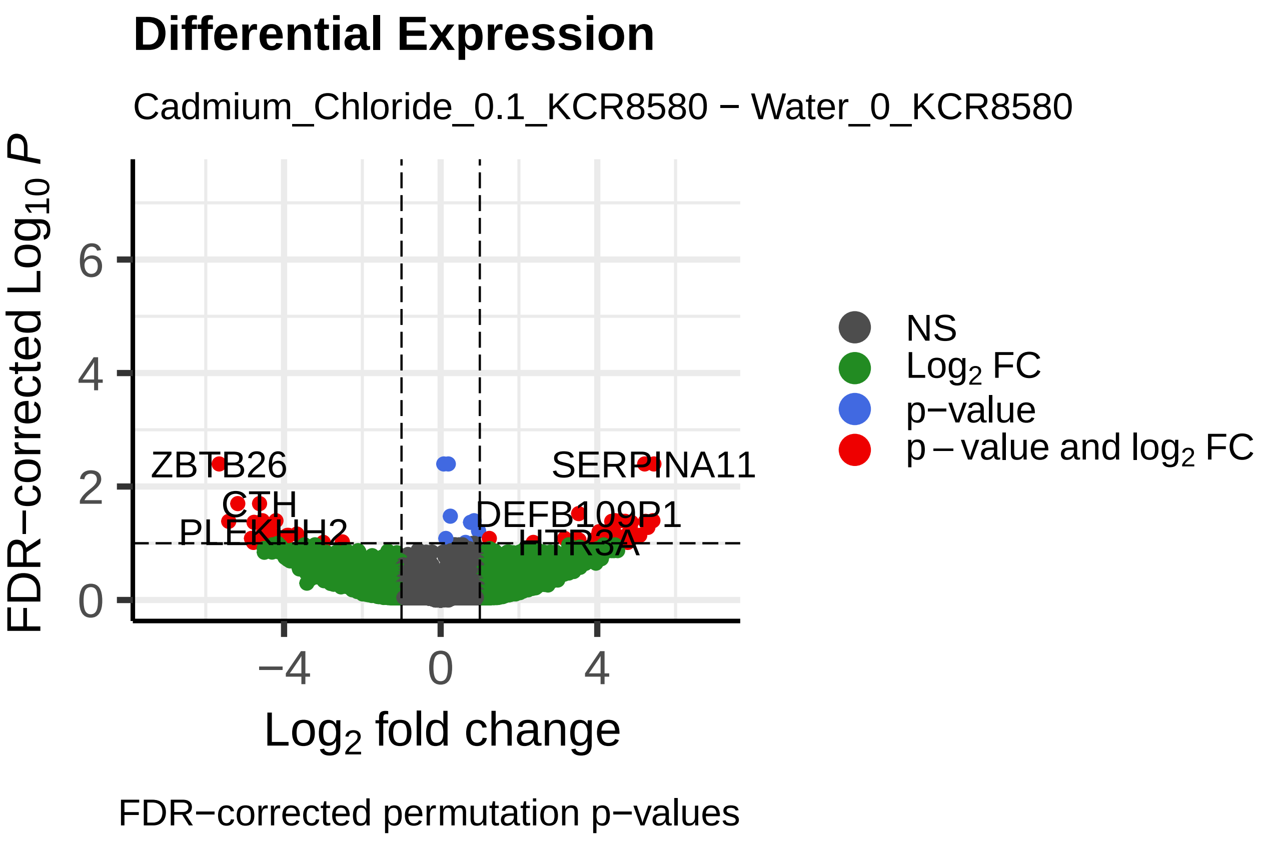
Differential Expression
Cadmium_Chloride_0.1_KCR8580 − Water_0_KCR8580
ZBTB26
CTH
PLEKHH2
SERPINA11
DEFB109P1
HTR3A
FDR−corrected Log10 P
Log2 fold change
NS
Log2 FC
p−value
p − value and log2 FC
FDR−corrected permutation p−values

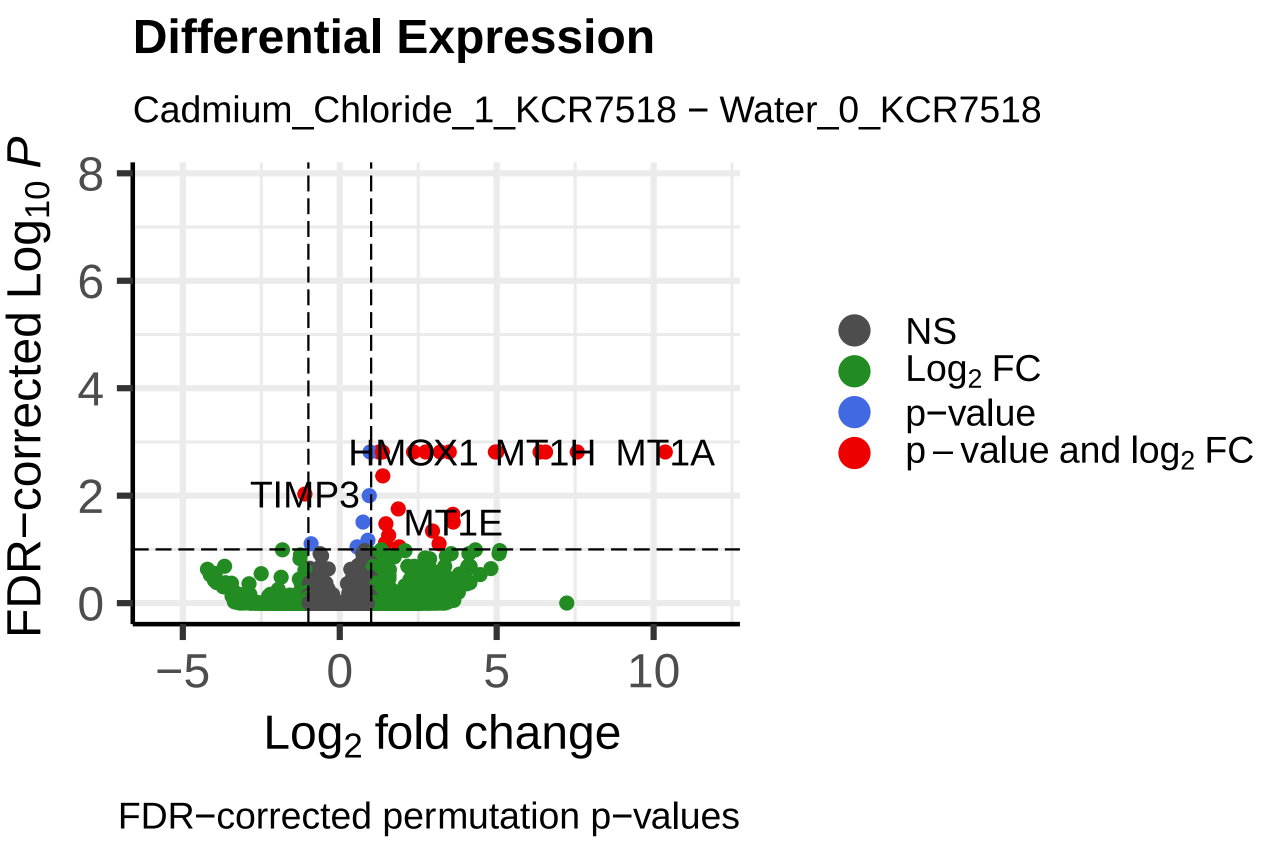
Differential Expression
Cadmium_Chloride_1_KCR7518 − Water_0_KCR7518
FDR−corrected Log10 P
Log2 fold change
HMOX1
MT1H
MT1A
TIMP3
MT1E
NS
Log2 FC
p−value
p − value and log2 FC
FDR−corrected permutation p−values

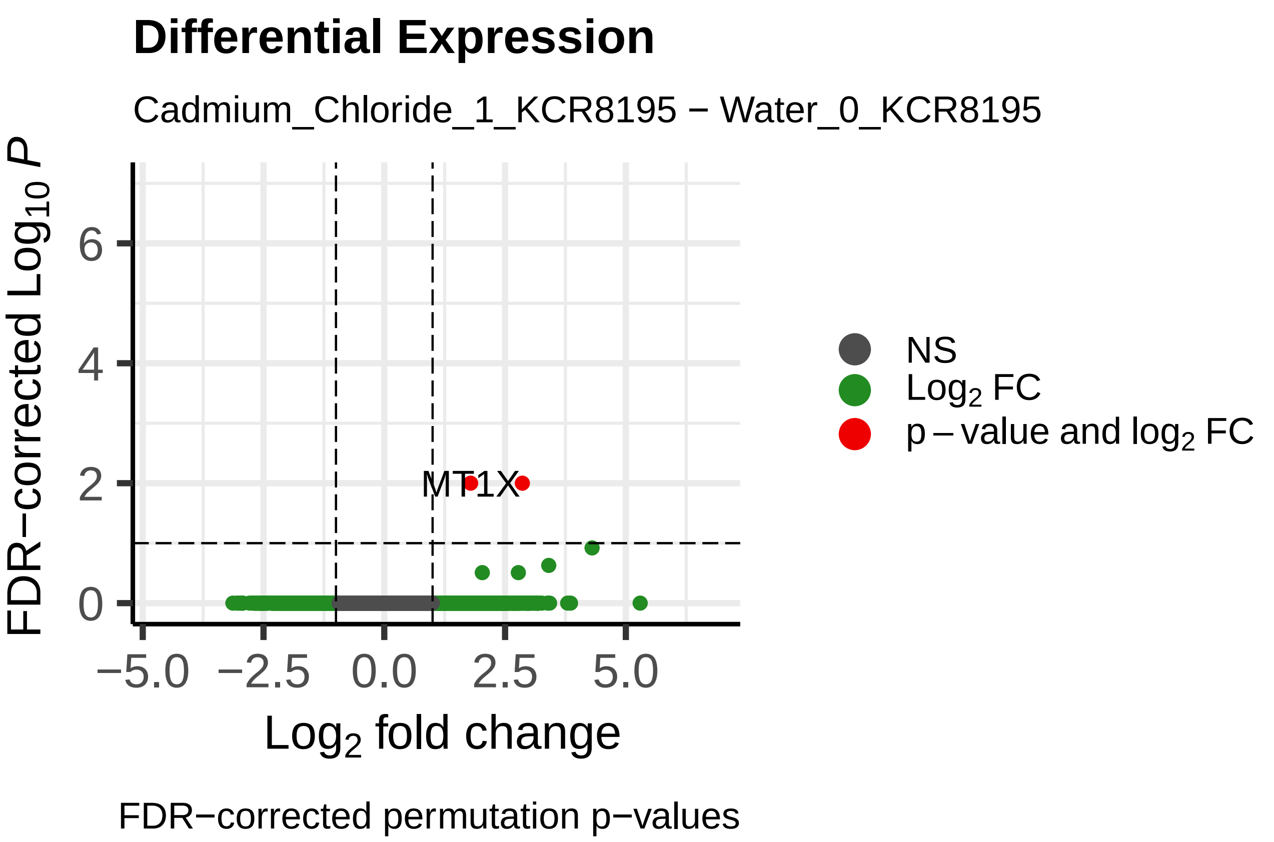
Differential Expression
Cadmium_Chloride_1_KCR8195 − Water_0_KCR8195
FDR−corrected Log10 P
Log2 fold change
MT1X
NS
Log2 FC
p − value and log2 FC
FDR−corrected permutation p−values

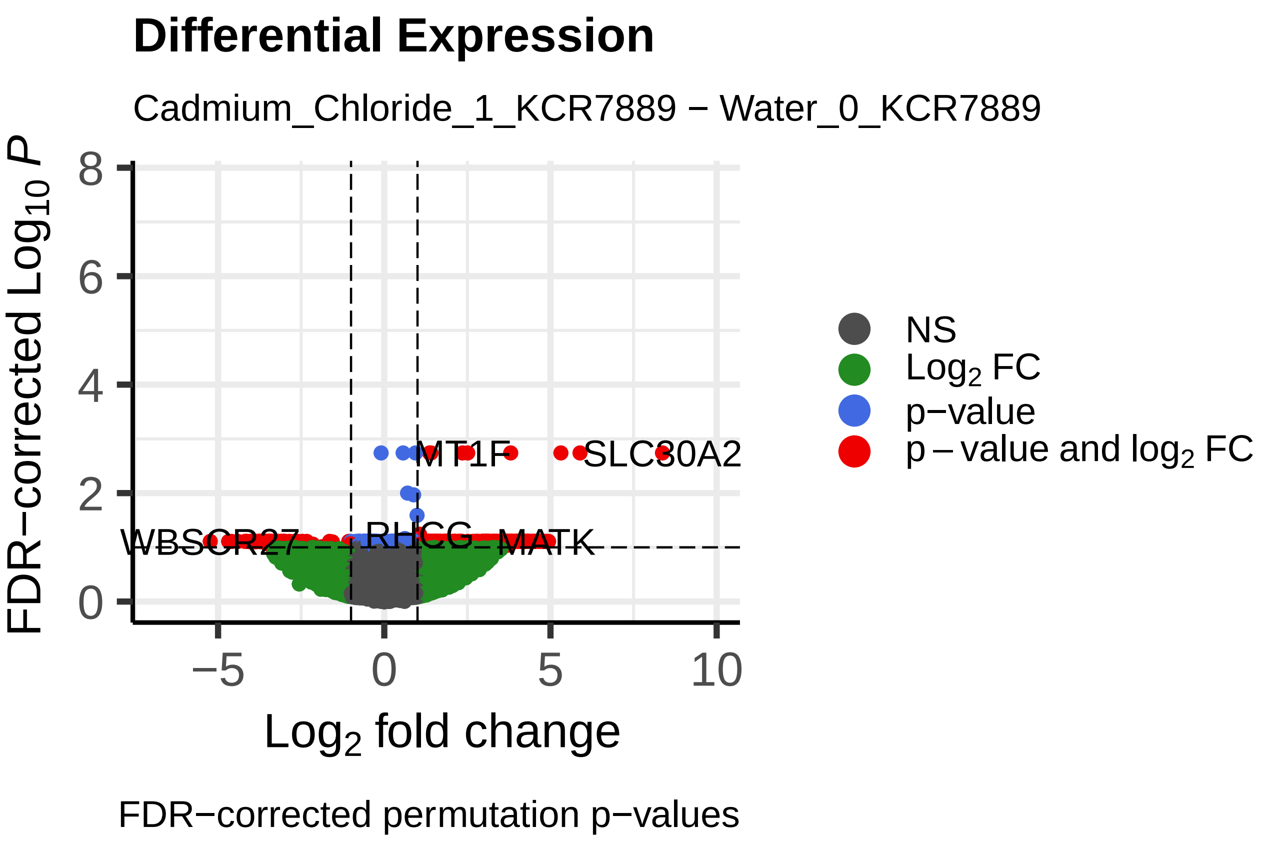
Differential Expression
Cadmium_Chloride_1_KCR7889 − Water_0_KCR7889
FDR−corrected Log10 P
Log2 fold change
MT1F
SLC30A2
WBSCR27
RHCG
MATK
NS
Log2 FC
p−value
p − value and log2 FC
FDR−corrected permutation p−values

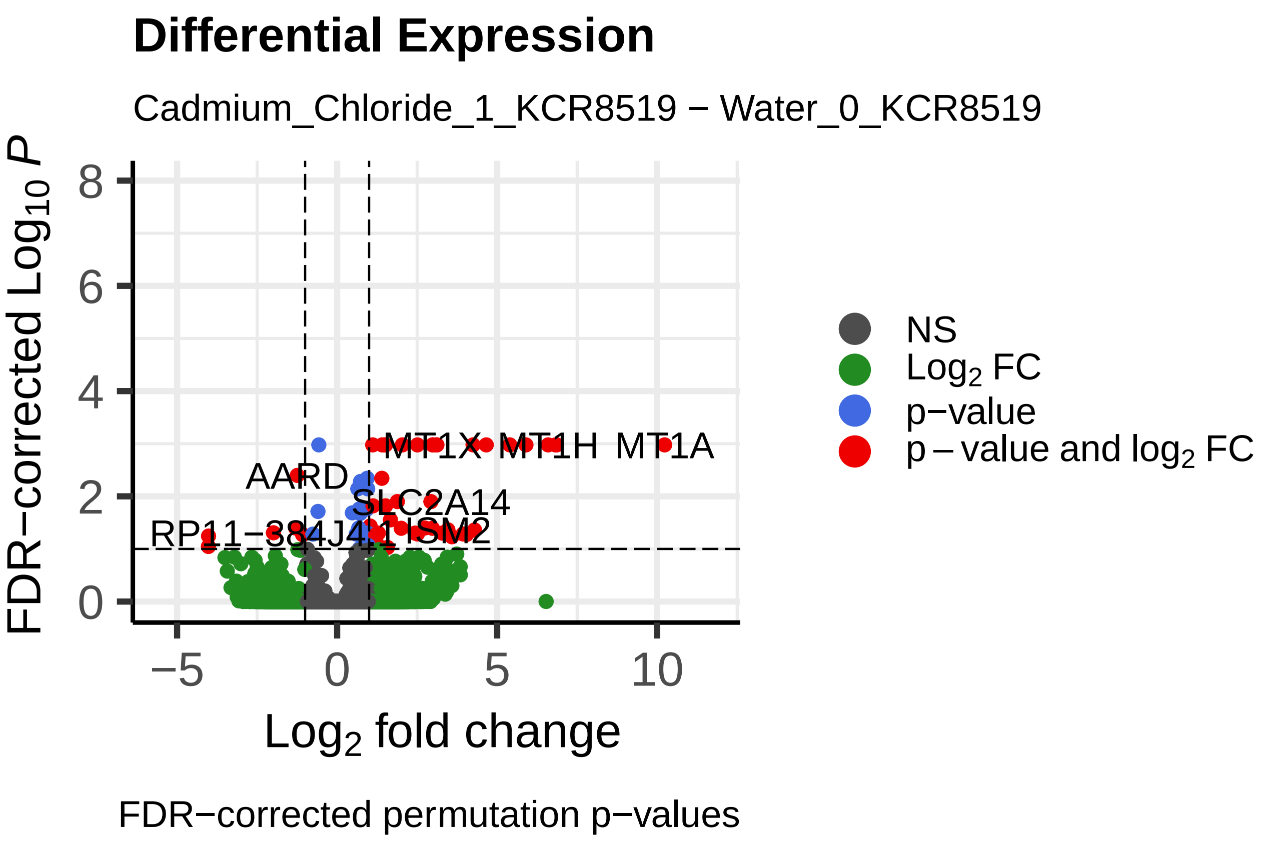
Differential Expression
Cadmium_Chloride_1_KCR8519 − Water_0_KCR8519
FDR−corrected Log10 P
Log2 fold change
MT1X
MT1H
MT1A
AARD
SLC2A14
NS
Log2 FC
p−value
p − value and log2 FC
FDR−corrected permutation p−values

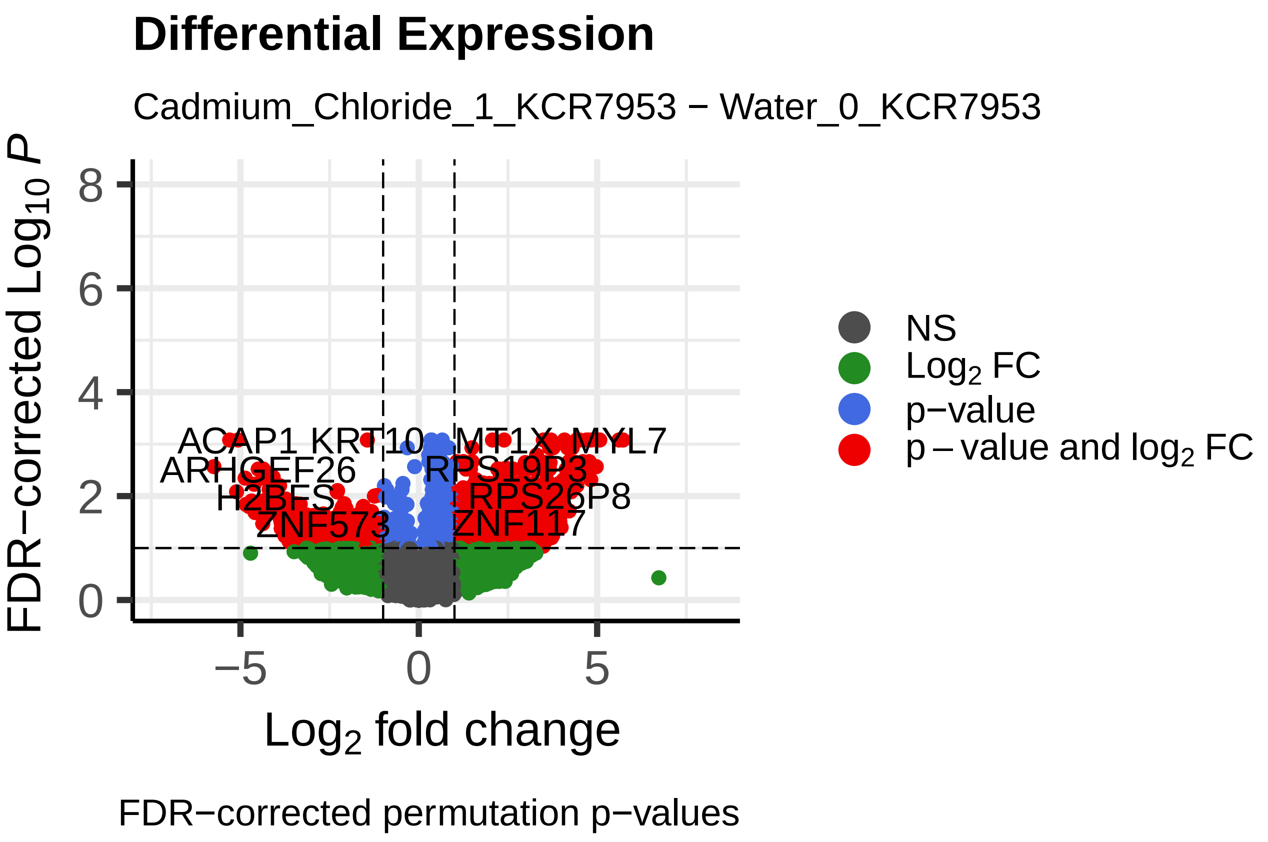
Differential Expression
Cadmium_Chloride_1_KCR7953 − Water_0_KCR7953
FDR−corrected Log10 P
Log2 fold change
ACAP1
MT1X
MYL7
RPS19P3
RPS26P8
ZNF573
ZNF117
NS
Log2 FC
p−value
p − value and log2 FC
FDR−corrected permutation p−values

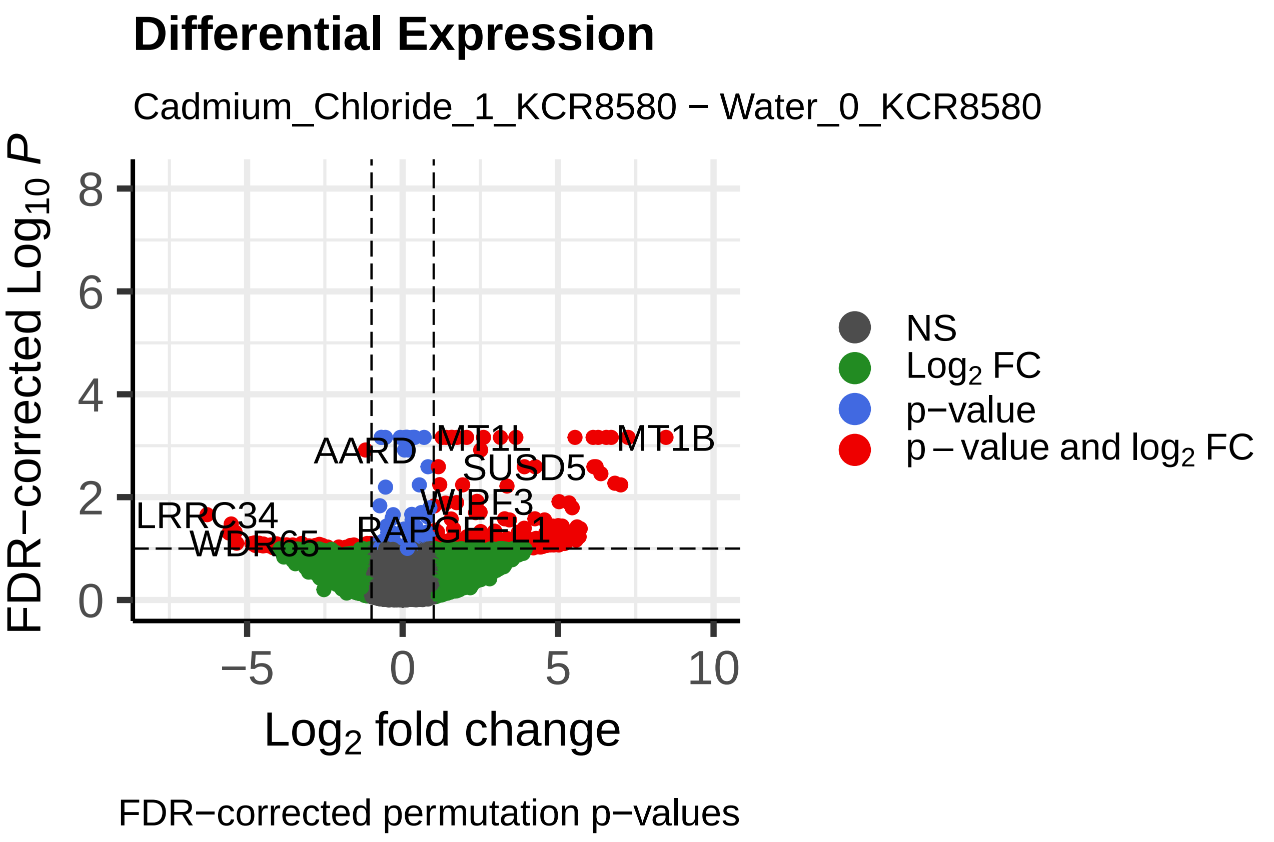
Differential Expression
Cadmium_Chloride_1_KCR8580 − Water_0_KCR8580
FDR−corrected Log10 P
Log2 fold change
MT1B
AARD
SUSD5
LRRC34
WDR65
NS
Log2 FC
p−value
p − value and log2 FC
FDR−corrected permutation p−values

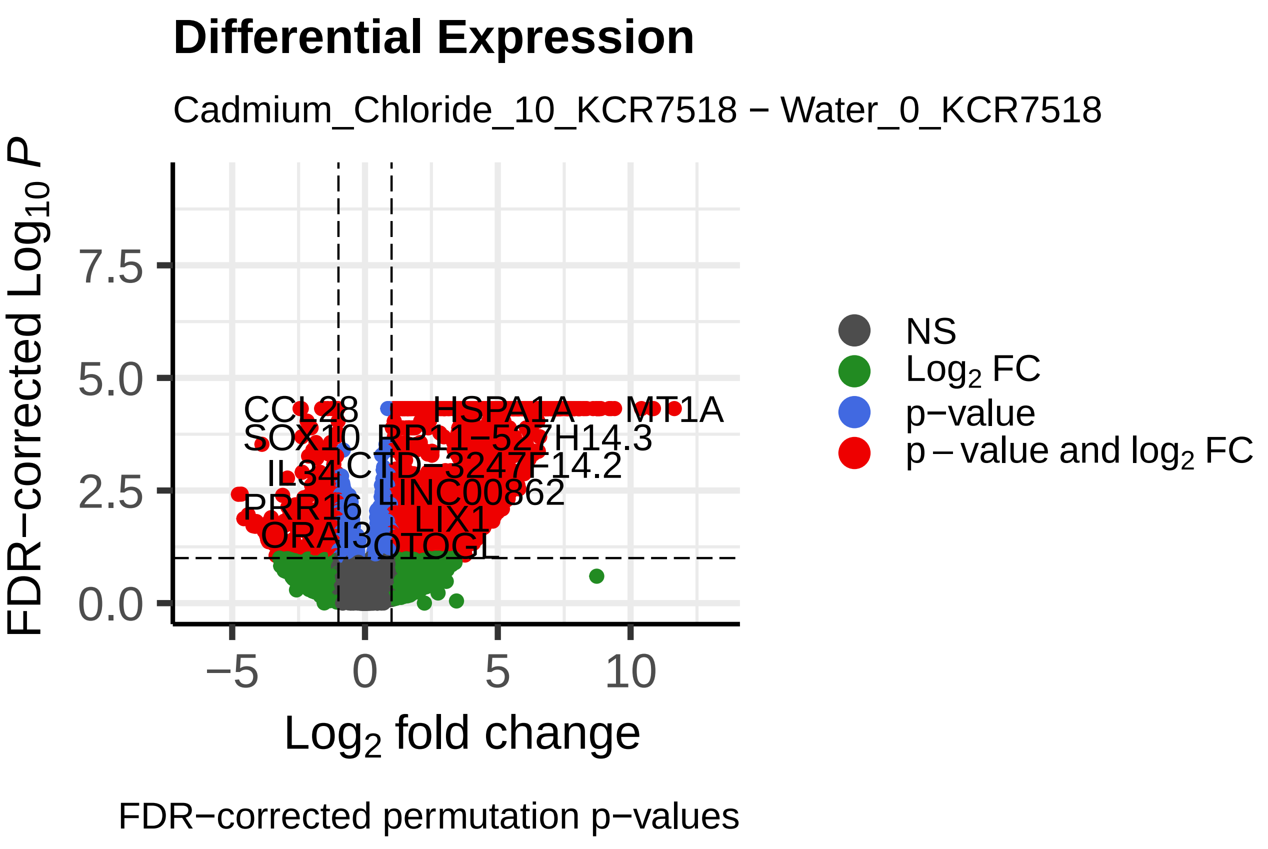
Differential Expression
Cadmium_Chloride_10_KCR7518 − Water_0_KCR7518
CCL28
SOX10
IL34
PRR16
ORAI3
HSPA1A
MT1A
RP11−527H14.3
CTD−3247F14.2
LINC00862
LIX1
OTOGL
NS
Log2 FC
p−value
p − value and log2 FC
FDR−corrected Log10 P
Log2 fold change
FDR−corrected permutation p−values

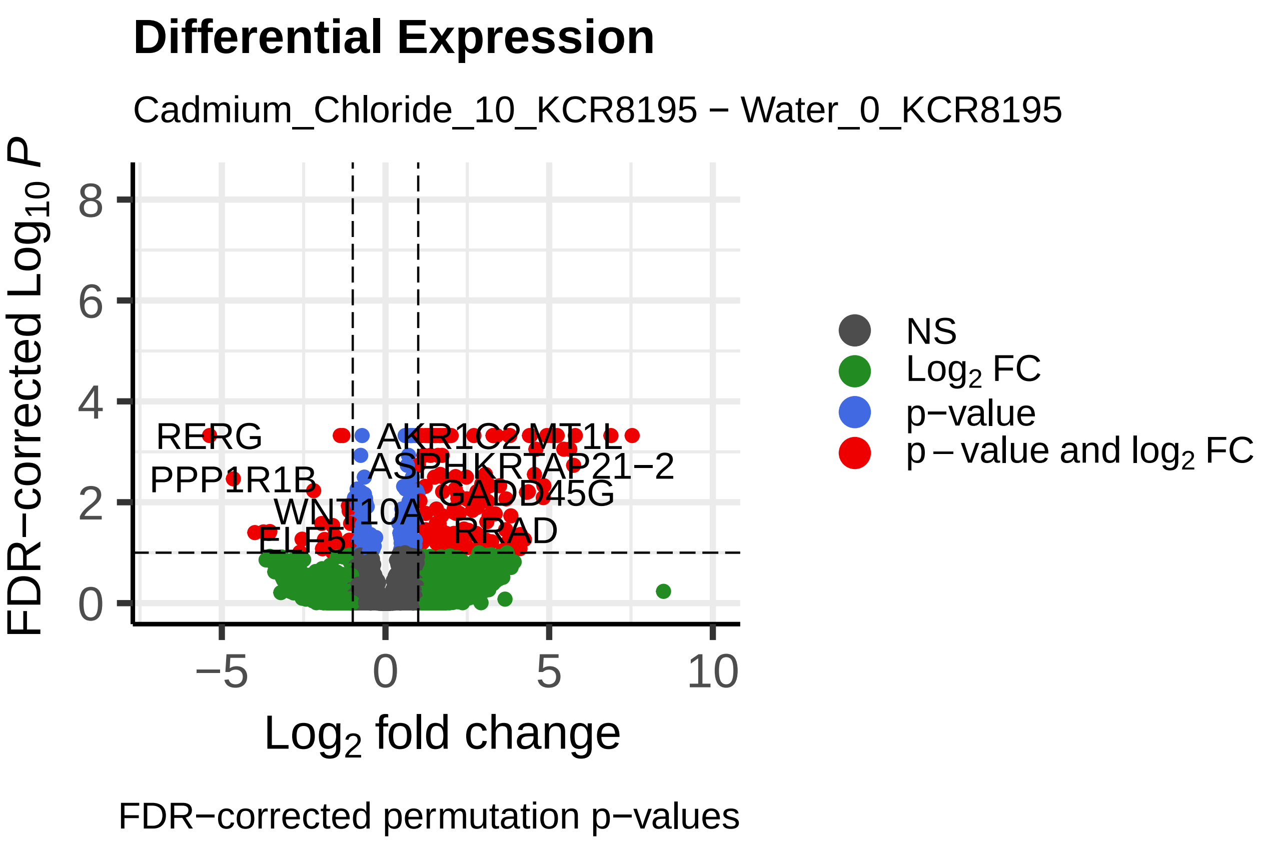
Differential Expression
Cadmium_Chloride_10_KCR8195 − Water_0_KCR8195
RERG
PPP1R1B
WNT10A
ELF5
AKR1C2
MT1L
ASPHKRTAP21−2
GADD45G
RRAD
NS
Log2 FC
p−value
p − value and log2 FC
FDR−corrected Log10 P
Log2 fold change
FDR−corrected permutation p−values

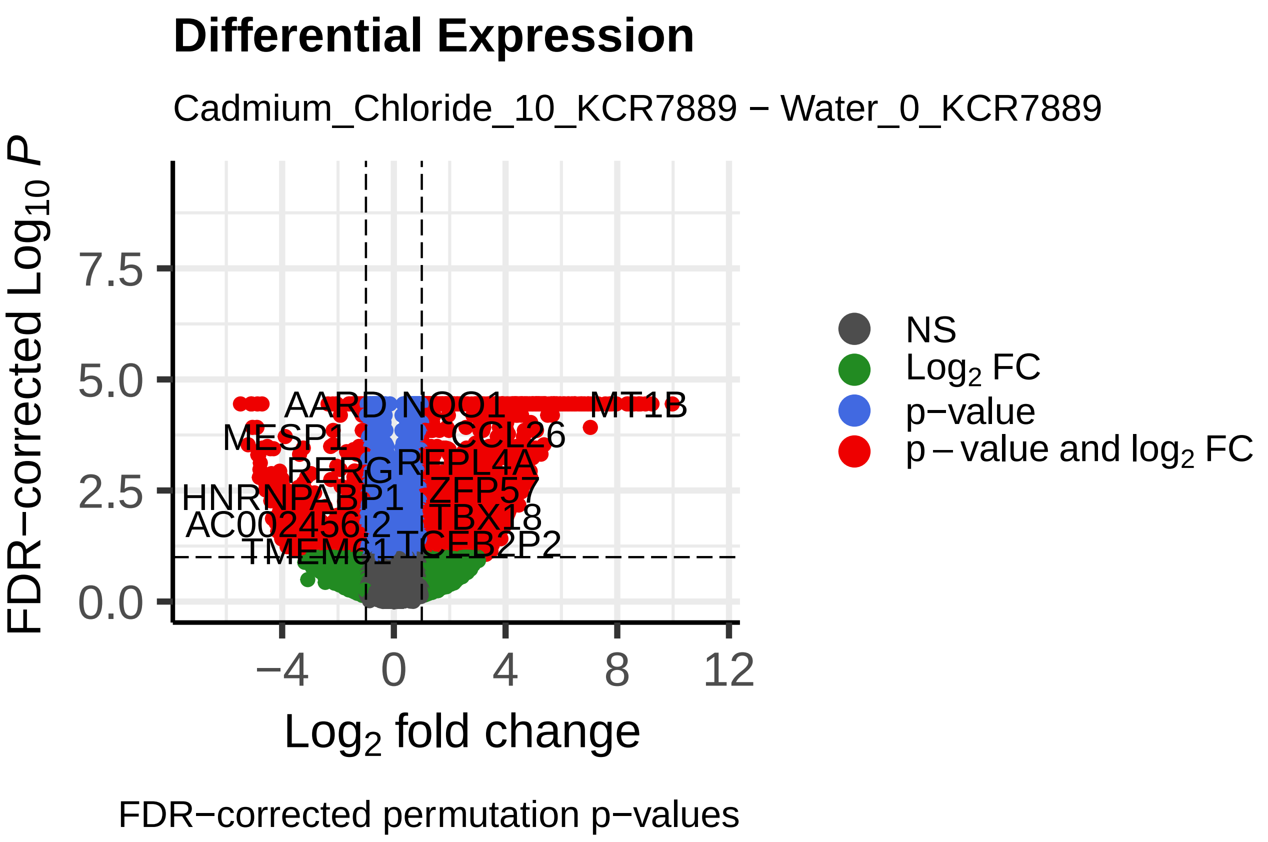
Differential Expression
Cadmium_Chloride_10_KCR7889 − Water_0_KCR7889
AARD
NQO1
MT1B
MESP1
CCL26
RERG
RFPL4A
HNRNPABP1
ZFP57
AC002456.2
TBX18
TMEM61
TCEB2P2
NS
Log2 FC
p−value
p − value and log2 FC
FDR−corrected Log10 P
Log2 fold change
FDR−corrected permutation p−values

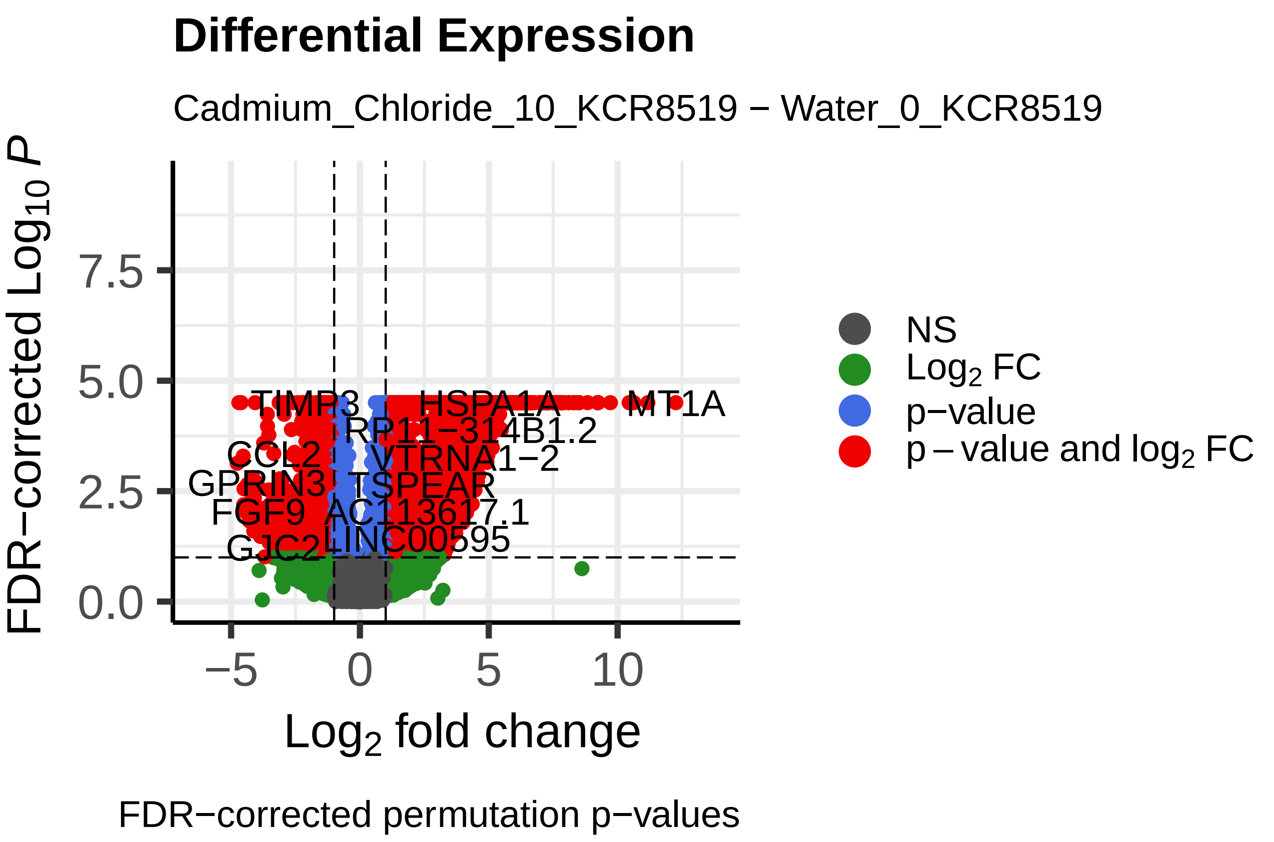
Differential Expression
Cadmium_Chloride_10_KCR8519 − Water_0_KCR8519
TIMP3
HSPA1A
MT1A
RP11−314B1.2
CCL2
VTRNA1−2
GPRIN3
TSPEAR
FGF9
AC113617.1
GJC2
LINC00595
NS
Log2 FC
p−value
p − value and log2 FC
FDR−corrected Log10 P
Log2 fold change
FDR−corrected permutation p−values

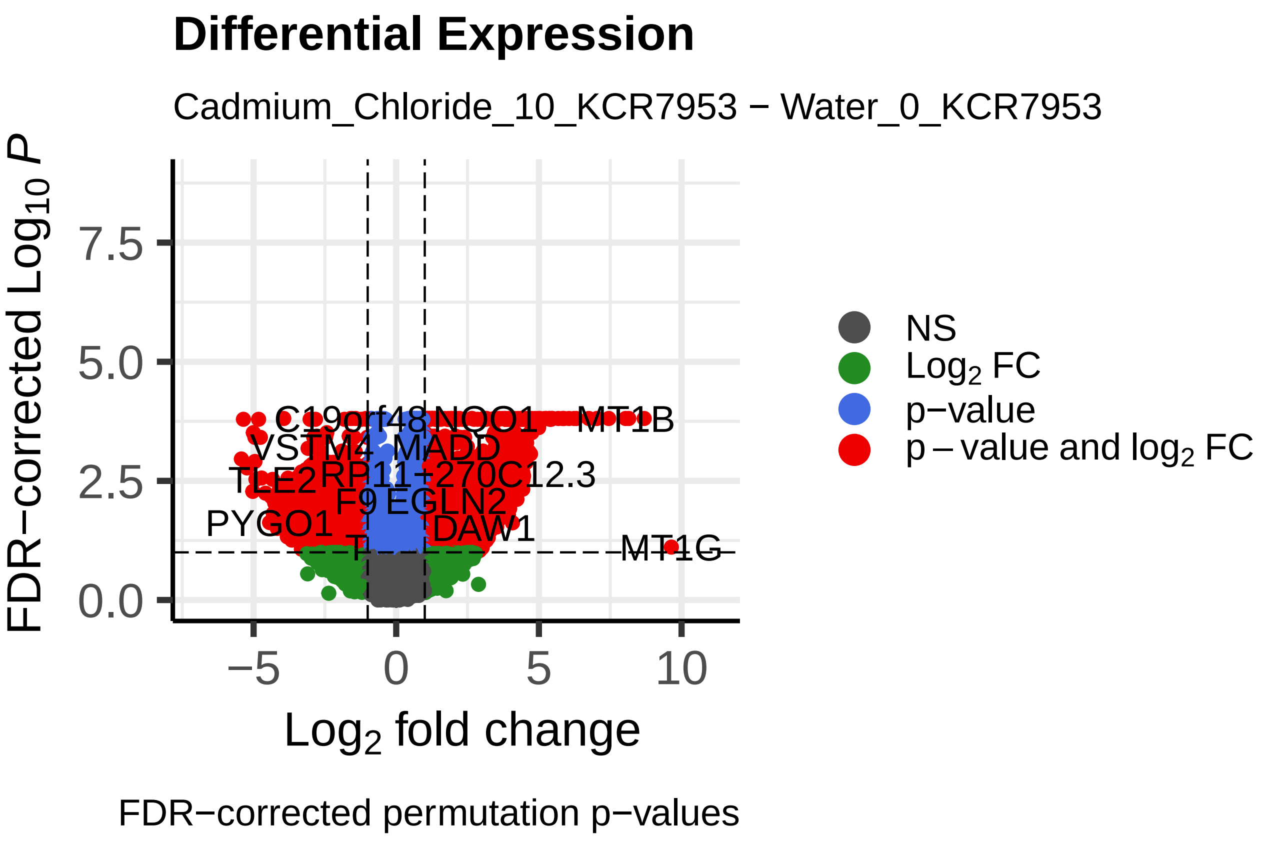
Differential Expression
Cadmium_Chloride_10_KCR7953 − Water_0_KCR7953
C19orf48
NQO1
MT1B
VSTM4
MADD
TLE2
RP11−270C12.3
F9
EGLN2
PYGO1
T
DAW1
MT1G
NS
Log2 FC
p−value
p − value and log2 FC
FDR−corrected Log10 P
Log2 fold change
FDR−corrected permutation p−values

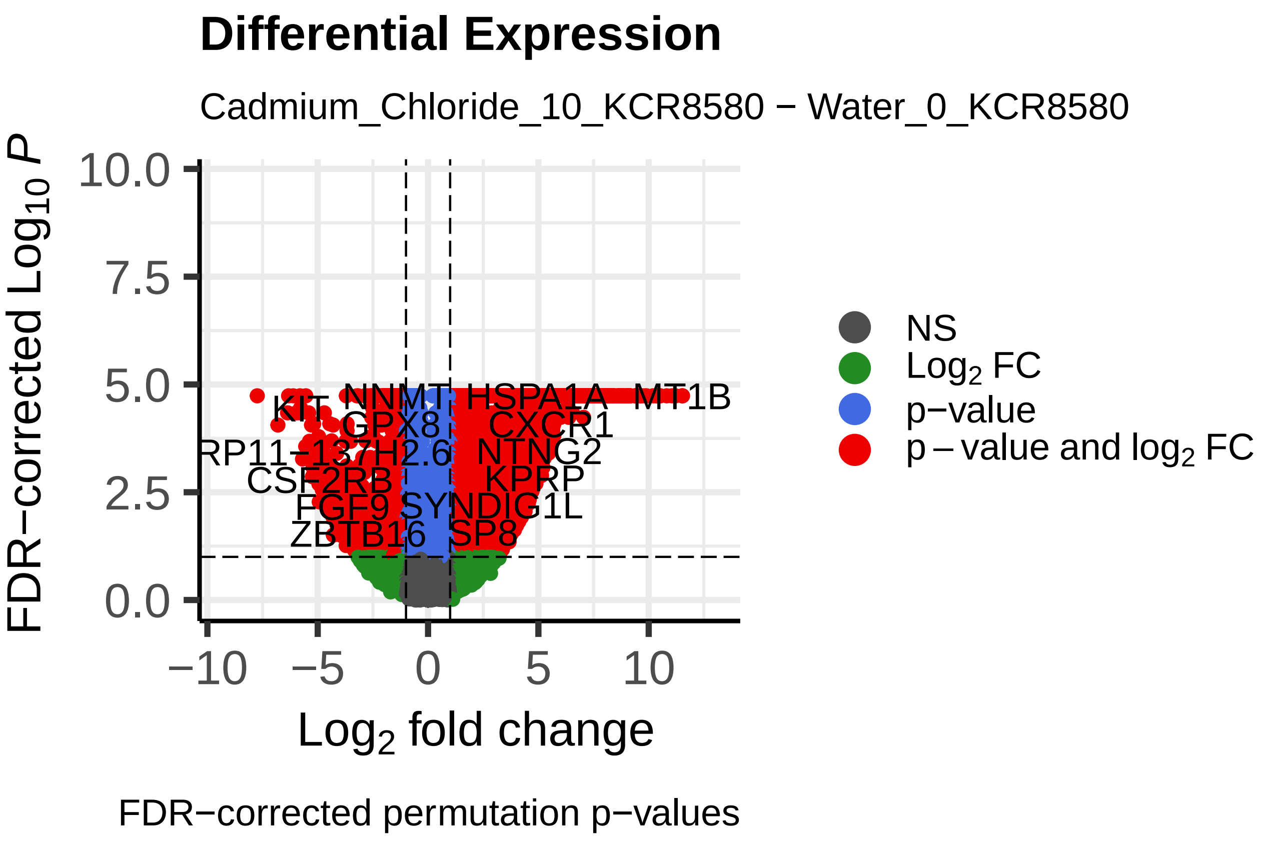
Differential Expression
Cadmium_Chloride_10_KCR8580 − Water_0_KCR8580
KIT
NNMT
HSPA1A
MT1B
GPX8
CXCR1
RP11−137H2.6
NTNG2
CSF2RB
KPRP
FGF9
SYNDIG1L
ZBTB16
SP8
NS
Log2 FC
p−value
p − value and log2 FC
FDR−corrected Log10 P
Log2 fold change
FDR−corrected permutation p−values
