## Supplementary material for "Exploring the Influence of Chemical Exposures in Breast Cancer Disparities: High-Throughput Transcriptomic Analysis in Normal Breast Cells from Diverse Donors": Table S-5

| **Sample** | **Chemical** | **0.1_vs_Ctrl_Down** | **0.1_vs_Ctrl_Up** | **0.1_vs_Ctrl_Total** | **1_vs_Ctrl_Down** | **1_vs_Ctrl_Up** | **1_vs_Ctrl_Total** | **10_vs_Ctrl_Down** | **10_vs_Ctrl_Up** | **10_vs_Ctrl_Total** |
| --- | --- | --- | --- | --- | --- | --- | --- | --- | --- | --- |
| KCR7518 | BPA | 681 | 786 | 1,467 | 196 | 384 | 580 | 1 | 21 | 22 |
| KCR7889 | BPA | 30 | 142 | 172 | 54 | 391 | 445 | 32 | 131 | 163 |
| KCR7953 | BPA | 74 | 334 | 408 | 29 | 93 | 122 | 27 | 39 | 66 |
| KCR8195 | BPA | 3 | 42 | 45 | 4 | 44 | 48 | 3 | 48 | 51 |
| KCR8519 | BPA | 0 | 0 | 0 | 0 | 0 | 0 | 0 | 7 | 7 |
| KCR8580 | BPA | 400 | 921 | 1,321 | 1 | 0 | 1 | 666 | 1,672 | 2,338 |
| KCR7518 | BPS | 0 | 3 | 3 | 0 | 5 | 5 | 0 | 0 | 0 |
| KCR7889 | BPS | 16 | 96 | 112 | 3 | 2 | 5 | 315 | 1,436 | 1,751 |
| KCR7953 | BPS | 121 | 255 | 376 | 13 | 25 | 38 | 39 | 200 | 239 |
| KCR8195 | BPS | 0 | 9 | 9 | 0 | 11 | 11 | 0 | 41 | 41 |
| KCR8519 | BPS | 0 | 0 | 0 | 0 | 0 | 0 | 0 | 0 | 0 |
| KCR8580 | BPS | 563 | 1,488 | 2,051 | 0 | 0 | 0 | 608 | 1,538 | 2,146 |
| KCR7518 | DDE | 13 | 26 | 39 | 4 | 45 | 49 | 0 | 6 | 6 |
| KCR7889 | DDE | 0 | 0 | 0 | 0 | 0 | 0 | 961 | 4,632 | 5,593 |
| KCR7953 | DDE | 0 | 2 | 2 | 1 | 2 | 3 | 17 | 10 | 27 |
| KCR8195 | DDE | 5 | 63 | 68 | 1 | 44 | 45 | 10 | 82 | 92 |
| KCR8519 | DDE | 0 | 0 | 0 | 0 | 1 | 1 | 0 | 0 | 0 |
| KCR8580 | DDE | 79 | 61 | 140 | 1 | 0 | 1 | 0 | 0 | 0 |
| KCR7518 | PFNA | 0 | 4 | 4 | 0 | 6 | 6 | 0 | 9 | 9 |
| KCR7889 | PFNA | 1 | 9 | 10 | 22 | 150 | 172 | 15 | 222 | 237 |
| KCR7953 | PFNA | 5 | 3 | 8 | 50 | 83 | 133 | 44 | 50 | 94 |
| KCR8195 | PFNA | 2 | 37 | 39 | 1 | 27 | 28 | 2 | 33 | 35 |
| KCR8519 | PFNA | 0 | 0 | 0 | 1 | 3 | 4 | 0 | 2 | 2 |
| KCR8580 | PFNA | 634 | 1,359 | 1,993 | 17 | 31 | 48 | 326 | 774 | 1,100 |
| KCR7518 | Sodium  Arsenite | 1 | 3 | 4 | 56 | 108 | 164 | 1,399 | 1,374 | 2,773 |
| KCR7889 | Sodium  Arsenite | 589 | 4,625 | 5,214 | 516 | 4,912 | 5,428 | 1,415 | 8,144 | 9,559 |
| KCR7953 | Sodium  Arsenite | 7,434 | 3,996 | 11,430 | 7,513 | 7,269 | 14,782 | 6,066 | 9,135 | 15,201 |
| KCR8195 | Sodium  Arsenite | 0 | 1 | 1 | 404 | 1,446 | 1,850 | 1,834 | 7,370 | 9,204 |
| KCR8519 | Sodium  Arsenite | 0 | 0 | 0 | 34 | 57 | 91 | 1,143 | 1,343 | 2,486 |
| KCR8580 | Sodium  Arsenite | 217 | 408 | 625 | 647 | 1,063 | 1,710 | 1,559 | 3,606 | 5,165 |
| KCR7518 | Lead  Acetate | 0 | 0 | 0 | 10 | 25 | 35 | 1 | 33 | 34 |
| KCR7889 | Lead  Acetate | 437 | 4,759 | 5,196 | 737 | 5,192 | 5,929 | 442 | 4,328 | 4,770 |
| KCR7953 | Lead  Acetate | 5,008 | 5,055 | 10,063 | 996 | 2,816 | 3,812 | 1,837 | 3,761 | 5,598 |
| KCR8195 | Lead  Acetate | 1 | 3 | 4 | 1 | 7 | 8 | 9 | 26 | 35 |
| KCR8519 | Lead  Acetate | 0 | 1 | 1 | 5 | 30 | 35 | 12 | 82 | 94 |
| KCR8580 | Lead  Acetate | 5 | 8 | 13 | 250 | 667 | 917 | 259 | 622 | 881 |
| KCR7518 | Copper  Chloride | 0 | 0 | 0 | 0 | 0 | 0 | 0 | 0 | 0 |
| KCR7889 | Copper  Chloride | 155 | 1,890 | 2,045 | 485 | 3,614 | 4,099 | 474 | 4,656 | 5,130 |
| KCR7953 | Copper  Chloride | 1,777 | 2,808 | 4,585 | 6,942 | 3,220 | 10,162 | 7,339 | 4,912 | 12,251 |
| KCR8195 | Copper  Chloride | 1 | 0 | 1 | 1 | 0 | 1 | 6 | 18 | 24 |
| KCR8519 | Copper  Chloride | 0 | 0 | 0 | 0 | 0 | 0 | 0 | 0 | 0 |
| KCR8580 | Copper  Chloride | 2 | 3 | 5 | 84 | 294 | 378 | 95 | 297 | 392 |
| KCR7518 | Cadmium  Chloride | 0 | 0 | 0 | 2 | 27 | 29 | 1,161 | 2,511 | 3,672 |
| KCR7889 | Cadmium  Chloride | 357 | 2,540 | 2,897 | 537 | 2,196 | 2,733 | 1,745 | 3,650 | 5,395 |
| KCR7953 | Cadmium  Chloride | 340 | 3,134 | 3,474 | 374 | 1,468 | 1,842 | 4,612 | 3,284 | 7,896 |
| KCR8195 | Cadmium  Chloride | 0 | 1 | 1 | 0 | 2 | 2 | 67 | 195 | 262 |
| KCR8519 | Cadmium  Chloride | 0 | 0 | 0 | 9 | 59 | 68 | 1,812 | 2,299 | 4,111 |
| KCR8580 | Cadmium  Chloride | 30 | 40 | 70 | 309 | 871 | 1,180 | 3,259 | 4,735 | 7,994 |
